# GRable version 1.0: A software tool for site-specific glycoform analysis with improved MS1-based glycopeptide detection with parallel clustering and confidence evaluation with MS2 information

**DOI:** 10.1101/2023.10.30.564073

**Authors:** Chiaki Nagai-Okatani, Daisuke Tominaga, Azusa Tomioka, Hiroaki Sakaue, Norio Goda, Shigeru Ko, Atsushi Kuno, Hiroyuki Kaji

**Affiliations:** Molecular and Cellular Glycoproteomics Research Group, Cellular and Molecular Biotechnology Research Institute, National Institute of Advanced Industrial Science and Technology (AIST), Tsukuba, Ibaraki 305-8565, Japan; Department of Systems Medicine, Keio University School of Medicine, Shinjuku, Tokyo 160-8582, Japan; Institute for Glyco-core Research (iGCORE), Nagoya University, Furo-cho, Chikusa, Nagoya, Aichi 464-8601, Japan; Education and Research Center for Pharmacy, Meiji Pharmaceutical University, Kiyose, Tokyo 204-8588, Japan; Center for Preventive Medicine, Keio University, Shinjuku, Tokyo 160-8582, Japan

**Keywords:** glycoproteomics, site-specific glycoform, glycan heterogeneity, retention time, software

## Abstract

High-throughput intact glycopeptide analysis is crucial for elucidating the physiological and pathological status of the glycans attached to each glycoprotein. Mass spectrometry-based glycoproteomic methods are challenging because of the diversity and heterogeneity of glycan structures. Therefore, we have developed an MS1-based site-specific glycoform analysis method named “Glycan heterogeneity-based Relational IDentification of Glycopeptide signals on Elution profile (Glyco-RIDGE)” for a more comprehensive analysis. This method detects glycopeptide signals as a cluster based on the mass and chromatographic properties of glycopeptides and then searches for each combination of core peptides and glycan compositions by matching their mass and retention time differences. Here we developed a novel browser-based software named GRable for semi-automated Glyco-RIDGE analysis with significant improvements in glycopeptide detection algorithms, including “parallel clustering.” This unique function improved the comprehensiveness of glycopeptide detection and allowed the analysis to focus on specific glycan structures, such as pauci-mannose. The other notable improvement is evaluating the “confidence level” of the GRable results, especially using MS2 information. This function facilitated reduced misassignment of the core peptide and glycan composition and improved the interpretation of the results. Additional improved points are: “correction function” for accurate monoisotopic peak picking; one-to-one correspondence of clusters and core peptides even for multiply sialylated glycopeptides; and “inter-cluster analysis” function for understanding the reason for detected but unmatched clusters. The significance of these improvements was demonstrated using purified and crude glycoprotein samples, showing that GRable allowed site-specific glycoform analysis of intact sialylated glycoproteins on a large scale and in depth. Therefore, this software will help to analyze the status and changes in glycans to obtain biological and clinical insights into protein glycosylation by complementing the comprehensiveness of MS2-based glycoproteomics. GRable can run freely online using a web browser via the GlyCosmos Portal (https://glycosmos.org/grable).

**Graphical Abstract:** 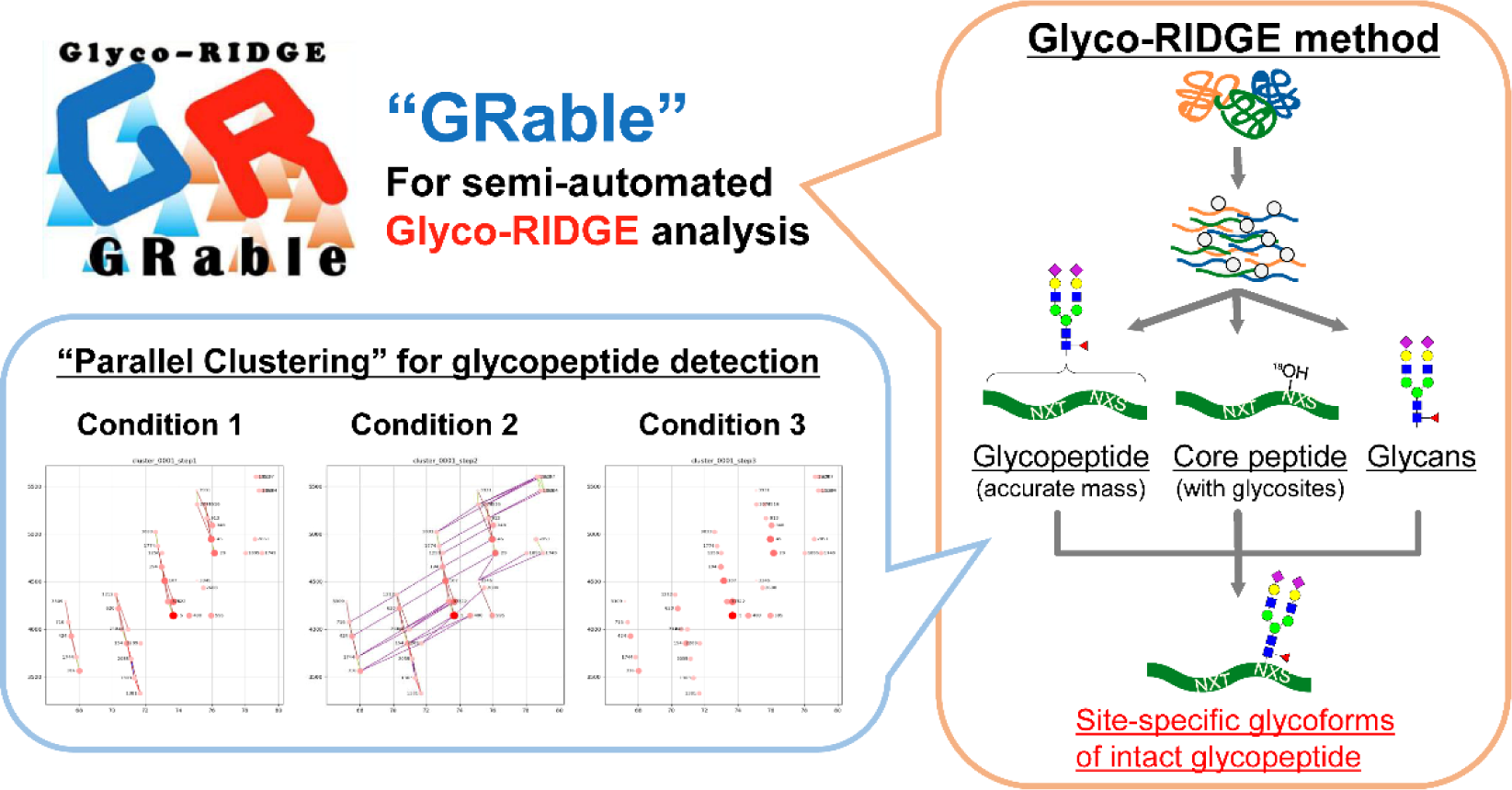

## INTRODUCTION

Protein glycosylation is a common and complex post-translational modification in eukaryotes (1) that plays pivotal roles in various biological processes (2). This post-translational modification regulates the glycoprotein function and localization by modulating their tertiary structures and interactions with other molecules (2). However, revealing the glycan structure-function relationship is challenging because glycan structures are highly diverse and heterogeneous and differ in organisms, cells, proteins, and the attached sites (3). Furthermore, the post-translational modification alters the cell state change through differentiation, carcinogenesis, infection, nutrition, medicine, and stimuli (3). Glycans are naturally heterogeneous, and elucidating their function of glycans by uncovering the state and changes in glycans with heterogeneities at each level is important. Heterogeneous levels include the micro (glycan variation of one glycosite), macro (with or without glycosylation of one glycosite), and meta (glycan variation of the entire glycoprotein molecule) levels, which are important features responsible for the presence of different proteoforms (4).

Although current improvements in mass spectrometry (MS) have made it possible to perform an in-depth structural analysis of glycopeptide peptide backbones and glycomes, elucidating the glycan compositions of specific sites for proteins with multiple glycosylated sites (i.e., micro-heterogeneity) is challenging (5). First, the glycan compositions of each site must be determined to estimate the site-specific glycoforms. For this purpose, analysis of only glycomes and deglycosylated peptides is insufficient; thus, intact glycopeptides must be analyzed directly. Current standard glycoproteomic approaches rely on MS2 spectra to identify site-specific glycoforms (6). For MS2-based glycoproteomics, the commercial software Byonic (7) is often used (6) and many in-house software programs have been developed (8–20). However, these MS2-based methods have a critical problem in that they require glycopeptide fragmentation, leading to lower detection sensitivity than the corresponding naked peptides. The decreased detection sensitivity results from the lower ionization efficiency of glycopeptides, which is notable, especially for sialylated glycopeptides because of their negative properties. In addition, glycan heterogeneity decreases the abundance of each glycopeptide. Therefore, in some MS1 spectra, many signals are not selected for MS2 analysis, and frequently, MS2 spectra do not contain sufficient information for confident glycopeptide identification.

To overcome this limitation, we developed an MS1-based glycoproteomic approach named “Glycan heterogeneity-based Relational IDentification of Glycopeptide signals on Elution profile” (Glyco-RIDGE) (21), which was first introduced in 2015 (22). One limitation of MS1-based approaches is to “estimate” combinations of core peptides and glycans, having a possibility of misassignment because of isobaric combination issues caused by adducts and core peptide modifications observed even in MS2-based approaches (23, 24). To minimize such errors, a general workflow of the Glyco-RIDGE analysis using liquid chromatography/mass spectrometry (LC/MS) data of glycopeptide samples requires information on core peptides present in the analyte, as well as presumed glycans (Figure 1); these sequences and *N-*glycosites can be identified by isotope-coded glycosylation site-specific tagging (IGOT)-LC/MS/MS (25, 26), in which N-glycosites are specifically labeled by peptide-*N*-glycosidase F (PNGase F)-mediated incorporation of a stable isotope tag (^18^O). In the Glyco-RIDGE workflow, MS1 signals derived from glycopeptides with the same core peptide are assigned as a cluster in the LC/MS data based on their mass differences and closeness of retention times (RTs). Then, the combination of the core peptide and glycan composition for these glycopeptides is annotated by matching the observed accurate glycopeptide mass with the sum of the masses of the pre-identified core peptide candidates and presumed glycans. A notable feature of Glyco-RIDGE is the detection of glycopeptide signals based only on MS1 data, in contrast to other MS1-based approaches that employ MS2 data to identify at least one glycopeptide for each core peptide (27–31). With this feature, the Glyco-RIDGE method is expected to achieve a more high-sensitive and comprehensive analysis of protein glycosylation than current fragmentation-dependent identification methods. Notably, a previous study (32) has demonstrated that among the MS2 spectra assigned using the Glyco-RIDGE method and Byonic search, the ratio of the same assignment was quite high (349/353 = 98.9%), demonstrating that the reliability of this method is similar to that of Byonic.

**Figure 1.**
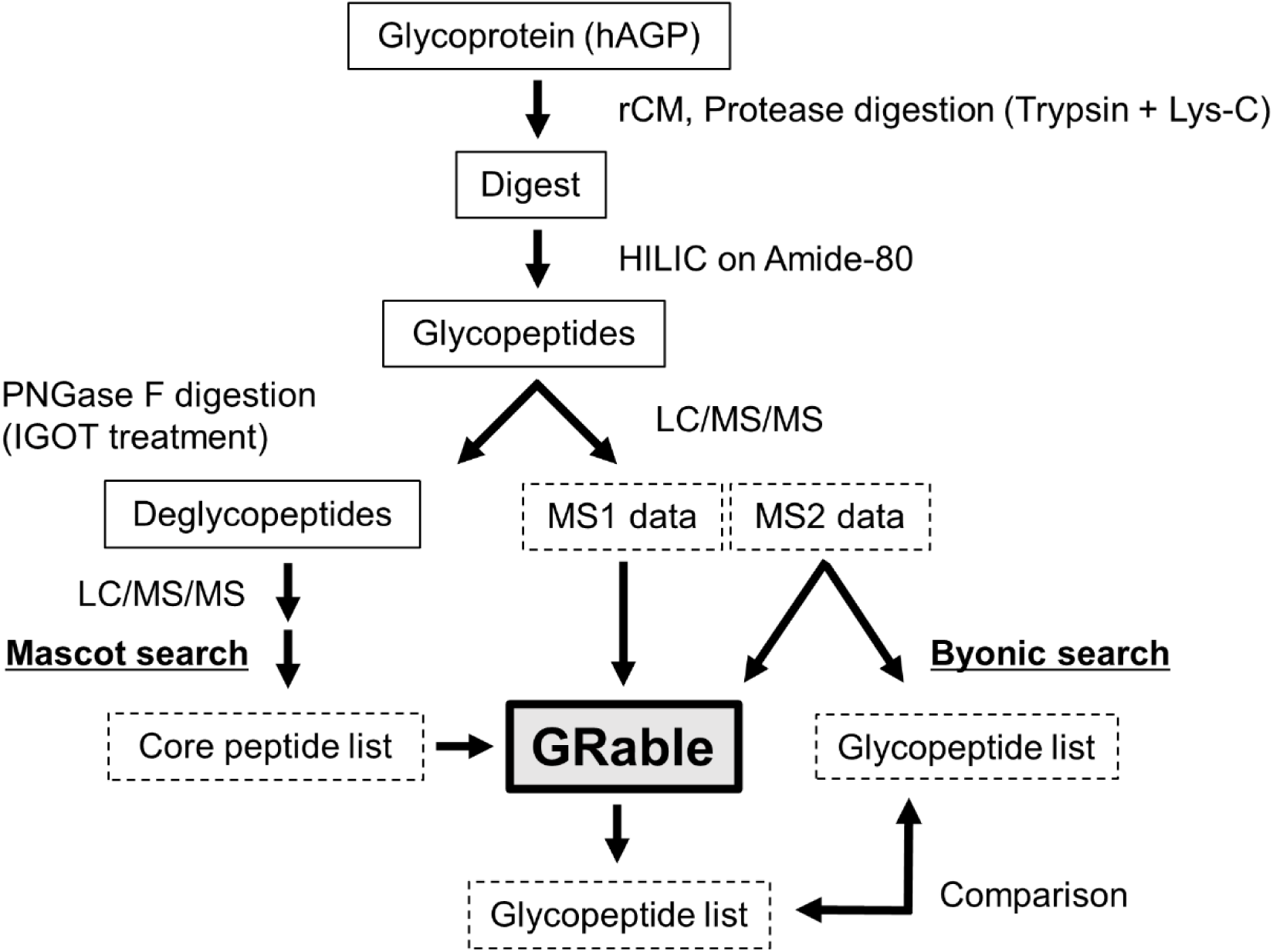
Workflow of glycoproteomic analysis for hAGP using GRable. hAGP was subjected to reduction and carbamidomethylation (rCM) using dithiothreitol and iodoacetamide, respectively, and digested with trypsin and Lys-C endopeptidase. The digest was used in hydrophilic interaction chromatography (HILIC) on an Amide-80 column to capture glycopeptides. A small aliquot of the glycopeptide fraction was treated with peptide-*N*-glycosidase F (PNGase F) in ^18^O-labeled water to remove *N*-glycans and to label deglycosylated Asn as ^18^O-labeled Asp (isotope-coded glycosylation site-specific tagging; IGOT) (25, 26). The IGOT-treated deglycopeptides were analyzed using liquid chromatography-tandem mass spectrometry (LC/MS/MS), and the MS2 data were used for database search using Mascot to prepare a core peptide list. In parallel, another aliquot of the glycopeptide fraction was analyzed through LC/MS/MS. MS1 data of the glycopeptide analysis underwent GRable analysis. GRable used the core peptide (Data S1) and glycan composition (Data S2) lists to assign each glycopeptide signal, whereas MS2 data are used for adding the confidence level to the GRable results. Glycopeptide analysis MS2 data was also used for glycopeptide estimation using Byonic. The resulting glycopeptide lists of GRable and Byonic analyses were compared.

The feasibility and usefulness of the Glyco-RIDGE method have been demonstrated using in-house software by applying it to the analysis of specific glycan motifs (i.e., Lewis X and polylactosamine)-containing glycopeptides in complex glycoprotein mixtures after the removal of sialic acids (22, 32). Then, this method has been used also for site-specific glycoform analysis of intact (i.e., sialylated) glycopeptides (33–36), revealing site-specific “glycostem” and “glycoleaf” features (36). However, the prototype software has several limitations. One limitation was the misfinding of monoisotopic signals from MS1 data, causing a decreased number of detected glycopeptide signals. Regarding clustering, glycopeptides with the same core peptide and different numbers of sialic acid residues were detected as discrete clusters, impairing the detection and interpretation of microheterogeneity. Another issue was the presence of glycopeptide clusters that were detected but not assigned, which might result from adducts or peptide modifications; however, the reason is unclear and thereby remains to be improved. In addition, selecting the most plausible match for each glycopeptide cluster as the final assignment and evaluating the assignments for each glycopeptide were performed manually by searching the MS2 spectra, which is time-consuming. Accordingly, it is also difficult to optimize search parameter settings and acceptance criteria while balancing the coverage and accuracy of the analysis, which is an important consideration also in MS2-based approaches (37).

The present study introduces a novel software named “GRable” to semi-automatically execute the Glyco-RIDGE analysis with some technical improvements for the above issues. The notable features are “parallel clustering” for efficient and comprehensive finding of glycopeptide clusters and adding a Selection step for selecting and evaluating assignments using MS2 information as a “confidence level.” Additional improved points are: “correction function” for accurate monoisotopic peak picking; one-to-one correspondence of clusters and core peptides even for multiply sialylated glycopeptides; and “inter-cluster analysis” function for understanding the reason for detected but unmatched clusters; To demonstrate the feasibility and utility of this software, we analyzed MS data obtained with intact glycopeptides prepared from human serum α1-acid glycoprotein (hAGP), a glyco-biomarker candidate for liver fibrosis (38), as a model of targeted analysis. In addition, we used data on glycopeptides obtained from human promyelocytic leukemia-60 (HL-60) cell lysates (32), as a crude sample model.

## EXPERIMENTAL PROCEDURES

GRable version 1.0 requires the following four files: LC/MS data of glycopeptides after deconvolution (.mzML); a core peptide list (.xlsx); a glycan point list (.xlsx); and LC/MS/MS data for glycopeptides (.mgf) (Figure 2). Details of the input files and their preparation are described below and in the user manual (Document S1). Note that this version of GRable is guaranteed only for Thermo Fisher Scientific data.

**Figure 2.**
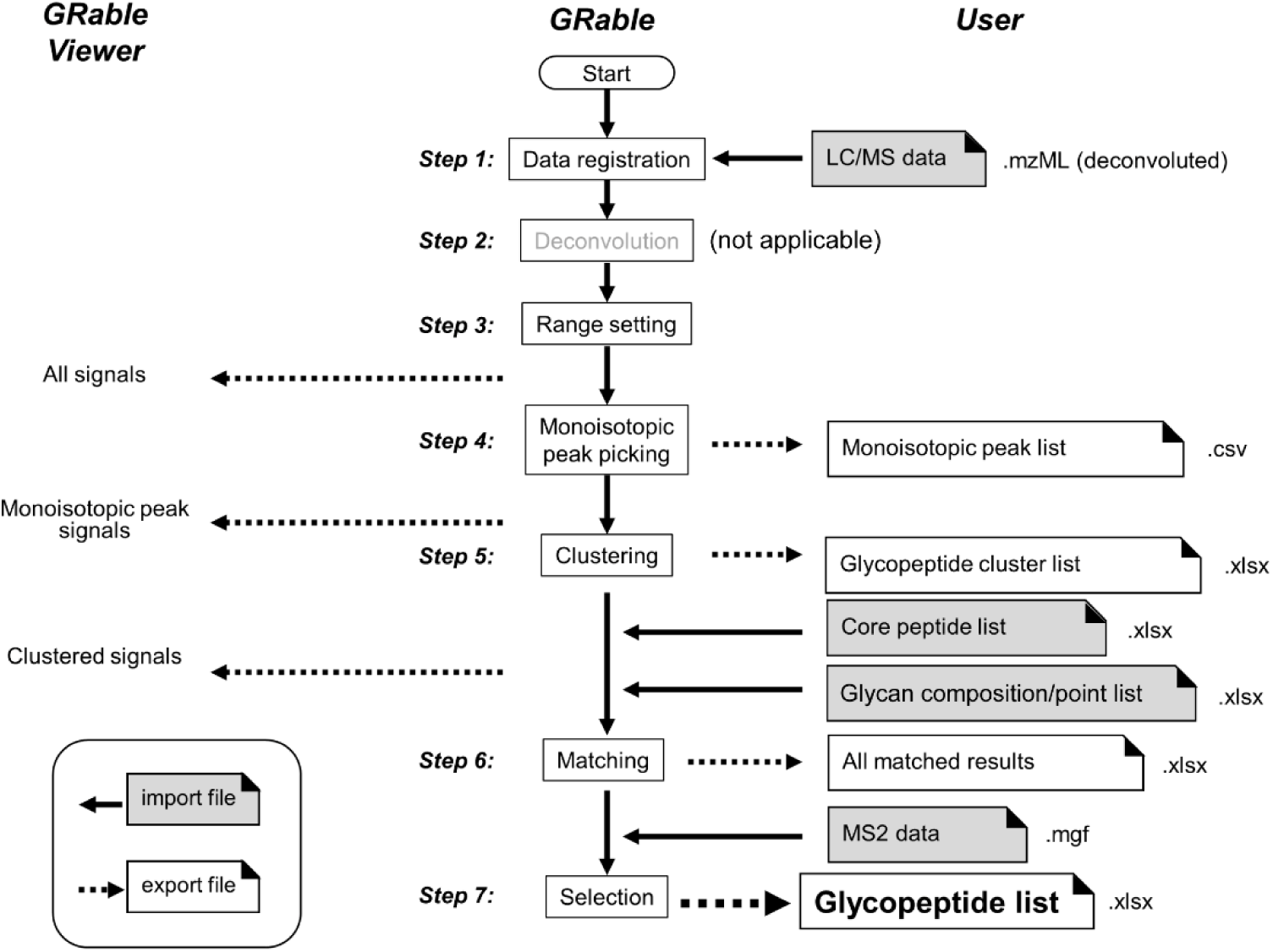
Overview of data processing by GRable. GRable proceeds in seven steps using the four files indicated in gray font. In Step 1, GRable allows the registration of the LC/MS data of glycopeptides in the mzML format, which ensures great versatility and ease of data processing. In the current version, the deconvolution function (Step 2) is not applicable although it is visible in the user interface for future implementation. Step 3 was designed to set the RT range, mass range, and minimum signal intensity (threshold) over which the analysis was performed. Step 4 was intended to find and group signals of an identical ion based on three parameters: time (scan), mass (MH+), and intensity, and to obtain the monoisotopic mass of each signal group at the peak time, using our unique algorithms. In Step 5, a series of signals from the glycopeptide group were found as a cluster based on the elution behavior and mass difference of its members. In Step 6, GRable searches for a combination of core peptides and glycan compositions that match the mass of the putative glycopeptide within the allowed mass error (user setting) according to the following equation: Observed M(glycopeptide) = calculated M(core peptide identified) + M(Hex)*i + M(HexNAc)*j + M(dHex)*k + M(NeuAc)*l (M is a mass value, and i, j, k, and l are integers). In Step 7, the most plausible combination among multiple candidate combinations suggested for one glycopeptide cluster was selected. Then, their reliability at the cluster level as well as the single glycopeptide level was evaluated using information from the results and additional MS2 information. The results of Steps 3–5 were visually confirmed using a viewer in the main window of the software. The detailed results of Steps 4–7 can be exported as an Excel file with each setting, as shown for hAGP (Data S5 and S6) and HL-60 cell lysates (Data S7 and S8).

### Preparation of hAGP glycopeptide samples

To prepare large-scale analytes for repeated verification, 5 mg of freeze-dried hAGP (Sigma-Aldrich, St.Louis, MO, USA) was dissolved in Milli-Q water (500 μL) and the aliquot was denatured with 0.1% Rapigest (Waters, Milford, MA, USA) in 50 mM Tris-HCl, reduced with dithiothreitol (Fujifilm Wako, Osaka, Japan), alkylated with iodoacetamide (Fujifilm Wako), and digested with Lys-C (1/200 weight of protein; Fujifilm Wako) and sequencing grade modified trypsin (1/100 weight of protein; Promega, Madison, WI, USA) at 37 °C overnight in 0.1 % Rapigest. After digestion, Rapigest was cleaved by acidification with 0.1% trifluoroacetic acid (TFA; Fujifilm Wako). Glycopeptides from hAGP tryptic digests were then captured using an Amide-80 column (TSK-gel Amide-80, 2 × 50 mm; TOSOH, Tokyo, Japan) equilibrated with 75% acetonitrile (Merck, Darmstadt, Germany)/0.1% TFA, and eluted isocratically with 50 % acetonitrile/0.1% TFA after washing.

### LC/MS/MS analysis of hAGP glycopeptide samples

Glycopeptides were analyzed using an LC-electrospray ionization-MS system equipped with a nanoflow LC system (UltiMate-3000; Thermo Fisher Scientific, Waltham, MA, USA) and a tandem mass spectrometer (Orbitrap Fusion Tribrid mass spectrometer; Thermo Fisher Scientific). Glycopeptides were trapped on a C18 cartridge column (Acclaim PepMap100 C18, 0.3 mm I.D. × 5 mm; Thermo Fisher Scientific) and separated on a C18 tip column (75 µm I.D. × 15 cm, 3 µm particle; Nikkyo Technos, Tokyo, Japan) using 2–36% acetonitrile/0.1% formic acid (Fujifilm Wako) gradient (90 min) at a flow rate of 300 nL/min. The ionization voltage was 2.0 kV (positive), and the temperature of the ion transfer tube was 275 °C. Data were acquired using the data-dependent mode (cycle time: 2.5 s) with internal mass calibration (lock mass) at 445.12003. The mass range was 377–2,000 m/z (MS1) and 135–2,000 m/z (MS2). The mass resolutions were 120,000 (MS1, Orbitrap) and 15,000 (MS2, Orbitrap). The fragmentation mode was high-energy collision-induced dissociation with stepped collision energies of 25, 30, and 35%. The acquired raw data file was deconvoluted using the Xtract of Proteome Discoverer (ver. 2.4, Thermo Fisher Scientific), and exported as an mzML file for GRable analysis.

### IGOT-LC/MS/MS analysis of hAGP glycopeptide samples

IGOT-LC/MS analysis was performed for core peptide identification as reported previously (25, 26). Glycopeptides were treated with PNGase F (Takara Bio, Shiga, Japan) in 50 mM Tris-HCl, pH 8.5, prepared with H_2_^18^O (isotope purity: 95%; Taiyo Nippon Sanso, Tokyo, Japan) at 37 °C overnight. Deglycosylated peptides (IGOT peptides) were analyzed before glycopeptide analysis using LC/MS/MS under the same conditions as for the glycopeptide sample.

### Preparation of a core peptide list

The format of the core peptide list required by the software was fixed and created in a manner similar to that used in this study (Data S1). The details of the preparation are documented in the manual (Document S1, Section 4). Items minimally required for preparing a core peptide list are the calculated mass, retention time of each peptide and at least one peptide identifier such as CP (core peptide) number. These information are used to detect the combinations of glycopeptides and core peptides by calculating the mass differences within the limited RT range in the Matching step. For the identification of core peptides, IGOT-LC/MS analysis, as described above, is recommended to confirm their *N*-glycosylations, but it is possible to list the peptides identified by proteomic analysis. Confirmation of the presence of core peptides for glycoproteins is crucial for crude samples. For the analysis of a highly purified glycoprotein, it is also possible to list the peptides predicted based on protease digestion patterns.

For hAGP, the raw data acquired by IGOT-LC/MS/MS were converted to mgf using Mascot Distiller (ver. 2.7.0; Matrix Science, Boston, MA, USA), and searched using Mascot (ver. 2.5.1; Matrix Science) using a human protein sequence file (SwissProt_UniProtKB_isoform; downloaded in April 2019; entry: 42,431). The search parameter settings were as follows: enzyme: trypsin (full or semi); missed cleavage: 2; mass tolerance: 7 ppm (MS1) and 0.02 Da (MS2); fixed modification: carbamidomethyl (C); variable modifications: Gln to pyro-Glu (peptide N-term, Q), ammonia-loss (peptide N-term, carbamidomethyl C), oxidation (M), and Delta: H(-1)N(-1)18O(1)(N); and target false discovery rate: 1%. Mascot search results with a peptide rank of 1 and peptide expectation value of <0.05 were selected. Matched sequences containing IGOT modifications at Asn on the consensus sequence for *N*-glycosylation (Asn-Xaa-[Ser/Thr]; Xaa is not Pro) obtained using trypsin (full) and trypsin (semi) were combined. The resulting glycopeptide list was used to create a core peptide list for GRable analysis. To ensure the comprehensive analysis, hAGP peptide sequences that were not identified in the data used here but found in other analyses were manually added to the list with observed or predicted RTs (Data S1). That is, five core peptides with predicted hAGP sequences were added to the list, and three core peptides were matched with glycopeptides as clusters.

It is noteworthy that considering oxidized Met in the deglycopeptide search for improving coverage may cause incidental mismatching for one glycopeptide; for example, oxidation of the peptide and HexNAc and Hex and carbamidomethylation of the peptide (23). However, in the Glyco-RIDGE method, such a misassignment is removed at the selection step because the core peptide of each glycopeptide is evaluated as a glycopeptide cluster. Similarly, carbamidomethylation of the Met within a core peptide, which was not considered in the peptide list preparation and has the possibility of mismatching (23), should be omitted from the selection results. It should also be noted that the Mascot search for *O*-glycosylation of IGOT-treated *N*-deglycopeptides may be useful for evaluating the possibility of misinterpretation of the GRable results of site-specific N-glycoform analysis because this software is intended to assign glycopeptides with single glycosite.

### Preparation of a glycan composition list

The format of the core peptide list required by the software was fixed and created in a manner similar to that used in this study (Data S2). The details of the preparation are documented in the manual (Document S1). In the Matching setting, users can define the permissible glycan compositions (i.e., the type and number of each glycan unit). The default settings were as follows: Hex: 0–12, HexNAc: 1–12, dHex: 0–4, and NeuAc: 0–4. The maximum numbers of Hex and HexNAc were determined using the approximate masses of the ionizable glycopeptides. The maximum number of NeuAc was limited to four because our sample preparation conditions did not allow the remaining oligo- or polysialic acids. The number of dHex (fucose) was also limited to four or fewer because the mass of dHex(5) is similar to that of Hex(2)HexNAc(2) (difference = 0.025), corresponding to approximately 5 ppm of 5,000 Da. If the measurement is performed with an accuracy greater than 2 ppm, five or more fucoses may be distinguishable. Within the range of glycan compositions, users can provide arbitrary points for likely compositions in the glycan list. In contrast, among the possible compositions, many are considered unlikely based on biosynthetic pathways, and matches with such unusual compositions can be considered incorrect. Accordingly, users can also define unusual compositions in the glycan list and such incorrect matches are marked in the match results by giving negative points, and thus, will be considered in the selection.

The format of the glycan point list was fixed (Data S2). The list used in this study can be used to analyze human cell-derived protein samples and was not considered for NeuGc. In this list, 139 glycan compositions were assigned 1 point if the composition is matched. The list contained glycan compositions in *N*-glycan major biosynthetic pathways, from the glycan produced by oligosaccharide transferase to the glycan processed by many glycosyltransferases and glycosidases. The largest is Hex(7)HexNAc(6)dHex(4)NeuAc(4), which corresponds to a tetra-antennary glycan with four Fuc and four NeuAc. The smallest is HexNAc(1), a breakdown product remaining in GlcNAc on Asn. Unusual compositions were determined by considering *N*-glycan biosynthetic pathways.

This study considered four glycan units (Hex, HexNAc, dHex, and NeuAc) as the glycan components attached to hAGP. *N*-glycan compositions were expressed as the generic monosaccharide composition as “Hex(*)HexNAc(*)dHex(*)NeuAc(*)”, where * represents the number of individual monosaccharide residues. The glycan composition list for hAGP was almost identical to the human *N*-glycome list supplied by Byonic, except for one composition (Hex(7)HexNAc(6)) (Table S1), which was manually added because this composition is considered common in the human *N*-glycome. In the glycan composition list, unusual compositions were defined based on *N*-glycan biosynthetic pathways; however, negative points were not given for these unusual compositions in this study.

### Byonic search for hAGP data

Glycopeptide MS2 data were searched using the Byonic search engine ver. 2.15.7 (Protein Metrics, Cupertino, CA, USA) using the human protein sequence file from SwissProt (downloaded on May 19, 2020; entry: 42,296) and the glycan composition list (Table S1). Search parameter settings were as follows: enzyme: Trypsin_KR (Full or Semi), max missed cleavages: 2, static modifications: carbamidomethyl (C), dynamic modifications: ammonia-loss (N-term C), Gln to pyro-Glu (N-term Q), oxidation (M), peptide mass tolerance: ± 3 ppm, fragment mass tolerance: 0.02 Da. The search results used estimated glycopeptides with Confidence = High and Byonic Score ≥ 200 to compare GRable results. The two results obtained with Trypsin_KR (full) and Trypsin_KR (semi) were combined, and the resultant glycopeptide list was compared with the GRable results.

### GRable analysis of HL-60 cell data

LC/MS/MS data for HL-60 cell lysates were obtained from a previous study (Figure S1) (32). A protease digest of the HL-60 cell lysate was prepared in a manner similar to that for hAGP. An aliquot of the digest was acidified and heated to remove sialic acid, decreasing glycan heterogeneity and increasing the relative abundance of each glycopeptide. An aliquot of the desialylated glycopeptide fraction was subjected to HILIC to collect four glycopeptide fractions, which were then analyzed using LC/MS/MS and IGOT-LC/MS/MS to obtain glycopeptide and deglycopeptide MS data, respectively. The glycans obtained from the IGOT procedure were analyzed using matrix-assisted laser desorption/ionization time-of-flight (MALDI-TOF) MS to prepare a glycan composition list. Because the HL-60 cell data were for desialylated glycopeptides, only three glycan units (Hex, HexNAc, and dHex) were considered glycan components. The definition of unusual composition was the same as that used for the hAGP. The resulting core peptide list (Data S3) and glycan point list (Data S4) for fraction 2 were used for the GRable analysis.

### Software implementation

GRable was implemented as a web application by Hitachi Solutions Technology Ltd. The user interface was mostly written in JavaScript, data management in Java, and scientific calculations in Python. PostgreSQL was used as the background for the user and data file management. The software was developed and tested on Ubuntu Linux 22.04 LTS.

## RESULTSAND DISCUSSION

GRable is a web application that users can access through web browsers. It should be noted that the deconvoluted LC/MS data must be uploaded in the current version. For data processing (Figure 2), all steps were executed step-by-step after uploading the required data and setting the appropriate parameters. A detailed instruction manual for GRable is provided in Document S1. The data processing, search parameter settings, and results of each step are described below, primarily focusing on improved points compared with the prototype.

### Monoisotopic peak picking (Step 4) with improved accuracy using the correction function

After the ranges of RT and mass were set in the Range setting (Step 3), monoisotopic peaks were selected using the original algorithm in Step 4. In a previous in-house prototype, the search for monoisotopic peaks started from the highest signal, which is time-consuming. Therefore, GRable uses a new algorithm to improve data processing speed. First, a filter of five scans × 5 Da was used to search for a local peak in each group comprising a single ion. When the highest signal was centered in the filter, it was recorded as a local peak, and the other signals within the filter were excluded from the local peak candidates. The local peaks were used as the starting points for identifying the corresponding groups comprising the same ion. The local peak signal spectrum was integrated with the same ion spectra before and after the scans, and the resulting spectrum was used to determine the monoisotopic signal. This algorithm change greatly speeds up the monoisotopic peak-picking process.

In addition, the algorithms for finding monoisotopic signals were also modified to improve the accuracy of the monoisotopic assignment in GRable. The prototype used a monoisotopic signal assignment method with criteria based on the relative intensity of the isotope signals; however, the misdetection of monoisotopic ions was frequently observed. The accuracy of the monoisotopic ions is important for accurate glycopeptide assignment. Accordingly, GRable has a function that evaluates the selected monoisotopic peaks using a multinomial distribution based on averagine for a peptide as the abundance ratio of five isotopes, including C, H, N, O, and S. By fitting the observed spectrum with the calculated spectrum, the need to correct the monoisotopic signal obtained in the preceding step is suggested. This correction function facilitated an increased number of matched glycopeptide group members compared to the GRable results obtained without using this function. Users can confirm corrected monoisotopic peaks by the update flag=1 in a “Monoiso Peak List” sheet and in other results sheets of subsequent steps, where the peak no. was identical to that indicated in the monoisotopic peak list. For hAGP, 5,369 monoisotopic signals were detected, of which 2,140 were automatically corrected by the correction function (Data S6), demonstrating its utility for the accurate selection of monoisotopic signals.

### Clustering (Step 5) for efficient and comprehensive detection of glycopeptide signals using parallel clustering function

In this step, glycopeptide signals were detected from the monoisotopic peaks selected in Step 4 as a cluster, based on the relationship between the masses and retention times. The prototype could set a single RT range to find the glycopeptide group as a cluster because the time difference was considered for extending the neutral monosaccharides, namely Hex, HexNAc, and dHex. Glycopeptide groups with different numbers of acidic saccharides, such as NeuAc, were found as clusters (33), because adding neutral saccharides slightly shortens the RT, whereas adding acidic sugar units increases it (39). In the previous RT setting, as the acidic shift increased in proportion to the number of sialic acid units, four glycopeptide clusters with asialo-, monosialo-, disialo-, and trisialo-glycans were found separately in a representative hAGP cluster (Figure 3A). Therefore, glycopeptide groups with the same core but different numbers of acidic units cannot be combined. To improve this, GRable was constructed to allow setting the RT difference separately for each acidic and neutral saccharide and any motif, such as Hex + HexNAc (LacNAc). In the current version, four clusters with the same core peptide were successfully connected into a single cluster, because the range of RT shifts for each unit could be set separately (Figure 3B). This improvement facilitated a better understanding of the microheterogeneity of a glycosite by visualizing the signal intensity of each glycopeptide plot. Notably, a new tetrasialo-glycan was also detected. Such highly sialylated members could not be detected as a cluster because of the limited variety of the glycan stem portion with the four sialylations. Accordingly, this improvement facilitated the increased detection of glycan microheterogeneity for one glycosite. In addition, considering the RT difference separately for each glycan unit is also useful for reducing misdetection caused by isobaric combinations, especially neutral and acidic combinations, such as NeuAc + NH4^+^ and Hex + dHex (24).

**Figure 3.**
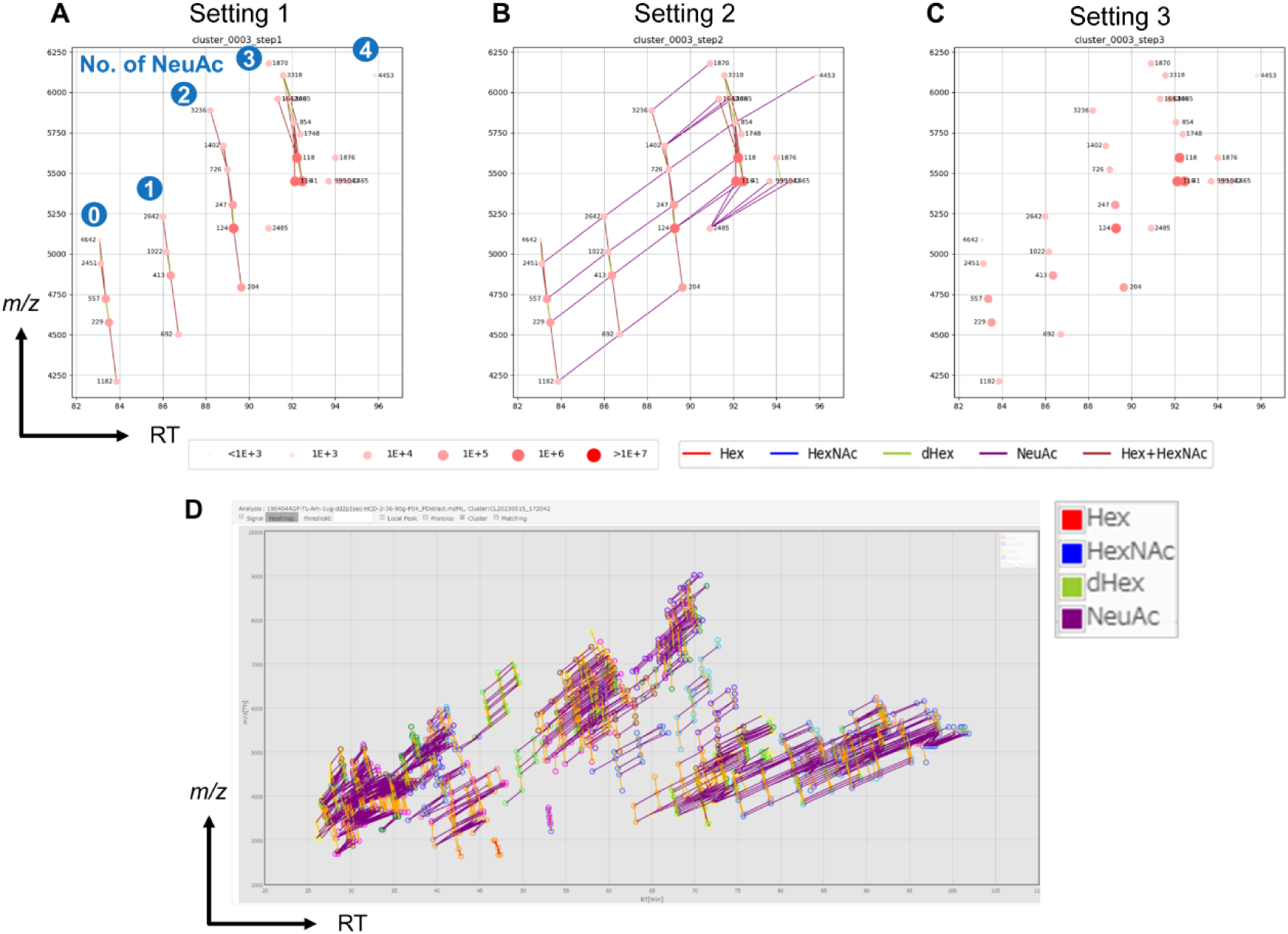
hAGP clustering results. Representative results (Cluster 3) of parallel clustering with three search parameter settings: **(A)** Setting 1 (a setting in our previous prototype), **(B)** Setting 2 (an extra setting for detecting sialylated glycopeptides), and **(C)** Setting 3 (an extra setting for detecting oligo-mannosylated glycopeptides). The detailed settings are as follows: for Setting 1, neutral saccharides (Hex, HexNAc, dHex, Hex+HexNAc): −1–0 (min), and the minimum number of members: 4 (setting for maximizing No. of detected clusters with minimized misdetection); for Setting 2, neutral saccharide: −1–0 (min), acidic saccharide (NeuAc for human): 1–4 (min), and minimum number of members: 10 (setting for guaranteeing the accuracy of assignment); and for Setting 3, neutral saccharide (Hex only): −1–0 (min), and minimum number of members: 3. Dots indicate the monoisotopic masses of the glycopeptide signals at each RT peak, and the size and color depth of the dot indicates the signal intensity. Deep red and larger dots indicate high intensity, and small pale red dots indicate low intensity. All the clusters for hAGP shown in Data S6 can be visualized in the viewer of the GRable main window **(D)**. The colors of lines that connect the signals indicate the type of glycan unit between the signals: red: Hex, blue: HexNAc, green: dHex, Hex + HexNAc: brown, and NeuAc: purple.

As is the case with MS2-based approaches, optimization of search parameter settings is crucial for obtaining reliable results in glycopeptide identification (40); thus, optimization is needed for the purpose of analysis. For this need, GRable can integrate multiple clustering results obtained in parallel under different (up to five) search parameter settings for more comprehensive detection of glycopeptide clusters. As the clustering results of each processing step were combined into a single cluster, each search parameter setting was independent of the processing order. This “parallel clustering” function facilitates optimization for the minimum No. of members for clustering, which is the most important parameter that should be optimized according to the purpose of the analysis; a smaller No. of members is expected to result in an increase in the detected clusters; however, the ratio of clusters without supportive information for evaluating certainty will also increase. For example, using this function, the clusters obtained using the previous setting with the No. of members =4 (setting 1; Figure 3A), and the current setting with No. of members =10 (setting 2; Figure 3B) can be easily compared. When the results with No. of members = 2, 3, and 4 were compared for hAGP, the No. of members = 2 resulted in an increased number of detected clusters, but most of the increased clusters were excluded in the subsequent selection step (Table S2). In addition, manual inspection after the selection step was required with No. of members = 2. In contrast, with No. of members = 3, more assignments were obtained without manual inspection after selection compared to No. of members = 4, demonstrating that the No. of members = 3 is the most suitable for hAGP analysis.

We often use a setting that considers only the number of Hex to find clusters, with the aim of detecting glycopeptide clusters with only high- or oligo-mannosylated glycans. There are many possible compositions of complex- and hybrid-type glycans; however, many oligo-mannosylated glycans are limited to M9–M5. Thus, the minimum cluster number was lowered to three to estimate oligo-mannosylated glycan carriers. In the hAGP samples, the abundance of oligo-mannosylated glycans was low, resulting in no detection in this setting (Setting 3; Figure 3C). This finding is consistent with previous reports that glycan structures attached to hAGP are branched, sialylated and fucosylated at the branches, and slightly extended by polylactosamine (41–44).

The individual time settings for each type of glycan unit and parallel clustering with different settings effectively increased the detection sensitivity, and multiple clusters with the same core but different sialic acid numbers could now be combined into one cluster. With parallel clustering of these three search parameter settings for hAGP, 65 clusters comprising 1,422 types of glycopeptides were detected (Figure 3D). Notably, among these clustered monoisotopic peaks of glycopeptides, 532 (37%) were corrected by correction functions, demonstrating their utility in improving glycopeptide detection efficiency. The average number of members in each cluster was 19.1. Among the detected monoisotopic signals, 2,309 signals were assigned to glycopeptide clusters.

To evaluate the usefulness of the parallel clustering function for detecting glycopeptides with a specific glycan structure, such as oligo-mannosylated glycopeptides, we used a dataset of HL-60 cell lysates (HILIC fraction 2) as a complex mixture of unknown proteins containing glycopeptides decorated with oligo-mannose (32). Because the sample glycopeptides used were desialylated, parallel clustering was performed under two search parameter settings: setting 1 (previous setting) and setting 3 (oligo-mannosylated glycans) for the hAGP sample. Figure 4A shows the clusters detected in setting 1, in which the minimum number of clusters was set to four. When setting 2 (only Hex was considered a glycan unit and the minimum number of clusters was three) was added to the setting 1 search, the number of detected clusters increased (Figure 4B). For clarity, only the increased numbers of clusters (No. of members: 3) were visualized (Figure 4C), demonstrating the presence of oligo-mannosylated glycopeptides. Thus, GRable can be applied to large-scale and in-depth analyses of glycoproteins, in which the comprehensiveness of glycan heterogeneity is improved, and glycan structure-specific search is allowed.

**Figure 4.**
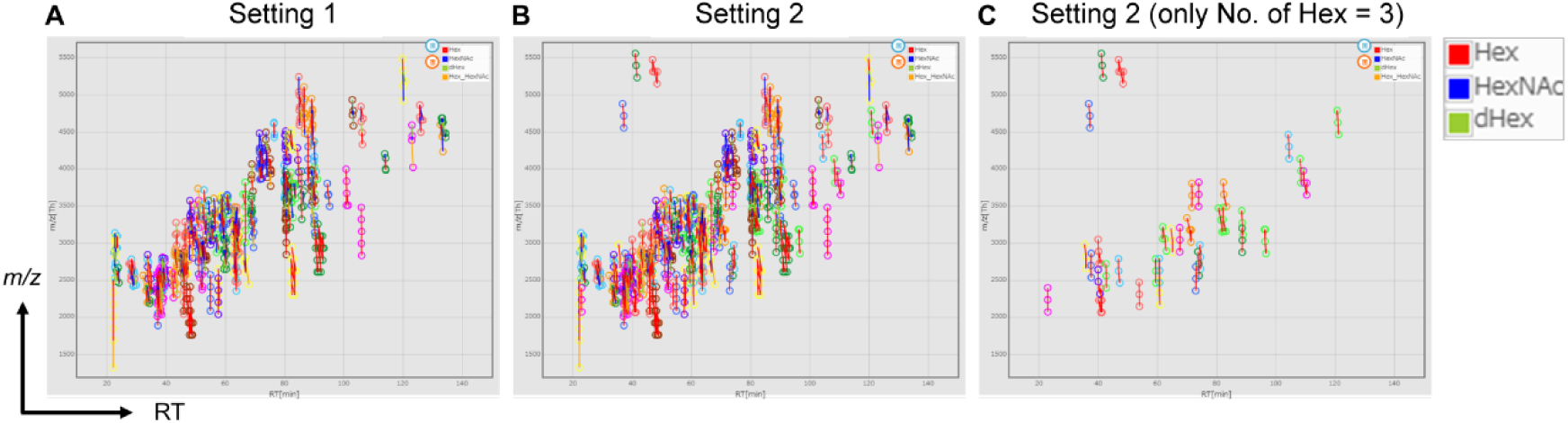
Clustering view of HL-60 cell lysate samples. All the clusters for HL-60 cell lysates (HILIC Fr.2) shown in Data S8 can be visualized in the viewer of the GRable main window. Parallel clustering was performed under two search parameter settings: **(A)** Setting 1 (a setting of the prototype) and (**B**) Setting 2 (a setting for detecting oligo-mannosylated glycopeptides). The detailed settings are as follows: for Setting 1, neutral saccharides (Hex, HexNAc, dHex, Hex+HexNAc): −1–0 (min), and minimum number of members: 4; for Setting 2, neutral saccharide (Hex only): −1–0 (min), and minimum number of members: 3 Among the detected clusters in Setting 2, only clusters with three members having oligo-mannosylated glycans is presented **(C)**.

### Matching (Step 6) with improved evaluation of the analysis using inter-clustering analysis function

In this step, GRable comprehensively searches all combinations of core peptide and glycan compositions for each glycopeptide cluster by consistency in their masses, based on the lists of possible peptide and glycan compositions provided by users. Accordingly, the results sheet shows information on the three matched components (glycopeptides, peptides, and glycans) and the differences from the calculated mass values for each cluster. Note that this step considers matching by mass alone; thus, all matched results that meet the criteria can be provided for a single cluster. In the subsequent step, the most plausible match was selected from among the matching candidates for each cluster, and the accuracy of the selected match was evaluated. The likelihood of the assigned glycan composition was crucial for this selection. Accordingly, positive points for likely compositions (and negative points for unusual compositions as an option), which can be defined in the glycan composition list, were assigned to the glycan compositions that matched with the matching result list, and the total score was calculated for each cluster as the sum of the points of all members.

Regarding this step, the previous Glyco-RIDGE analysis was limited by a low match rate. Mass calculations assume that peptides are ionized with protons such as MH^+^. Some clusters with no appropriate match were non-proton adduct clusters. Accordingly, GRable was equipped with an inter-cluster analysis function in the Matching step to detect clusters comprising glycopeptides ionized with non-proton cations such as iron (Fe^3+^) and ammonium (NH^4+^). In this inter-cluster analysis, the presence of such adducts was judged for each unit of the glycopeptide cluster and was unlikely to include unintended misdetection caused by isobaric combinations (e.g., NeuAc + NH_4_^+^ and Hex + dHex (24)). In the matching results for hAGP, using lists of 56 core peptides (Data S1) and 134 glycan compositions (Data S2), 65 clusters were identified. Among them, 46 clusters matched; however, 19 did not (Data S6). Over half of the clusters (i.e., 44 clusters) were related to the corresponding iron or ammonium adducts with the inter-cluster analysis function, as shown in the “relation between clusters” sheet (Data S6). This information facilitates technical evaluation because non-proton adducts cause sensitivity loss; thus, reducing these adducts during sample preparation and analysis is essential. The remaining unmatched clusters may be attributed to the lack of core peptides in the list; therefore, the inter-cluster analysis function provides clues as to how the list should be improved.

### Selection (Step 7) with improved selecting and evaluating of assignments using MS2 information

This last step is crucial because selecting the most plausible match is essential for guaranteeing the certainty of the Glyco-RIDGE analysis results, considering that the false discovery rate cannot be defined. Because this selection step was conducted with manual inspections in our previous procedure, using this step with clear criteria in GRable represents a large technical improvement. As previously described, a crucial criterion is the total score of each matched cluster. Accordingly, this step sets the total score threshold, and only matches within the threshold are shown in the selection result sheet. When the Clustering and Matching steps are performed correctly, all members match any composition, and many points are obtained based on the glycan point list. In addition to the total score, additional criteria were the delta mass and delta RT between the glycopeptide and the corresponding core peptide, albeit unused for automated selection by GRable. The usefulness of relative RT information has also been demonstrated in MS2-based *N*-glycoproteomics (24). Thus, all members within these criteria are highlighted in the results sheet, visually confirming the certainty of the results. Notably, identical results were obtained for hAGP analysis, even without the RT of the core peptide (i.e., RT=0 for all peptides in the core peptide list), suggesting that it is possible to use predicted peptide lists when an analyte is a purified glycoprotein.

The commercial hAGP glycoprotein specimen used here also contained serum glycoproteins such as haptoglobin (HPT) in the core peptide list (Data 1). For cluster 1, two candidate core peptides, hAGP isoform 1 (A1AG1) and HPT, were used. The total scores for A1AG1 and HPT were 28 and 11, respectively. However, the delta RTs of all members for HPT are over 15 min, whereas the delta RT of A1AG1 members is within −6.5–5.1 min, meeting the defined criteria. Accordingly, A1AG1 was selected as the core peptide for cluster 1. Similarly, the most plausible matching can be easily selected for all matched clusters.

As the other important improved point, this selection step allows evaluating the “confidence level” of the selected results by using MS2 information. This scrutiny and curation step to reduce misassignment and ambiguity is critical in current *N*-glycoproteomics (37). Therefore, as with many MS2-based glycopeptide identification methods (12), GRable first searches glycan fragment ions called “diagnostic ions” in MS2 spectra for cluster members to evaluate whether the spectrum is attributed to a glycopeptide. Diagnostic ions, such as fragments of HexNAc and HexNAc + Hex, were searched as defaults. The variation in the diagnostic ions can be set without restricting the number of ions. These diagnostic ions are not used for selection and are only added to the selection result list, allowing users to utilize this information for different purposes. In addition, glycopeptide fragments such as Y0, Y1, and Y2 were searched to estimate the mass of Y0, namely, the peptide moiety. Because the Y0 signals are often too weak to be detected, GRable is equipped with a function for predicting the Y0 mass based on the masses of other related signals if multiple signals are present. A matched peptide was considered correct if the mass value coincided with that of the matched peptide. This MS2 information is summarized for the selected clusters in the “MS2 info for Clusters” sheet, and all the search results are available in the “All MS2 info” sheet of an exported file (Data S6). The presence of such MS2 information is visible in the “Selection results” sheet and accessible directly via an intra-file link for all glycopeptides in the sheet.

For hAGP, all assigned glycopeptides showed MS2 spectra with HexNAc(204) and HexNAc + Hex(366) as diagnostic ions characteristic of glycopeptides, supporting the certainty of assignment. In addition, 45 of the 78 monoisotopic peaks in cluster 1 were accompanied by MS2 information, and the predicted peptide sequence was an A1AG1-derived peptide (WFYIASAFRNEEYNK). Therefore, the selection result for this cluster was confirmed using the MS2 information.

In this context, the Glyco-RIDGE results were presented with the “confidence level” using GRable. The probability/confidence of the assignment for core peptides is ranked as follows: 1) “High”; clusters having any member(s) showing Y0-related ions corresponding to the presumed core, 2) “High”; clusters having any member(s) identified as the same core using the MS2-based search engine, 3) “Medium”; clusters having any member(s) showing glycan diagnostic ions, 4) “Low”; clusters without any MS2 support. We believe that the confidence of the assignment is high even in the “Low” rank when their mass and RT differences fit under the threshold and the glycan point is the highest in the cluster. This was also supported when the assigned core peptide for a low-rank cluster was found in the core peptide list based on confident deglycopeptide identification for the identical samples with 1% FDR. Originally, GRable was designed to estimate signals that MS2 spectra could not identify; thus, cluster members without MS2 support and clusters with no MS2 data for all members are notable features of GRable. Such a glycopeptide detection methodology allows the evaluation of the certainty of MS2 information-lacking assignments based on their connected members in the same cluster. In this evaluation, visualization of the two-dimensional (i.e., RT and mass) relationship of each cluster member in the cluster sheet allows manual inspection of whether an unexpected glycan composition is likely; with this manual evaluation, unconvincing assignments can be excluded from final results.

### Improved in-depth glycoproteomics results using GRable

Two isoforms of hAGP exist, isoform 1 (A1AG1) and isoform 2 (A1AG2), of which A1AG1 is the major component. Based on the improved GRable procedure, five glycoproteins, including HPT, plasma protease C1 inhibitor, and lymphatic vessel endothelial hyaluronic acid receptor 1 as serum-derived contaminants, were assigned to the hAGP samples (Table S3). Among the 14 assigned clusters, seven (1, 4, 13, 23, 26, 43, and 48) and six (1, 3, 6, 11, 19, and 23) were assigned as A1AG1 and A1AG2, respectively. These clusters covered all five *N*-glycosylation potential sites of A1AG1 and A1AG2 (the two sites are common in A1AG1), except for site 93 of A1AG2; *N*-glycosylation impairment at this site is consistent with a previous report (44). The estimated core peptides for each cluster were the same as those estimated using the MS2 information utilization function of GRable and Byonic, increasing the confidence of the results. Each cluster strictly corresponded to one core peptide; thus, the clustering view facilitated effective glycan macroheterogeneity visualization for each glycosite (Figure 5). Notably, two *N*-glycosites (sites 33 and 103) were detected in almost the same core peptide, except for one residue between the two isoforms; however, their site-specific glycoforms could be visualized separately. This result highlights the usefulness of GRable for elucidating differences in the glycosylation status of target glycoproteins, even with other contaminated glycoproteins.

**Figure 5.**
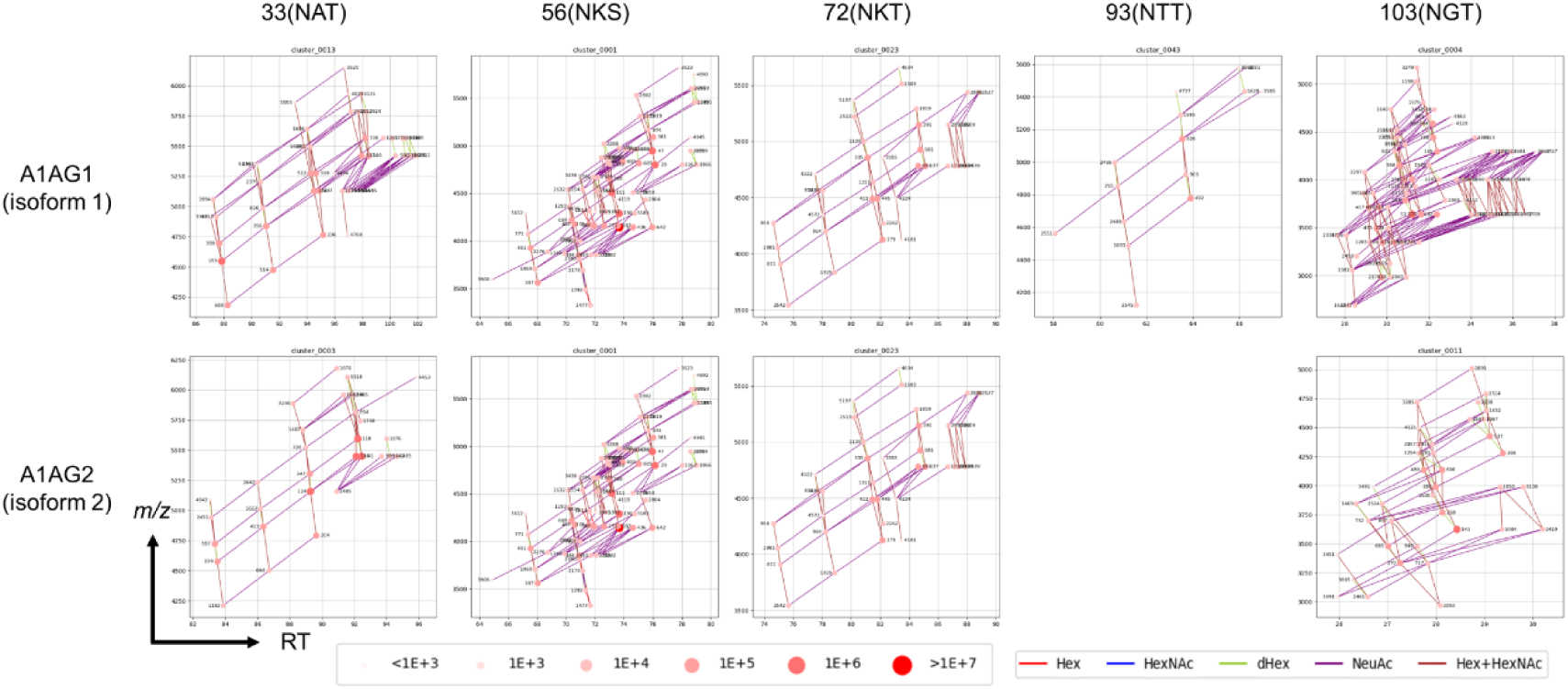
Glycopeptide clusters assigned for 5 *N*-glycosylation sites of hAGP isoforms. Corresponding core peptide sequences for each cluster are indicated in Table 1. Since the core peptide of two isoforms is identical for sites 56 and 72, the same clusters are indicated for both isoforms.

**Table 1.**
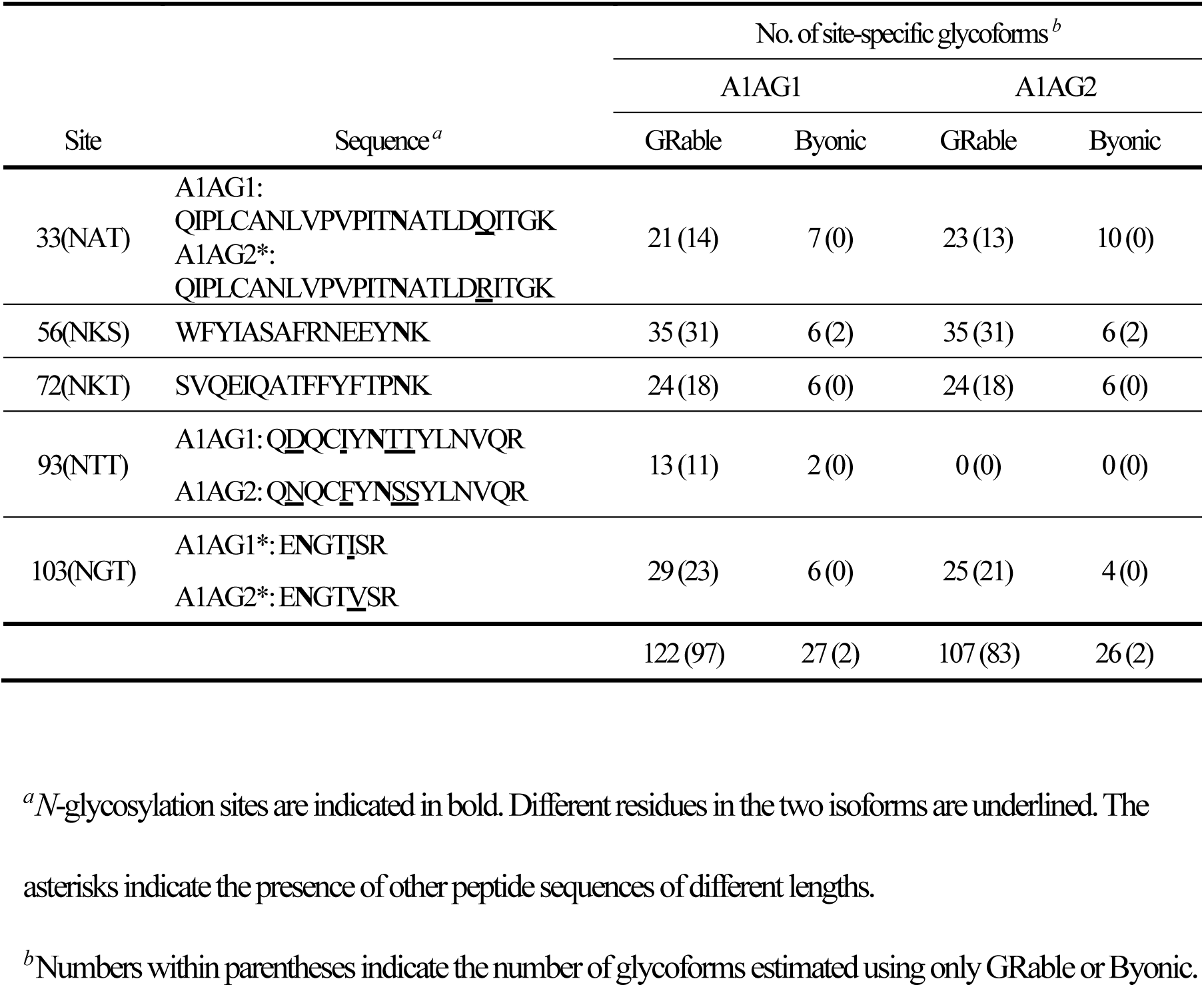
Site-specific hAGP glycoforms estimated using GRable and Byonic.

Using GRable, 123 and 108 site-specific glycoforms of A1AG1 and A1AG2 were estimated, respectively (Table 1 and Table S4). For some *N*-glycosites, the same glycan compositions were estimated for multiple core peptides, supporting the certainty of the results. In the case of site 103 of A1AG1, 24 of 29 compositions assigned for ENGTISR were also detected for the other two peptide sequences. hAGP had *N*-glycosylations with highly sialylated and fucosylated branches and extended with polylactosamine (43, 44). Similar results were obtained with GRable (Table S4); tetrasialo-glycans were estimated at all five A1AG1 sites, indicating high sialylation. Glycans with dHex ≥ 2 were estimated at four A1AG1 sites, indicating its fucosylation at both its core and branches. In addition, glycan components for polylactosamine (i.e., Hex + HexNAc ≥ 10) were estimated at four sites, indicating the presence of such extended glycans.

### Evaluation of the certainty of glycoproteomic results using GRable

To evaluate the suitability of the search parameter settings used here for accurate glycopeptide detection, we performed a GRable analysis of the IGOT-LC/MS data (i.e., *N*-deglycoproteomic data) used for preparing the core peptide list using the same search parameter settings as those for the *N*-glycoproteomic data. As expected, the detected clusters for *N*-deglycoproteomic data significantly decreased for both hAGP (from 65 to 4 clusters) and HL-60 (from 207 to 32 clusters), and all assigned glycopeptides were also observed for *N*-glycoproteomic data. These results indicate that the settings used for GRable analysis were appropriate.

In the hAGP analysis, the confidence level of all assigned clusters was High or Medium, indicating that at least one member had MS2 information for every glycopeptide cluster. To validate the certainty of the obtained results for hAGP analysis, 110 MS2 spectra of all the assigned glycopeptides with MS2 information were manually checked (Figure S2), confirming that the core peptides were correctly assigned. It is noteworthy that NeuAc-derived diagnostic ions were observed for several glycopeptides without NeuAc owing to insufficient isolation, highlighting the risk of evaluating the accuracy of glycoforms based only on diagnostic ions.

In addition, to evaluate the certainty and significance of the Glyco-RIDGE results obtained using GRable, the results were compared with those obtained using the representative MS2-based software Byonic. Notably, GRable could estimate more than four-fold more site-specific glycoforms than Byonic and covered most of the site-specific glycoforms estimated by Byonic. Only two of the site-specific glycoforms (Hex(2)HexNAc(1) and Hex(3)HexNAc(2)) were estimated by Byonic but not by GRable at site 56) because one component was absent (Hex(3)HexNAc(3) or Hex(3)HexNAc(2)NeuAc(1)). As expected, the MS2-based estimation by Byonic succeeded on higher-intensity glycopeptides (Data S9 and S10). Notably, such highly sialylated and polylactosamine-attached glycopeptides were not detected by Byonic (Table S4), and the median number of NeuAc for 38 glycopeptides assigned by GRable but not by Byonic was 3, whereas the median number of 27 glycopeptides assigned by Byonic was 2. Accordingly, it is likely that MS2 spectra was too poor to assign a glycopeptide, partly due to their low ionization efficiency. Thus, these results confirm the reliability of the Glyco-RIDGE results obtained by GRable, highlighting the usefulness of this method as a complementary method to the current gold standard, MS2-based glycoproteomics.

## CONCLUSION

We developed a novel software program, GRable, to enable semiautomated Glyco-RIDGE analysis. GRable version 1.0 can run online freely with a demo or user data using a web browser via the GlyCosmos Portal (https://glycosmos.org/grable). Note that the current version is guaranteed only for Thermo Fisher Scientific data. Some algorithms of the existing Glyco-RIDGE method were improved during the implementation of GRable version 1.0, which can be applied to in-depth site-specific glycoform analysis of intact sialylated glycopeptides derived from purified and crude glycoproteins. Thus, this software will help analyze the status and changes in glycans to obtain biological and clinical insights into protein glycosylation. The novel parallel clustering function enabled a targeted search focusing on multiple glycan structural layers, including the stem, branching, and terminal moieties, such as Lewis epitopes. This software also allows evaluating of a “confidence level” especially using MS2 information, for each glycopeptide obtained by MS1-based detection methodology. Using the MS2 utilization function opened doors to all glycoproteomics researchers who considered MS2-based glycoproteomics as the gold standard, allowing for the community evaluation of GRable by comparing it with other glycoproteomics software. The combined use of MS1-based glycoproteomics will be useful for expanding glycopeptide detection and providing supporting evidence for MS2-based glycoproteomics. Thus, the Glyco-RIDGE method with its unique MS1-based glycopeptide detection principle has complementary roles to MS2-based glycoproteomic methods, and GRable will be a powerful glycoproteomics tool when combined with currently developed MS2-based software. Accordingly, GRable will be updated by continuously improving the Glyco-RIDGE methodology and software while comparing the upcoming MS2-based methods and software.

## ACKNOWLEDGMENTS

This study was funded by a project for utilizing glycans in the development of innovative drug discovery technologies (grant number JP20ae0101021) from the Japan Agency for Medical Research and Development (AMED). We thank Dr. Mika Fujita and Ms. Masako Sukegawa for their technical assistance. We also appreciate Dr. Kiyoko F Aoki-Kinoshita and Dr. Akihiro Fujita of Soka University for releasing GRable at the GlyCosmos Portal (https://glycosmos.org/).

## DATA AVAILABILITY

The MS data set for hAGP presented in this study has been deposited in the ProteomeXchange Consortium via the jPOST partner repository with the dataset identifiers PXD046226 (ProteomeXchange) and JPST001876 (jPOST).

## AUTHOR INFORMATION

### Author Contributions

All authors contributed to writing the manuscript. All authors have approved the final version of the manuscript.

### Notes

The authors declare no competing financial interest.

## ABBREVIATIONS

Glyco-RIDGE: Glycan heterogeneity-based Relational IDentification of Glycopeptide signals on Elution profile
hAGP: serum α1-acid glycoprotein
HILIC: hydrophilic interaction chromatography
HPT: haptoglobin
HL-60: human promyelocytic leukemia-60
IGOT: isotope-coded glycosylation site-specific tagging
LC/MS: liquid chromatography/mass spectrometry
MALDI-TOF: matrix-assisted laser desorption/ionization time-of-flight
mgf: Mascot generic format
MS: mass spectrometry
PNGase F: peptide-N-glycosidase F
rCM: reduction and carbamidomethylation
RT: retention time
TFA: trifluoroacetic acid

