## Supporting Information (Figure S1 and S2) for "GRable version 1.0: A software tool for site-specific glycoform analysis with improved MS1-based glycopeptide detection with parallel clustering and confidence evaluation with MS2 information"

### Table of Contents


#### Supporting Figure (present in this file)

**Figure S1.** Workflow of glycopeptide analysis for HL-60 cell lysates using GRable.

**Figure S2.** MS2 spectra of glycopeptides assigned for hAGP.

#### Supporting Tables

**Table S1.** List of glycan compositions used for GRable and Byonic analyses of hAGP.

**Table S2.** Effects of changes in the minimum cluster member on hAGP analysis using GRable.

**Table S3.** Assigned site-specific glycoforms for the hAGP sample using GRable.

**Table S4.** Glycan compositions of each *N*-glycosite on hAGP estimated using GRable and Byonic.

#### Supporting Data

**Data S1.** Core peptide list for GRable analysis of hAGP.

**Data S2.** Glycan point list for GRable analysis of hAGP.

**Data S3.** Core peptide list for GRable analysis of HL-60 cell lysates.

**Data S4.** Glycan point list for GRable analysis of HL-60 cell lysates.

**Data S5.** Peak list of the GRable analysis for hAGP.

**Data S6.** Output file of the GRable analysis for hAGP after selection.

**Data S7.** Peak list of the GRable analysis for HL-60 cell lysates.

**Data S8.** Output file of the GRable analysis for HL-60 cell lysates after selection.

**Data S9.** Output file of the Byonic analysis (trypsin-full) for hAGP.

**Data S10.** Output file of the Byonic analysis (trypsin-semi) for hAGP.

#### Supporting Document

**Document S1.** Instruction manual for GRable version 1.0.

#### Supporting Figure

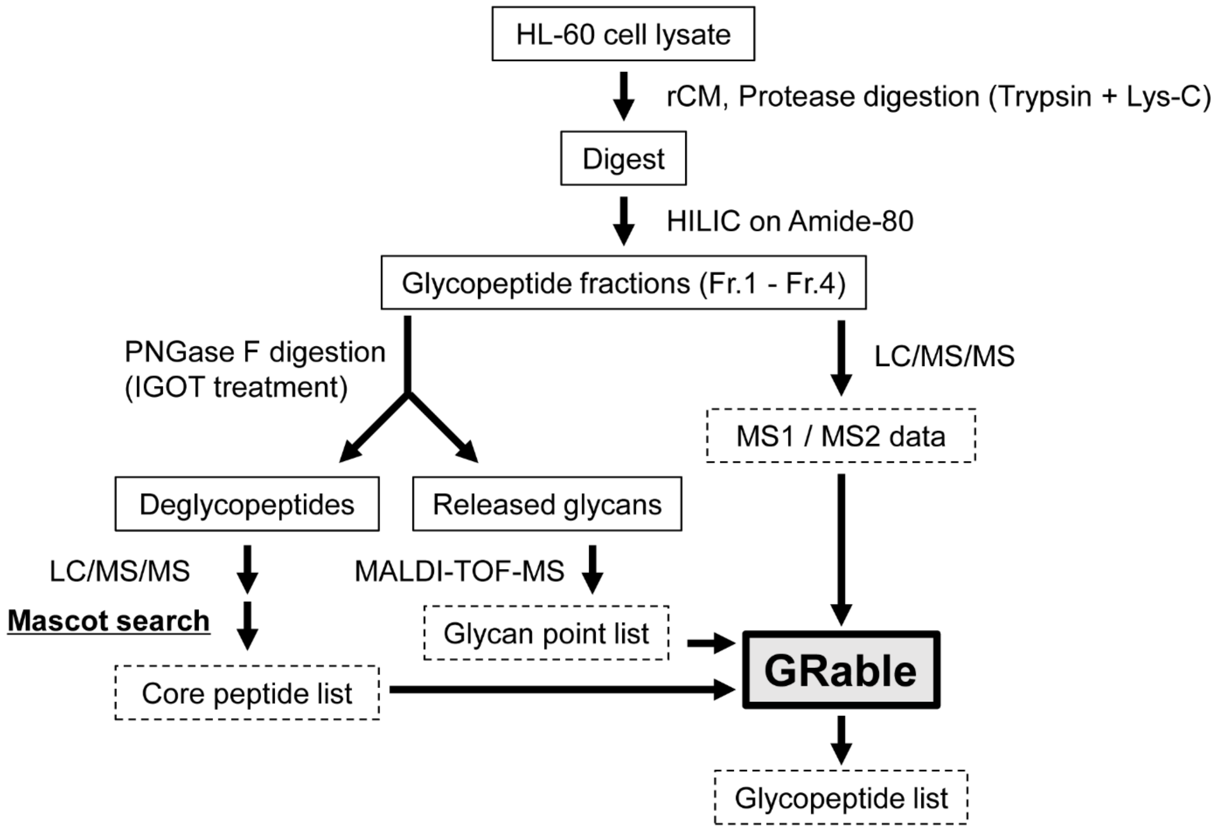


**Figure S1. Workflow of glycopeptide analysis for HL-60 cell lysates using GRable.**

The details of the workflow are documented in the Experimental Procedure. rCM, reduction and carbamidomethylation; HILIC, hydrophilic interaction liquid chromatography; IGOT, isotope-coded glycosylation site-specific tagging; LC/MS/MS, liquid chromatography-tandem mass spectrometry; MALDI-TOF-MS, matrix-assisted laser desorption/ionization time-of-flight mass spectrometry.

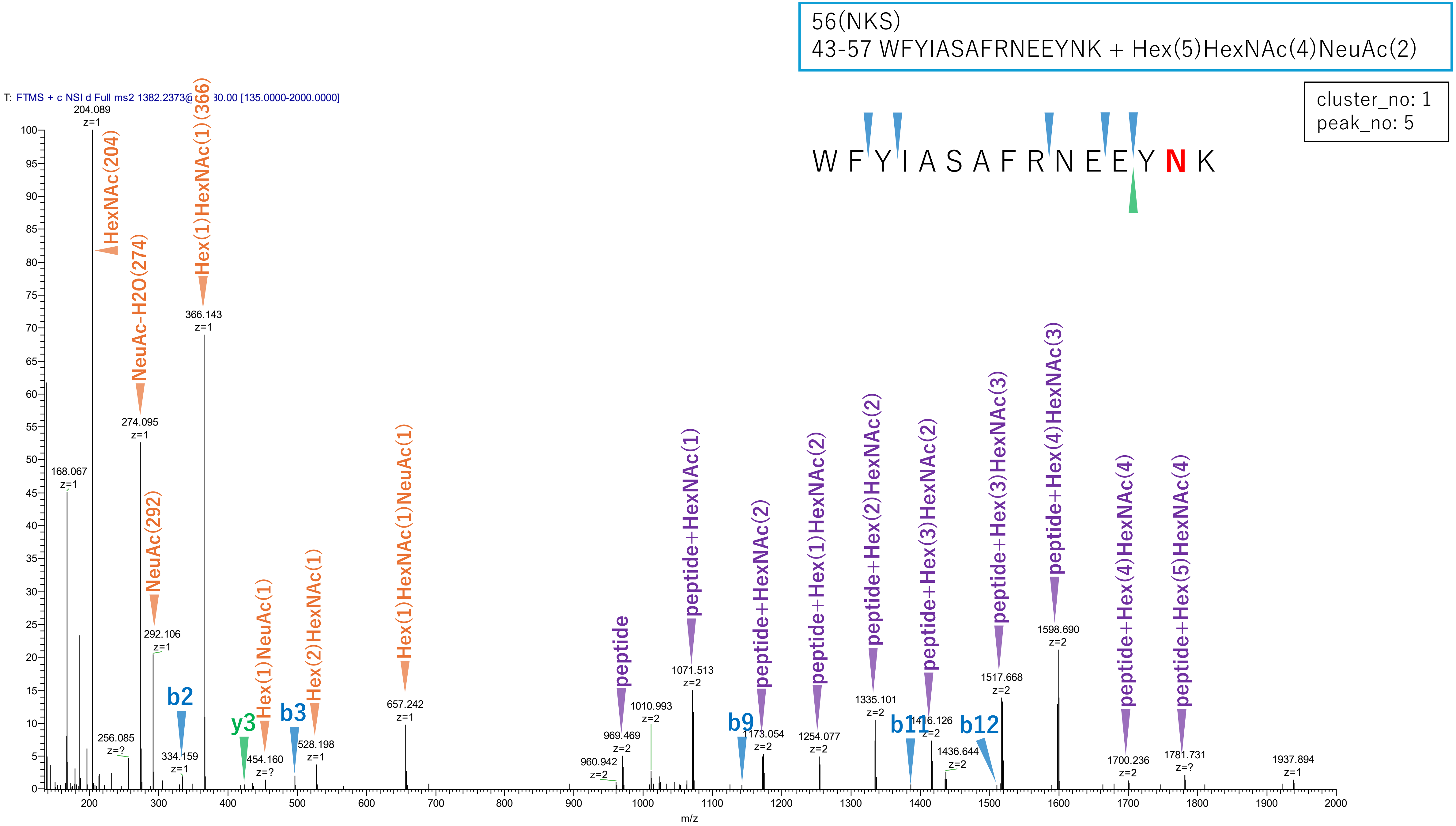


**Figure S2-1. MS2 spectra of glycopeptides assigned for hAGP.**

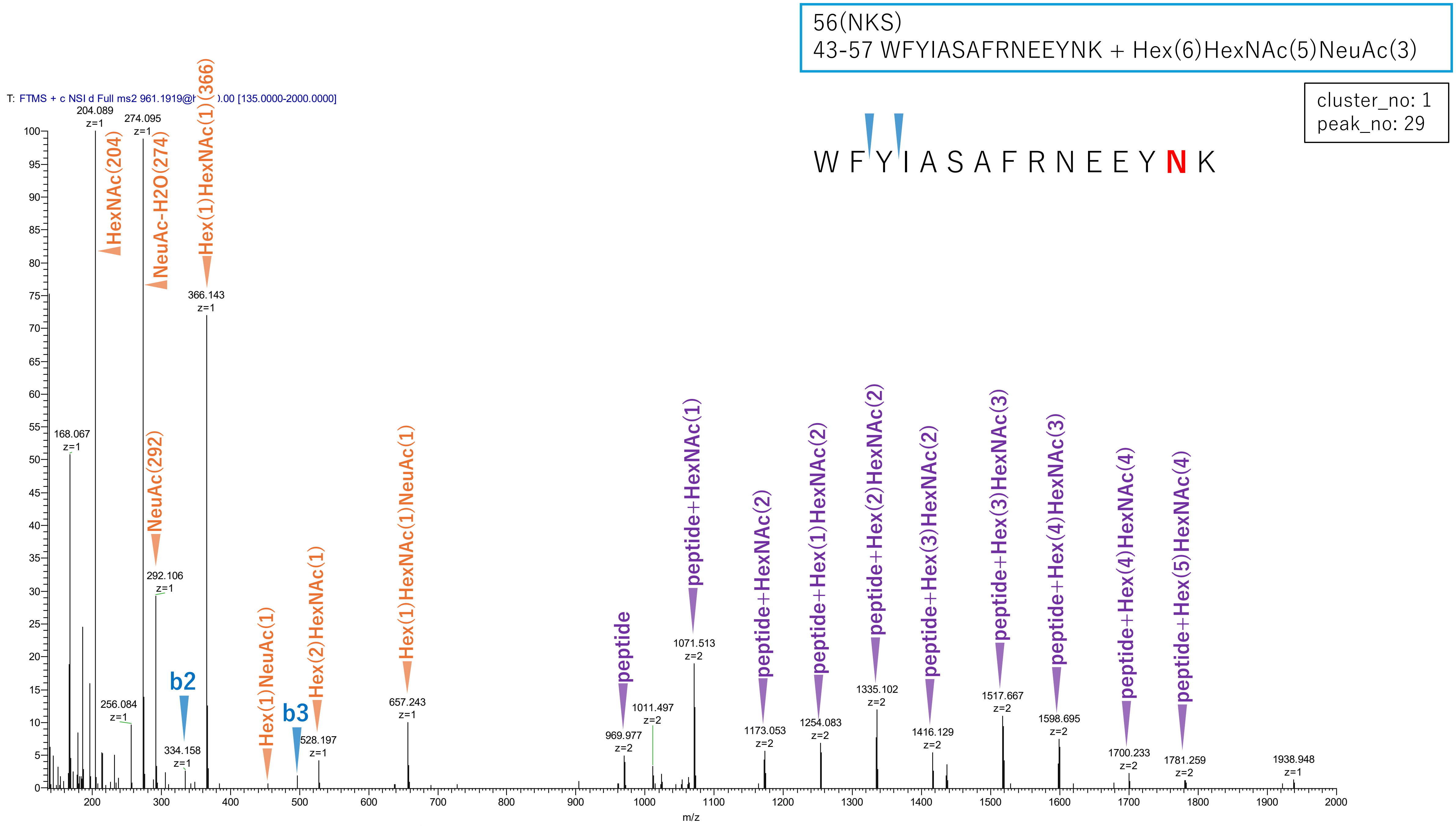


**Figure S2-2. MS2 spectra of glycopeptides assigned for hAGP.**

56(NKS)
43-57 WFYIASAFRNEEYNK + Hex(6)HexNAc(5)dHex(1)NeuAc(3)

cluster_no: 1
peak_no: 47

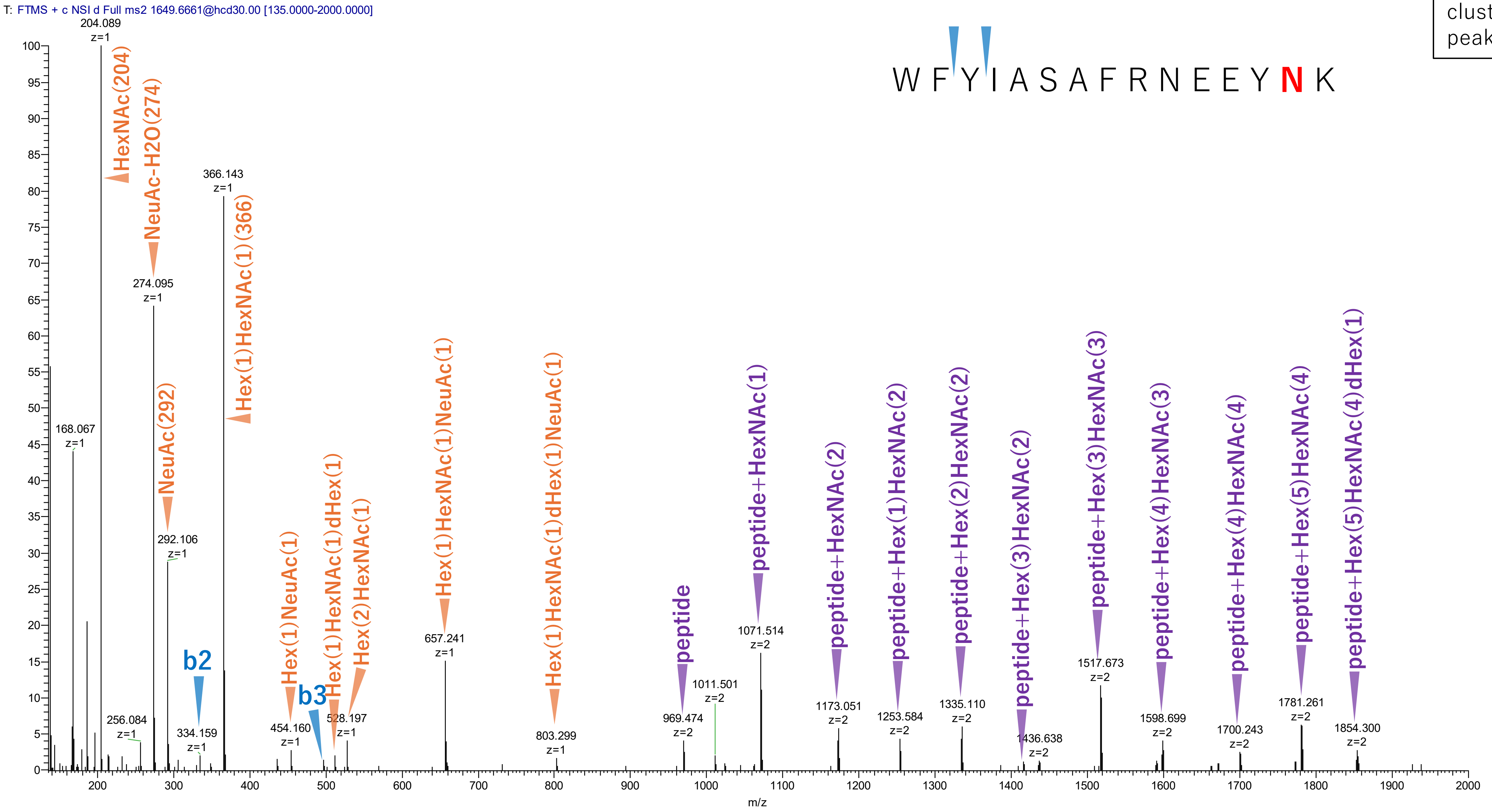


**Figure S2-3. MS2 spectra of glycopeptides assigned for hAGP.**

56(NKS)
43-57 WFYIASAFRNEEYNK + Hex(6)HexNAc(5)NeuAc(2)

cluster_no: 1
peak_no: 111

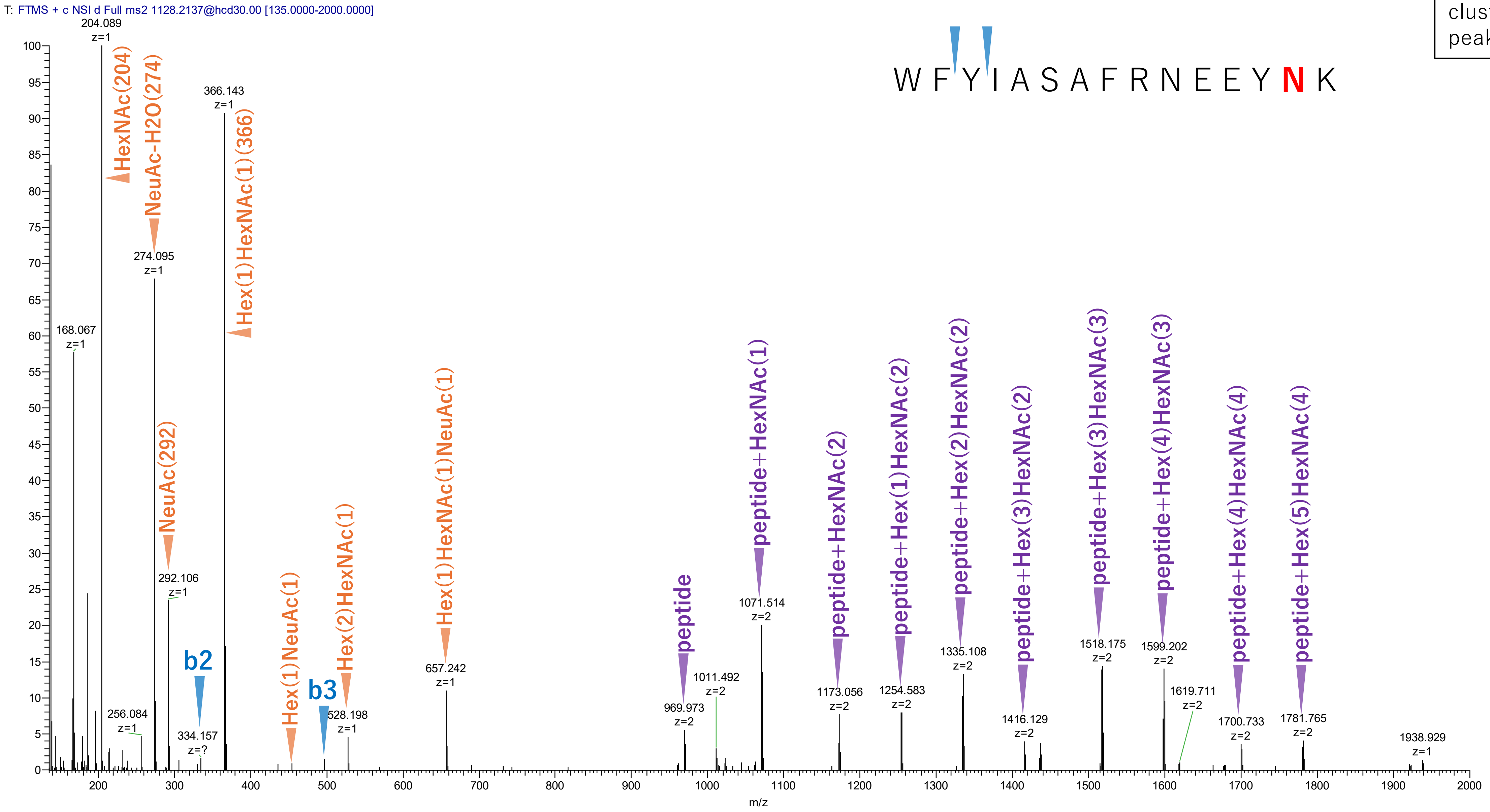


**Figure S2-4. MS2 spectra of glycopeptides assigned for hAGP.**

56(NKS)
43-57 WFYIASAFRNEEYNK + Hex(5)HexNAc(4)dHex(1)NeuAc(2)

cluster_no: 1
peak_no: 130

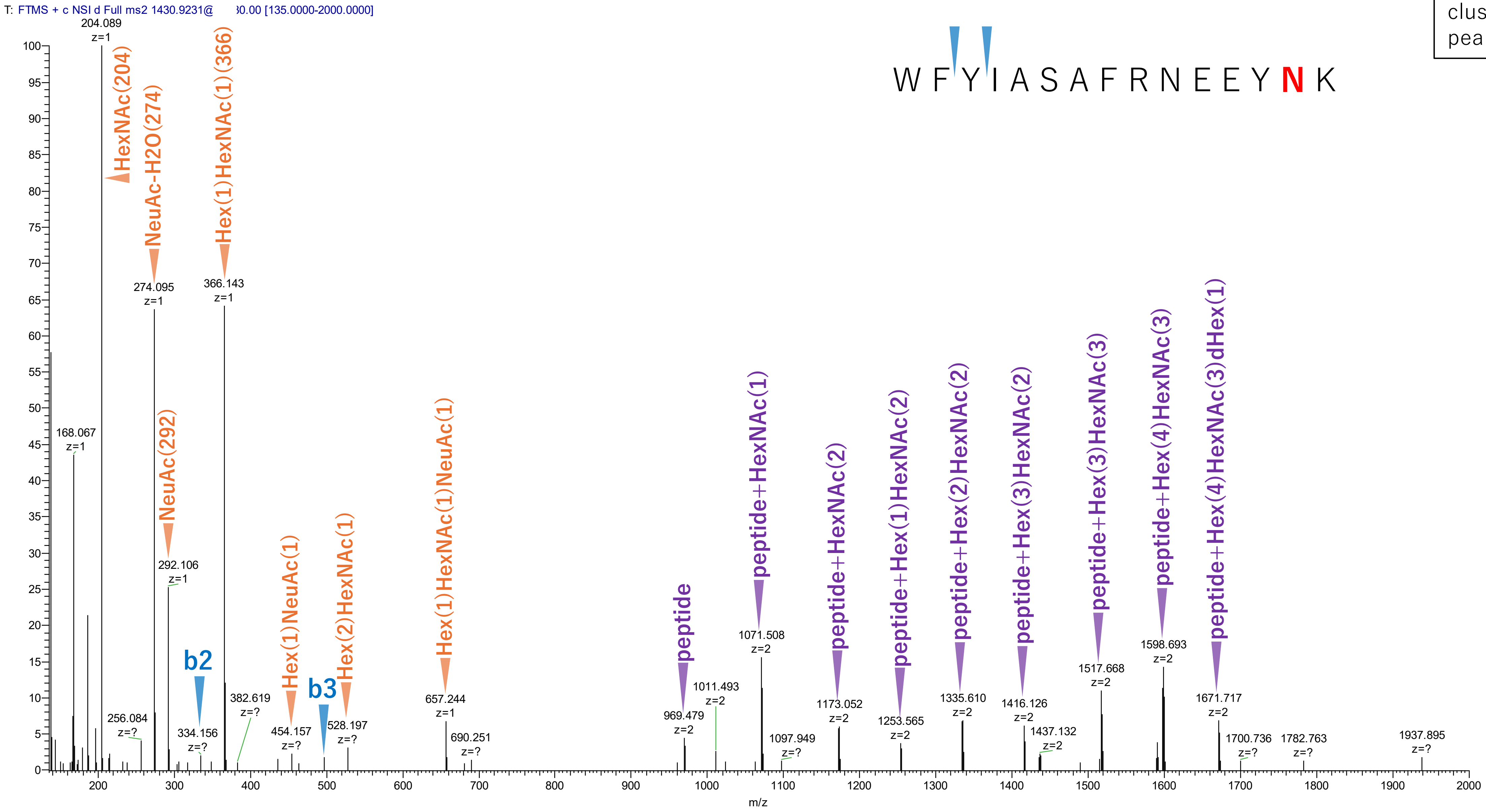


**Figure S2-5. MS2 spectra of glycopeptides assigned for hAGP.**

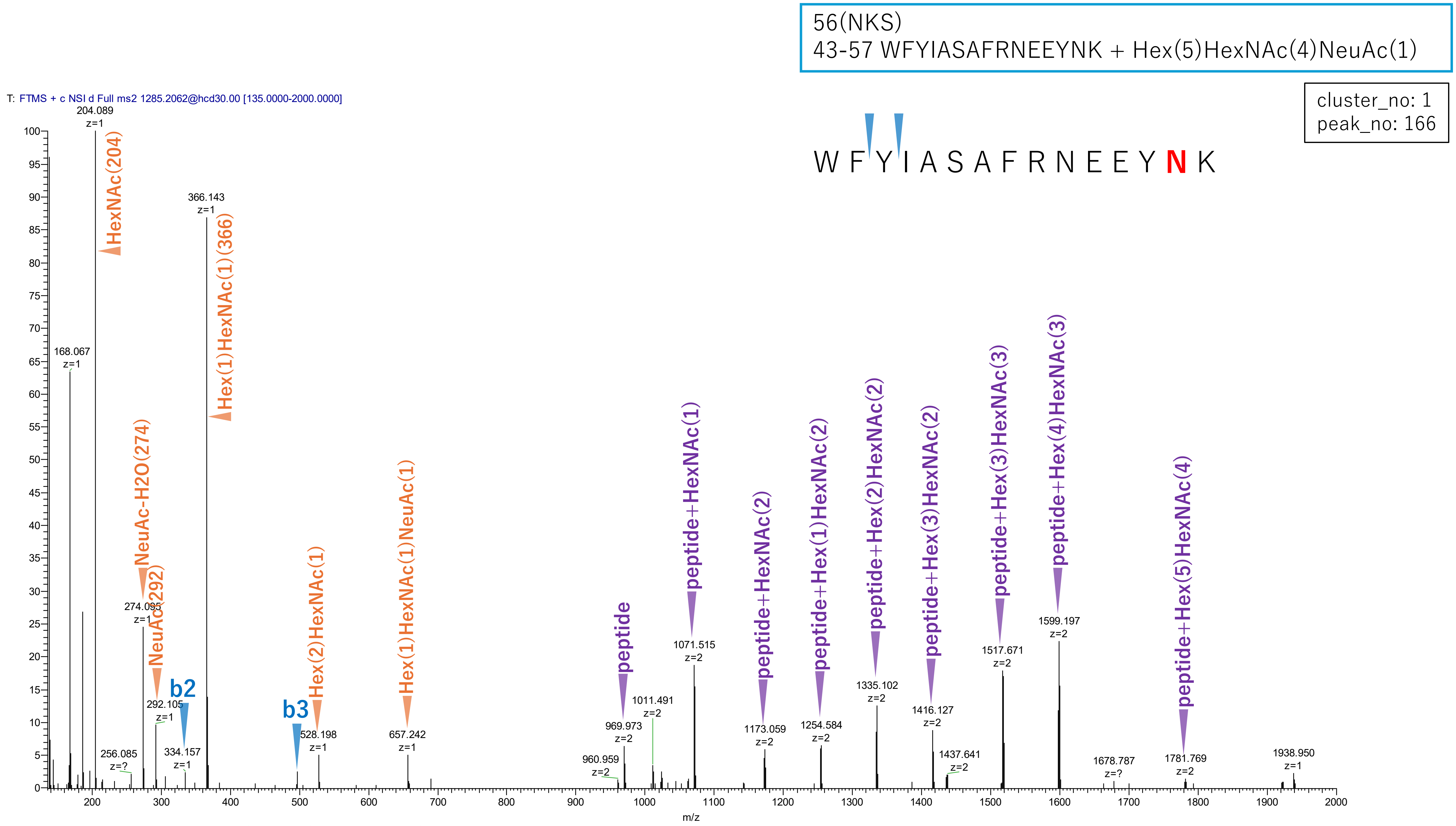


**Figure S2-6. MS2 spectra of glycopeptides assigned for hAGP.**

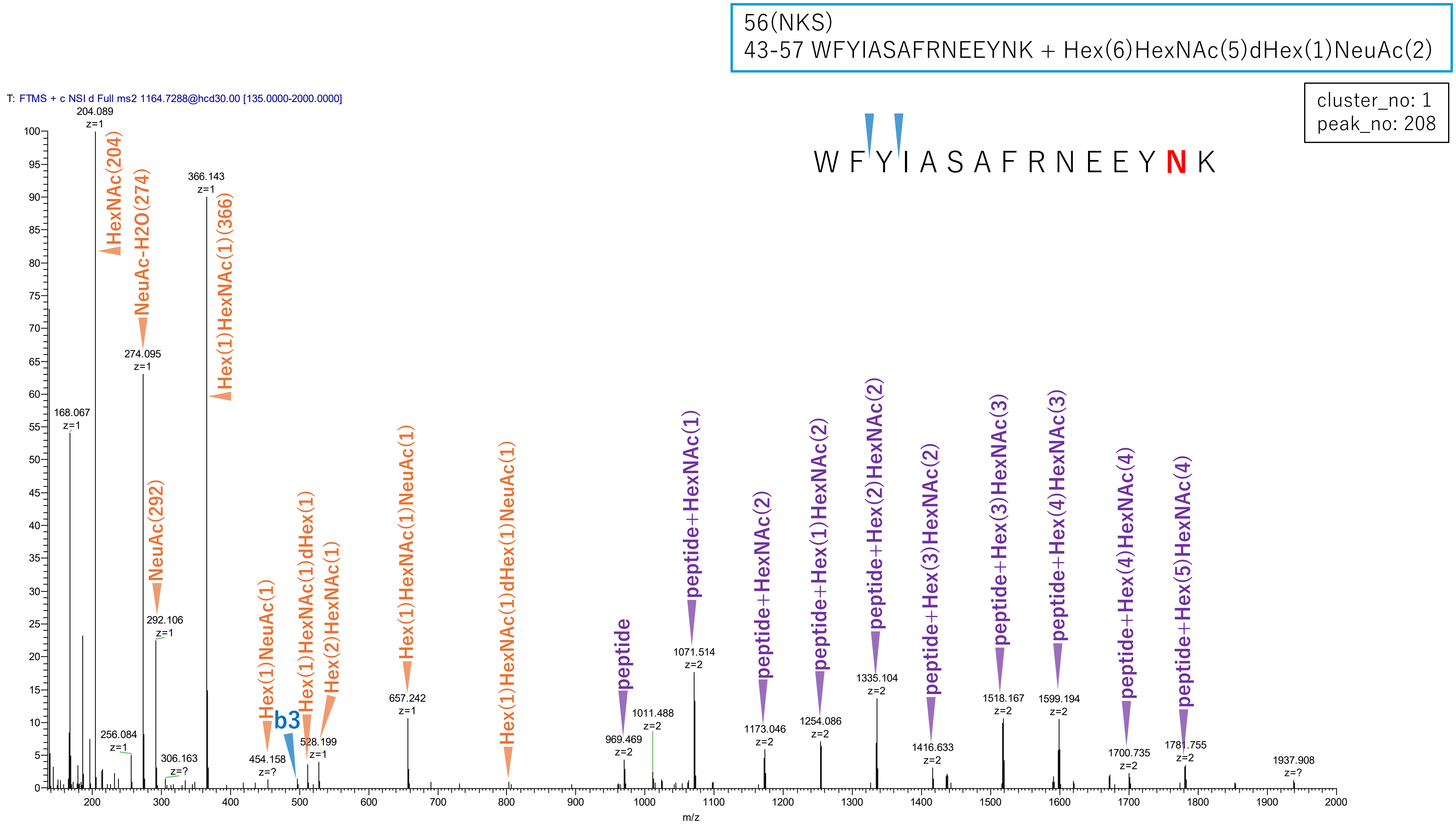


**Figure S2-7. MS2 spectra of glycopeptides assigned for hAGP.**

56(NKS)
43-57 WFYIASAFRNEEYNK + Hex(5)HexNAc(4)

cluster_no: 1
peak_no: 347

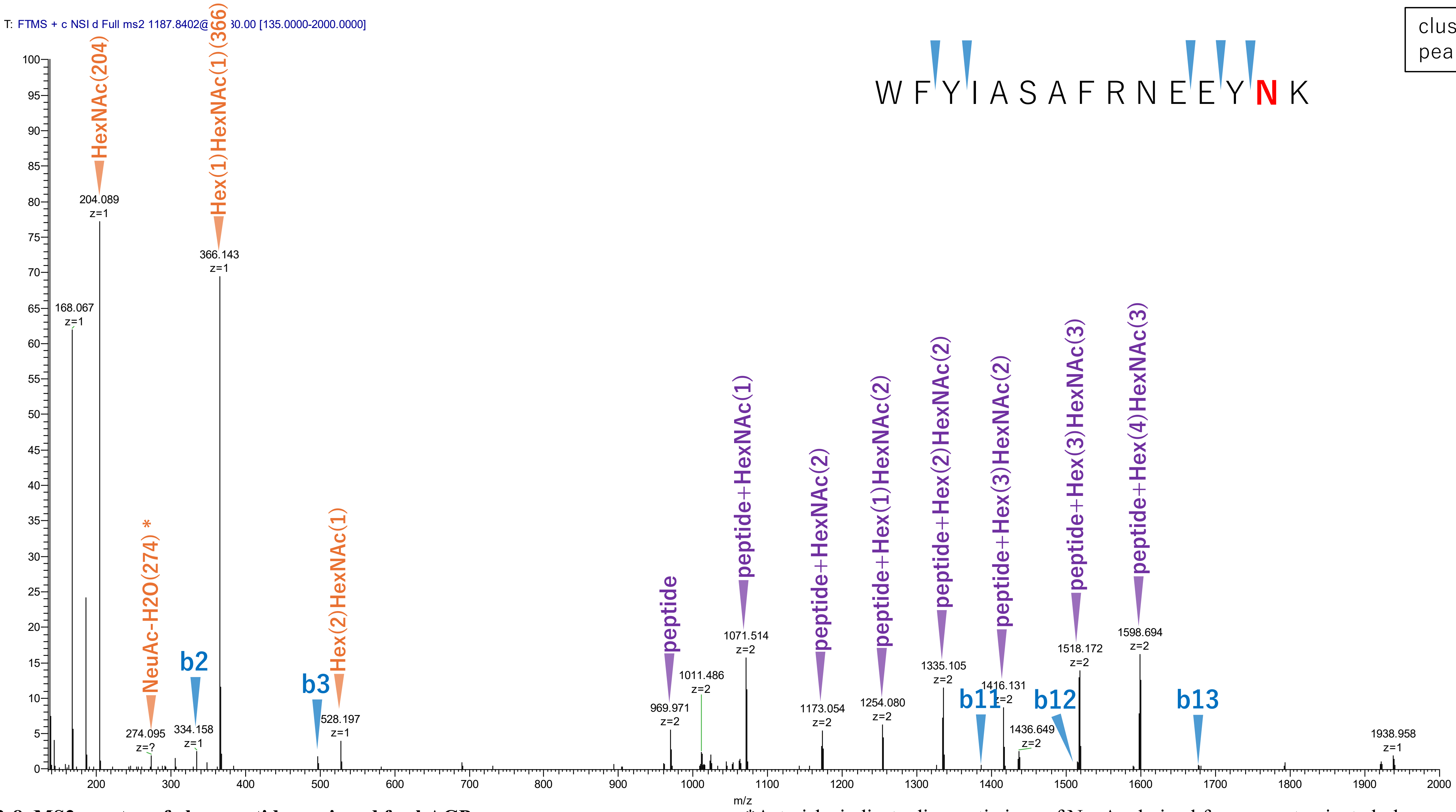


**Figure S2-8. MS2 spectra of glycopeptides assigned for hAGP.**

*Asterisks indicate diagnostic ions of NeuAc derived from a contaminated glycopeptide.

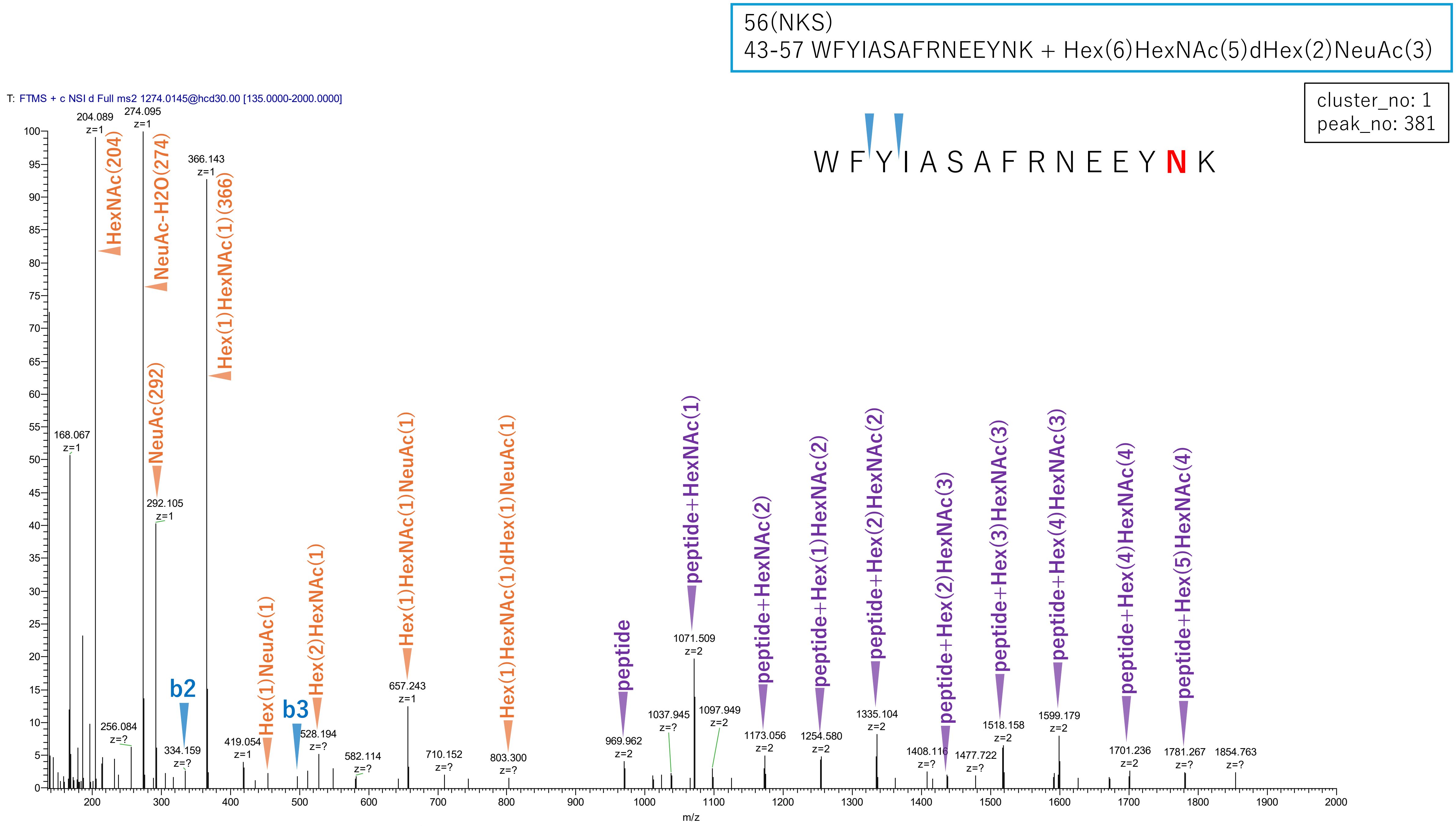


**Figure S2-9. MS2 spectra of glycopeptides assigned for hAGP.**

56(NKS)
43-57 WFYIASAFRNEEYNK + Hex(6)HexNAc(5)

cluster_no: 1
peak_no: 461

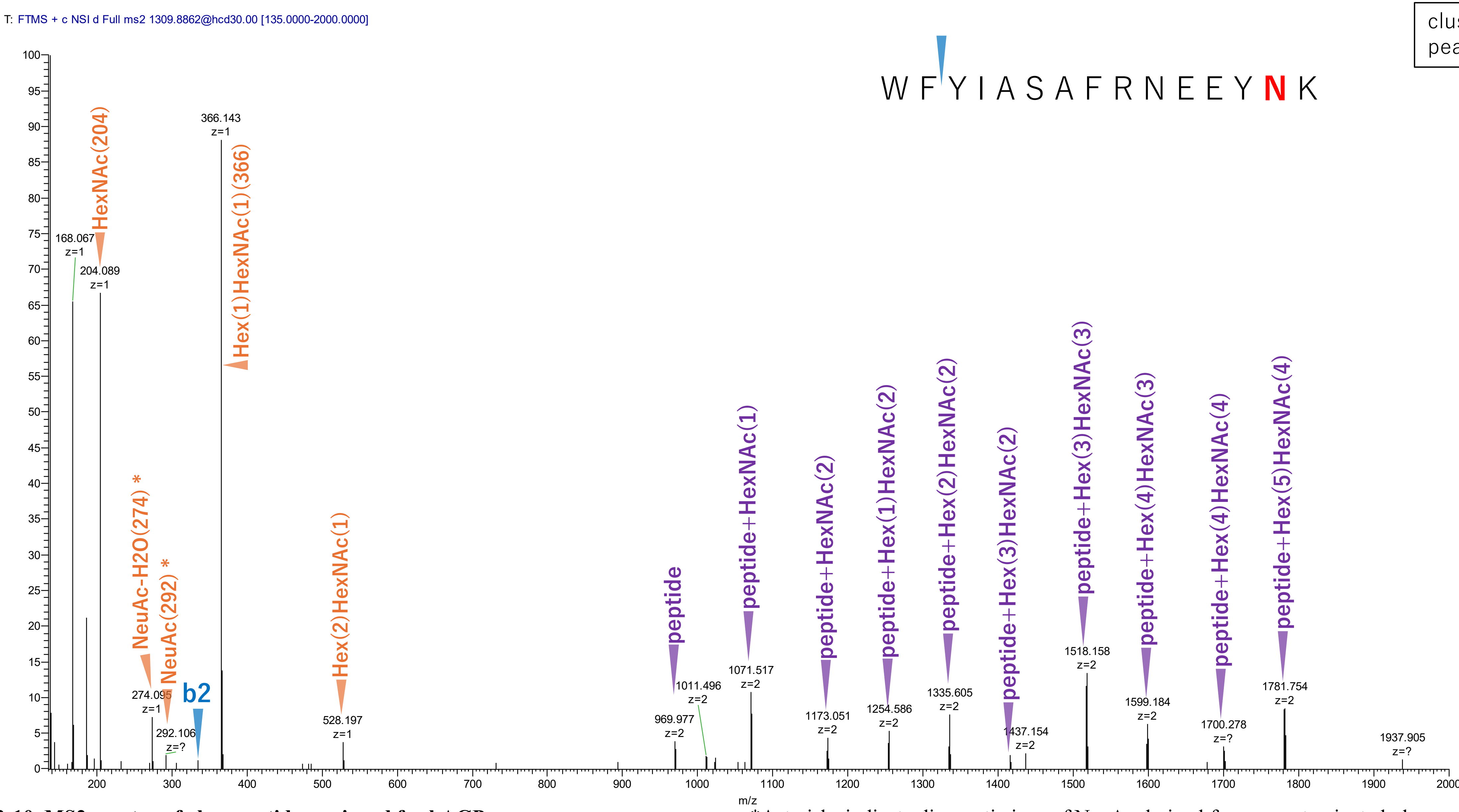


**Figure S2-10. MS2 spectra of glycopeptides assigned for hAGP.**

*Asterisks indicate diagnostic ions of NeuAc derived from a contaminated glycopeptide.

56(NKS)
43-57 WFYIASAFRNEEYNK + Hex(6)HexNAc(5)NeuAc(1)

cluster_no: 1
peak_no: 669

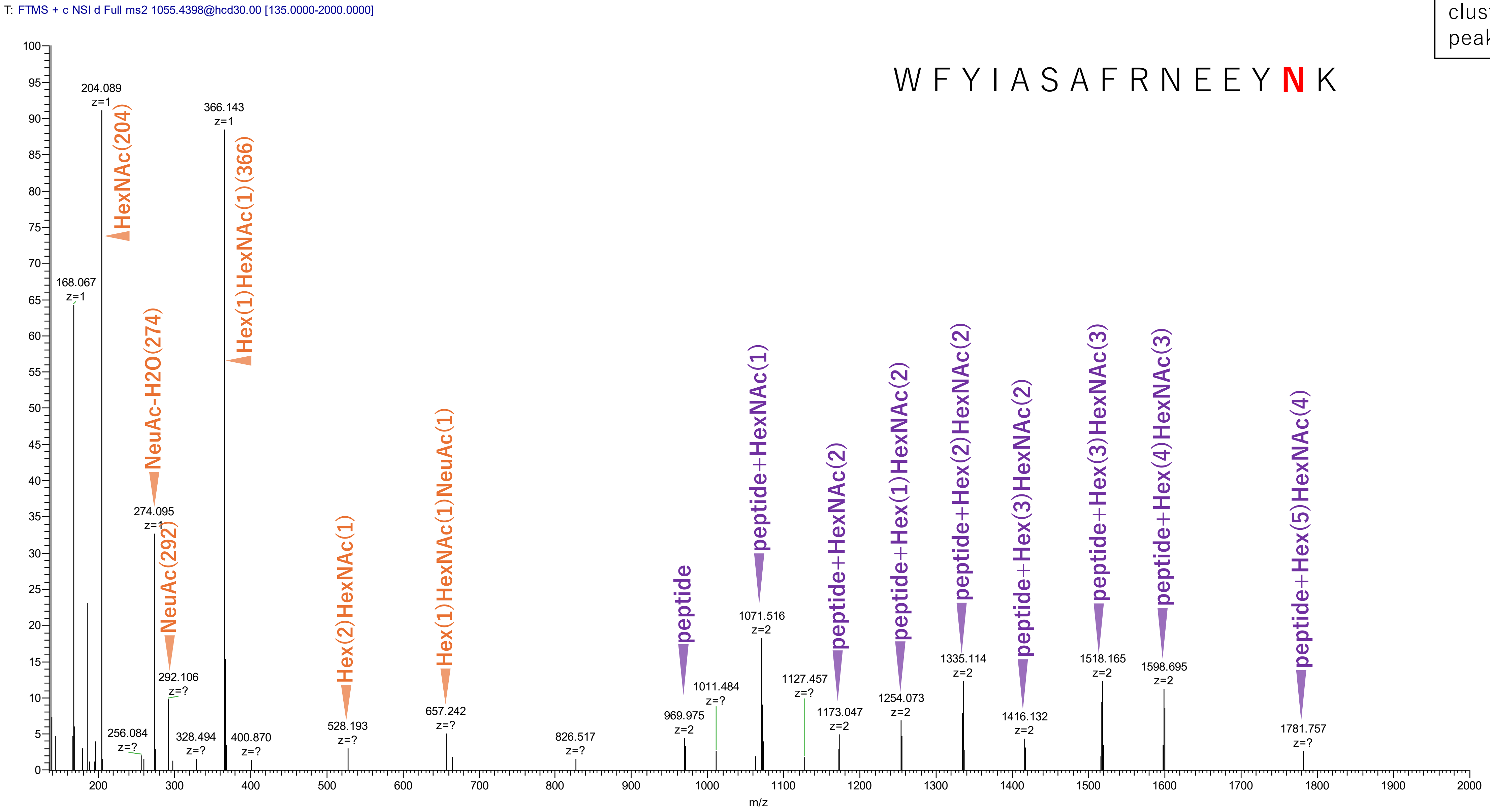


**Figure S2-11. MS2 spectra of glycopeptides assigned for hAGP.**

56(NKS)
43-57 WFYIASAFRNEEYNK + Hex(6)HexNAc(5)dHex(1)

cluster_no: 1
peak_no: 771

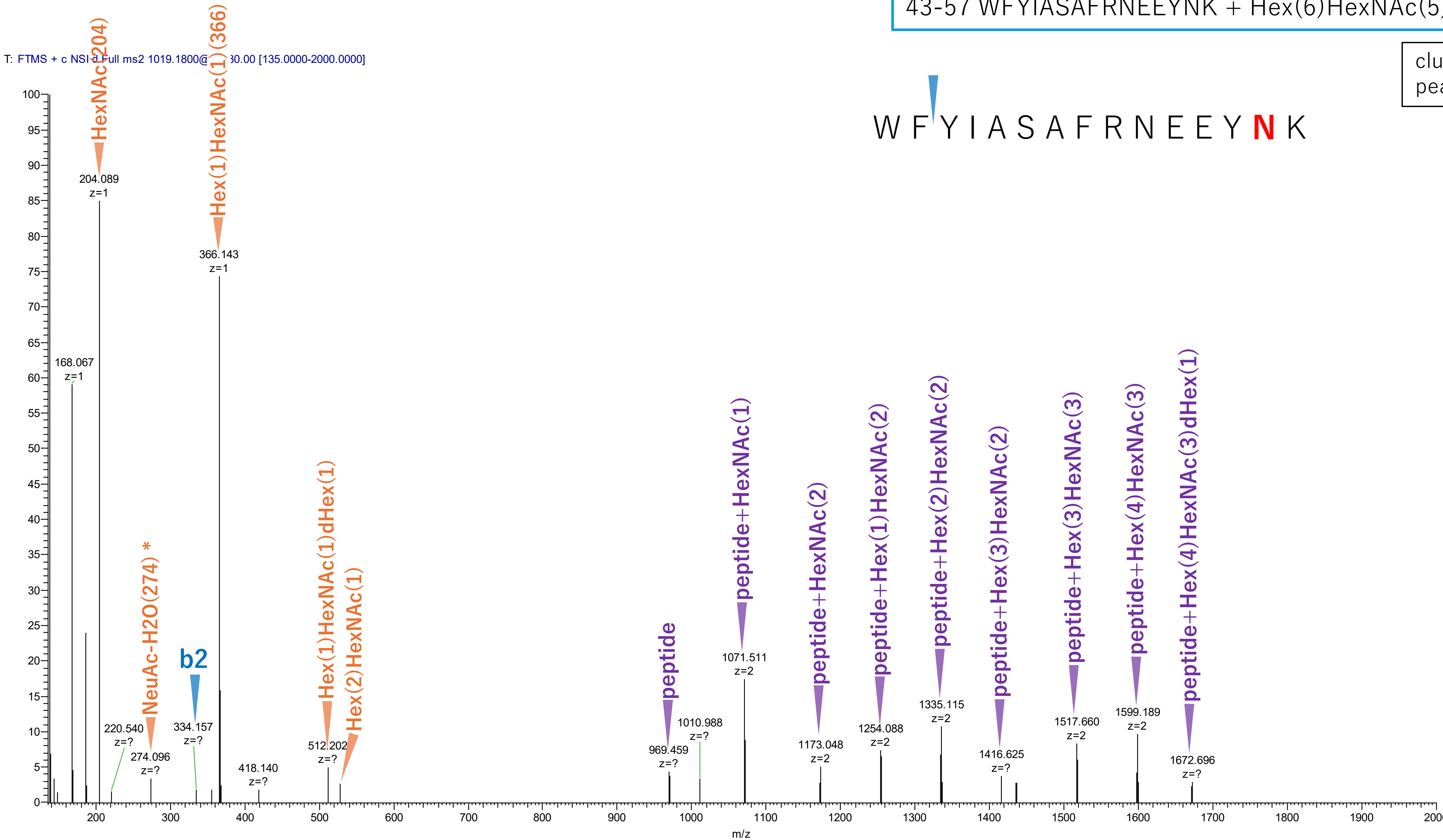


**Figure S2-12. MS2 spectra of glycopeptides assigned for hAGP.**

*Asterisks indicate diagnostic ions of NeuAc derived from a contaminated glycopeptide.

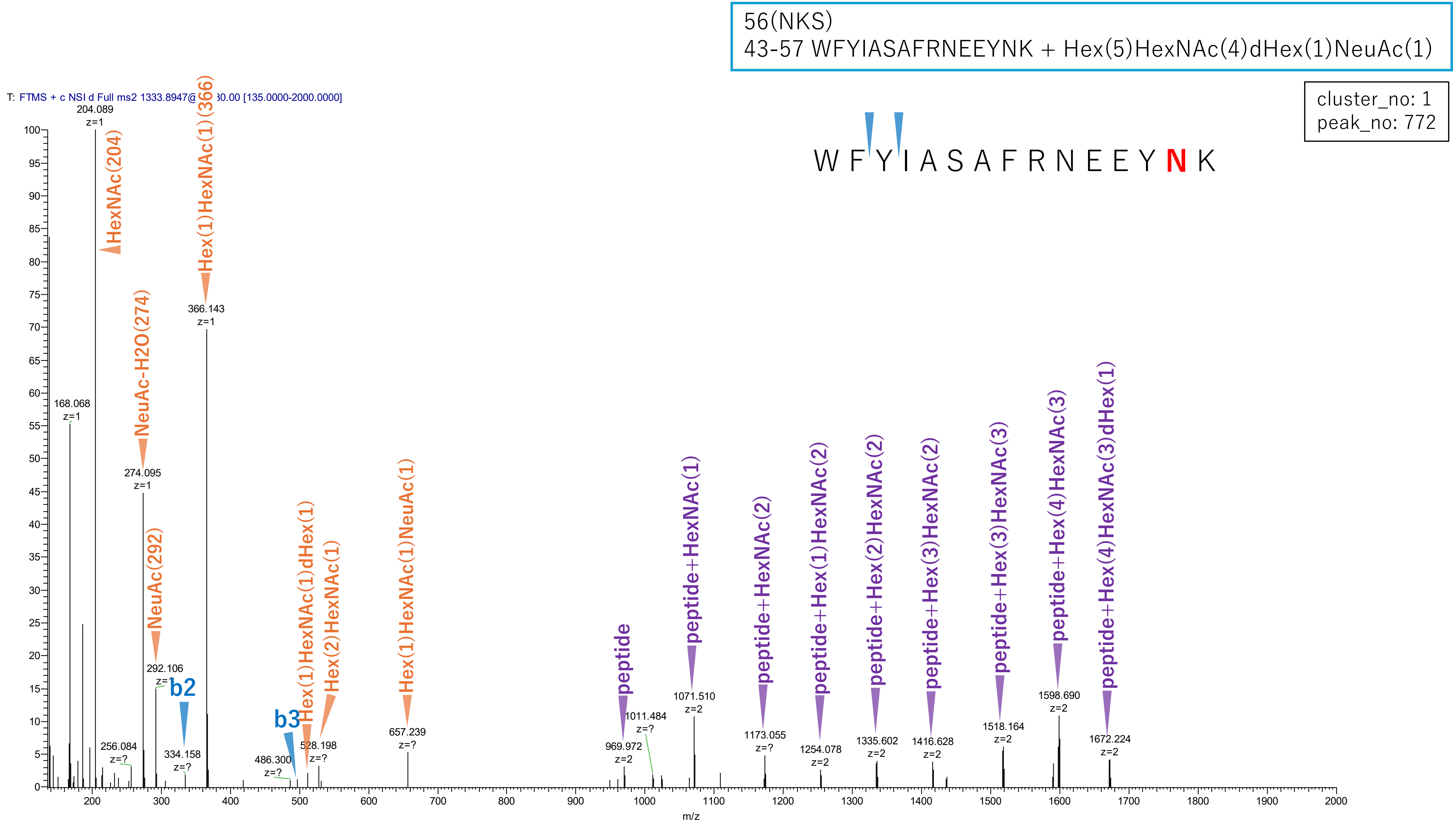


**Figure S2-13. MS2 spectra of glycopeptides assigned for hAGP.**

56(NKS)
43-57 WFYIASAFRNEEYNK + Hex(7)HexNAc(6)NeuAc(3)

cluster_no: 1
peak_no: 974

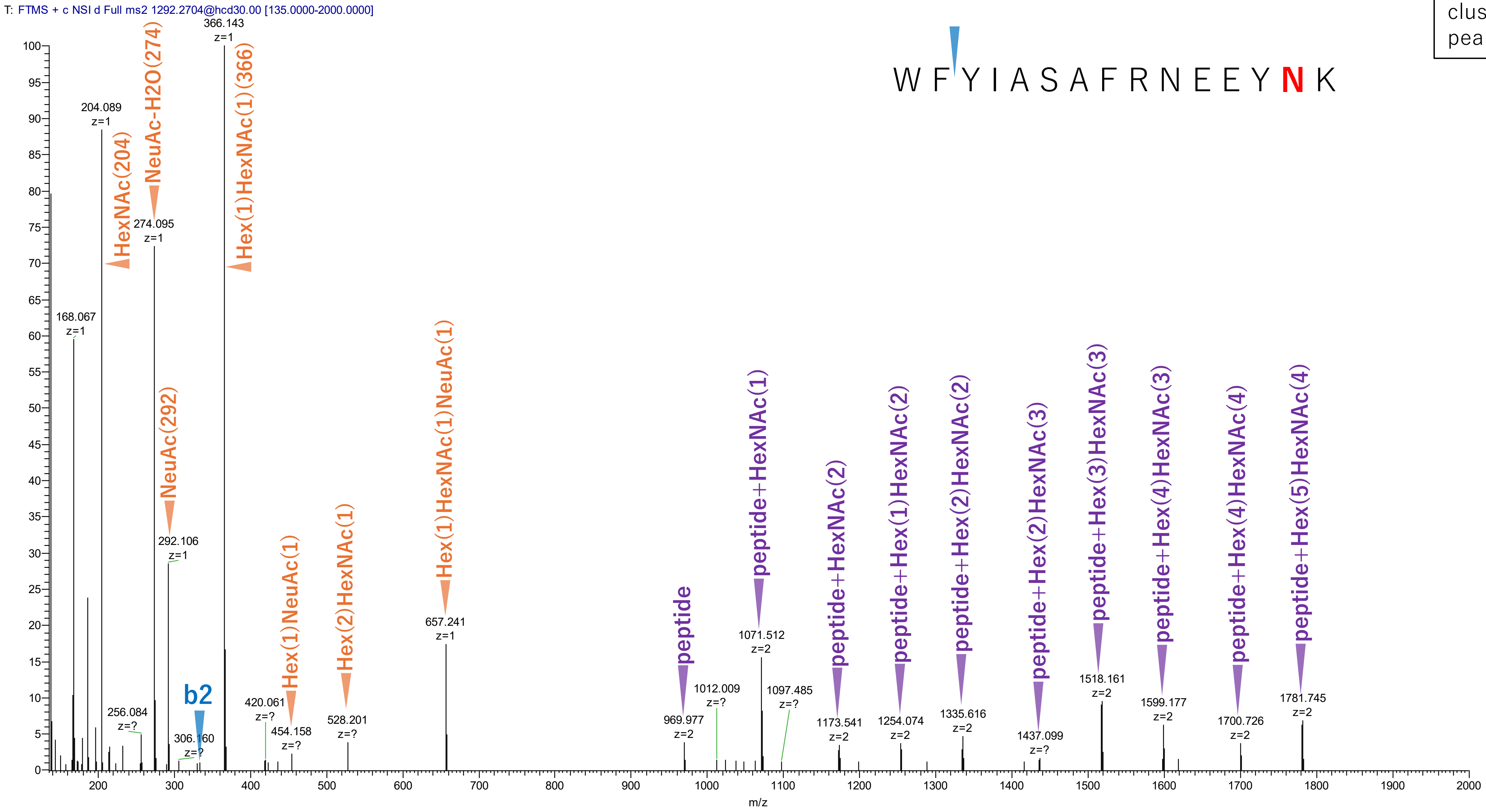


**Figure S2-14. MS2 spectra of glycopeptides assigned for hAGP.**

56(NKS)
43-57 WFYIASAFRNEEYNK + Hex(7)HexNAc(6)NeuAc(4)

cluster_no: 1
peak_no: 1249

W F Y I A S A F R N E E Y **N** K

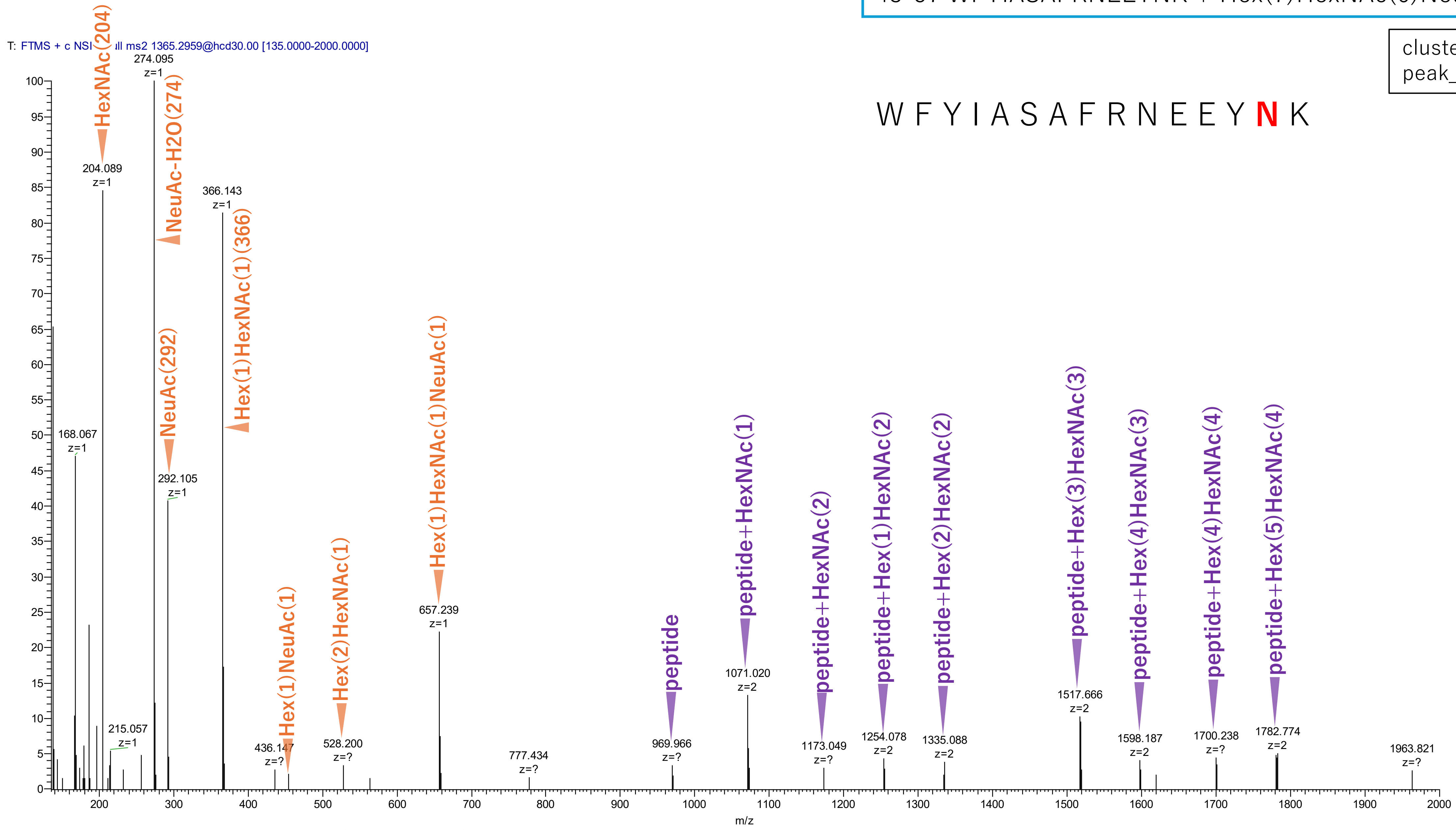


**Figure S2-15. MS2 spectra of glycopeptides assigned for hAGP.**

56(NKS)
43-57 WFYIASAFRNEEYNK + Hex(6)HexNAc(5)dHex(1)NeuAc(1)

cluster_no: 1
peak_no: 1293

W F Y I A S A F R N E E Y **N** K

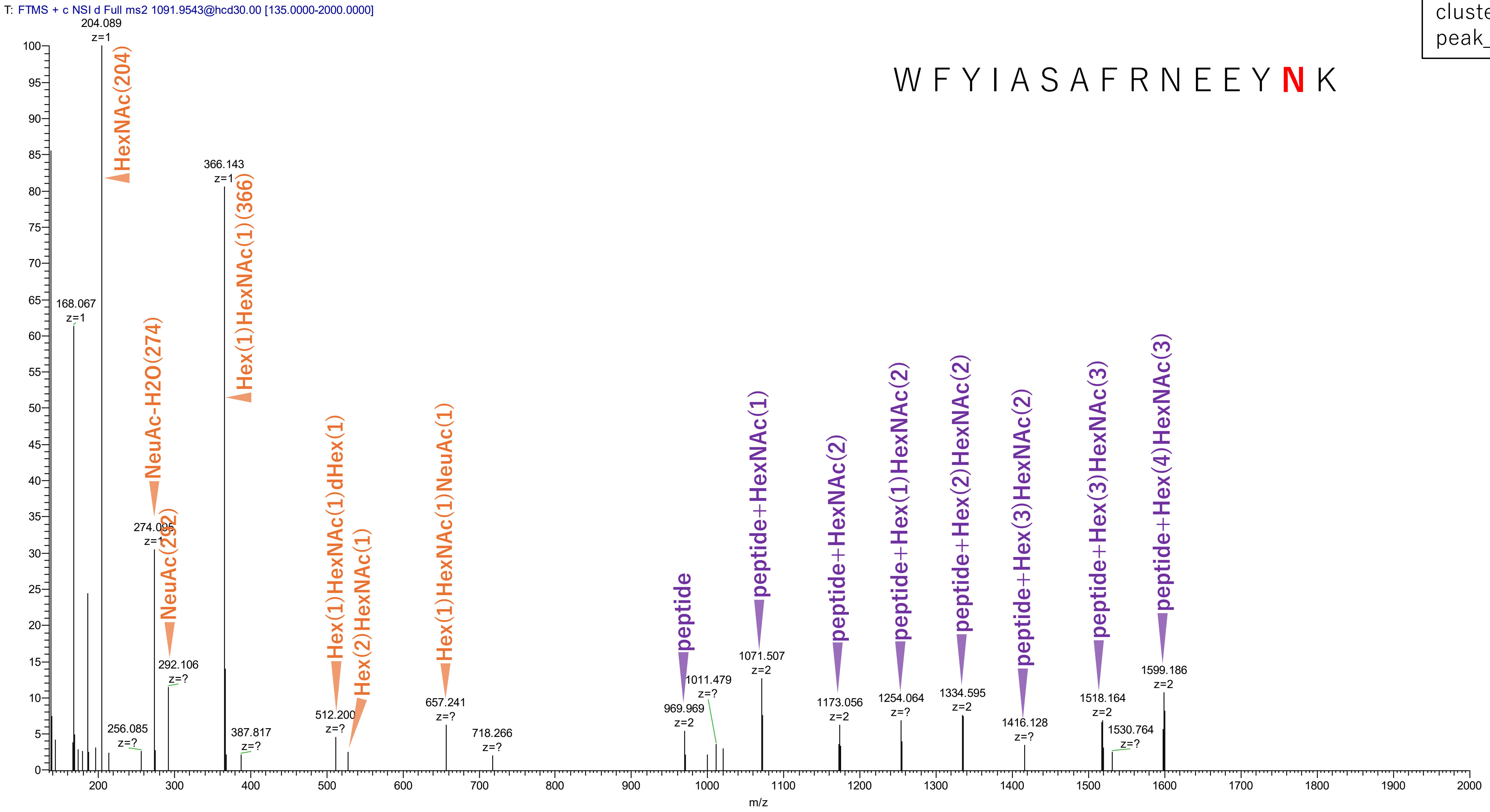


**Figure S2-16. MS2 spectra of glycopeptides assigned for hAGP.**

56(NKS)
43-57 WFYIASAFRNEEYNK + Hex(3)HexNAc(3)NeuAc(1)

cluster_no: 1
peak_no: 1477

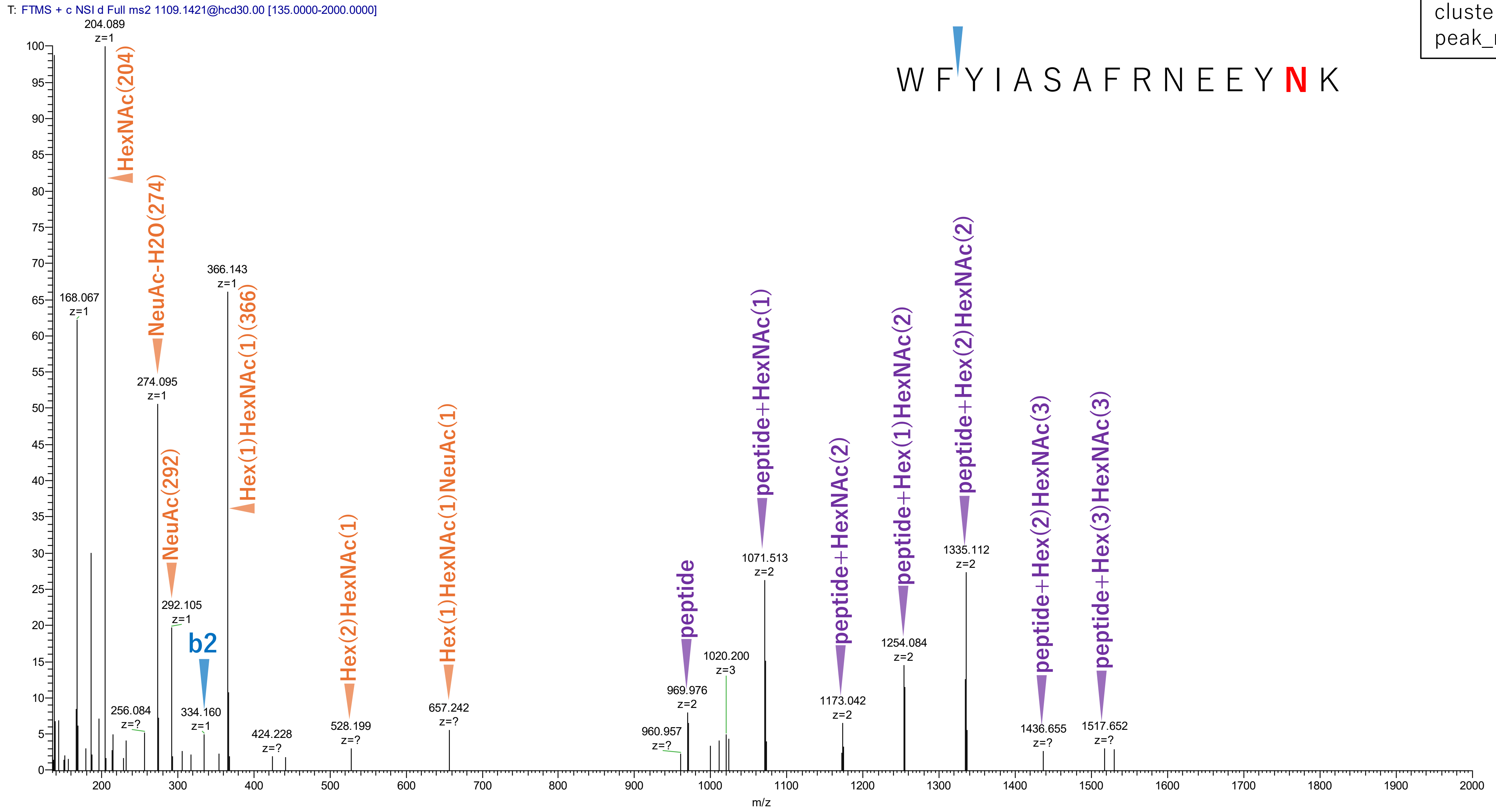


Figure S2-17. MS2 spectra of glycopeptides assigned for hAGP.

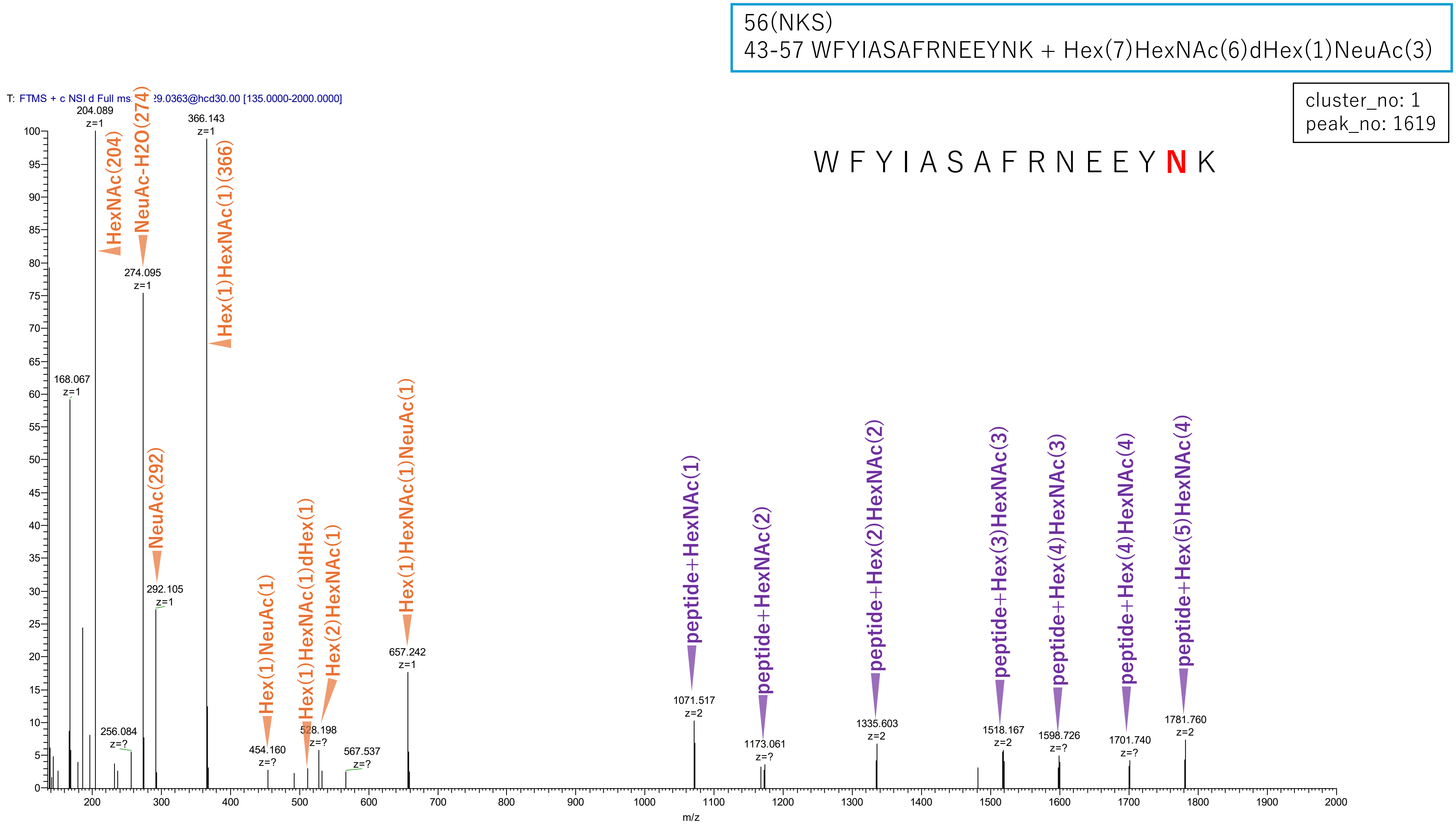


**Figure S2-18. MS2 spectra of glycopeptides assigned for hAGP.**

56(NKS)
43-57 WFYIASAFRNEEYNK + Hex(5)HexNAc(4)dHex(1)

cluster_no: 1
peak_no: 1860

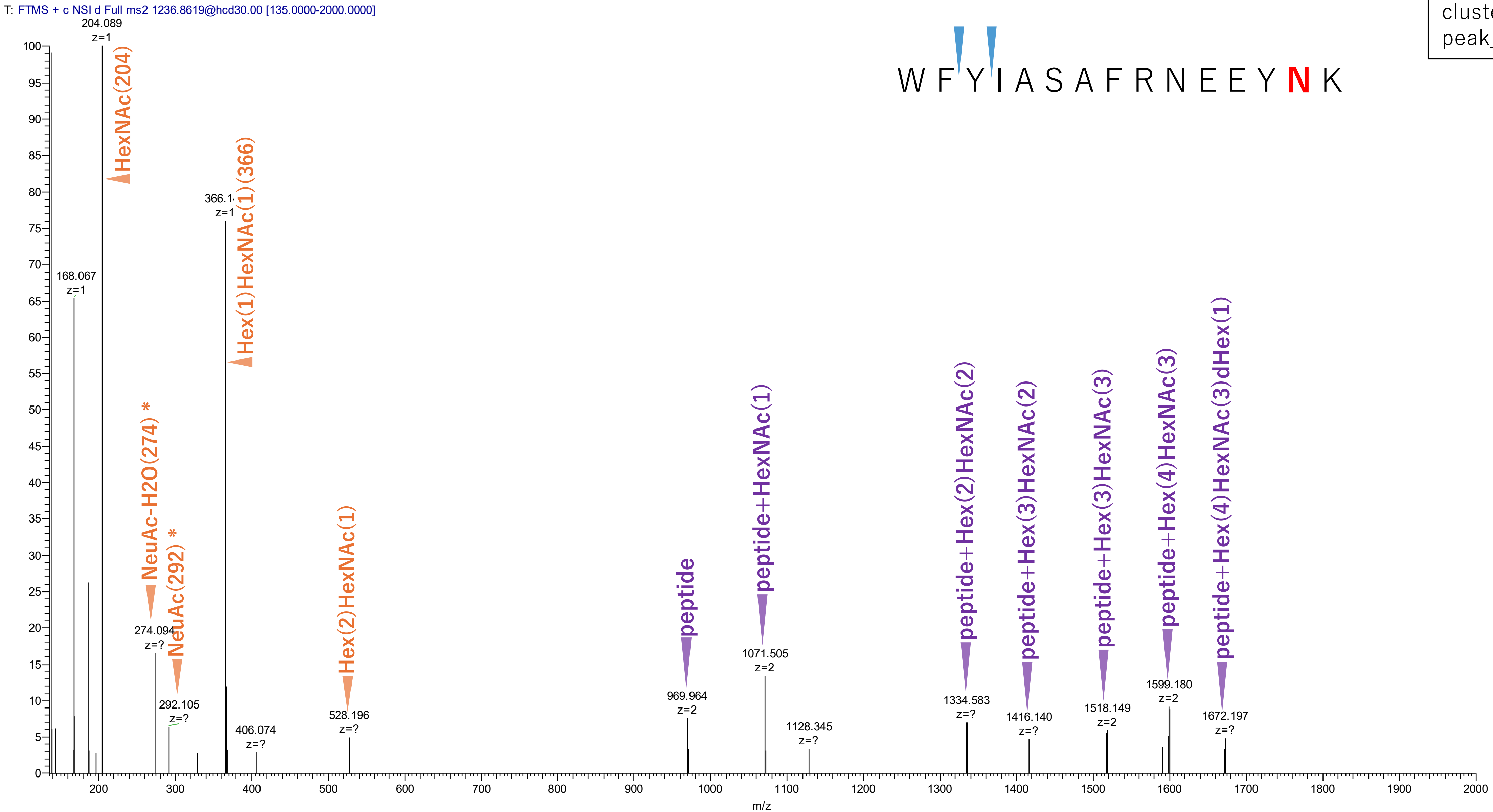


**Figure S2-19. MS2 spectra of glycopeptides assigned for hAGP.**

*Asterisks indicate diagnostic ions of NeuAc derived from a contaminated glycopeptide.

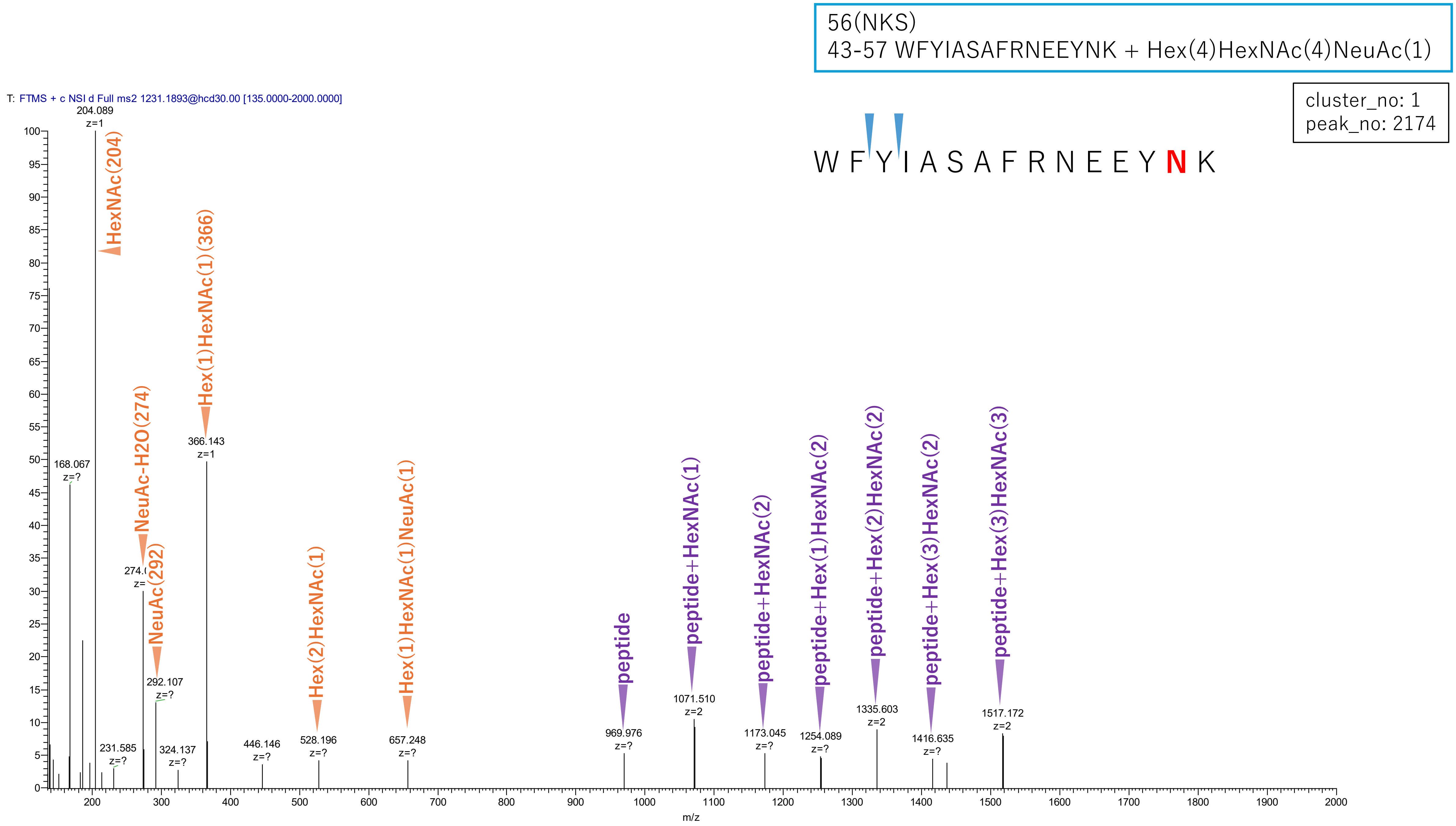


**Figure S2-20. MS2 spectra of glycopeptides assigned for hAGP.**

33(NAT)
19-42 QIPLCANLVPVPITNATLDRITGK + Hex(6)HexNAc(5)NeuAc(3)

cluster_no: 3
peak_no: 41

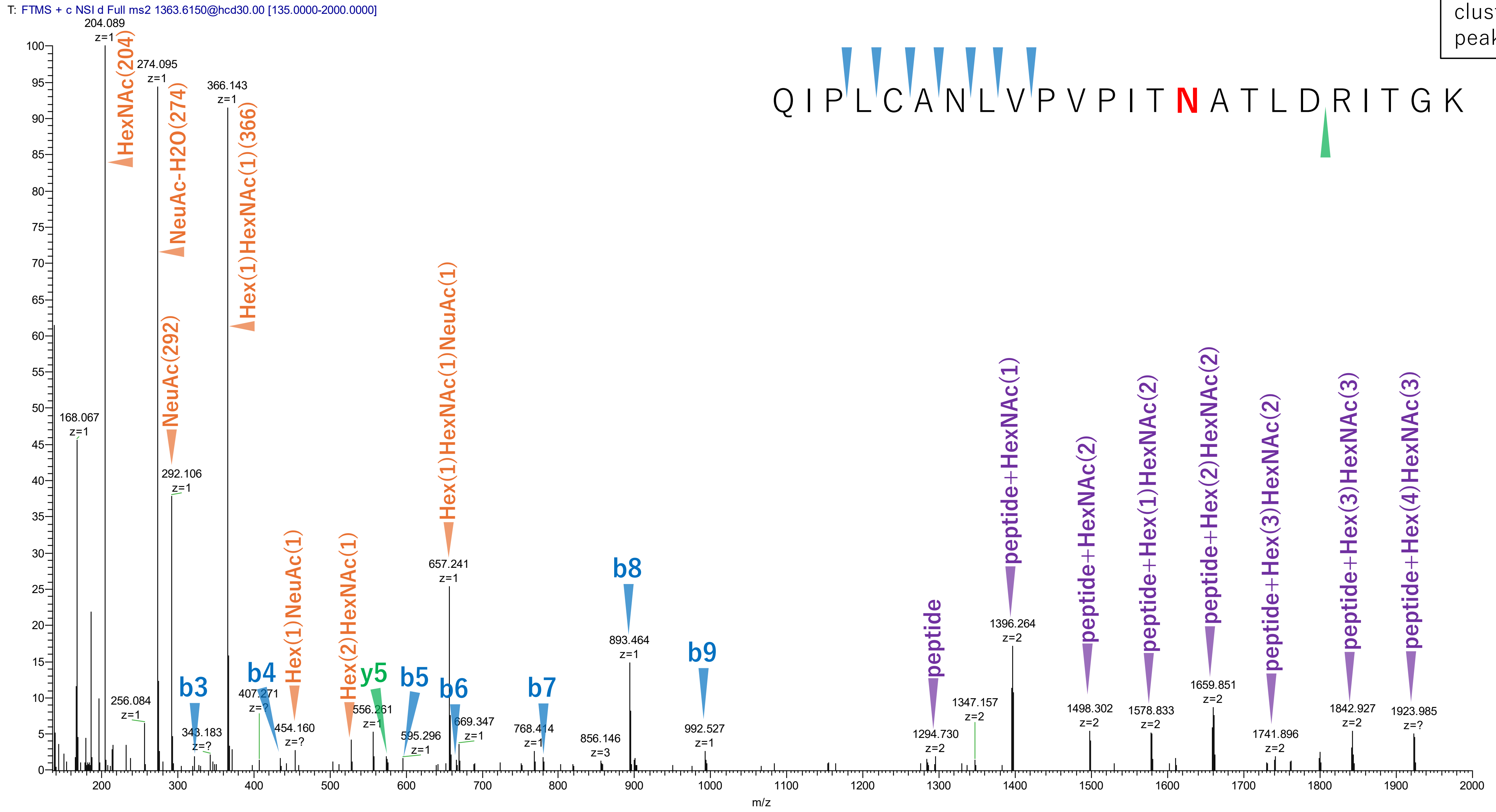


**Figure S2-21. MS2 spectra of glycopeptides assigned for hAGP.**

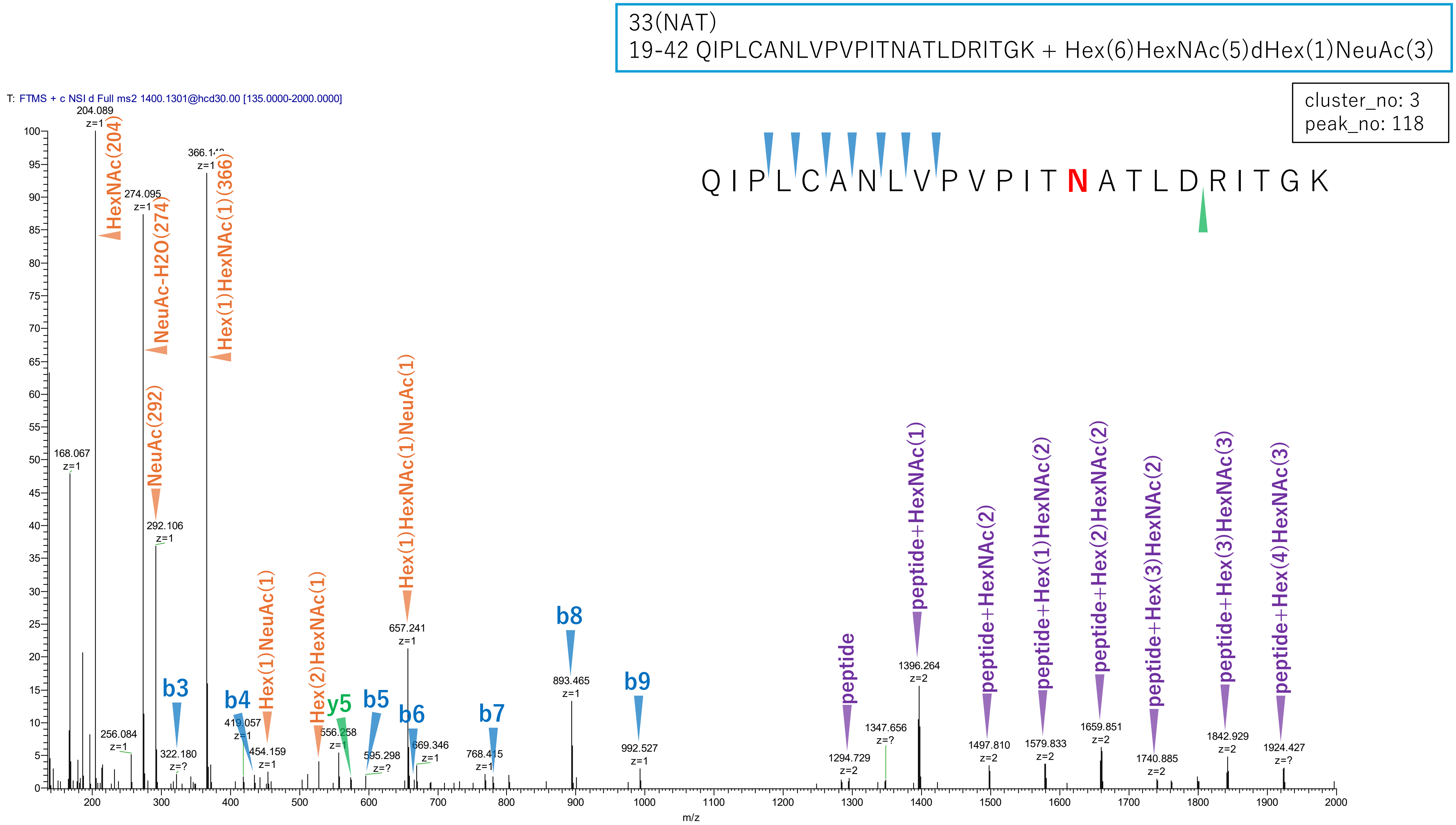


**Figure S2-22. MS2 spectra of glycopeptides assigned for hAGP.**

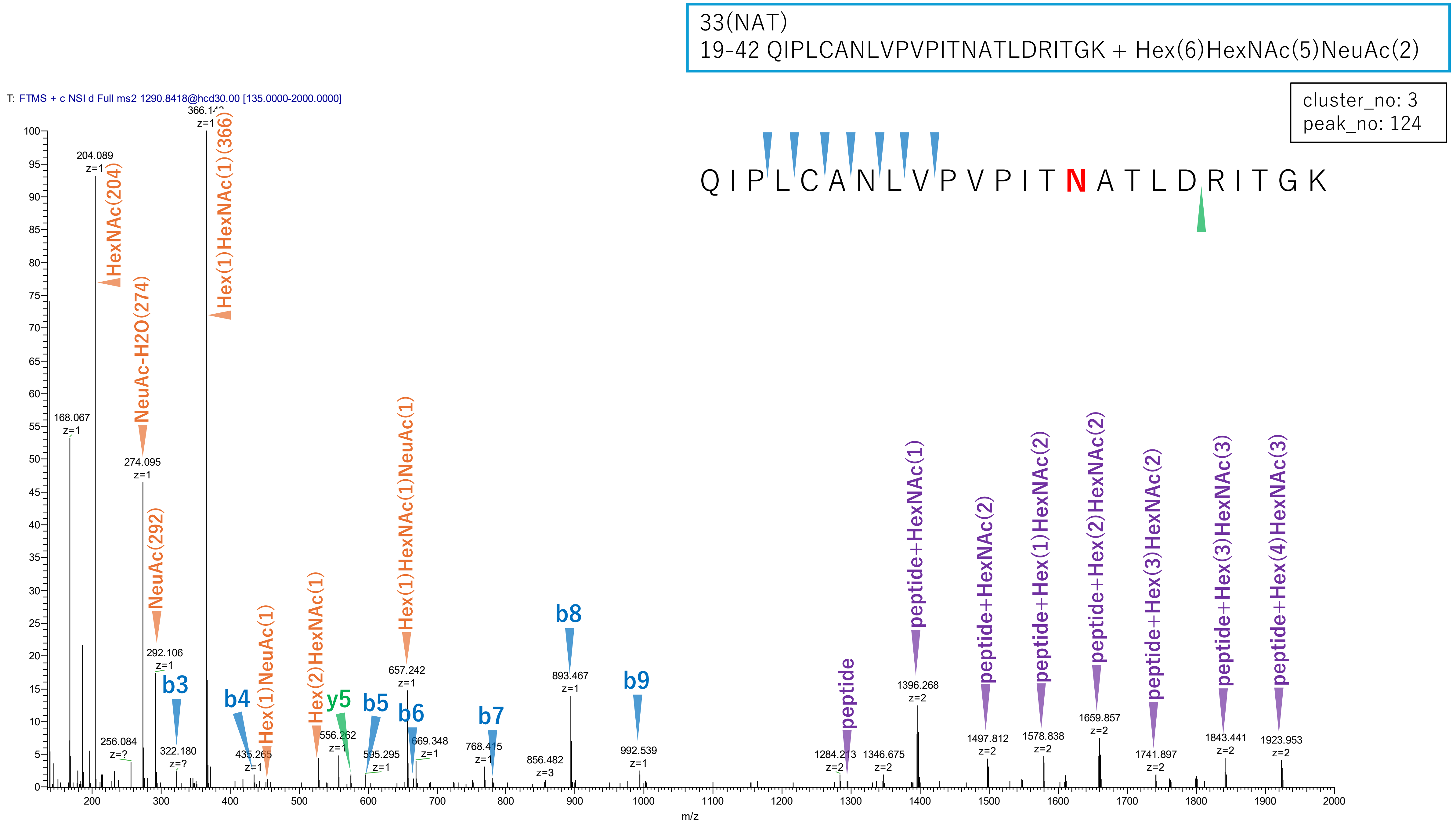


**Figure S2-23. MS2 spectra of glycopeptides assigned for hAGP.**

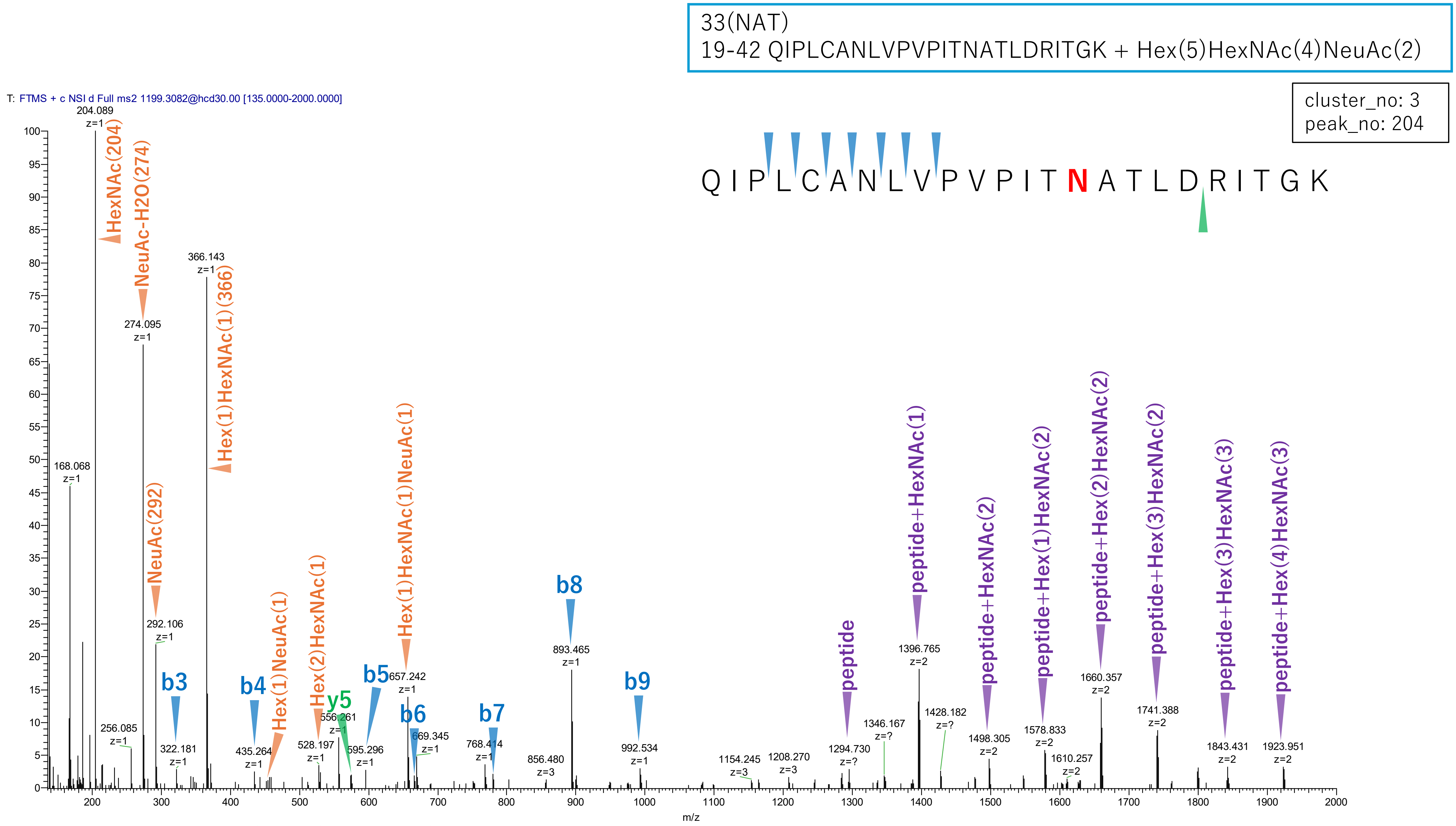


Figure S2-24. MS2 spectra of glycopeptides assigned for hAGP.

33(NAT)
19-42 QIPLCANLVPVPITNATLDRITGK + Hex(6)HexNAc(5)

cluster_no: 3
peak_no: 229

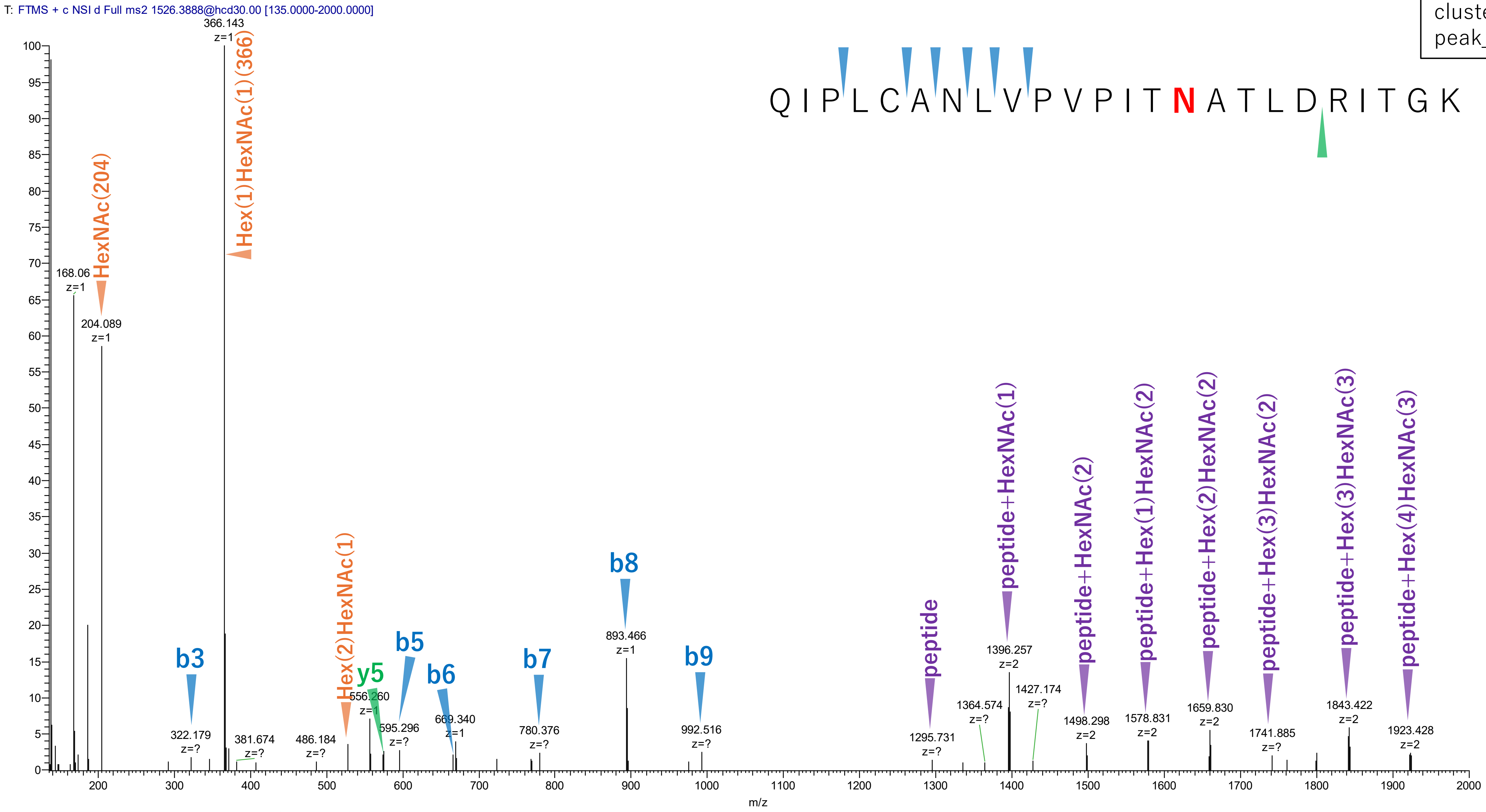


Figure S2-25. MS2 spectra of glycopeptides assigned for hAGP.

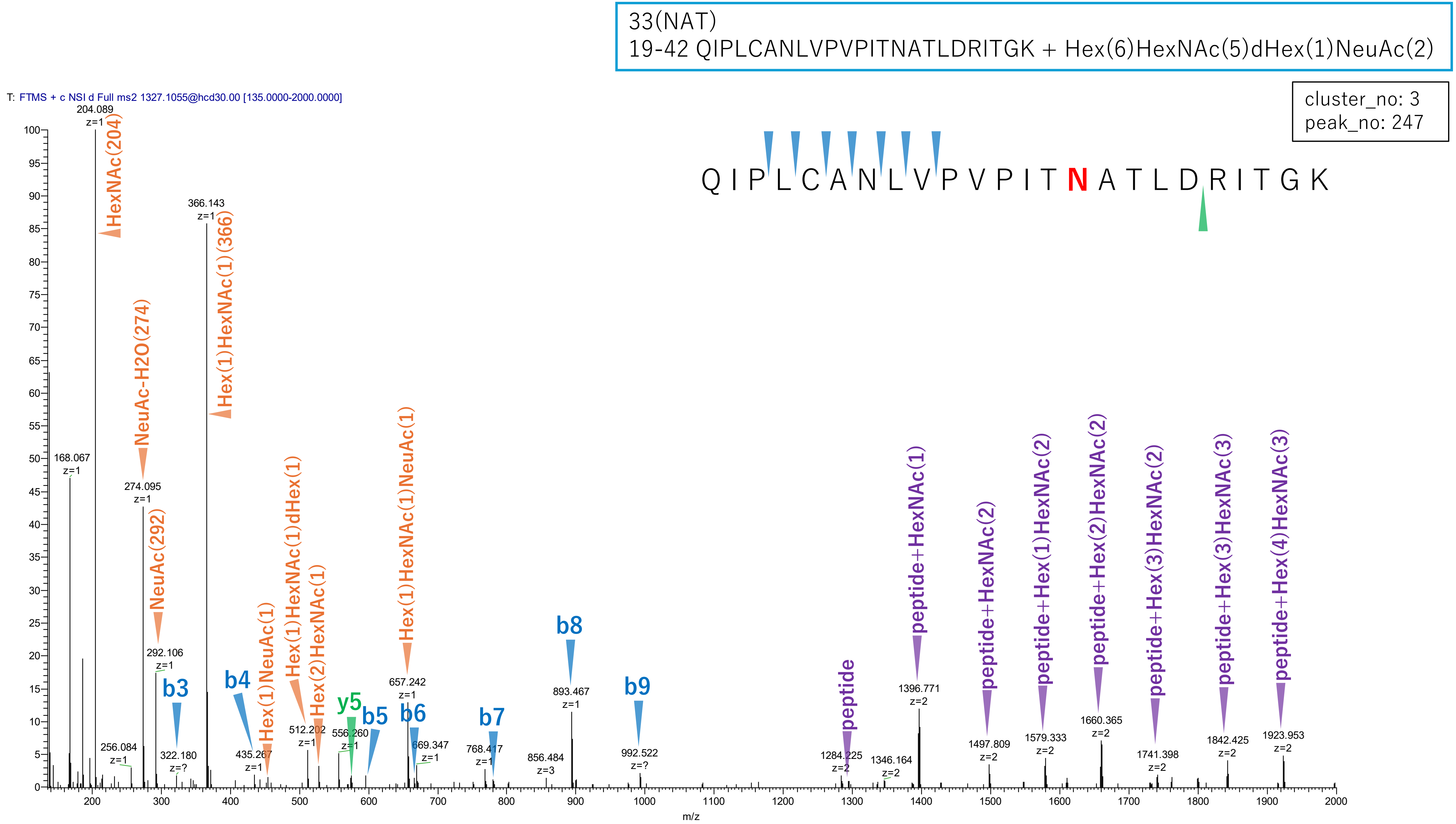


**Figure S2-26. MS2 spectra of glycopeptides assigned for hAGP.**

33(NAT)
19-42 QIPLCANLVPVPITNATLDRITGK + Hex(6)HexNAc(5)NeuAc(1)

cluster_no: 3
peak_no: 413

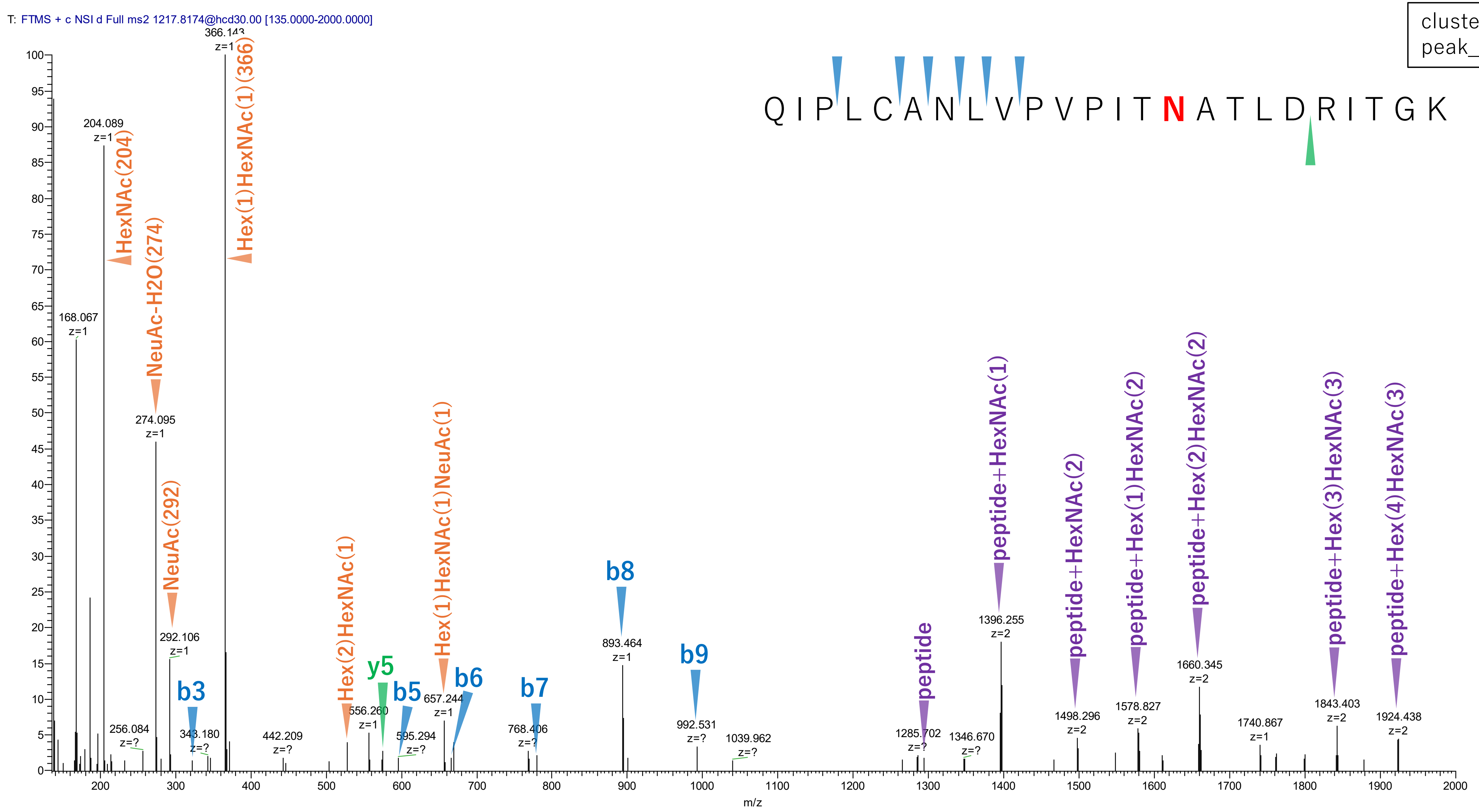


Figure S2-27. MS2 spectra of glycopeptides assigned for hAGP.

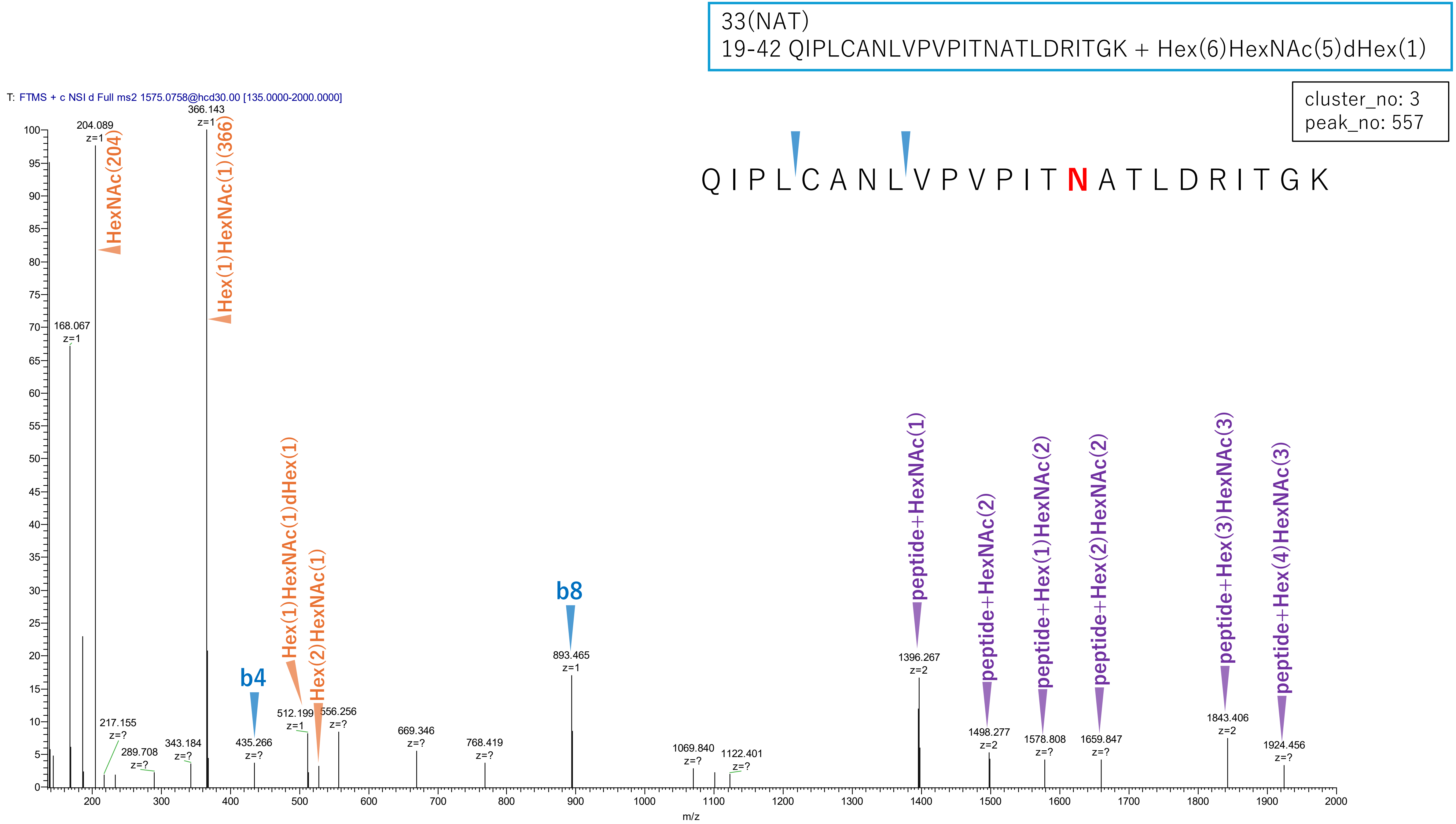


**Figure S2-28. MS2 spectra of glycopeptides assigned for hAGP.**

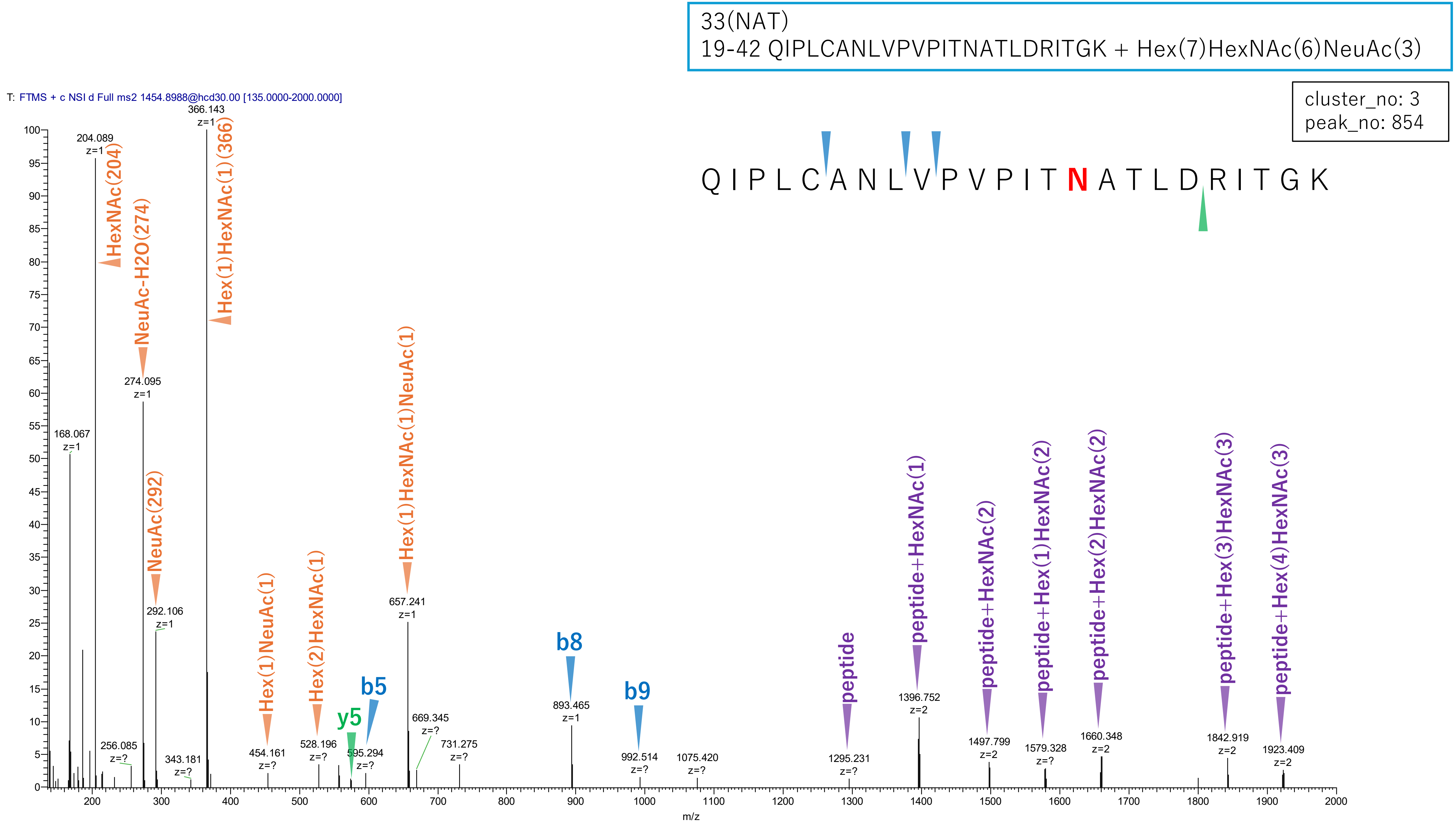


**Figure S2-29. MS2 spectra of glycopeptides assigned for hAGP.**

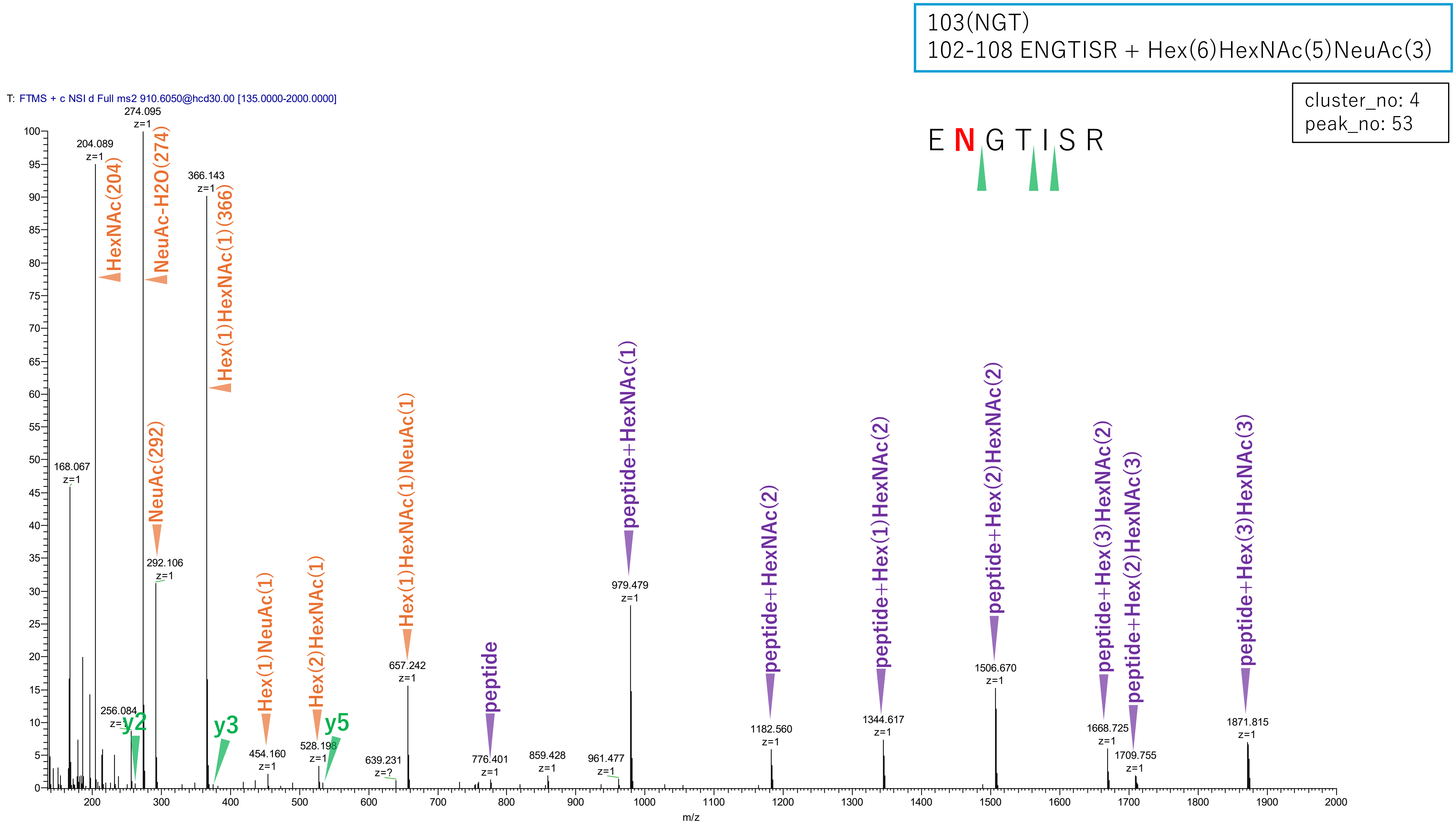


**Figure S2-30. MS2 spectra of glycopeptides assigned for hAGP.**

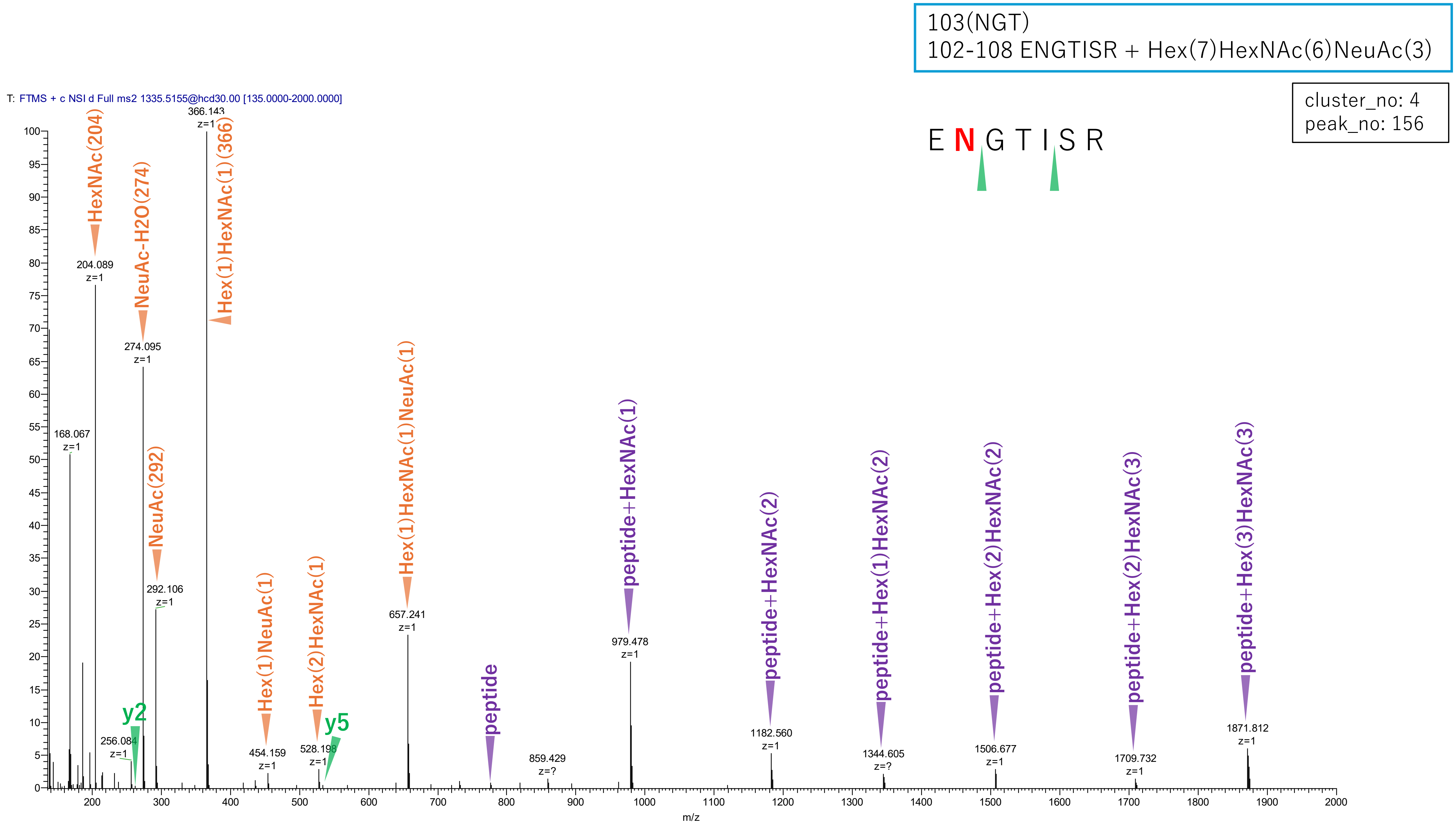


**Figure S2-31. MS2 spectra of glycopeptides assigned for hAGP.**

103(NGT)
102-108 ENGTISR + Hex(6)HexNAc(5)dHex(1)NeuAc(3)

cluster_no: 4
peak_no: 158

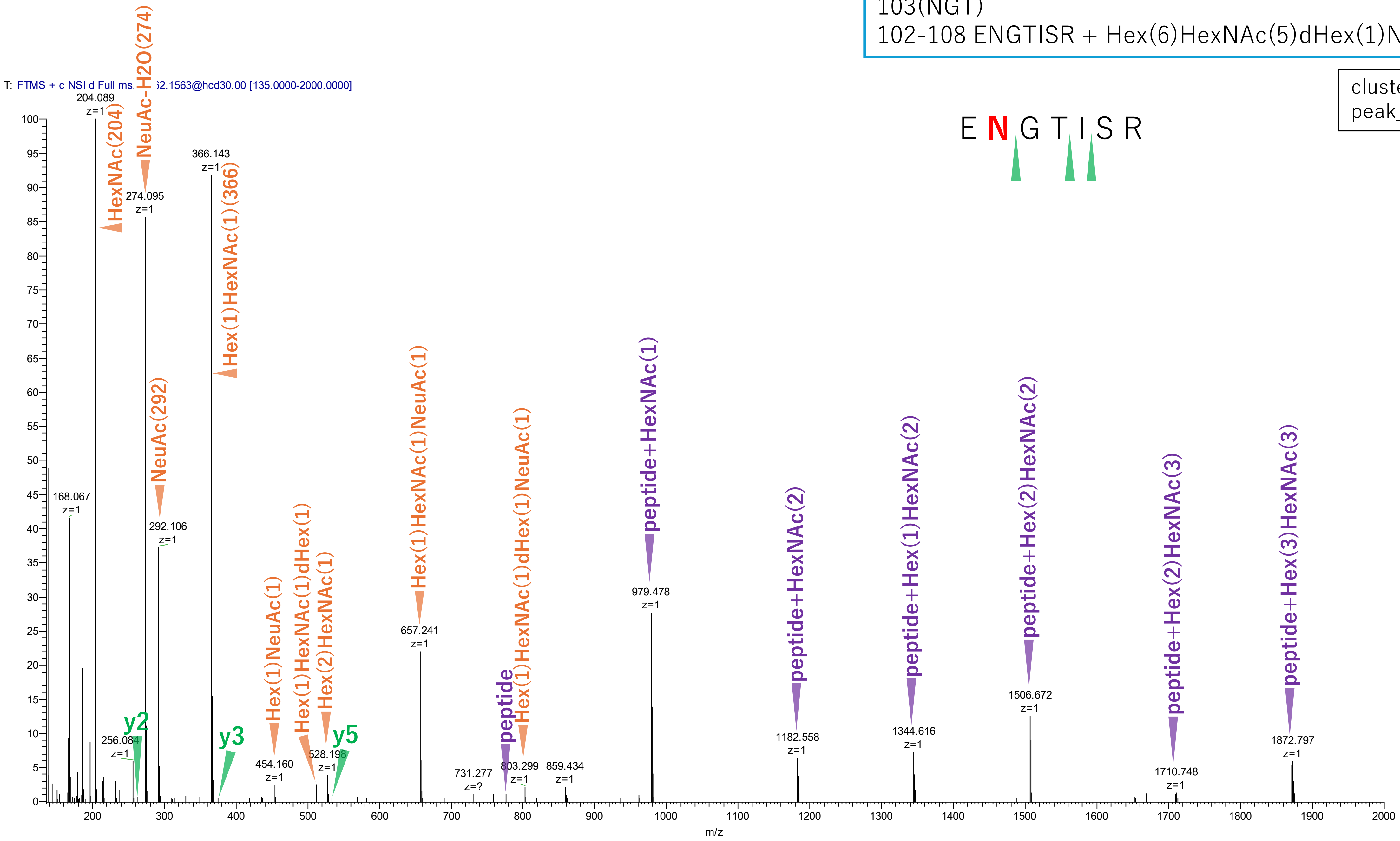


Figure S2-32. MS2 spectra of glycopeptides assigned for hAGP.

103(NGT)
102-108 ENGTISR + Hex(7)HexNAc(6)NeuAc(4)

cluster_no: 4
peak_no: 165

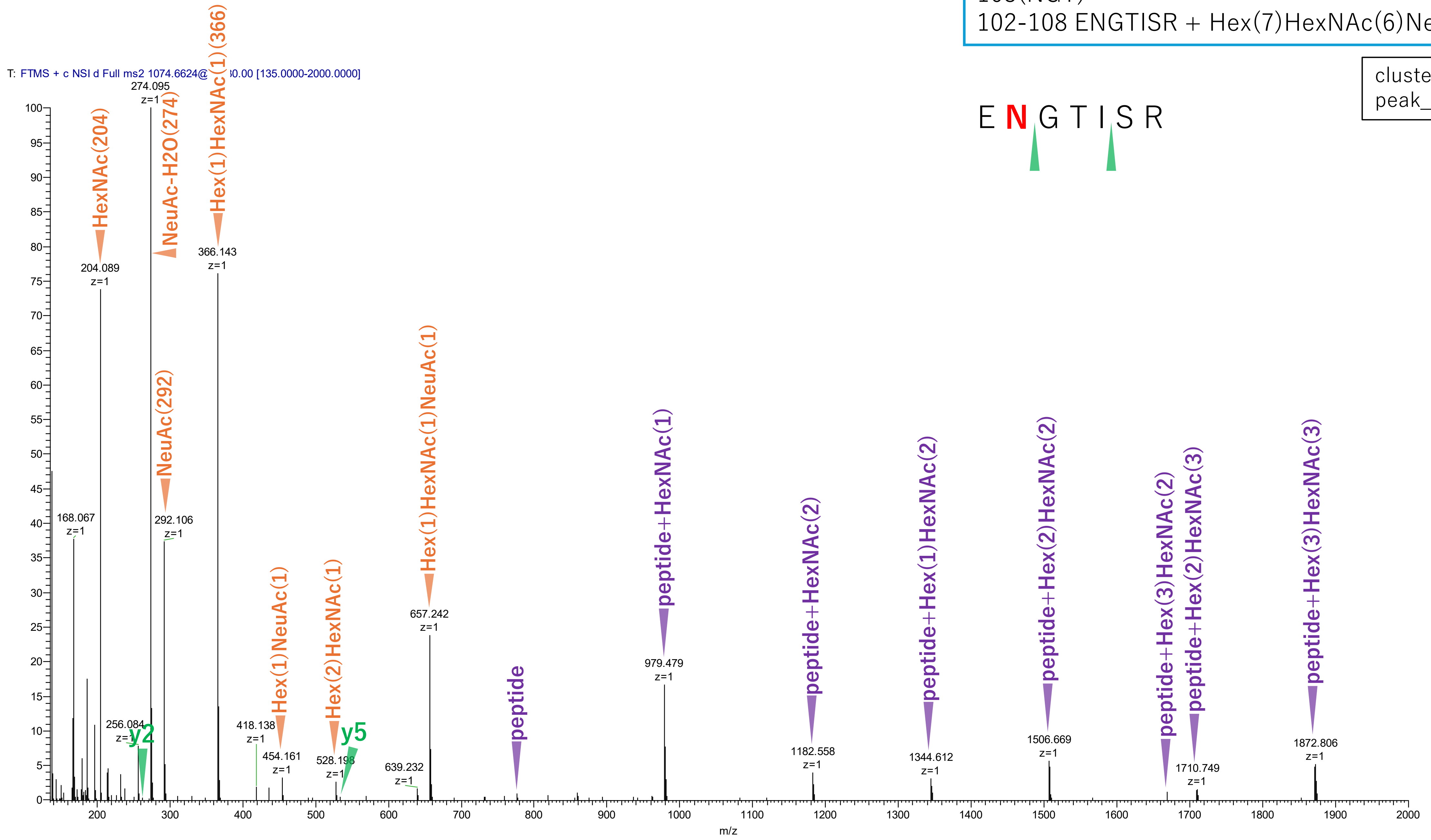


**Figure S2-33. MS2 spectra of glycopeptides assigned for hAGP.**

103(NGT)
102-108 ENGTISR + Hex(6)HexNAc(5)NeuAc(2)

cluster_no: 4
peak_no: 216

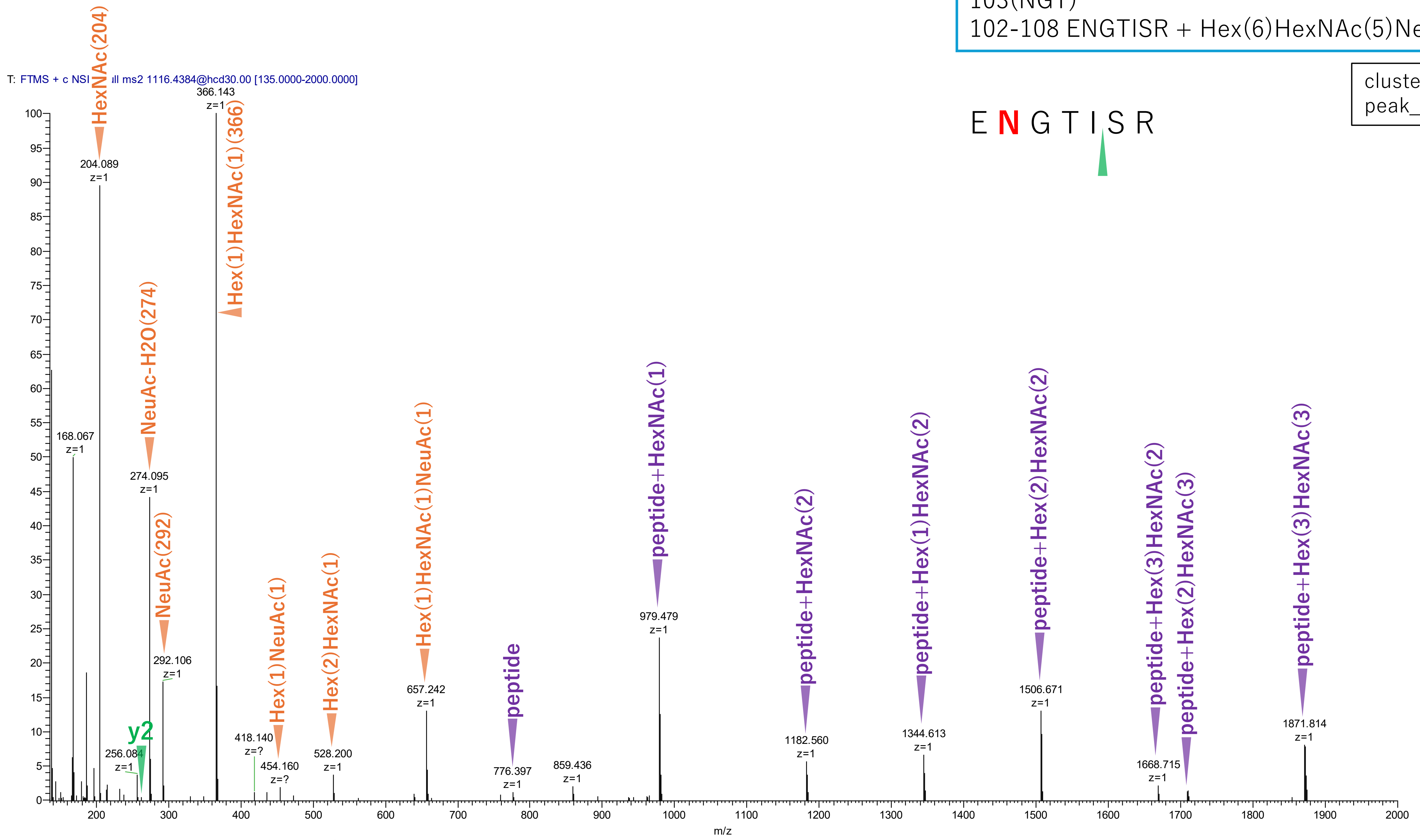


**Figure S2-34. MS2 spectra of glycopeptides assigned for hAGP.**

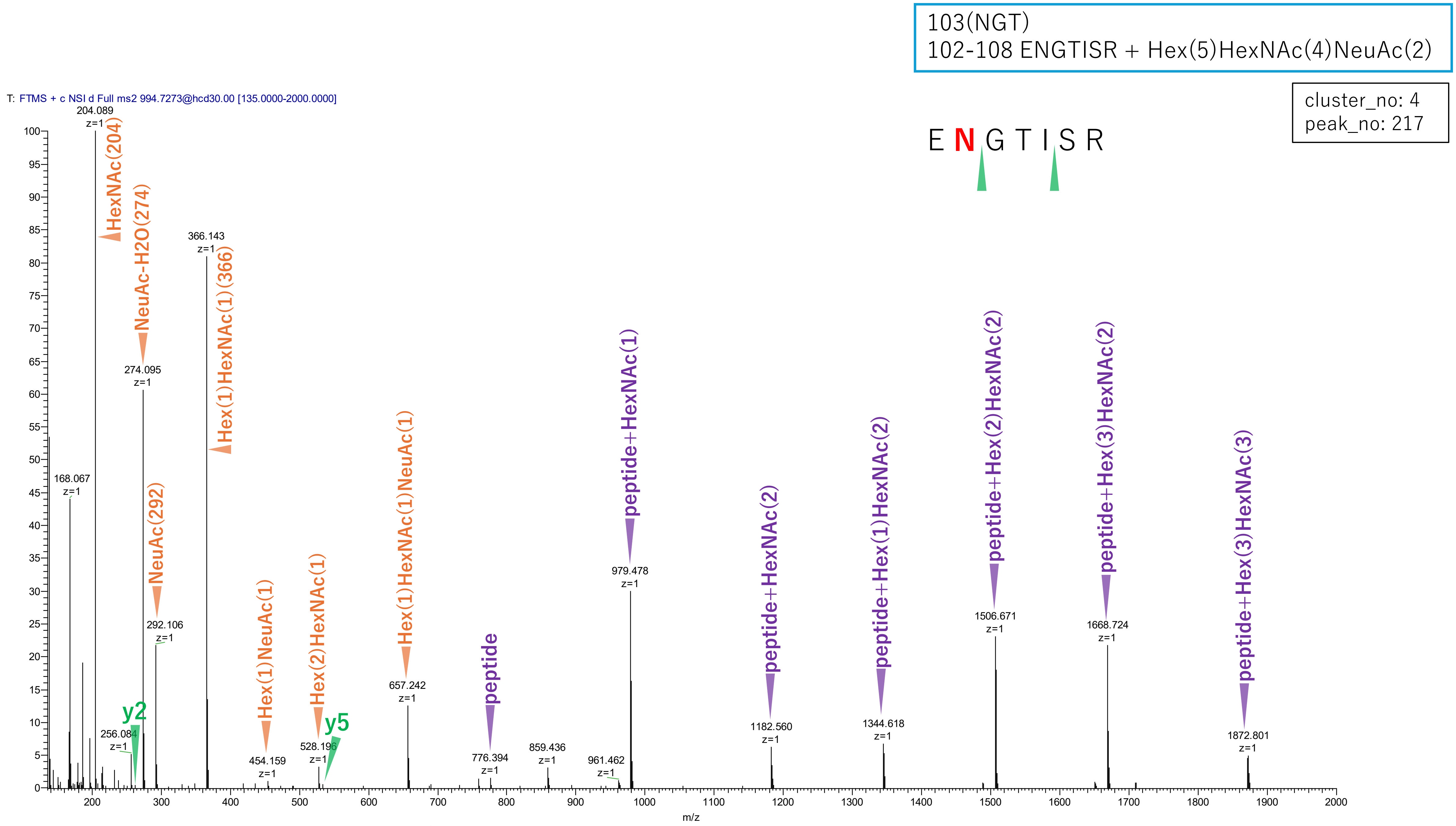


**Figure S2-35. MS2 spectra of glycopeptides assigned for hAGP.**

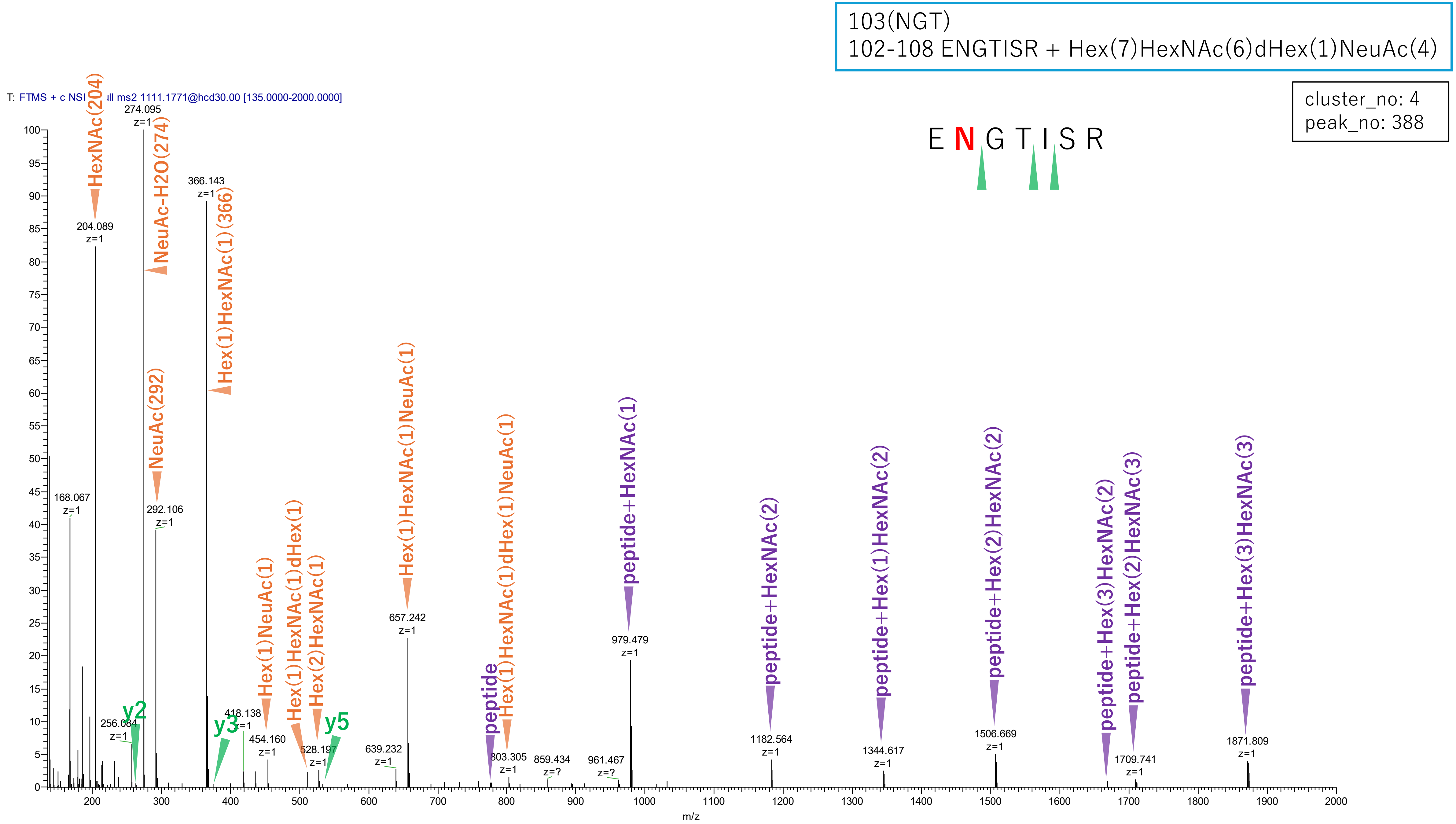


**Figure S2-36. MS2 spectra of glycopeptides assigned for hAGP.**

103(NGT)
102-108 ENGTISR + Hex(7)HexNAc(6)dHex(1)NeuAc(3)

cluster_no: 4
peak_no: 398

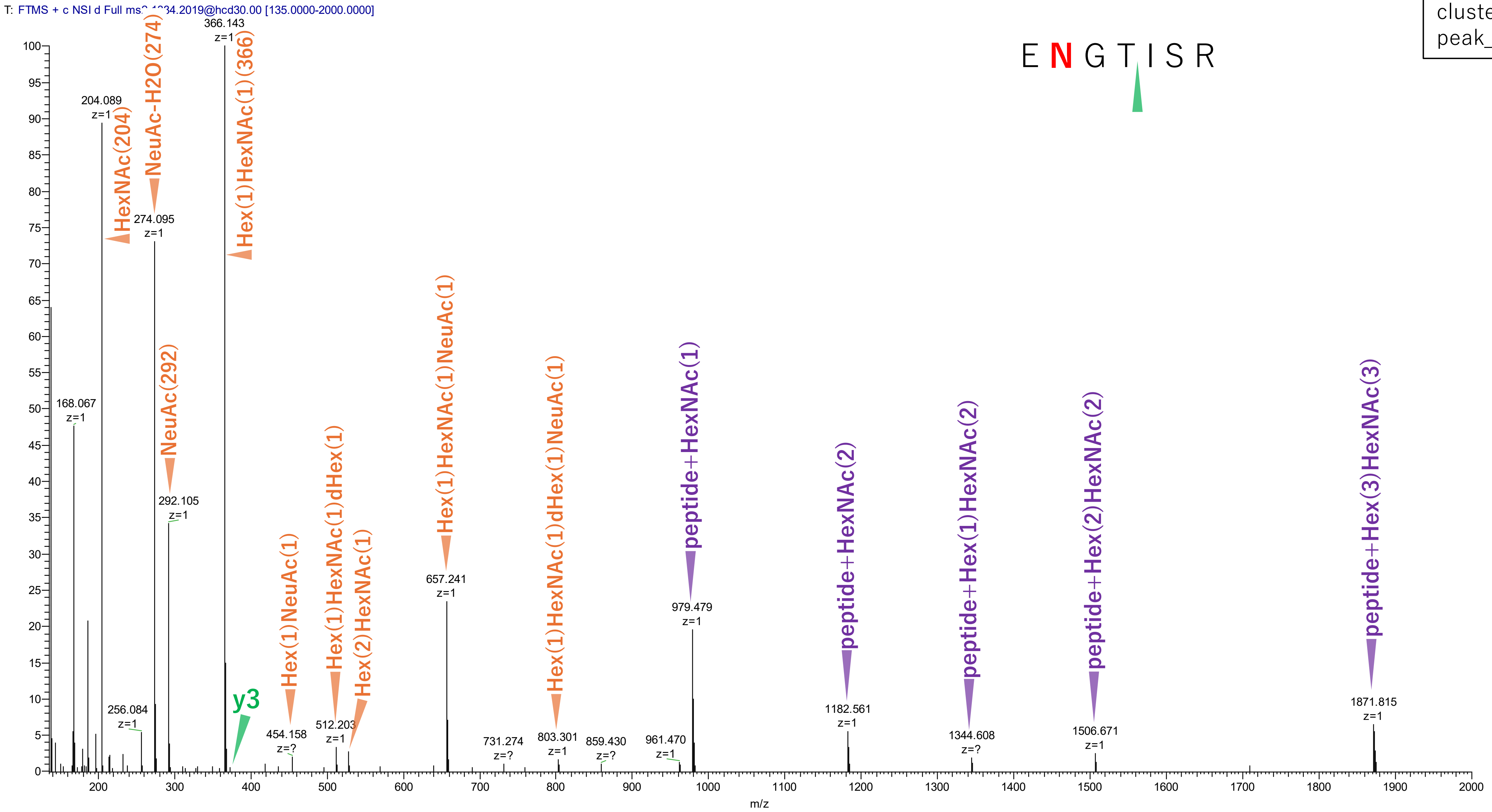


**Figure S2-37. MS2 spectra of glycopeptides assigned for hAGP.**

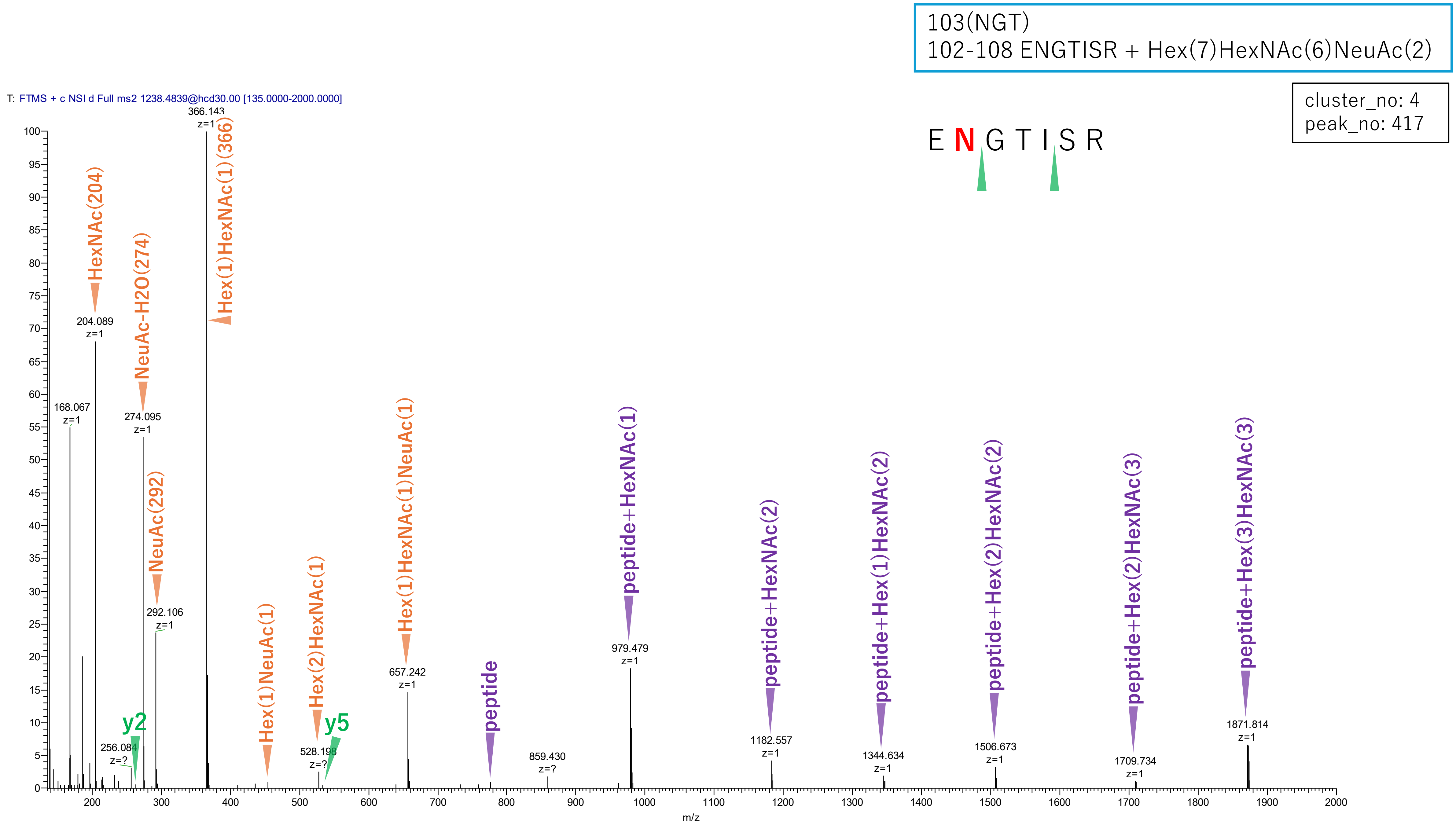


**Figure S2-38. MS2 spectra of glycopeptides assigned for hAGP.**

103(NGT)
102-108 ENGTISR + Hex(6)HexNAc(5)dHex(1)NeuAc(2)

cluster_no: 4
peak_no: 475

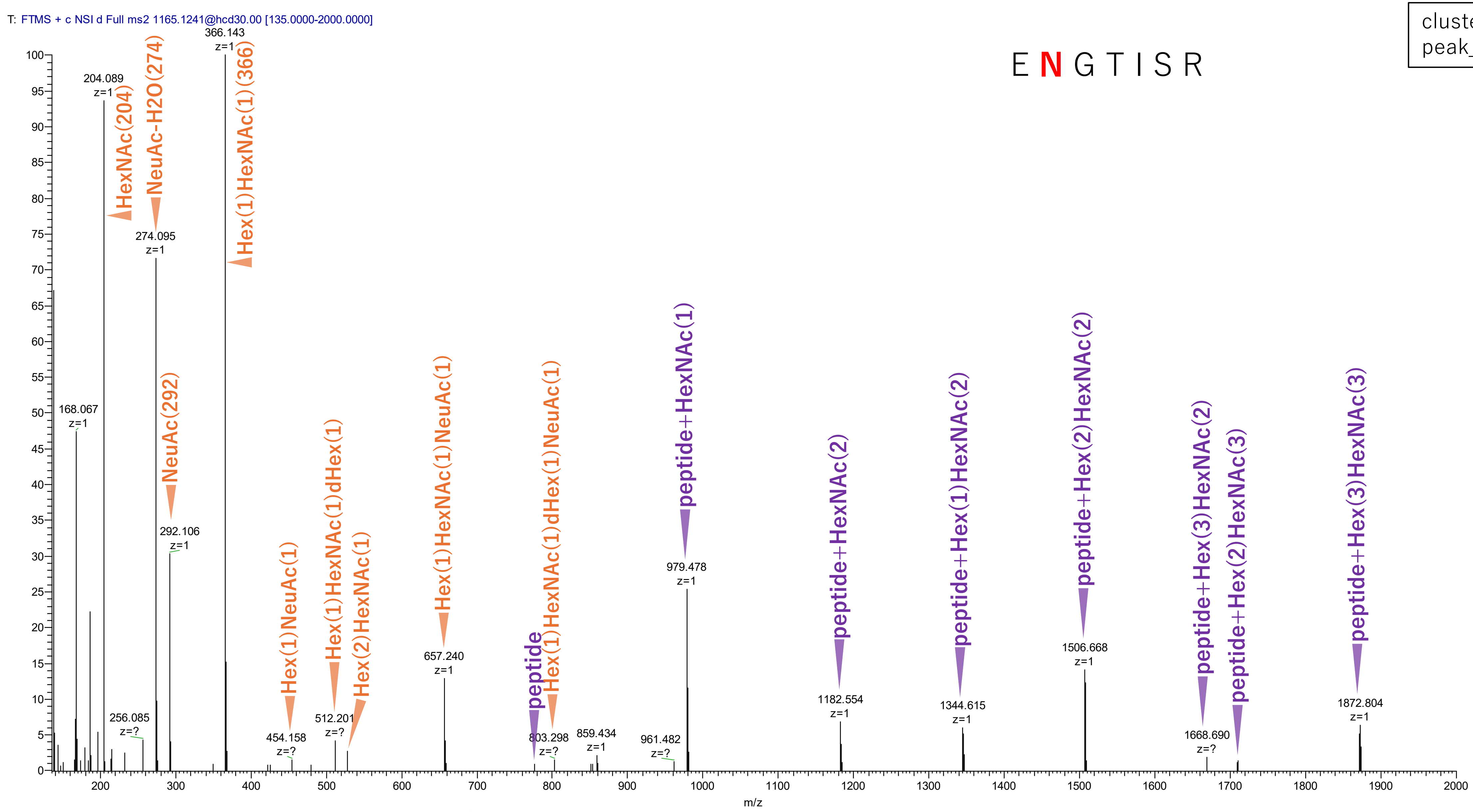


**Figure S2-39. MS2 spectra of glycopeptides assigned for hAGP.**

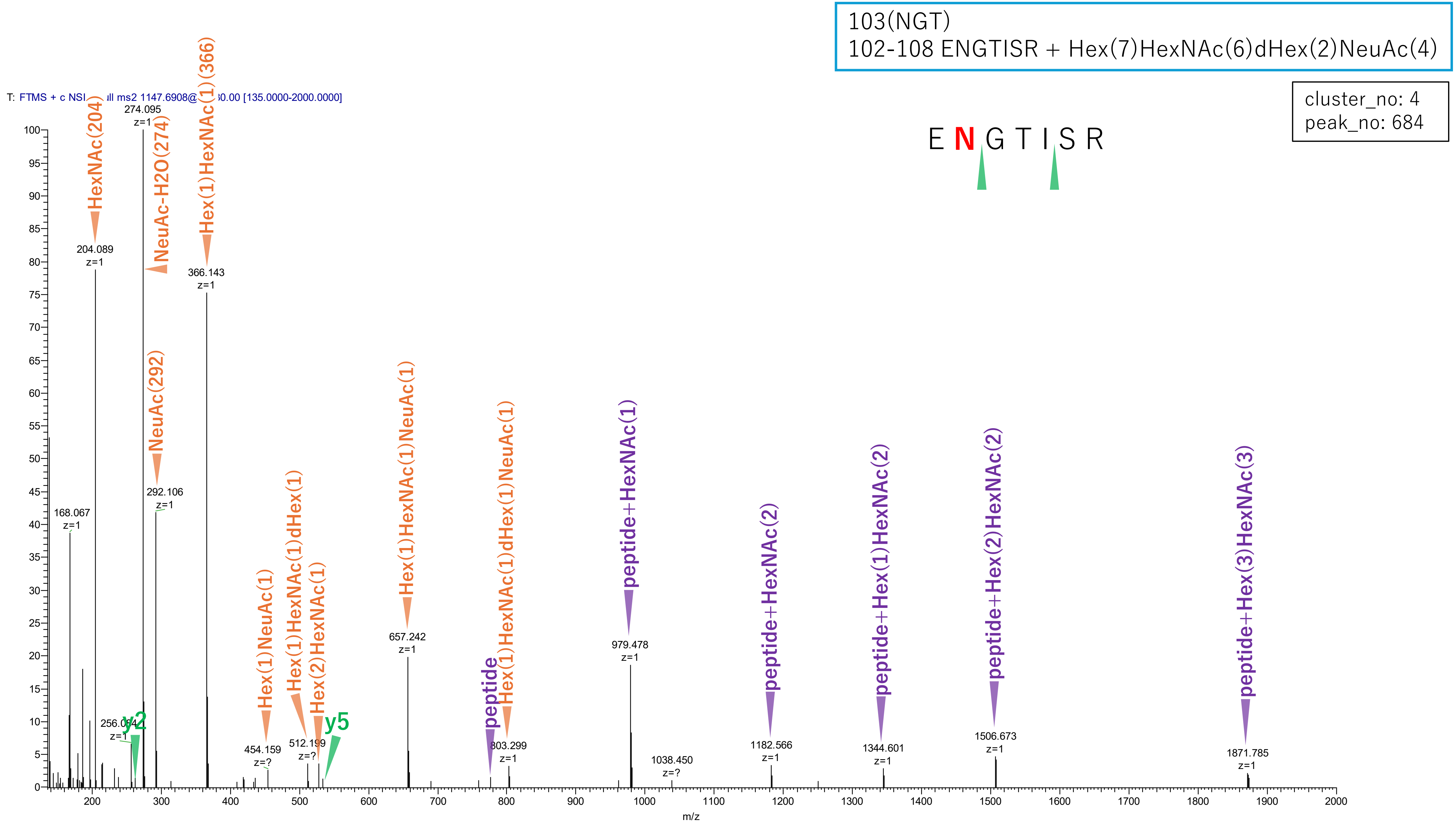


**Figure S2-40. MS2 spectra of glycopeptides assigned for hAGP.**

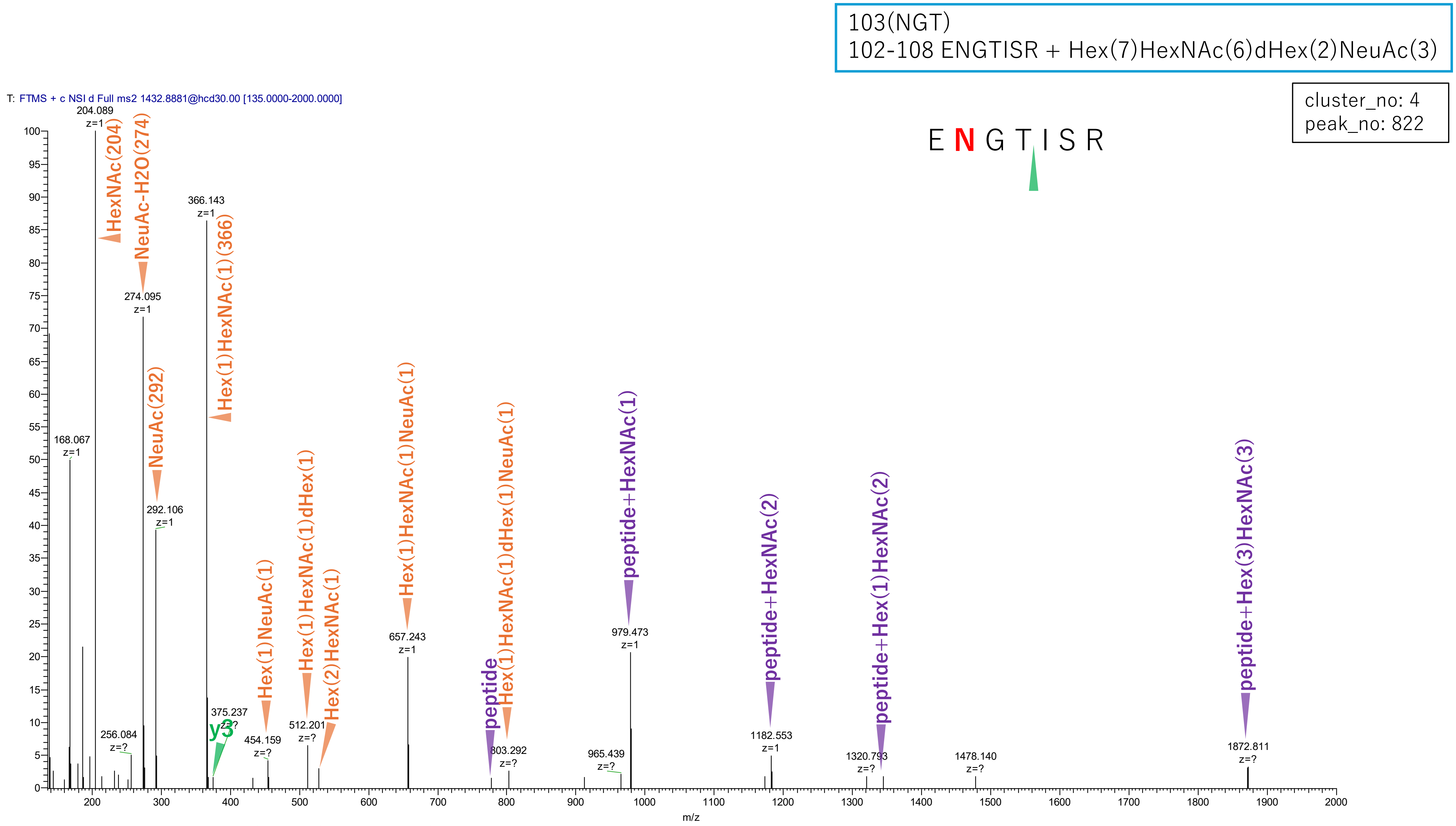


**Figure S2-41. MS2 spectra of glycopeptides assigned for hAGP.**

103(NGT)
102-108 ENGTISR + Hex(8)HexNAc(7)NeuAc(4)

cluster_no: 4
peak_no: 851

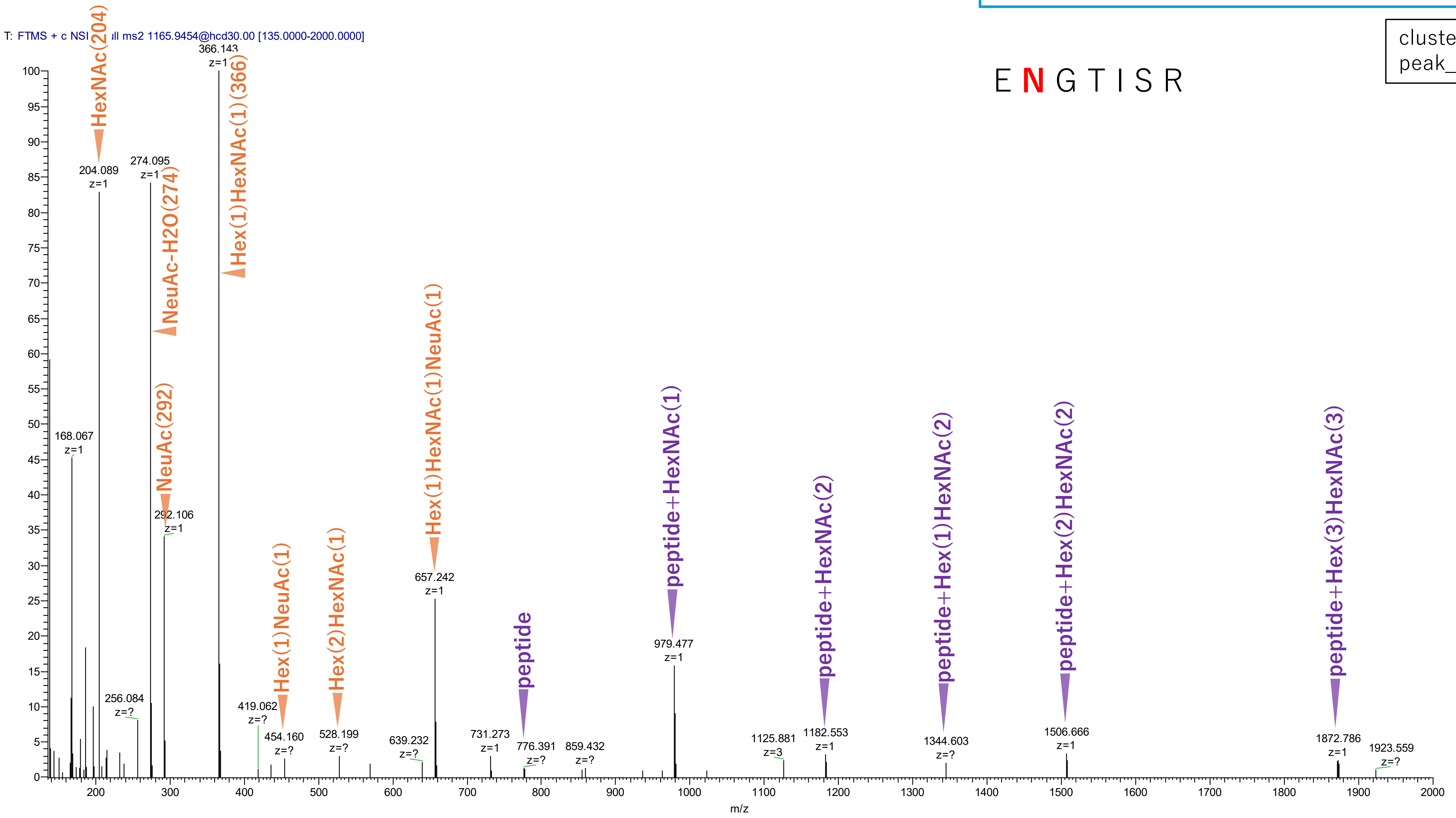

Figure S2-42. MS2 spectra of glycopeptides assigned for hAGP.

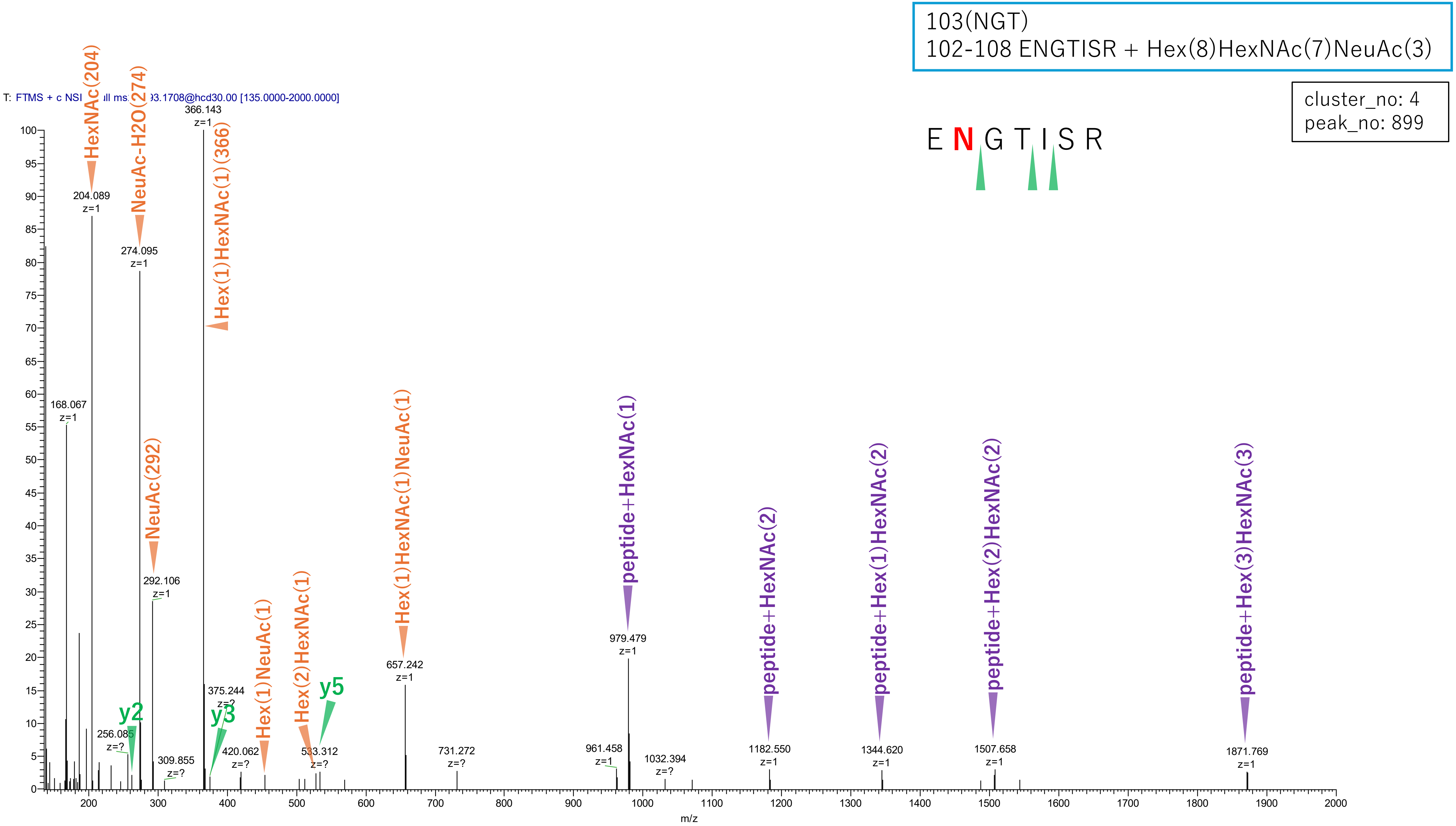


**Figure S2-43. MS2 spectra of glycopeptides assigned for hAGP.**

103(NGT)
102-108 ENGTISR + Hex(7)HexNAc(6)dHex(1)NeuAc(2)

cluster_no: 4
peak_no: 995

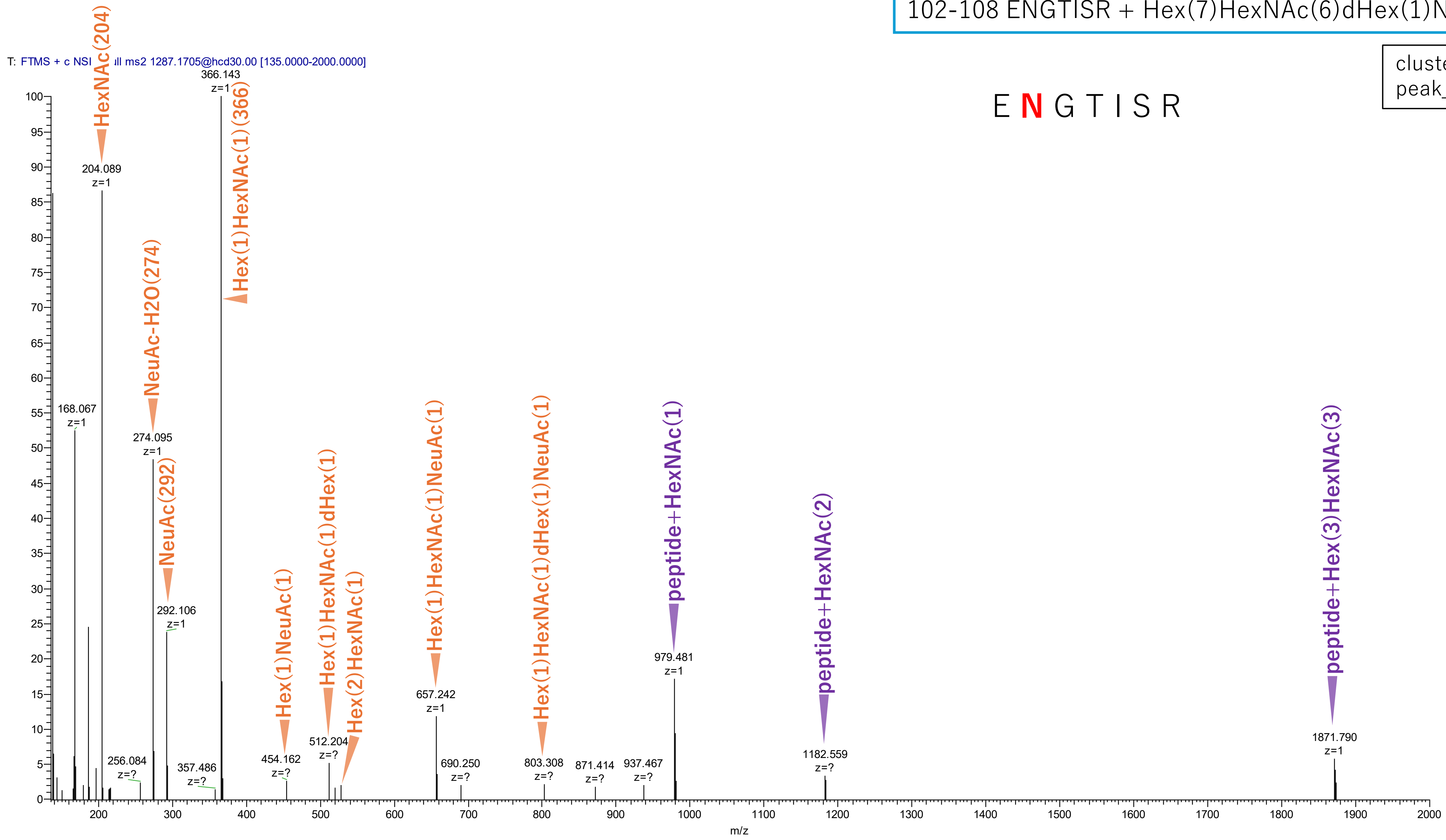


**Figure S2-44. MS2 spectra of glycopeptides assigned for hAGP.**

103(NGT)
102-108 ENGTISR + Hex(6)HexNAc(5)dHex(2)NeuAc(3)

cluster_no: 4
peak_no: 1118

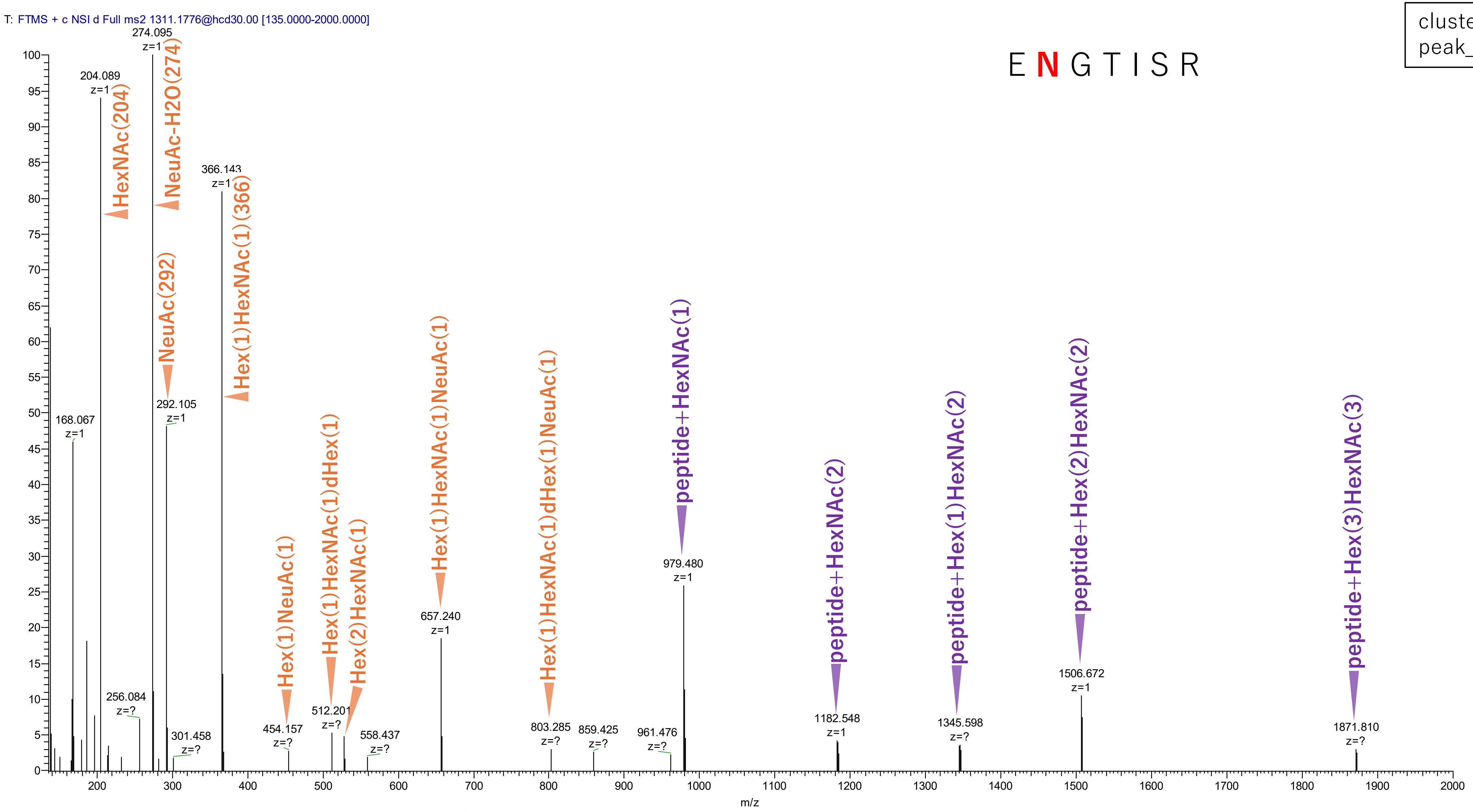


**Figure S2-45. MS2 spectra of glycopeptides assigned for hAGP.**

103(NGT)
102-108 ENGTISR + Hex(7)HexNAc(6)dHex(3)NeuAc(4)

cluster_no: 4
peak_no: 1141

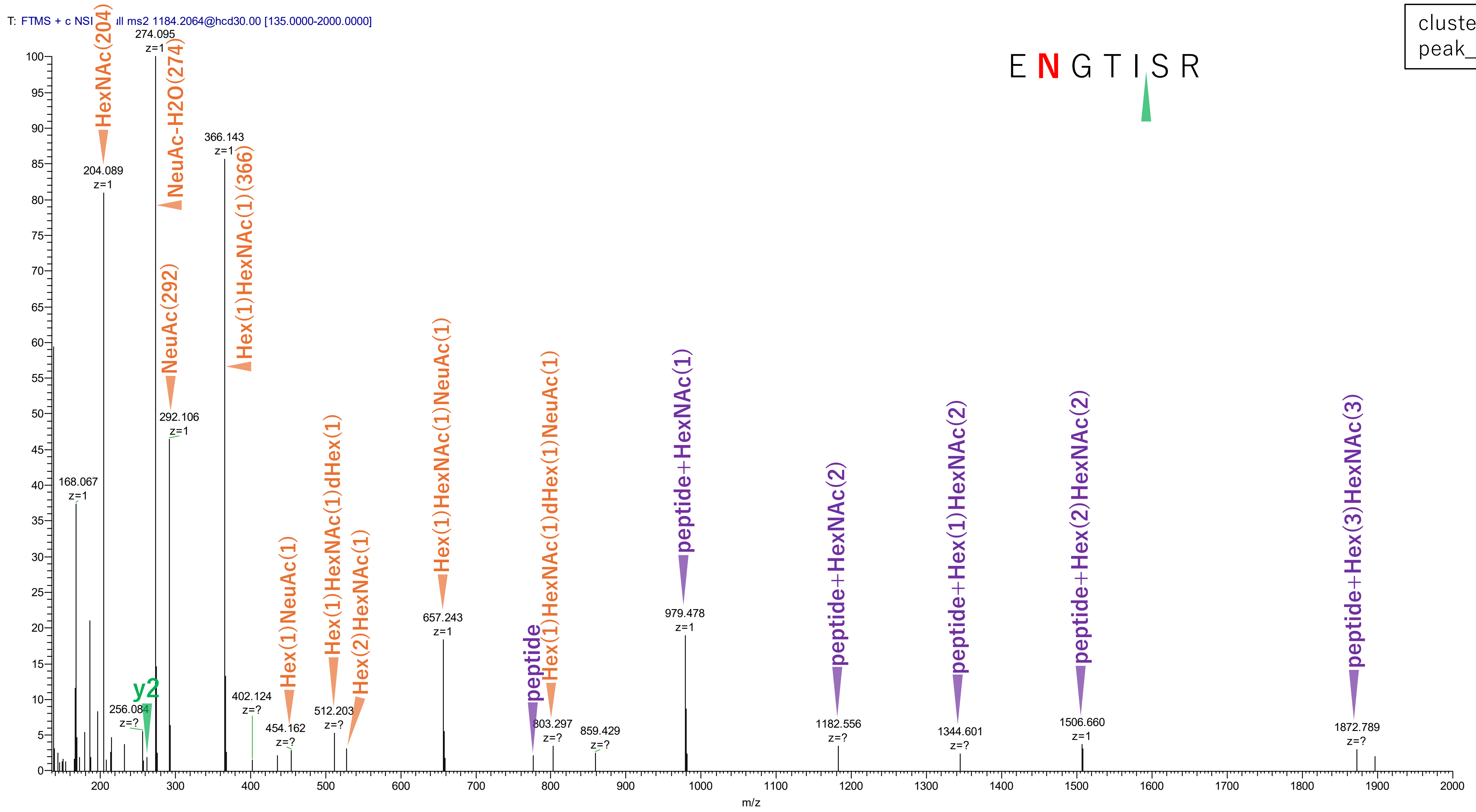


Figure S2-46. MS2 spectra of glycopeptides assigned for hAGP.

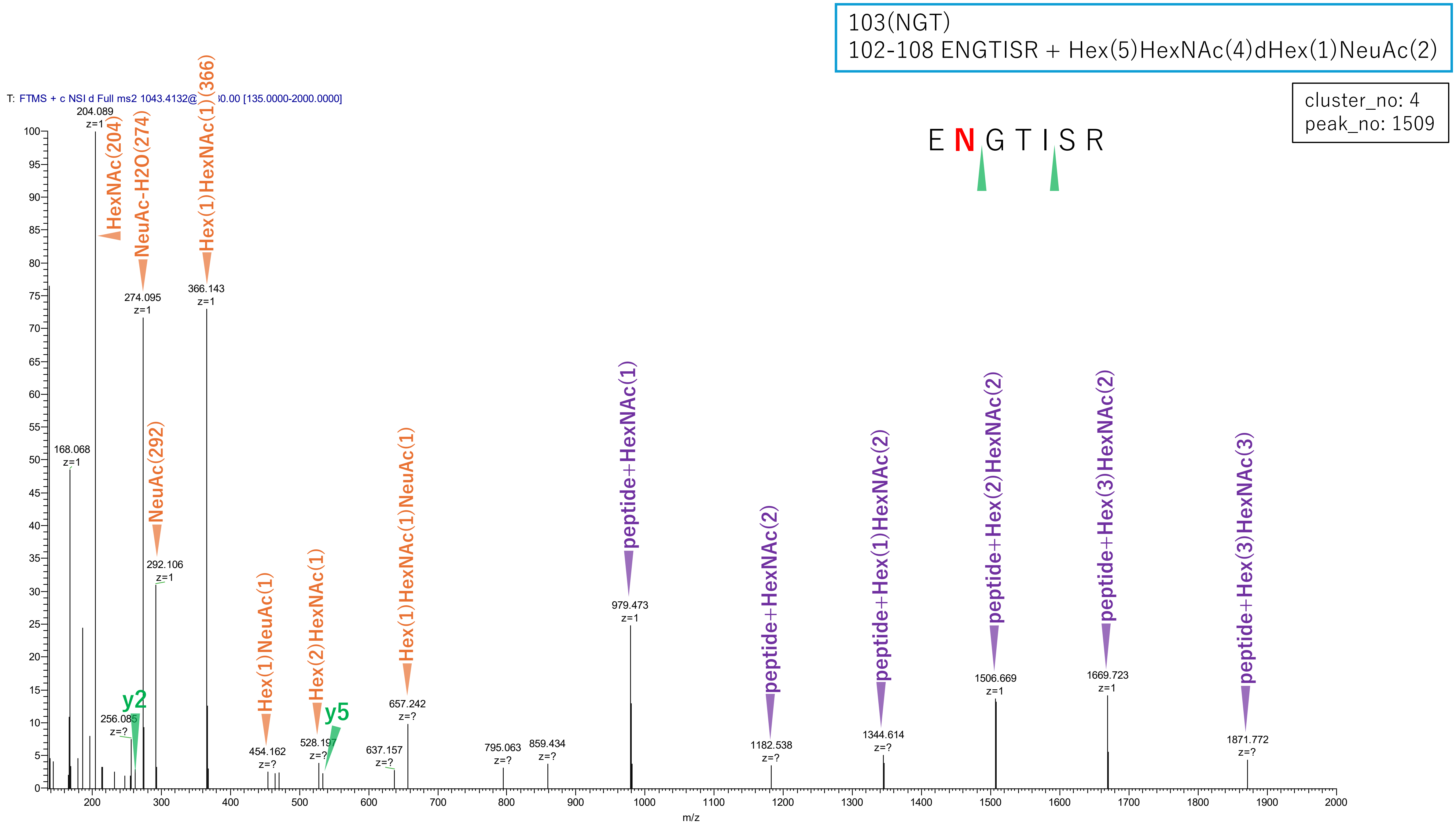


**Figure S2-47. MS2 spectra of glycopeptides assigned for hAGP.**

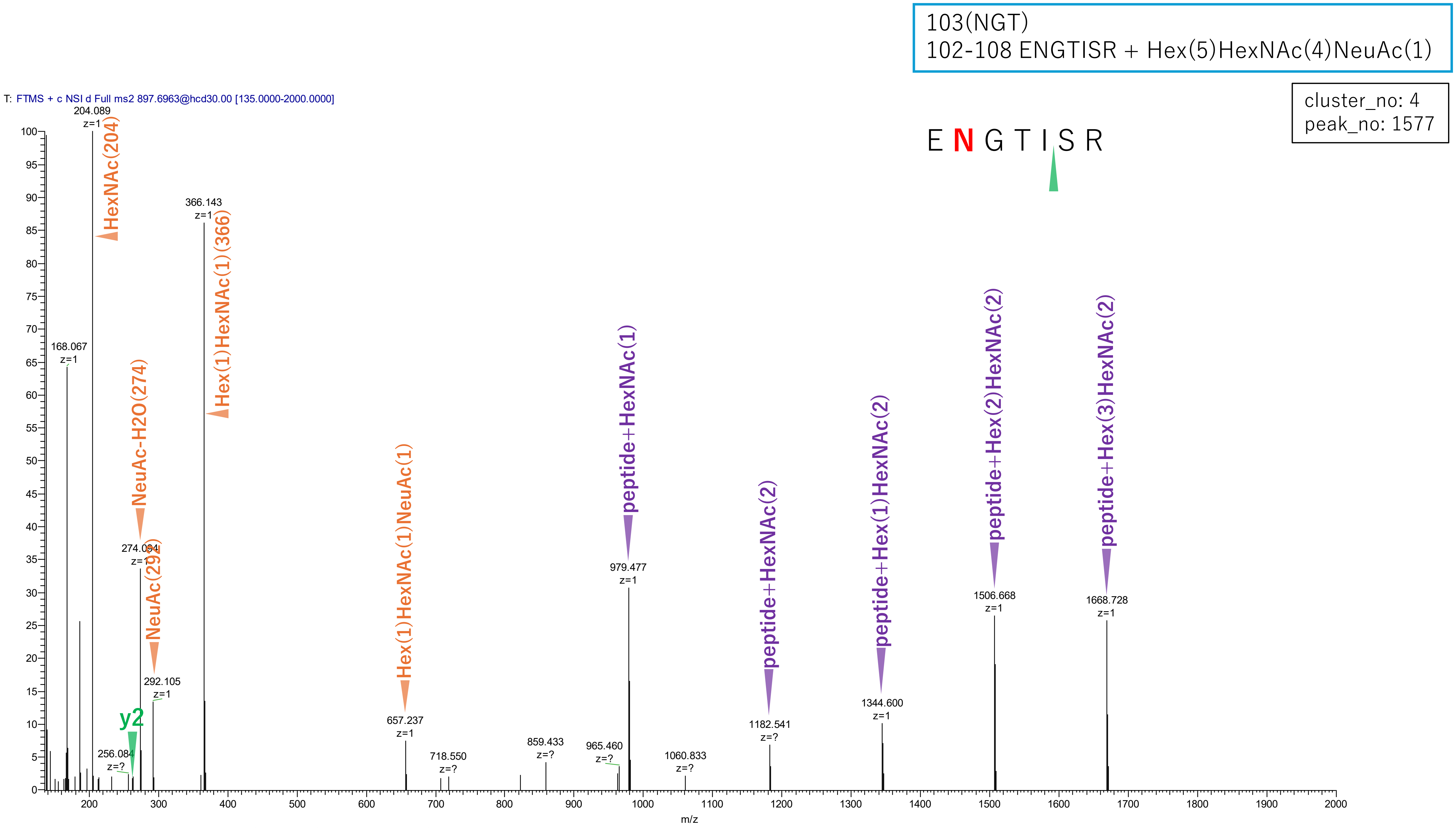


**Figure S2-48. MS2 spectra of glycopeptides assigned for hAGP.**

103(NGT)
102-108 ENGTISR + Hex(6)HexNAc(5)NeuAc(1)

cluster_no: 4
peak_no: 1582

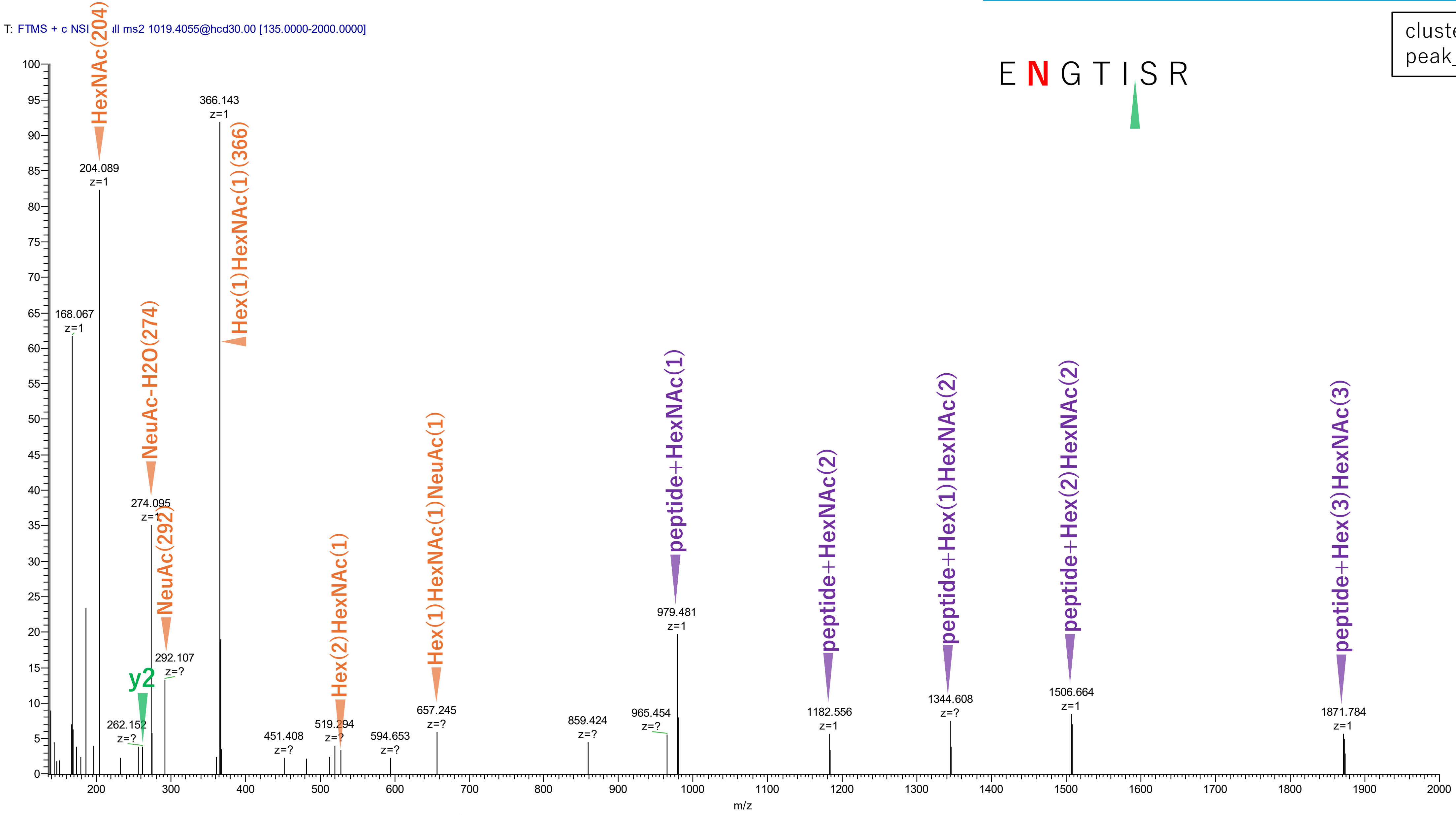


Figure S2-49. MS2 spectra of glycopeptides assigned for hAGP.

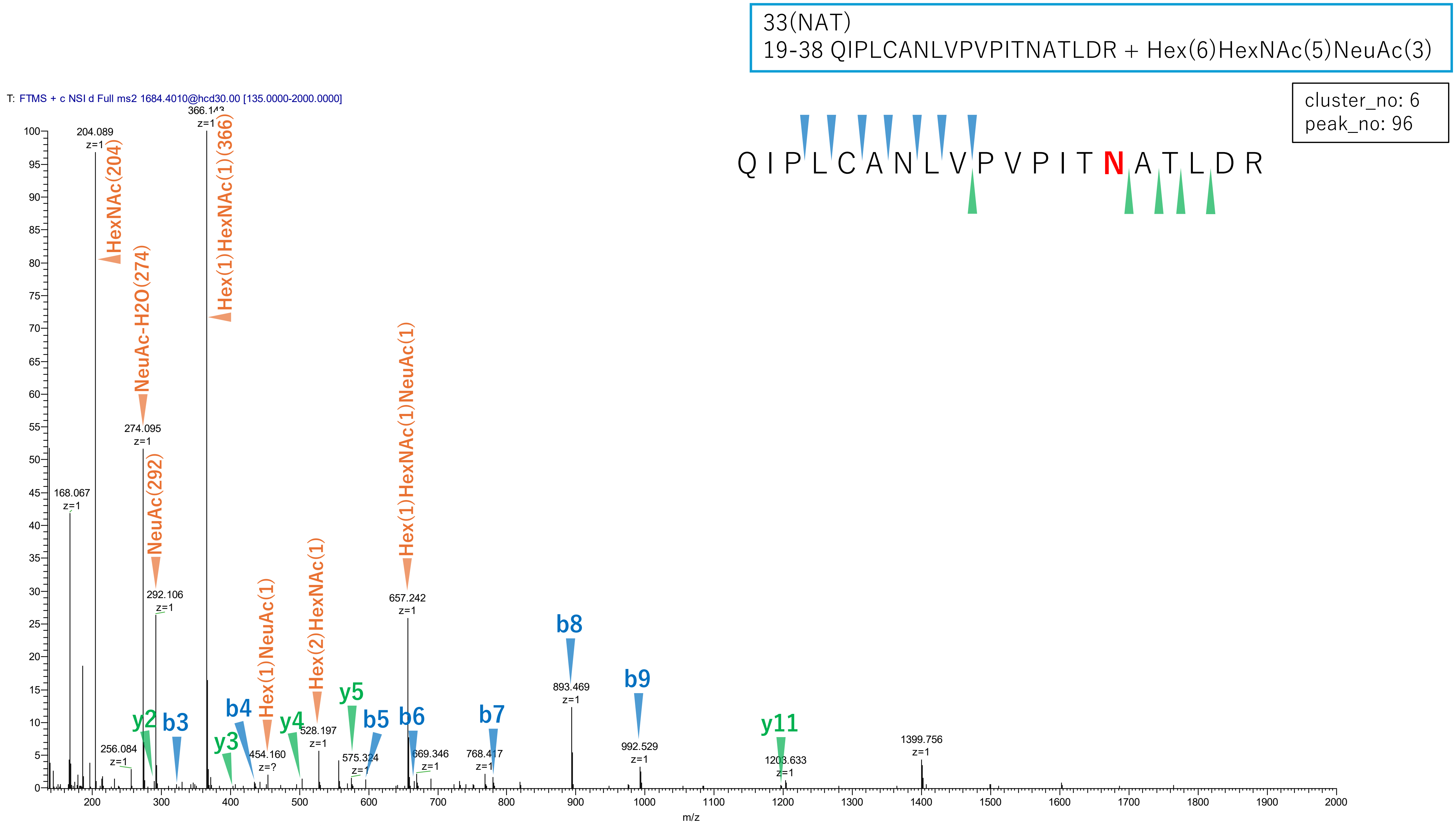


**Figure S2-50. MS2 spectra of glycopeptides assigned for hAGP.**

33(NAT)
19-38 QIPLCANLVPVPITNATLDR + Hex(6)HexNAc(5)NeuAc(2)

cluster_no: 6
peak_no: 107

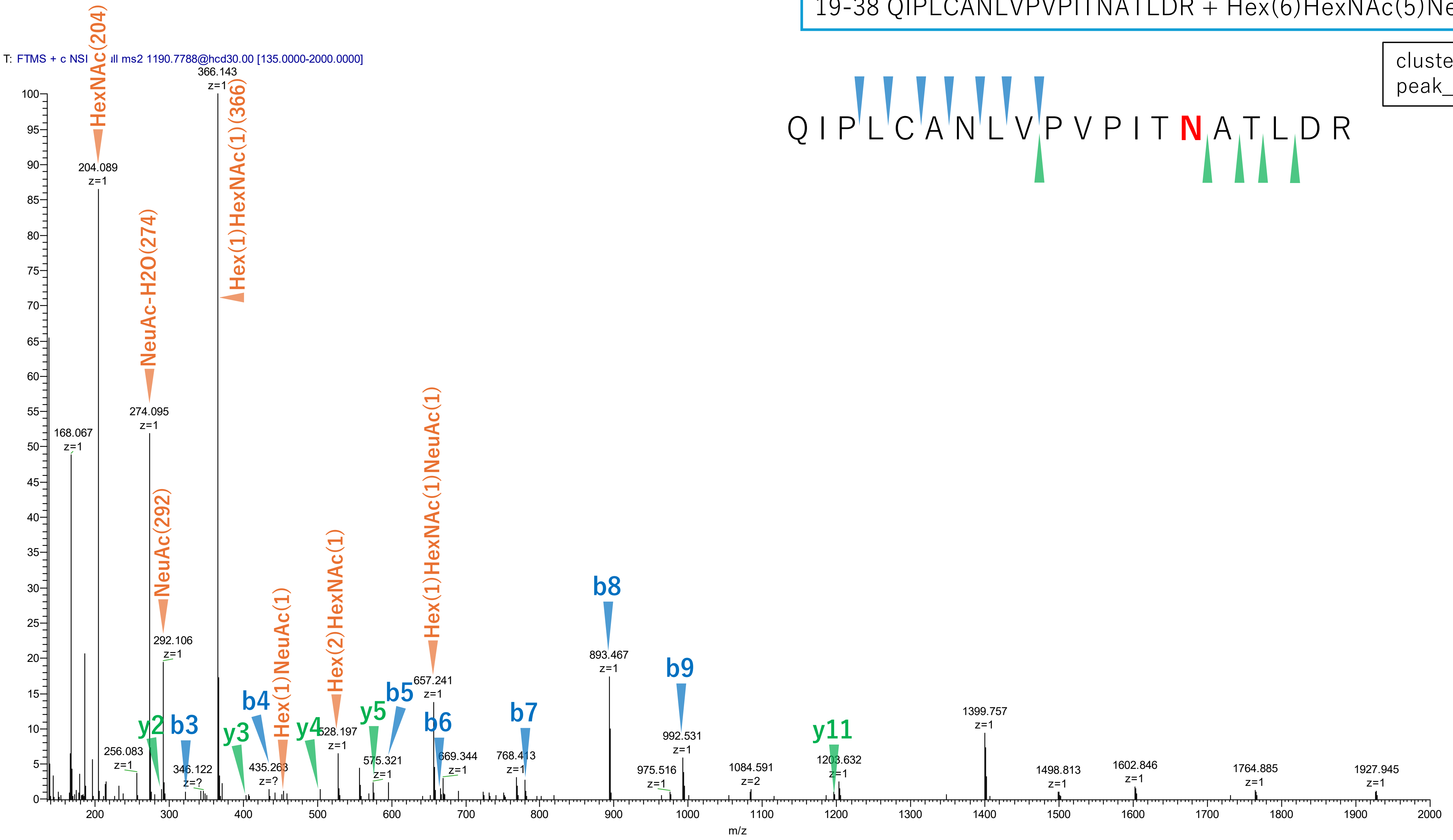


**Figure S2-51. MS2 spectra of glycopeptides assigned for hAGP.**

33(NAT)
19-38 QIPLCANLVPVPITNATLDR + Hex(5)HexNAc(4)NeuAc(2)

cluster_no: 6
peak_no: 155

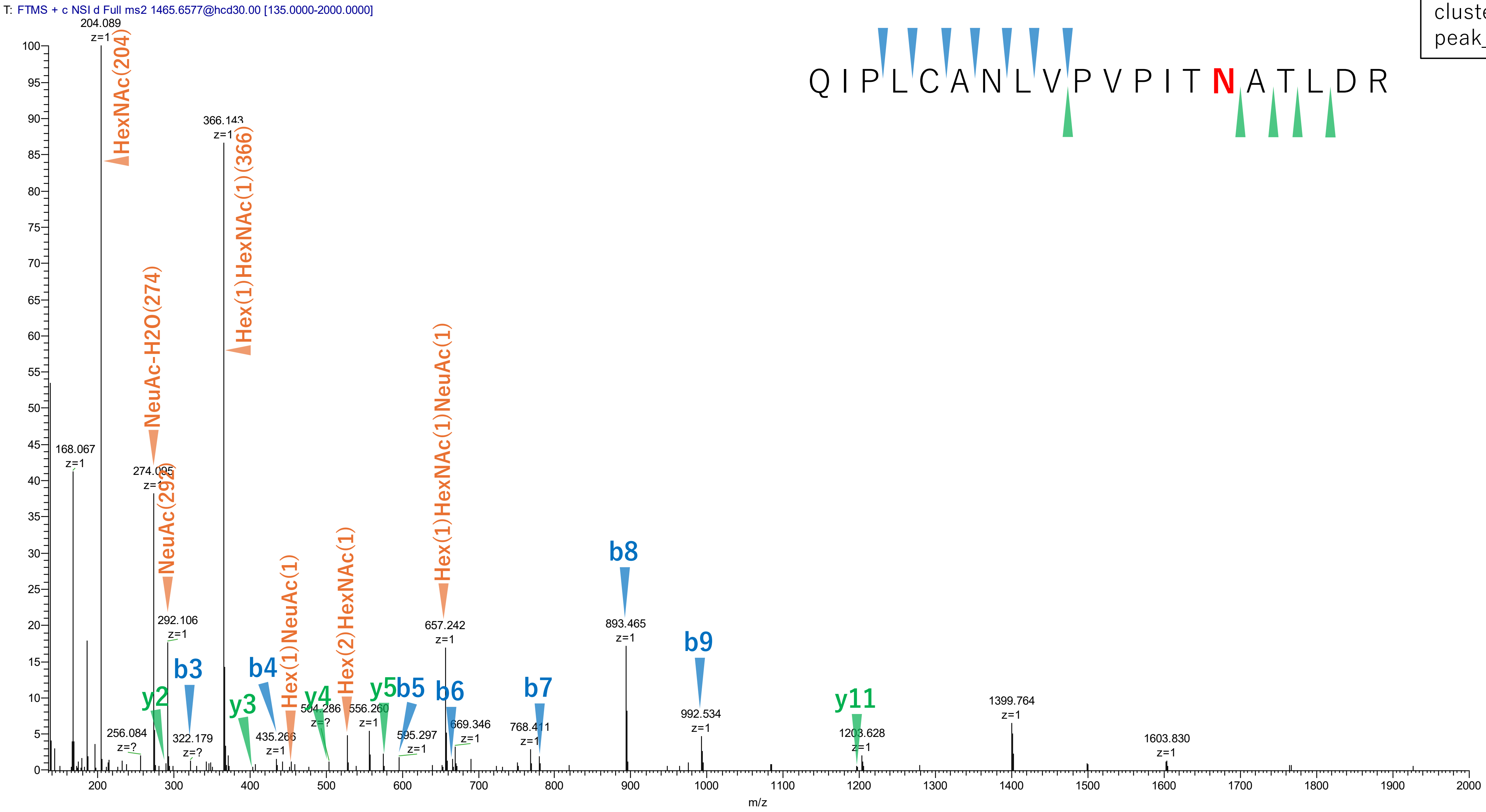


Figure S2-52. MS2 spectra of glycopeptides assigned for hAGP.

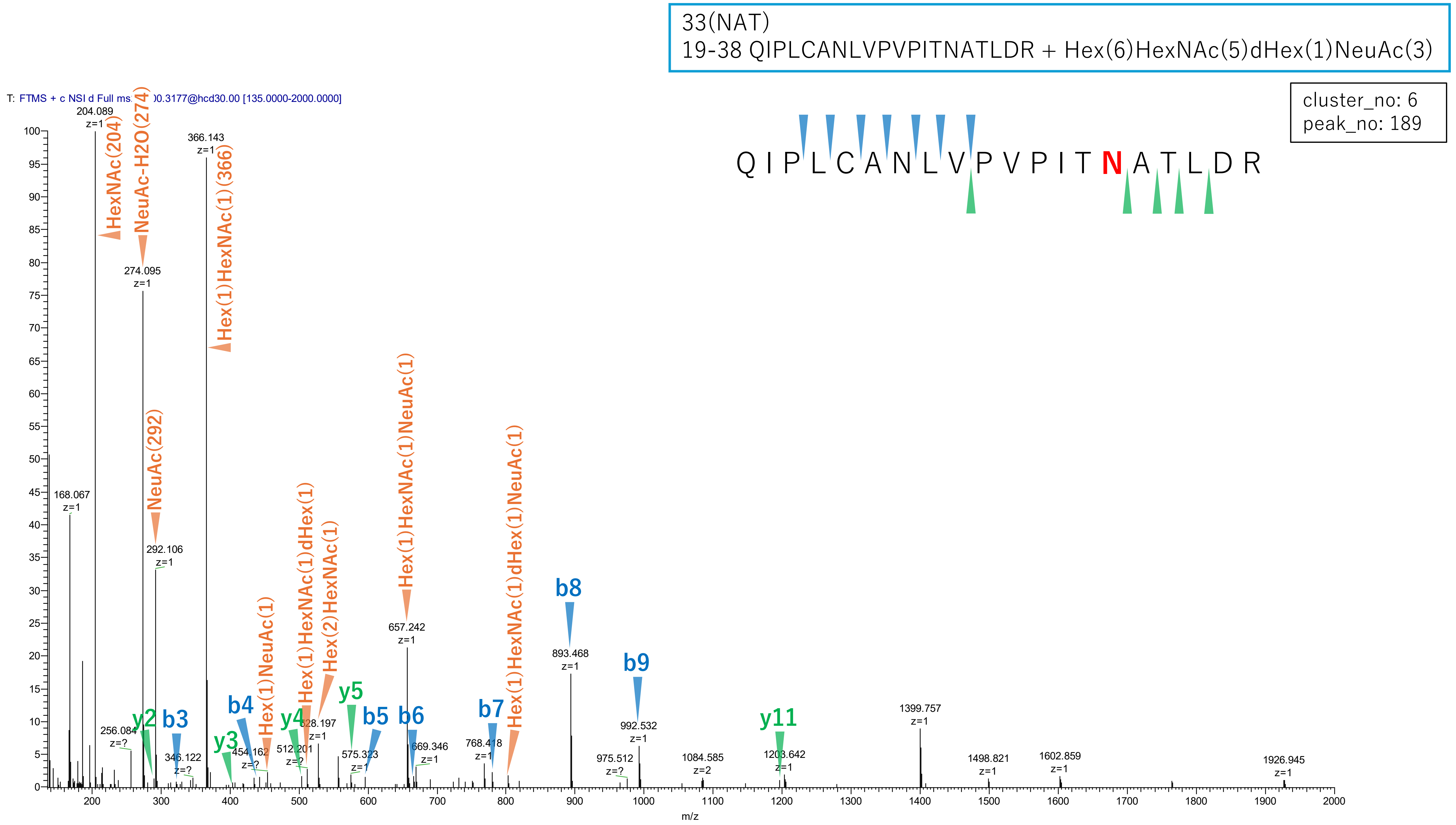


Figure S2-53. MS2 spectra of glycopeptides assigned for hAGP.

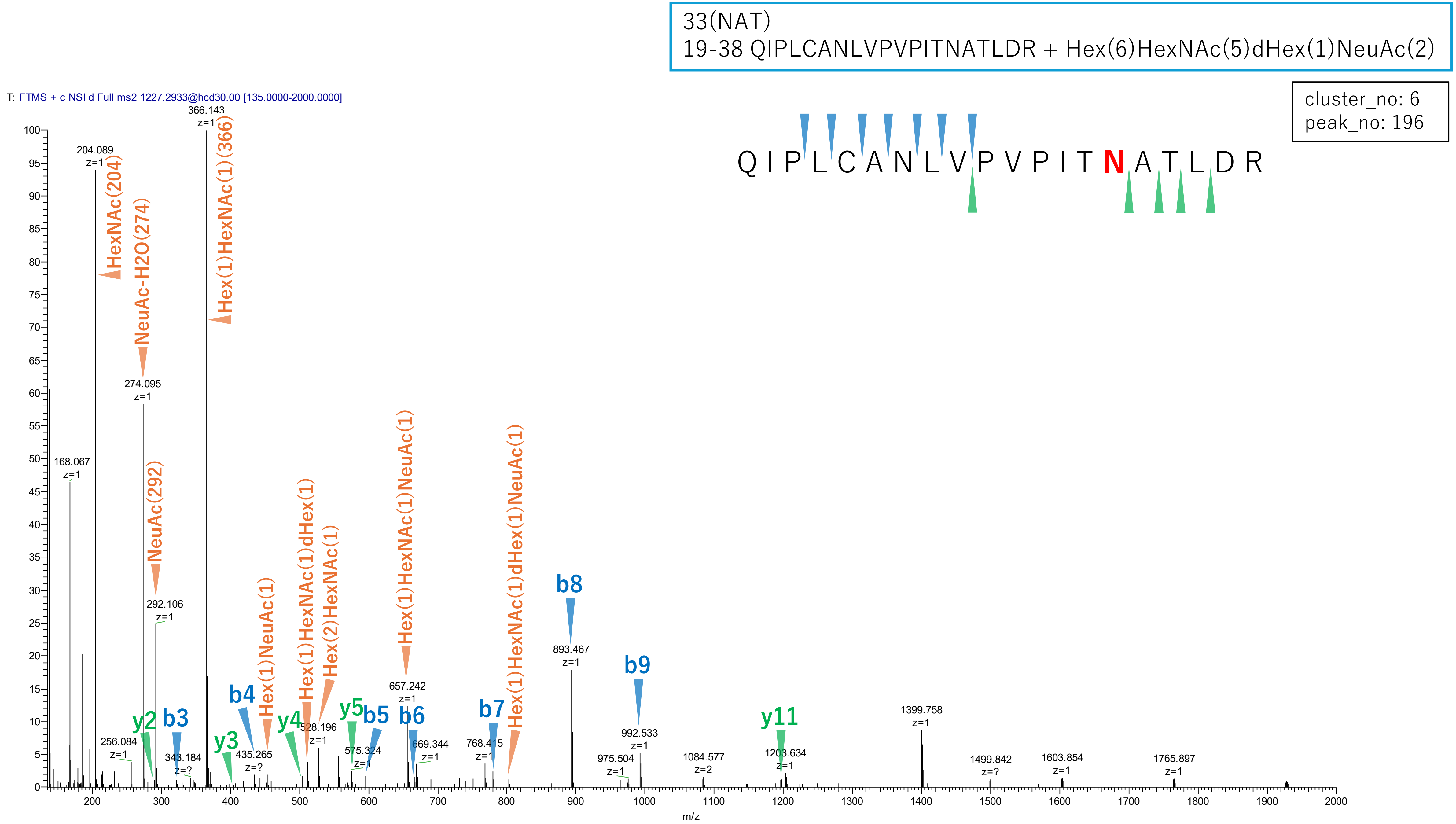


**Figure S2-54. MS2 spectra of glycopeptides assigned for hAGP.**

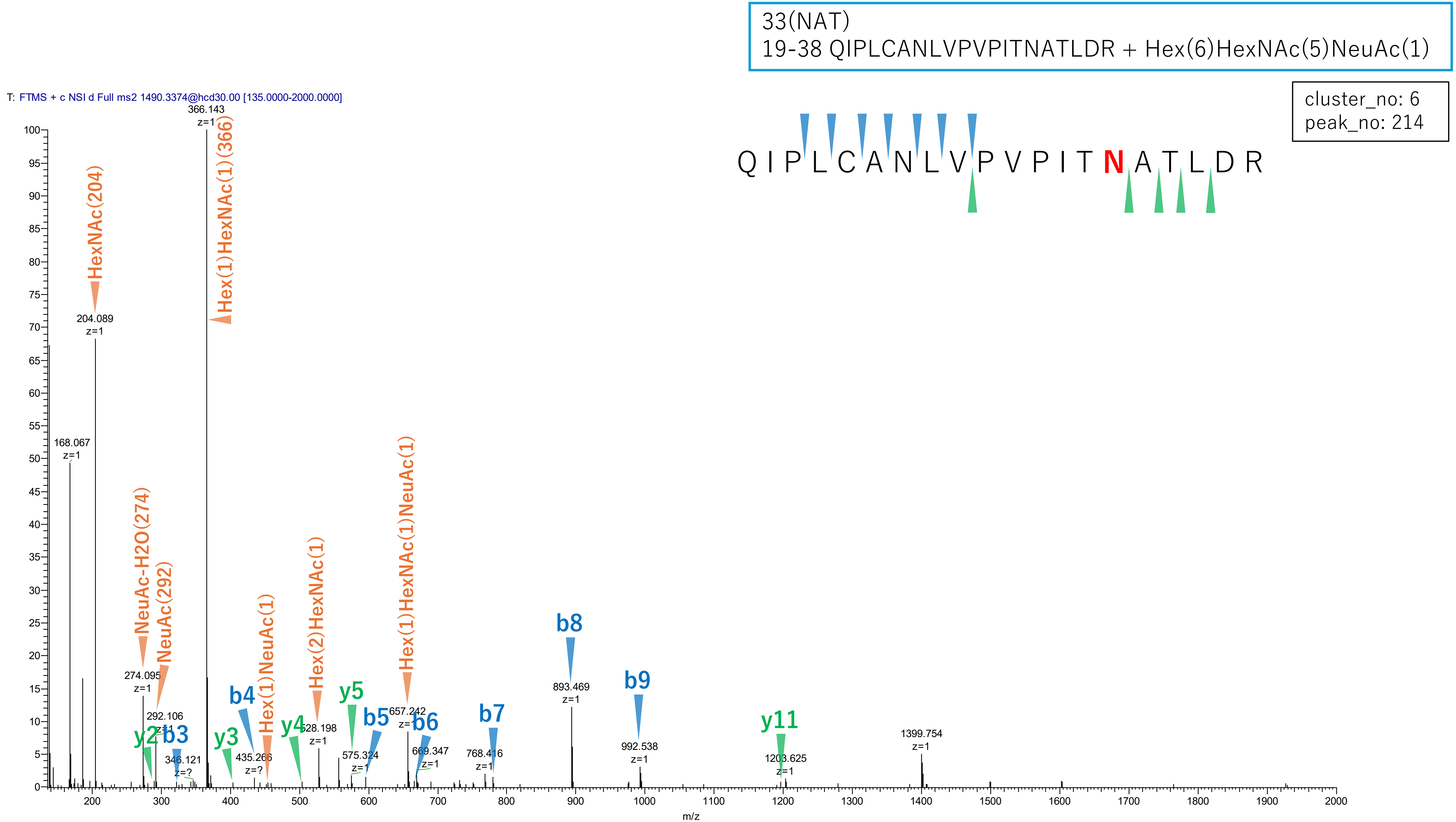


**Figure S2-55. MS2 spectra of glycopeptides assigned for hAGP.**

33(NAT)
19-38 QIPLCANLVPVPITNATLDR + Hex(5)HexNAc(4)NeuAc(1)

cluster_no: 6
peak_no: 423

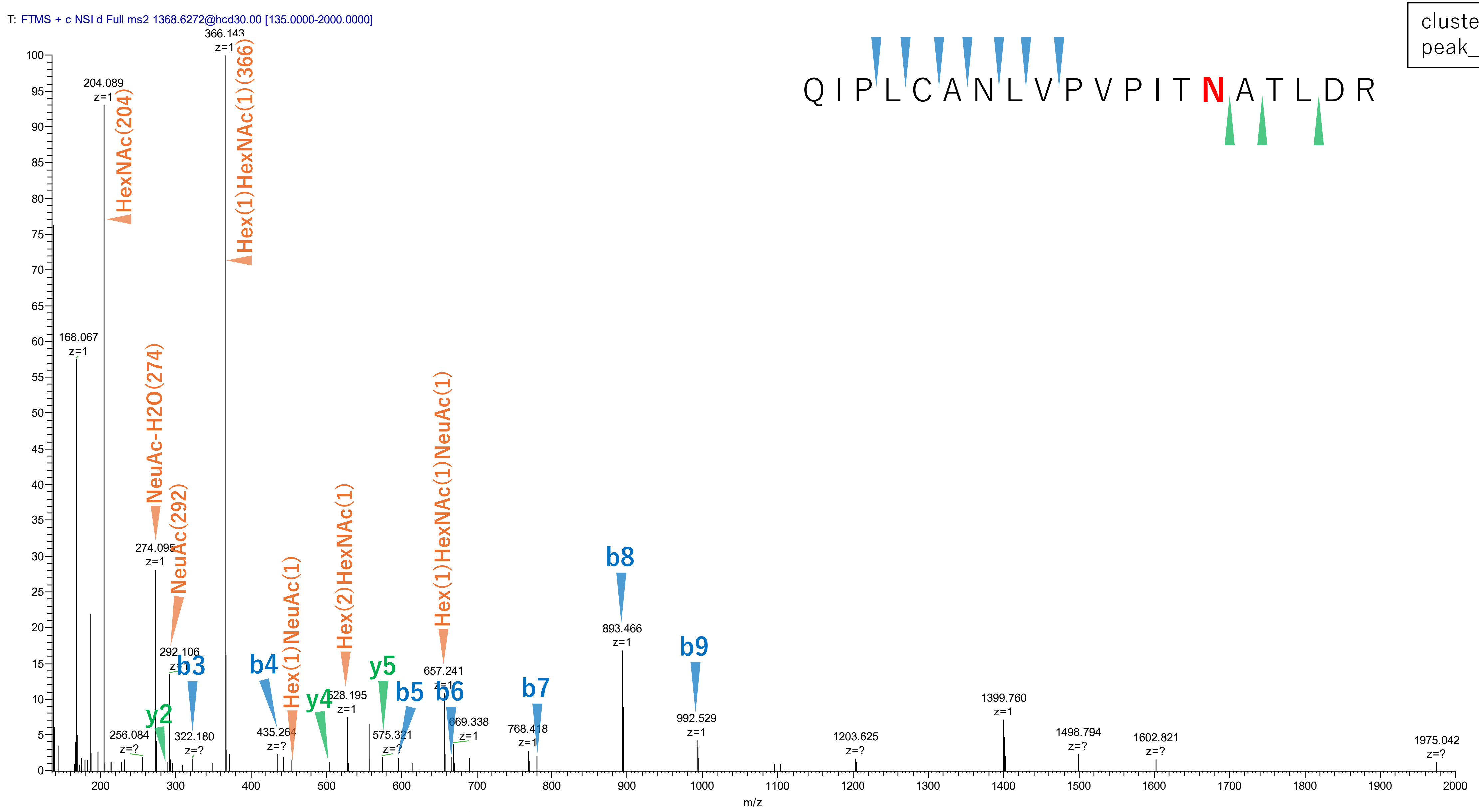


**Figure S2-56. MS2 spectra of glycopeptides assigned for hAGP.**

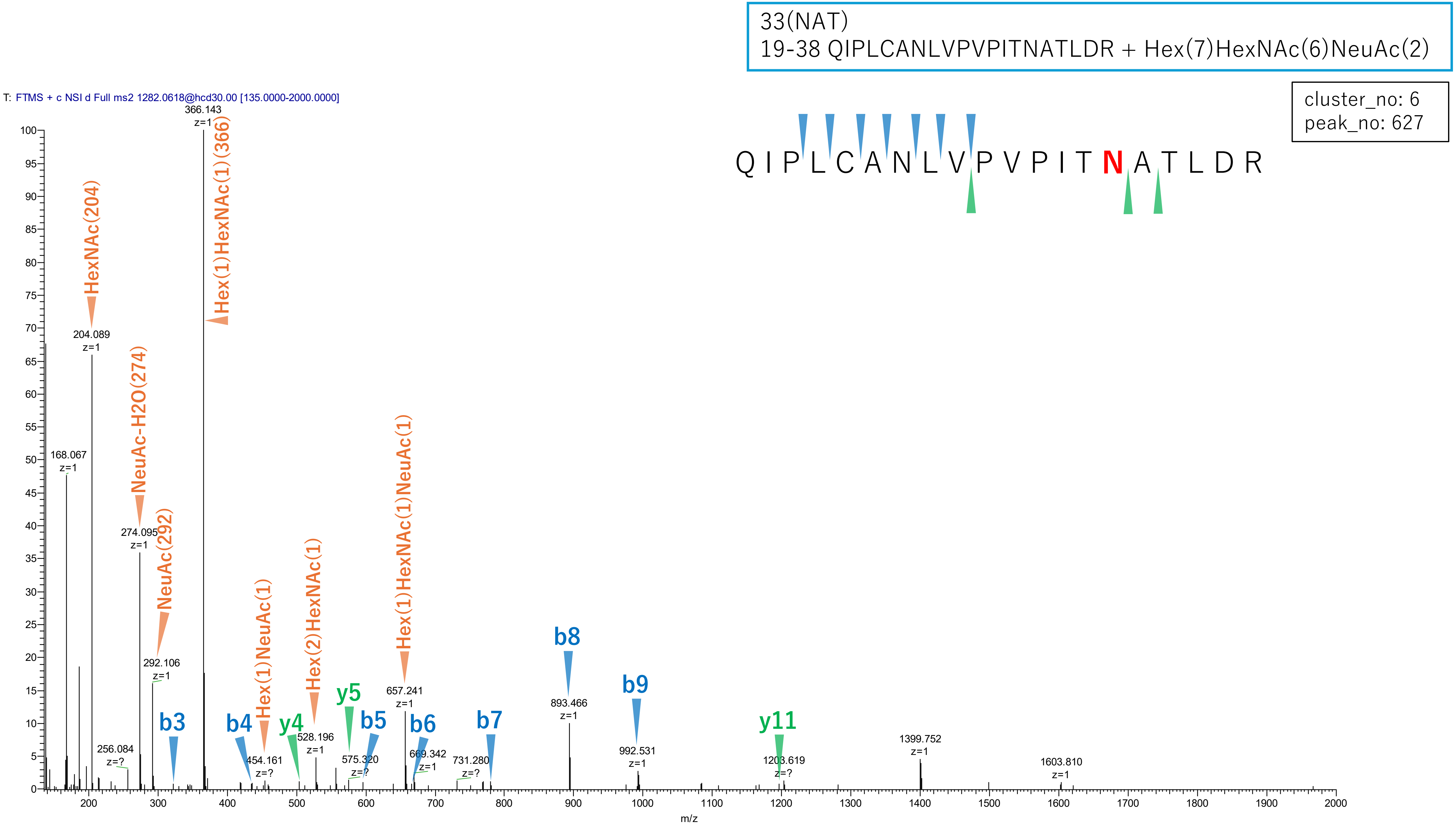


**Figure S2-57. MS2 spectra of glycopeptides assigned for hAGP.**

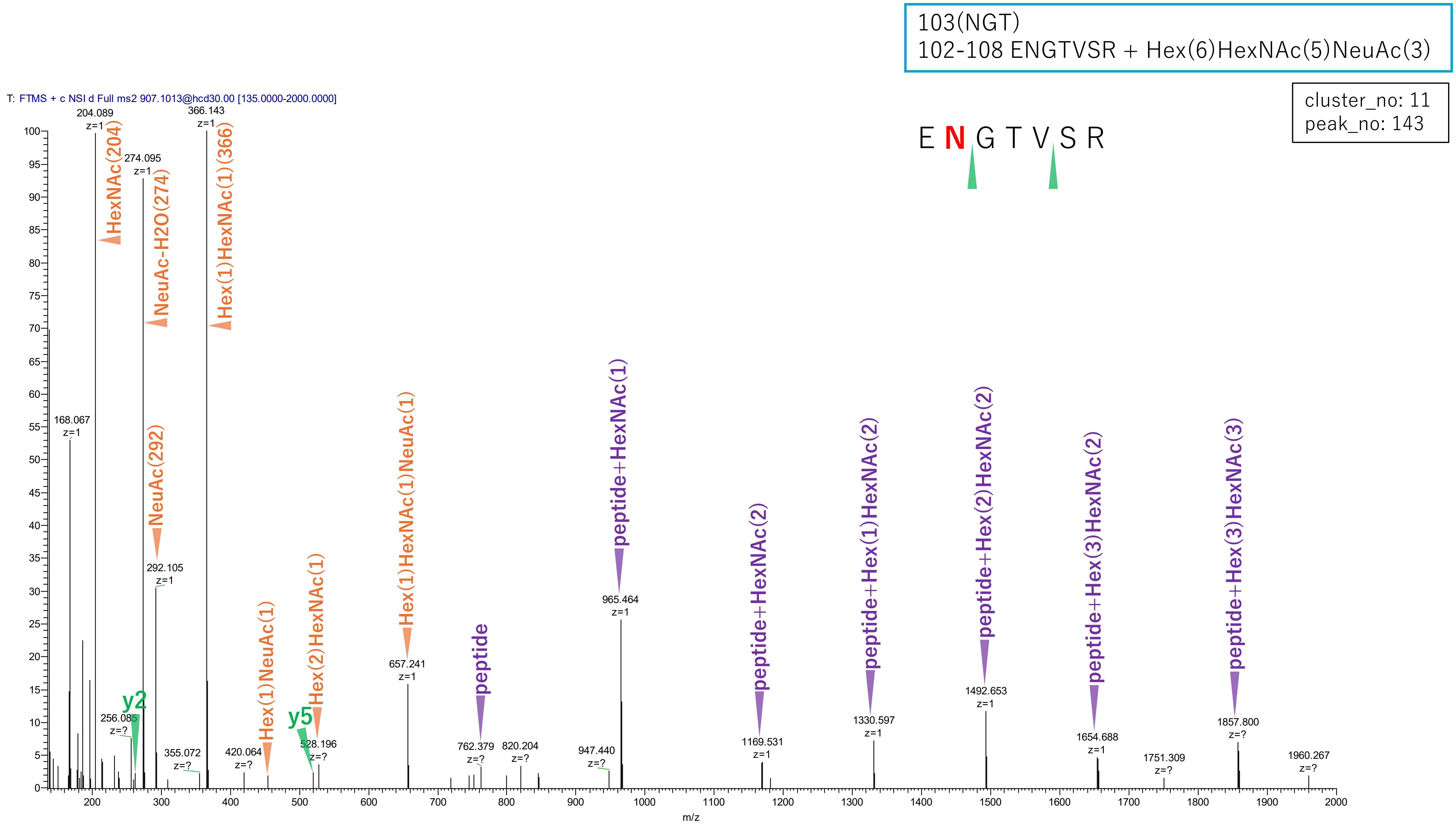


**Figure S2-58. MS2 spectra of glycopeptides assigned for hAGP.**

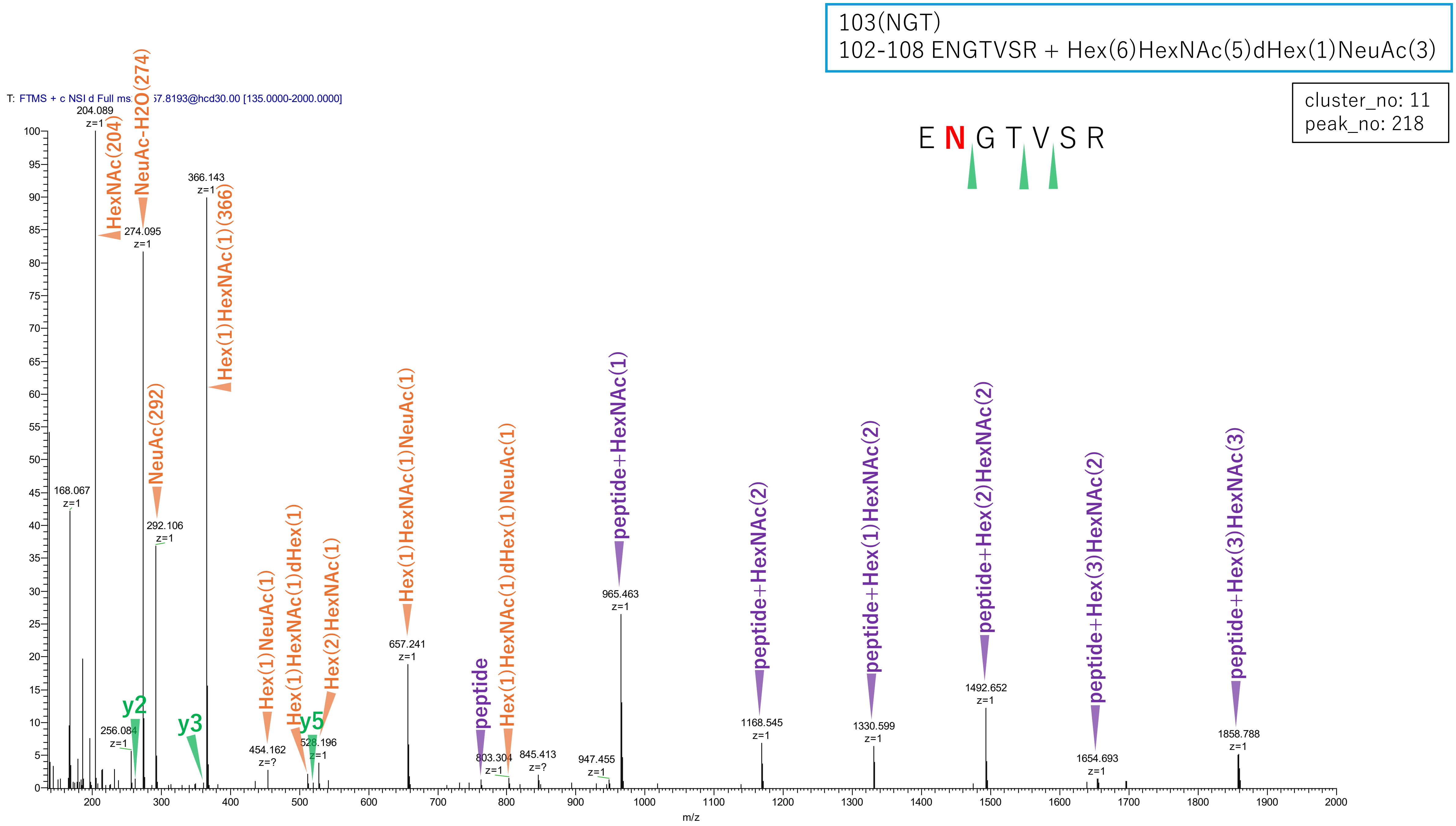


Figure S2-59. MS2 spectra of glycopeptides assigned for hAGP.

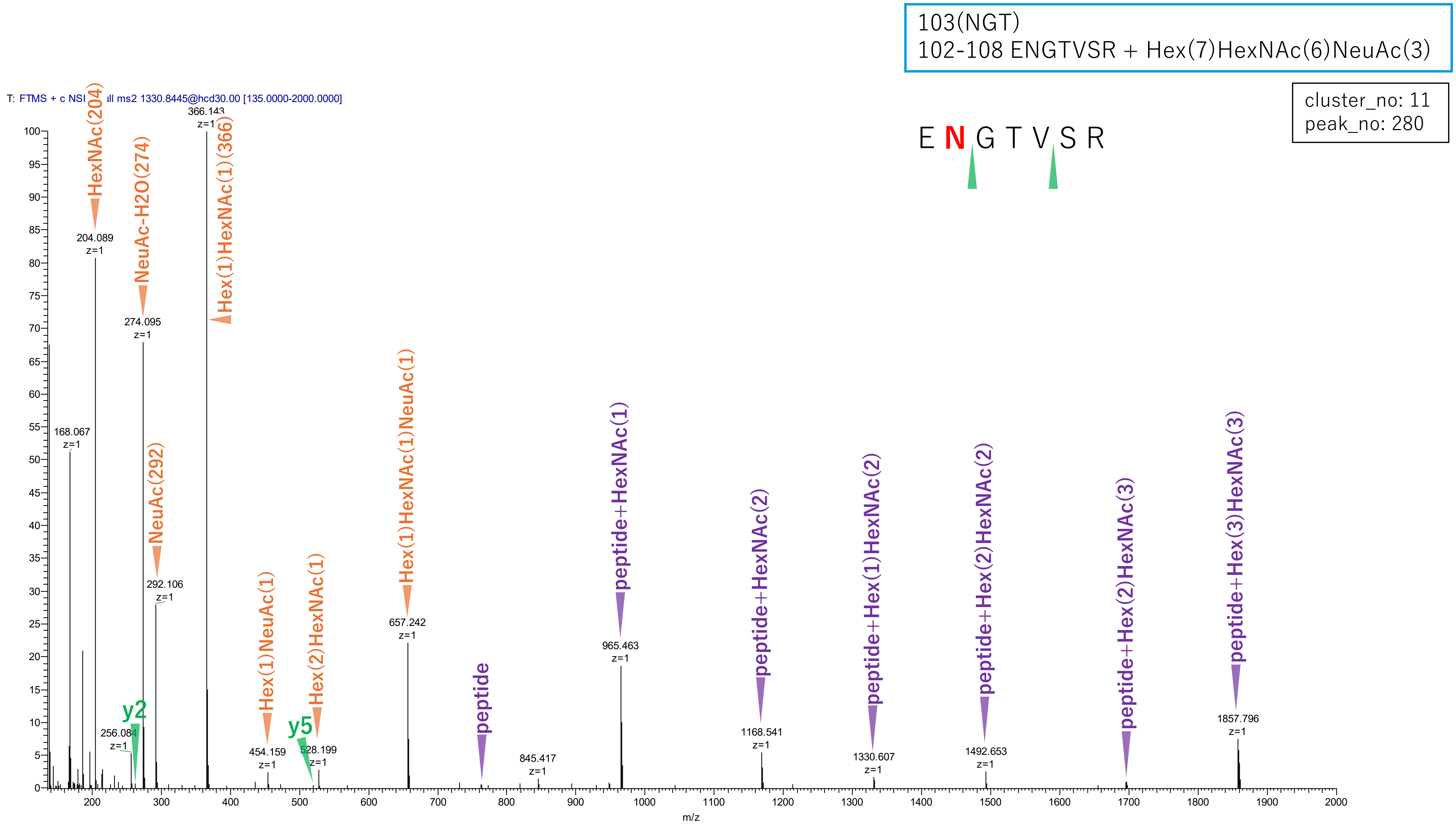


**Figure S2-60. MS2 spectra of glycopeptides assigned for hAGP.**

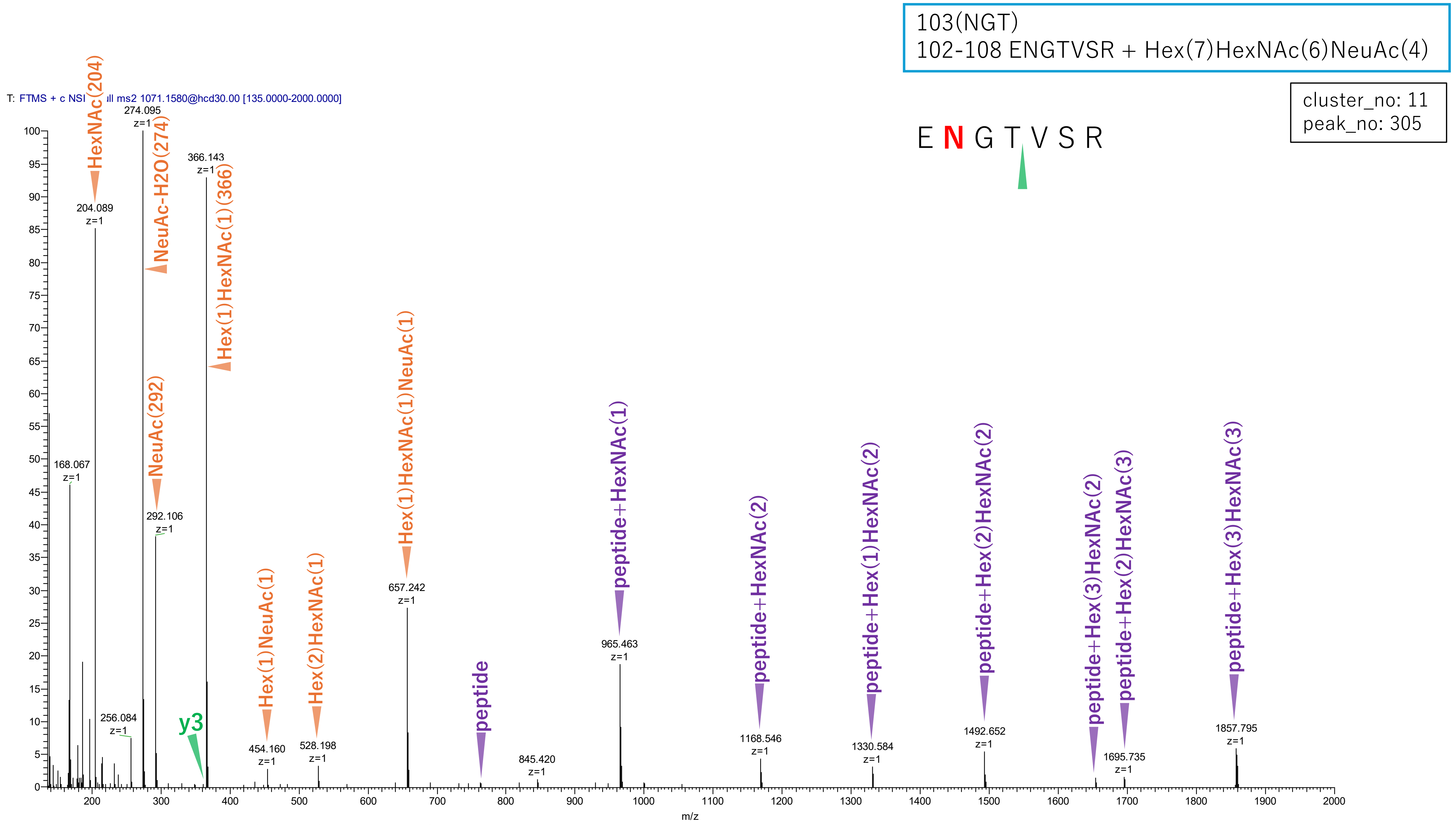


**Figure S2-61. MS2 spectra of glycopeptides assigned for hAGP.**

103(NGT)
102-108 ENGTVSR + Hex(6)HexNAc(5)NeuAc(2)

cluster_no: 11
peak_no: 370

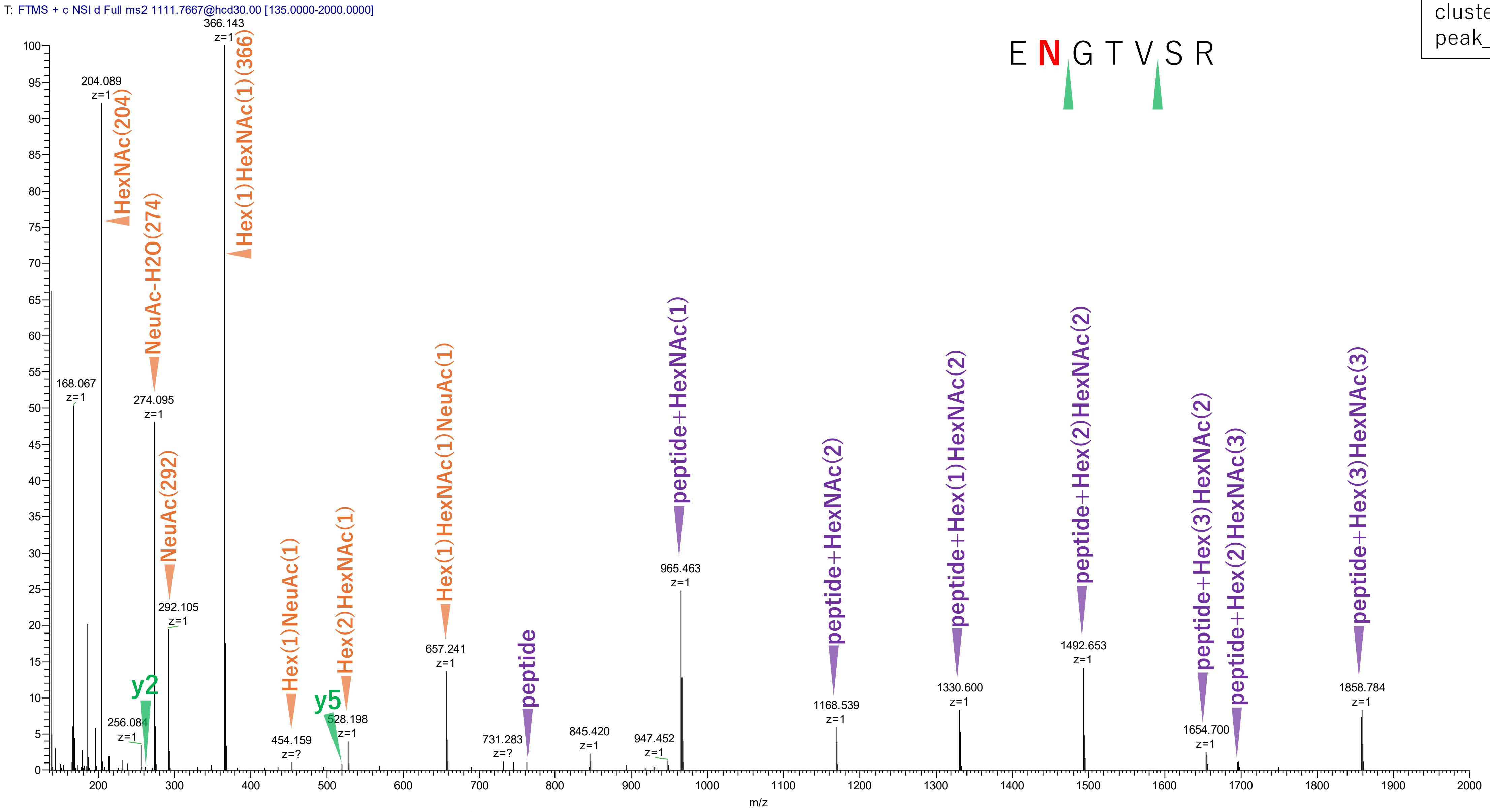


**Figure S2-62. MS2 spectra of glycopeptides assigned for hAGP.**

103(NGT)
102-108 ENGTVSR + Hex(7)HexNAc(6)dHex(1)NeuAc(3)

cluster_no: 11
peak_no: 480

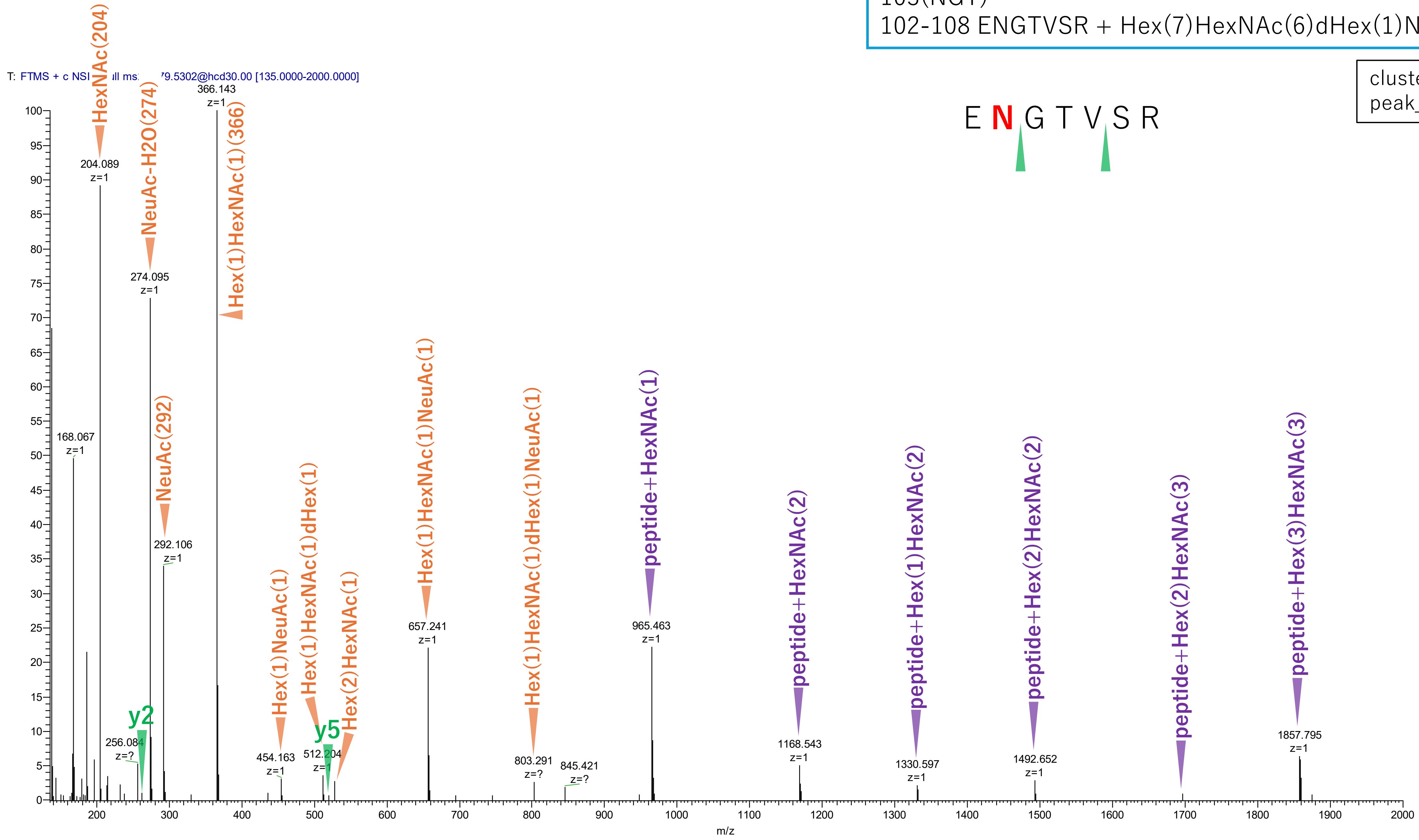


**Figure S2-63. MS2 spectra of glycopeptides assigned for hAGP.**

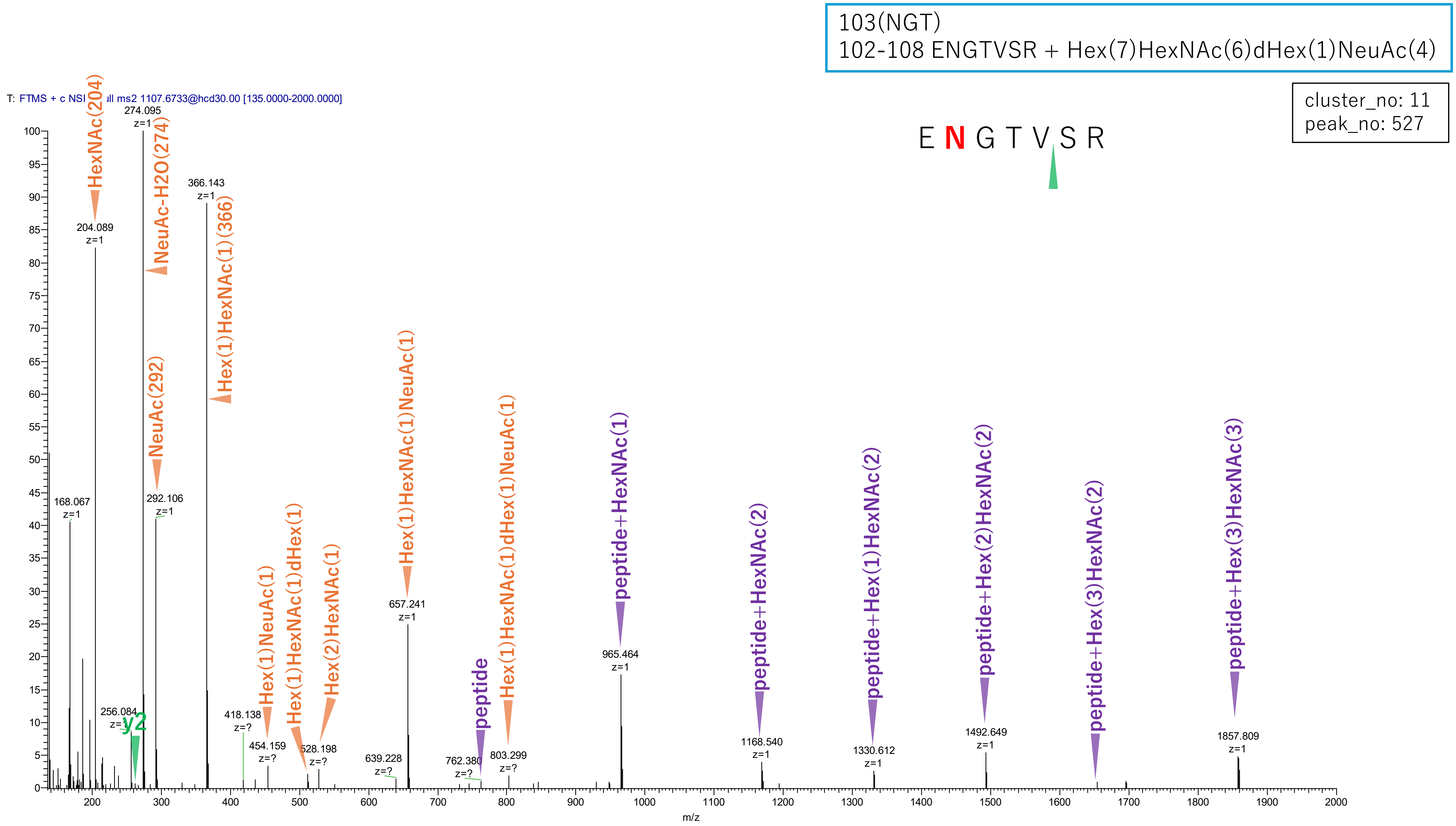


**Figure S2-64. MS2 spectra of glycopeptides assigned for hAGP.**

103(NGT)
102-108 ENGTVSR + Hex(6)HexNAc(5)dHex(1)NeuAc(2)

cluster_no: 11
peak_no: 685

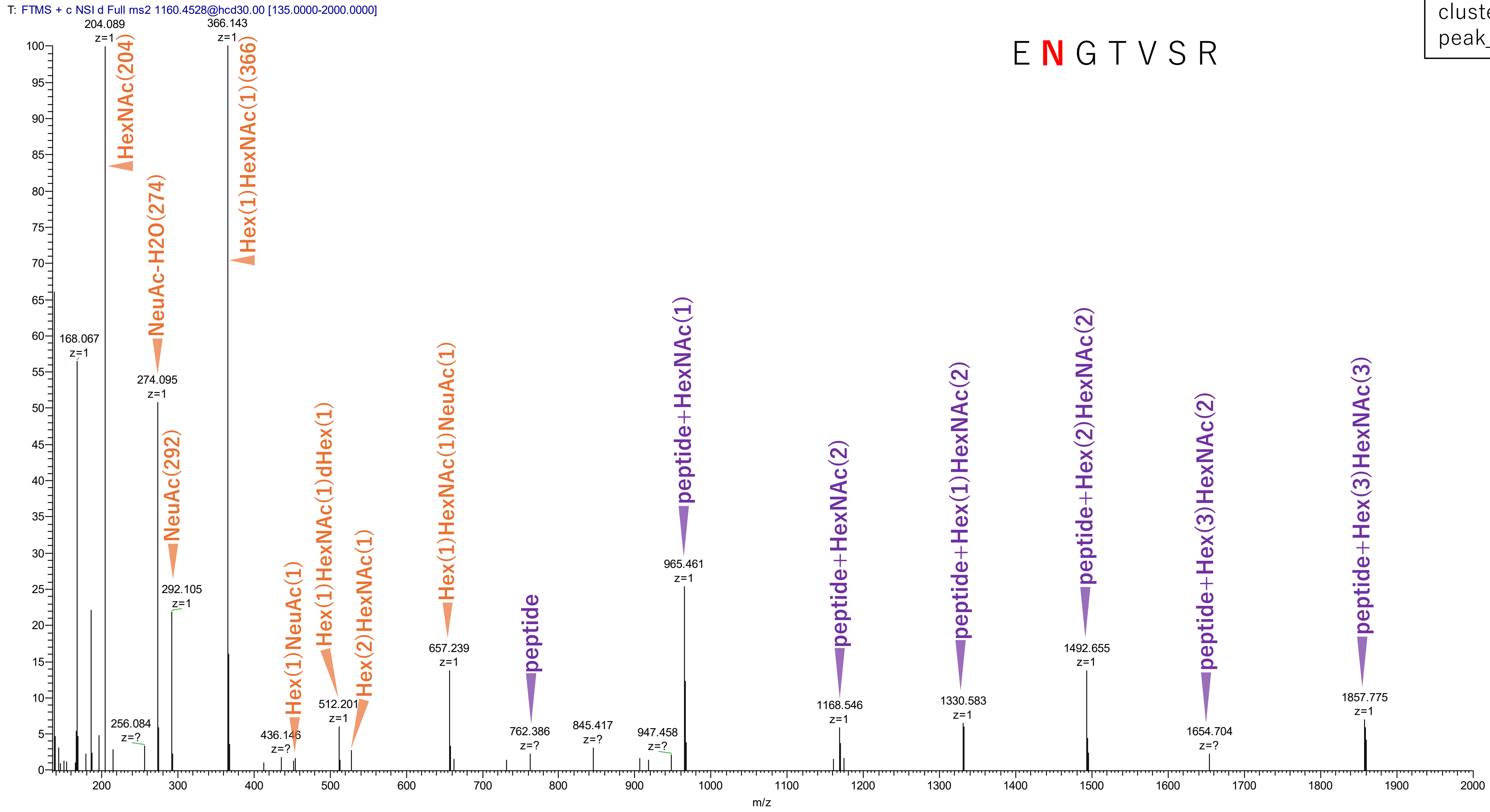


**Figure S2-65. MS2 spectra of glycopeptides assigned for hAGP.**

103(NGT)
102-108 ENGTVSR + Hex(7)HexNAc(6)NeuAc(2)

cluster_no: 11
peak_no: 762

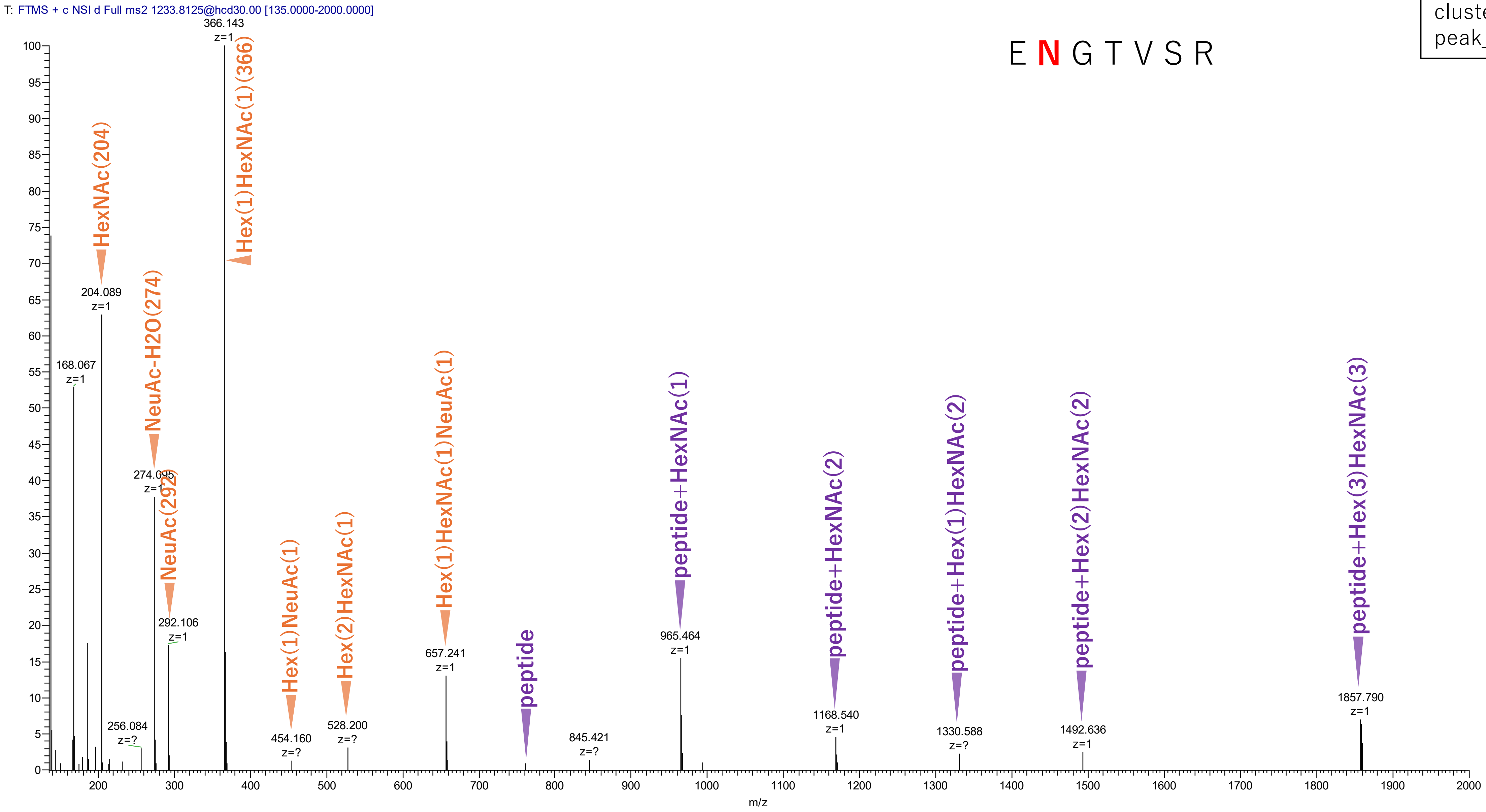


Figure S2-66. MS2 spectra of glycopeptides assigned for hAGP.

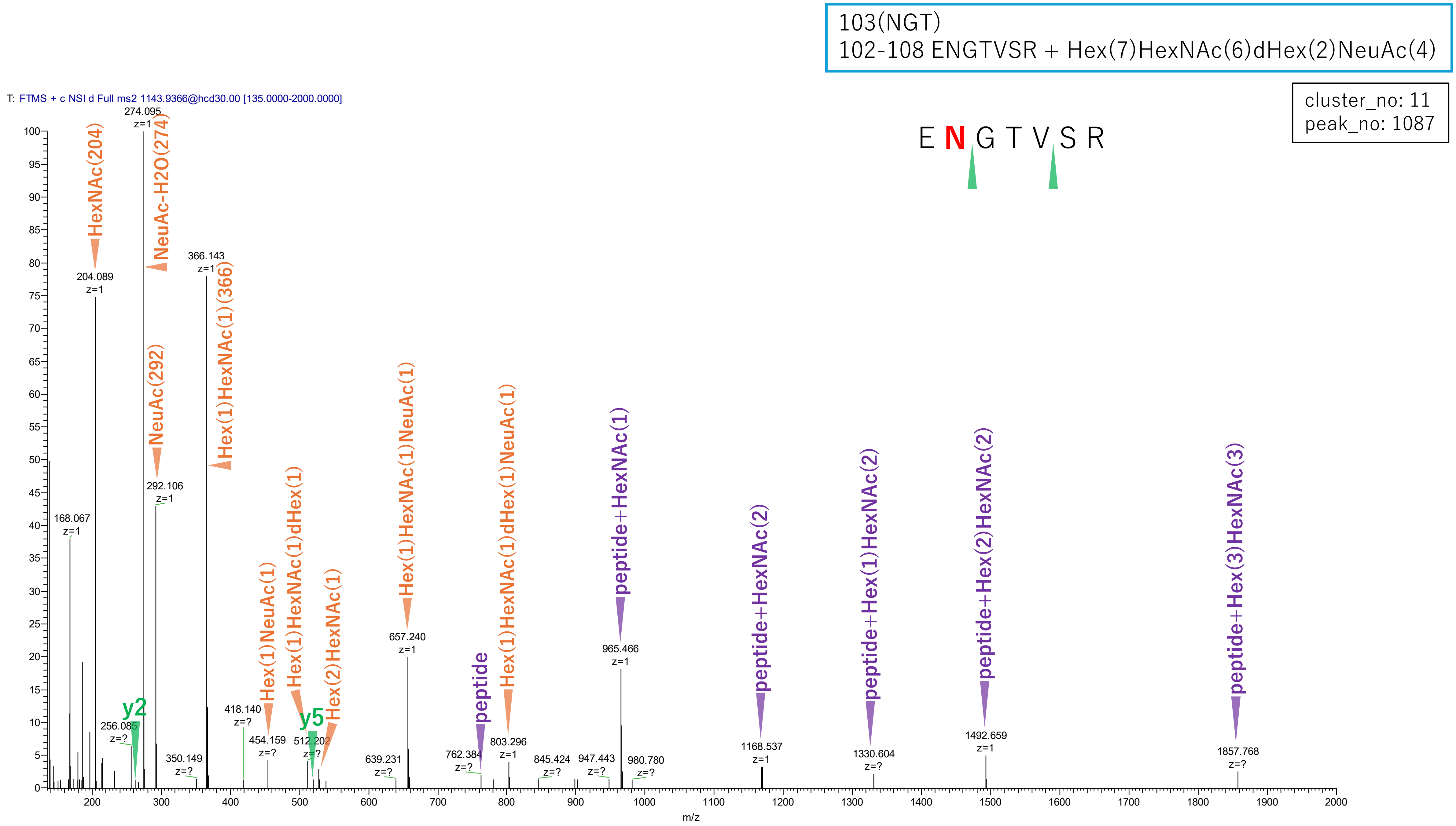


**Figure S2-67. MS2 spectra of glycopeptides assigned for hAGP.**

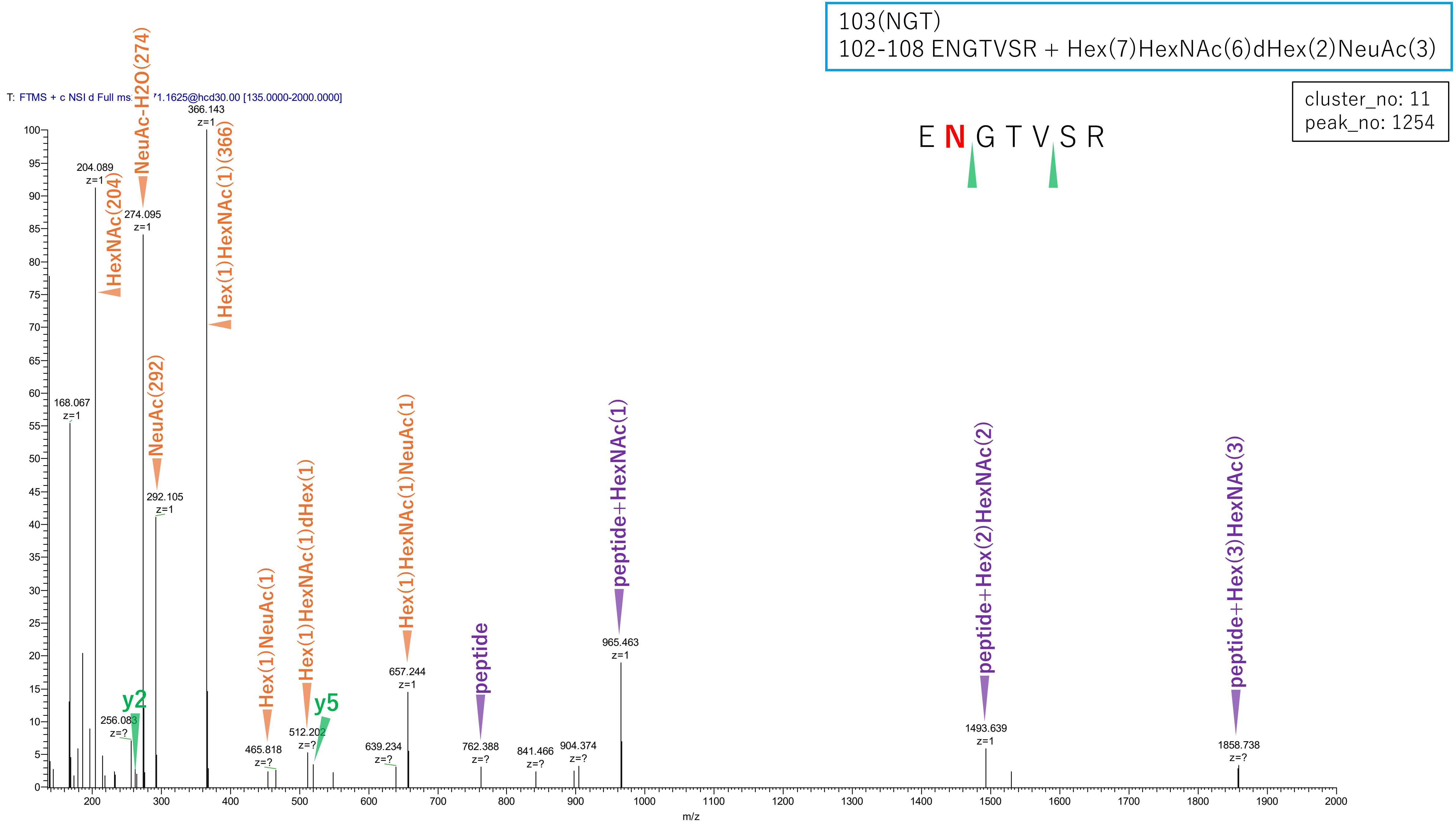


**Figure S2-68. MS2 spectra of glycopeptides assigned for hAGP.**

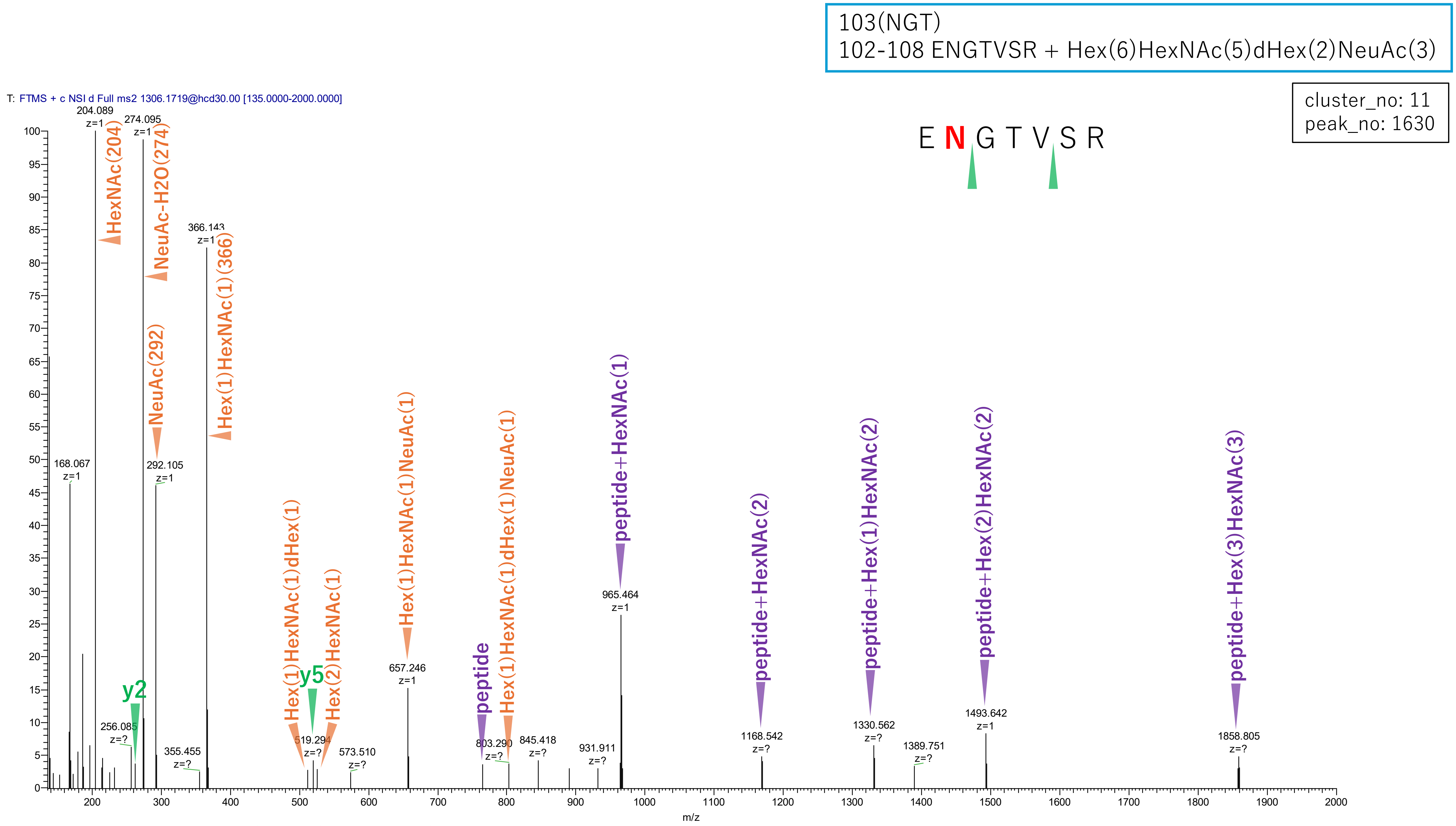


**Figure S2-69. MS2 spectra of glycopeptides assigned for hAGP.**

103(NGT)
102-108 ENGTVSR + Hex(8)HexNAc(7)NeuAc(4)

cluster_no: 11
peak_no: 1632

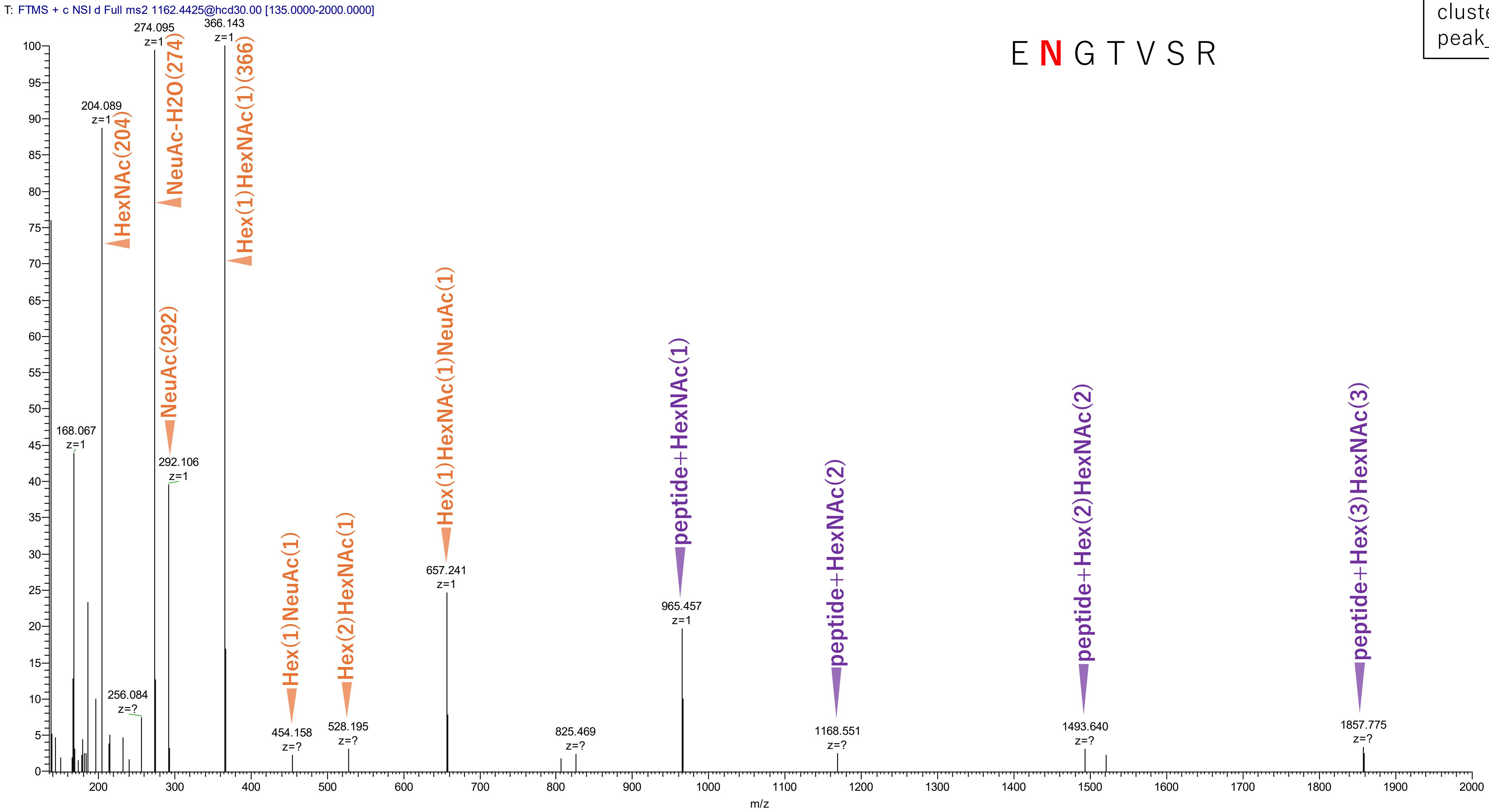


**Figure S2-70. MS2 spectra of glycopeptides assigned for hAGP.**

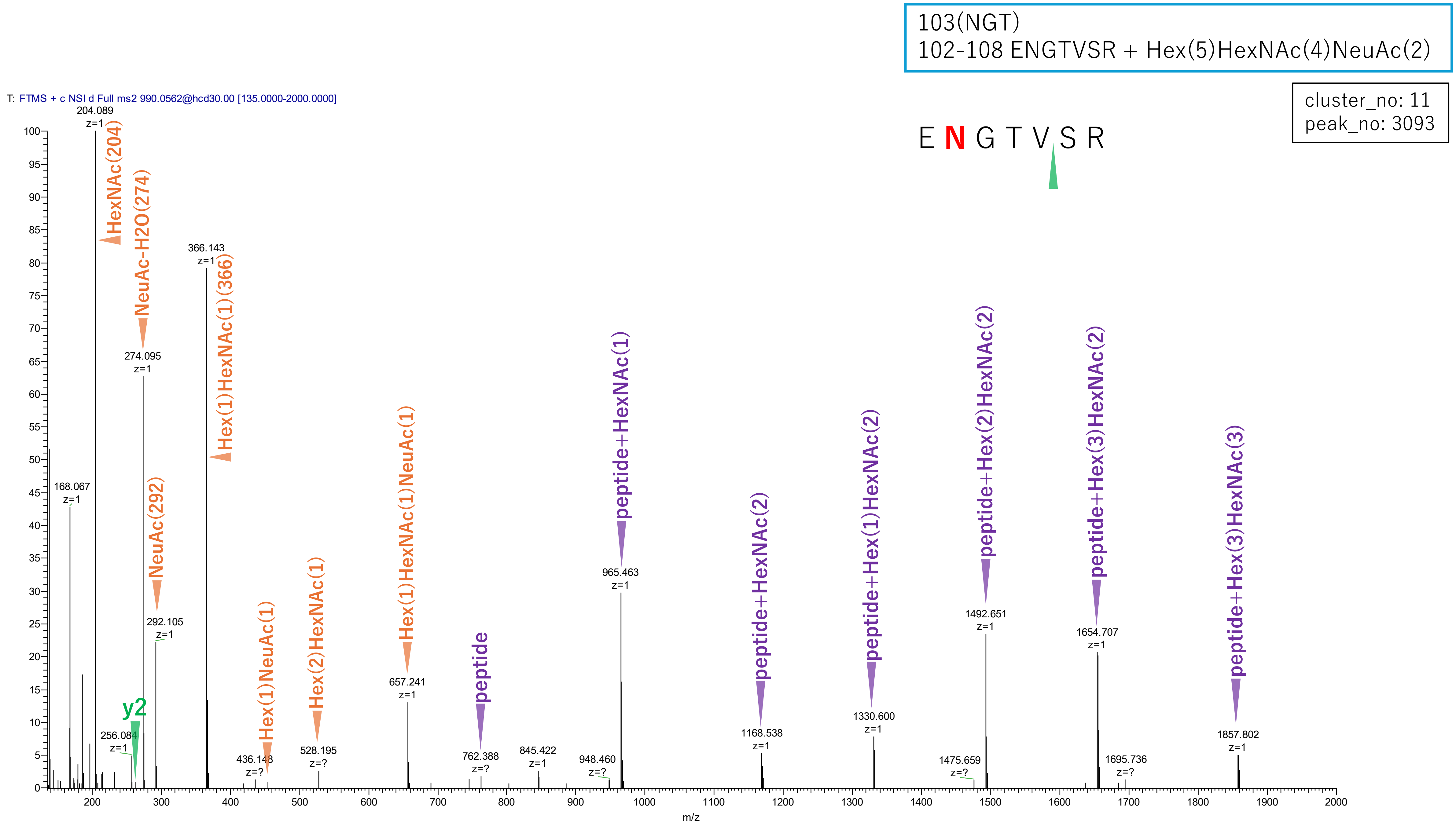


**Figure S2-71. MS2 spectra of glycopeptides assigned for hAGP.**

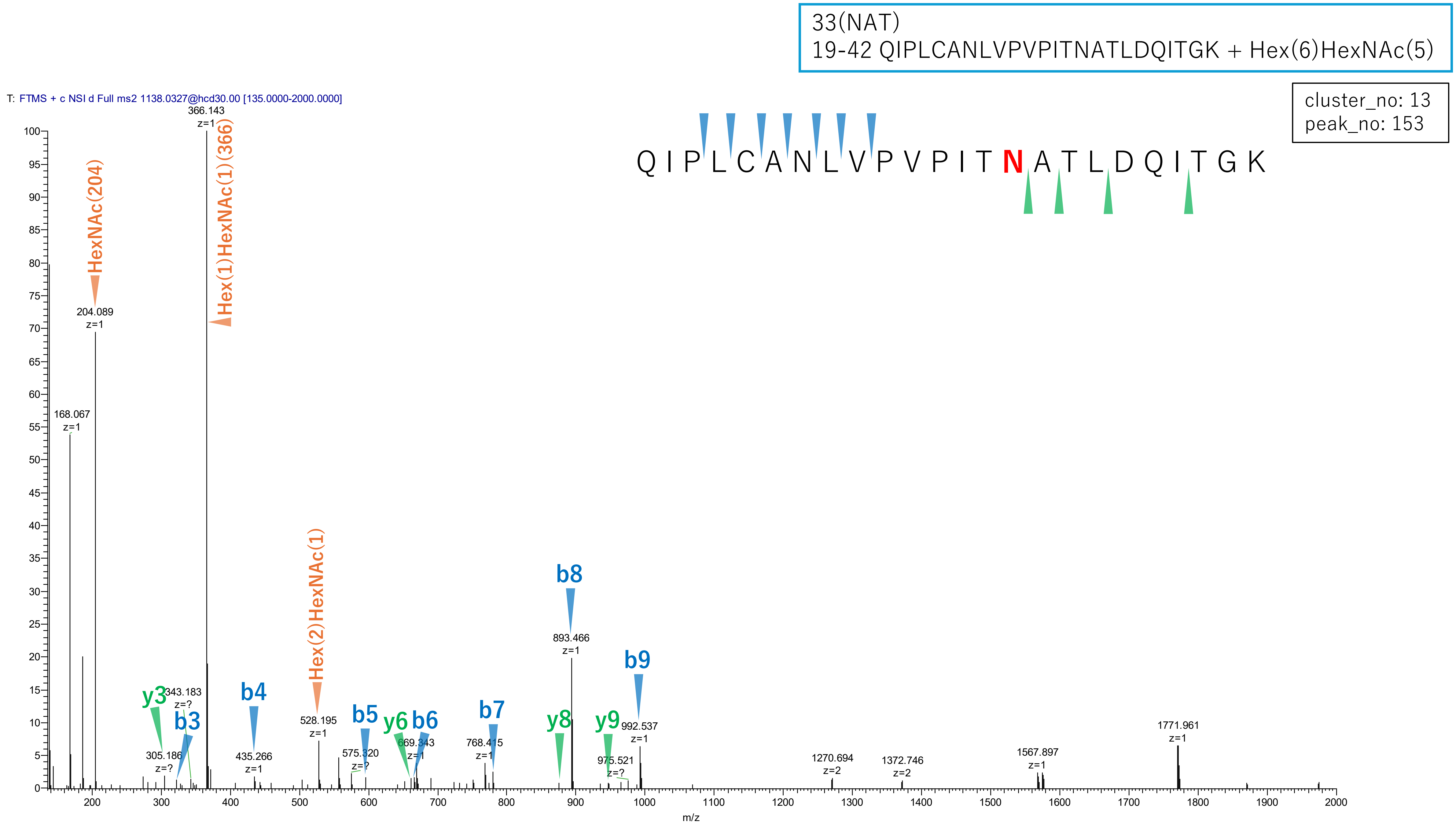


**Figure S2-72. MS2 spectra of glycopeptides assigned for hAGP.**

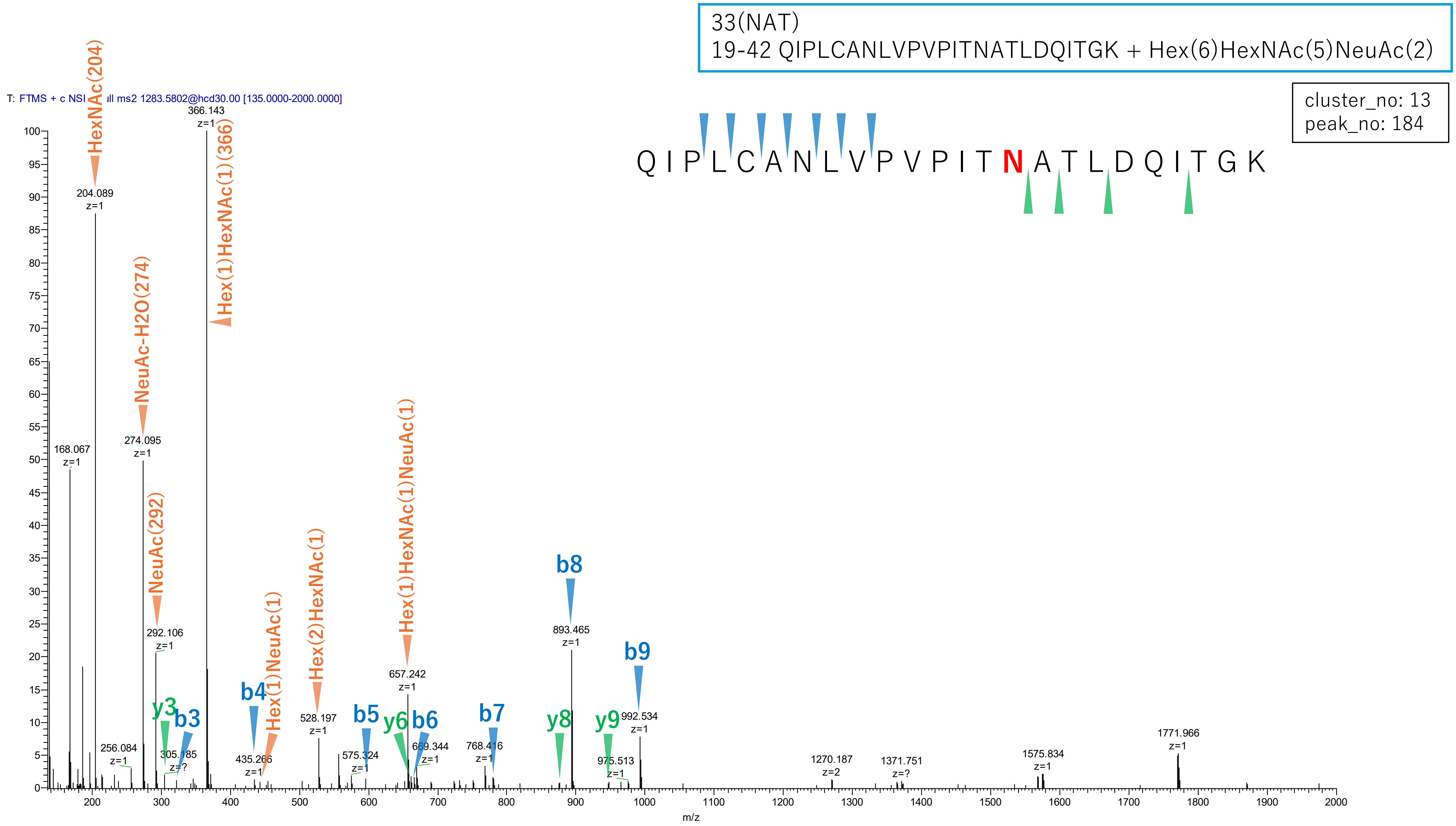


Figure S2-73. MS2 spectra of glycopeptides assigned for hAGP.

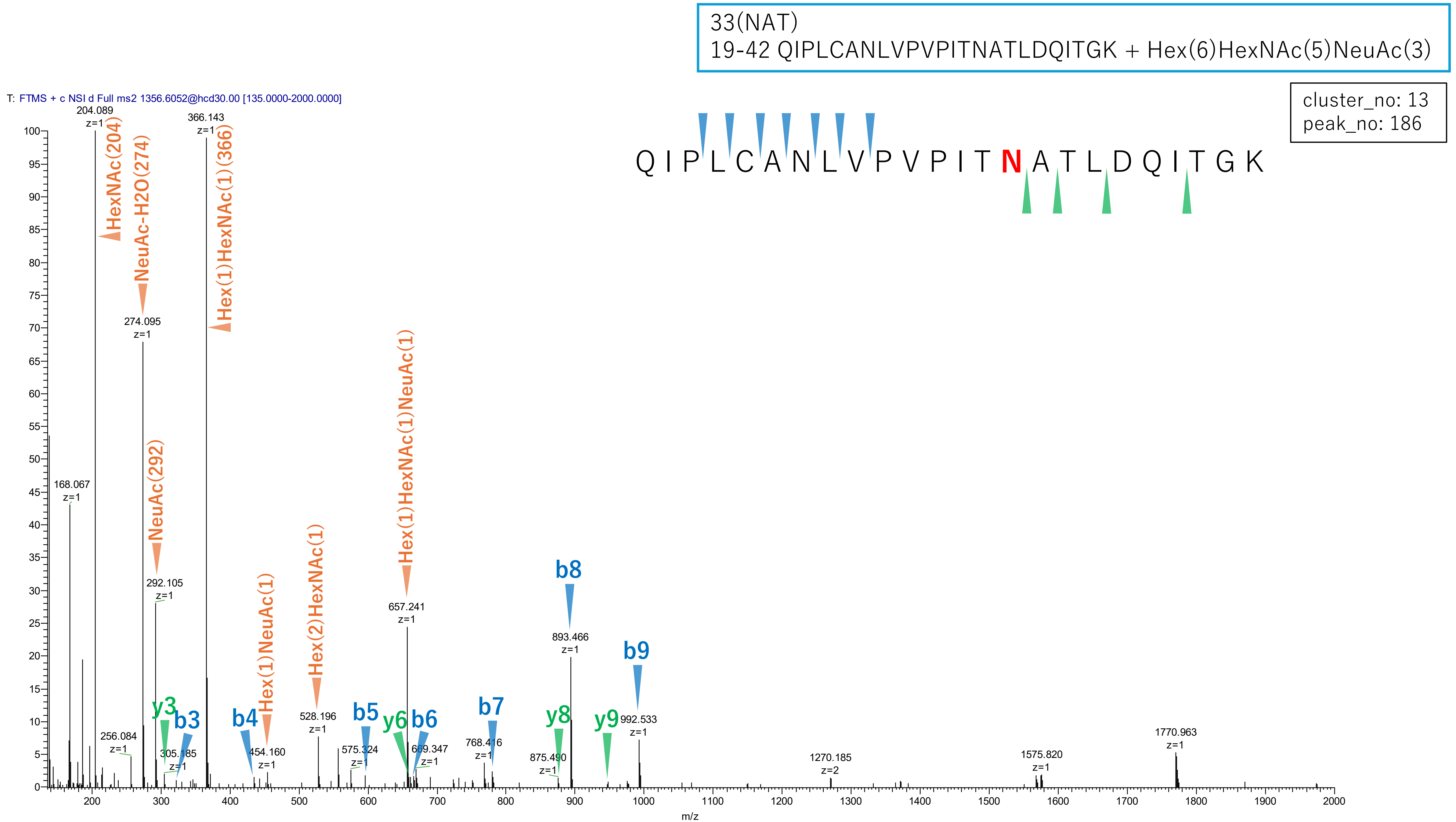


**Figure S2-74. MS2 spectra of glycopeptides assigned for hAGP.**

33(NAT)
19-42 QIPLCANLVPVPITNATLDQITGK + Hex(5)HexNAc(4)NeuAc(2)

cluster_no: 13
peak_no: 236

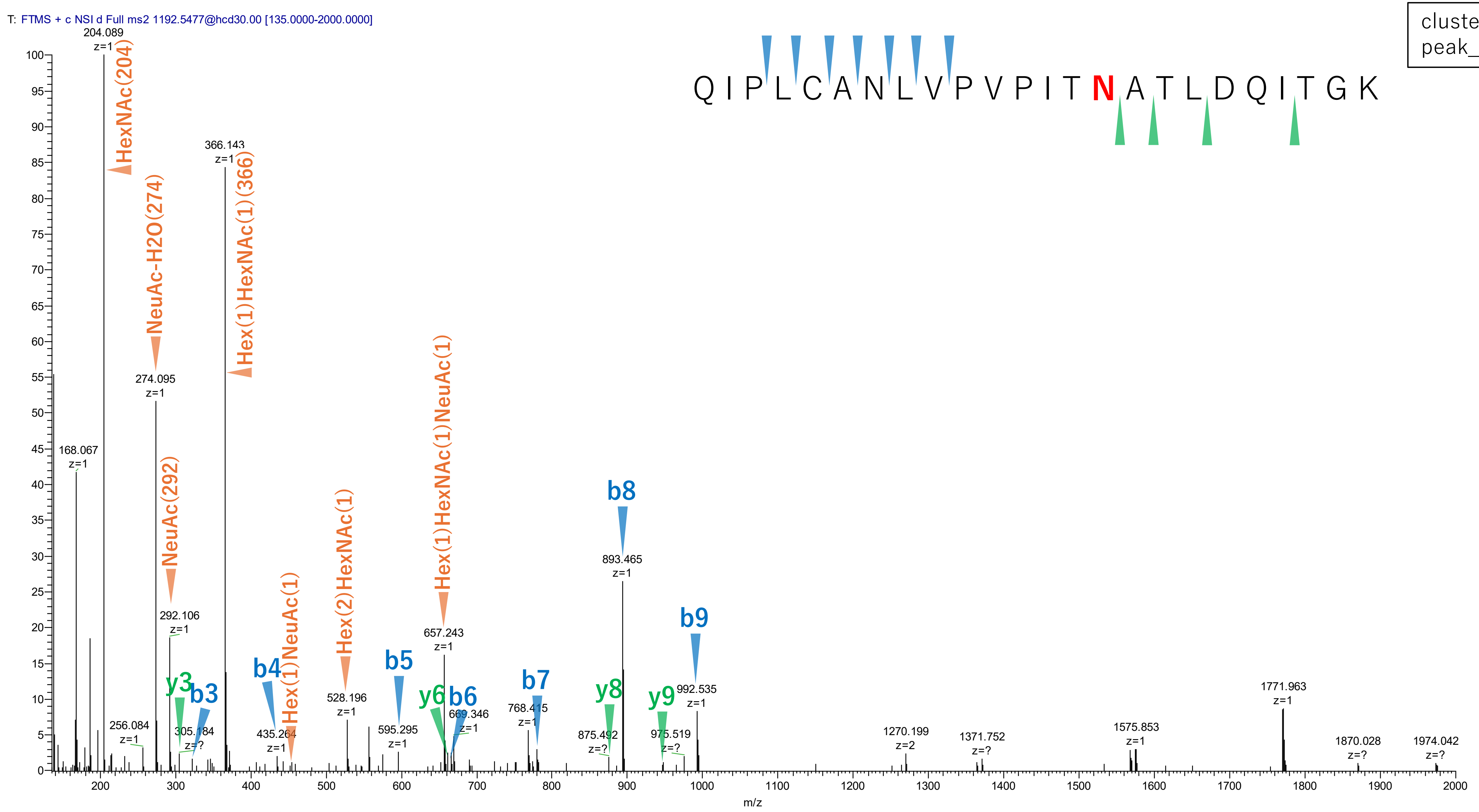


Figure S2-75. MS2 spectra of glycopeptides assigned for hAGP.

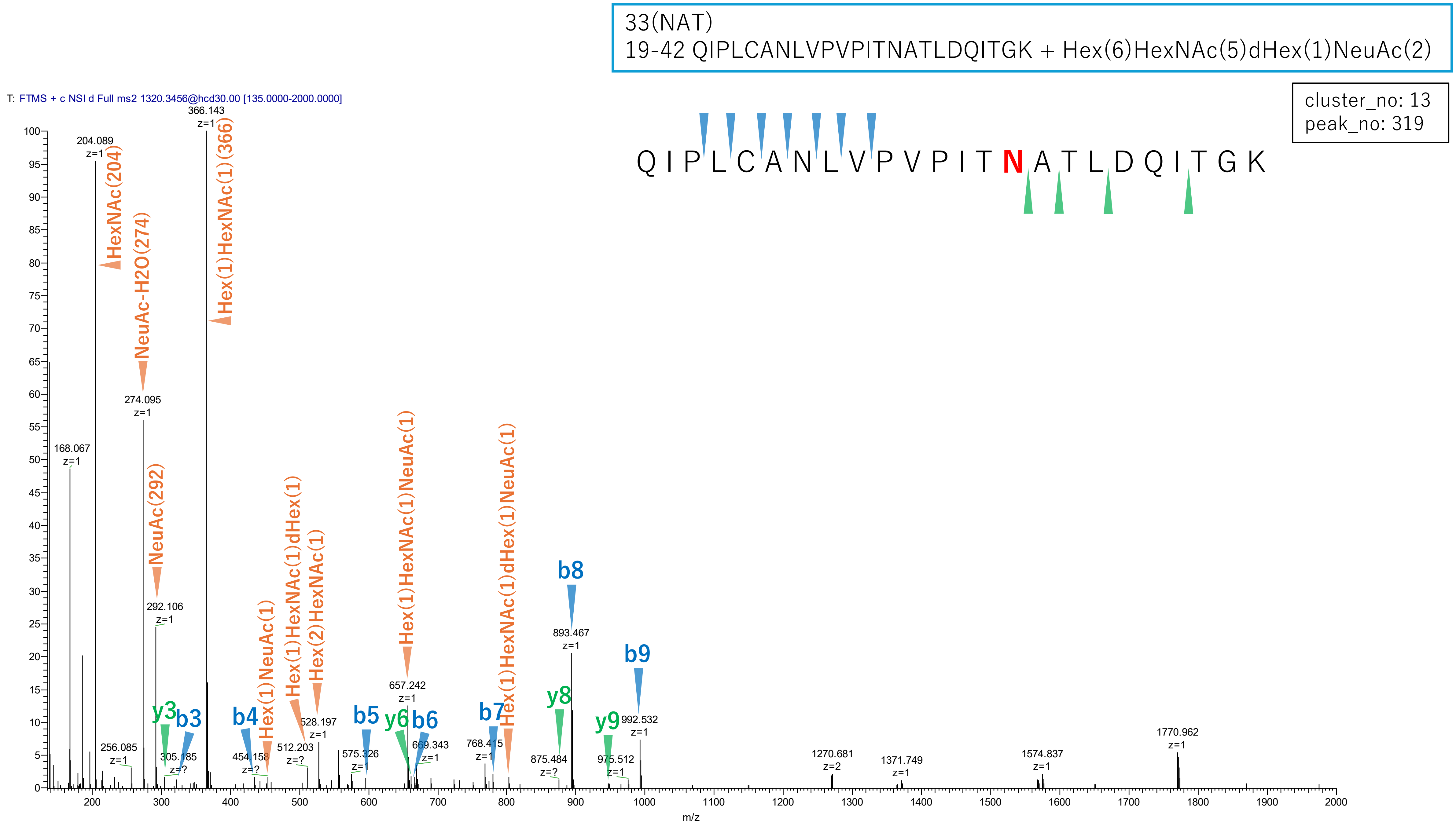


Figure S2-76. MS2 spectra of glycopeptides assigned for hAGP.

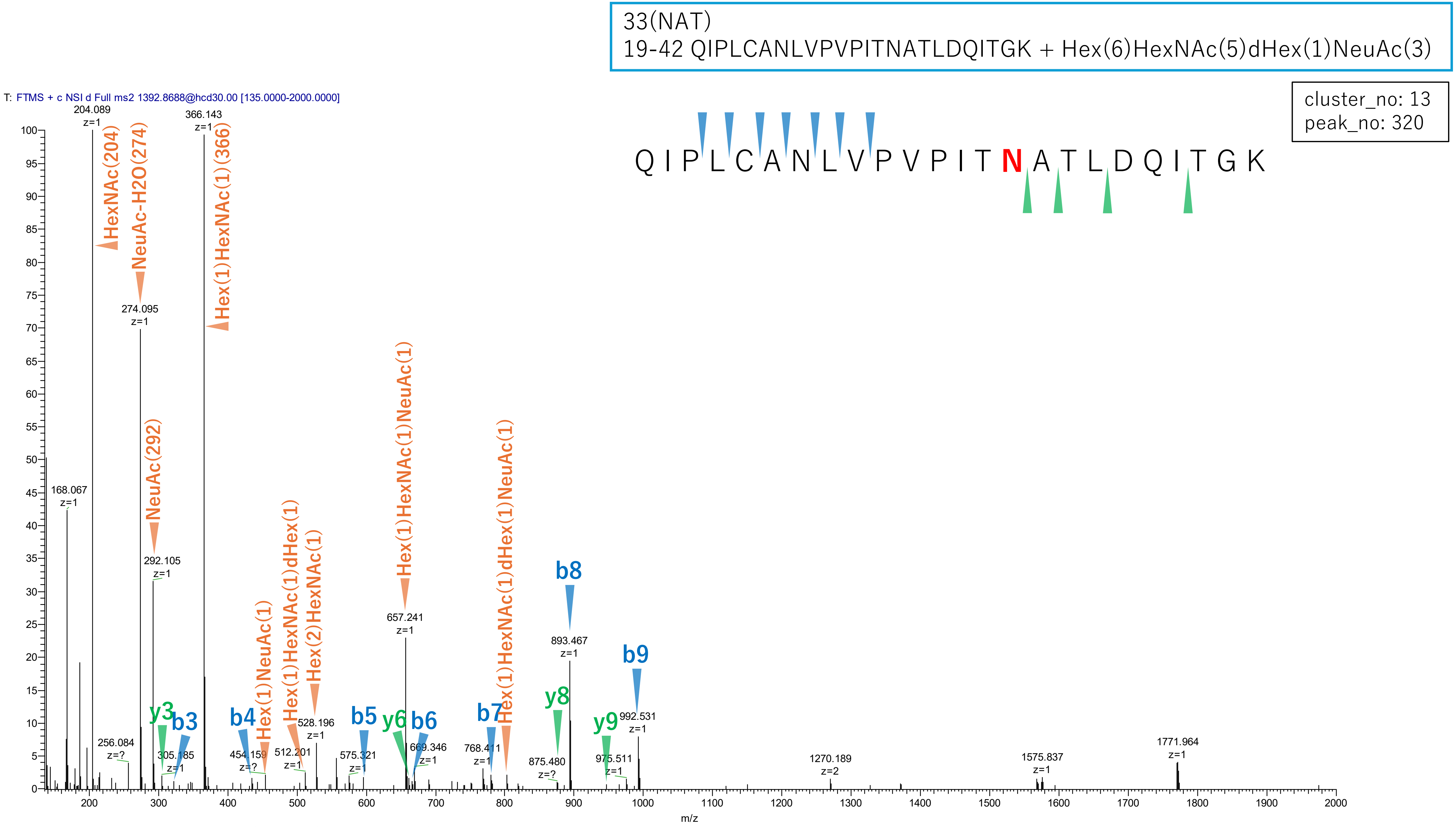


**Figure S2-77. MS2 spectra of glycopeptides assigned for hAGP.**

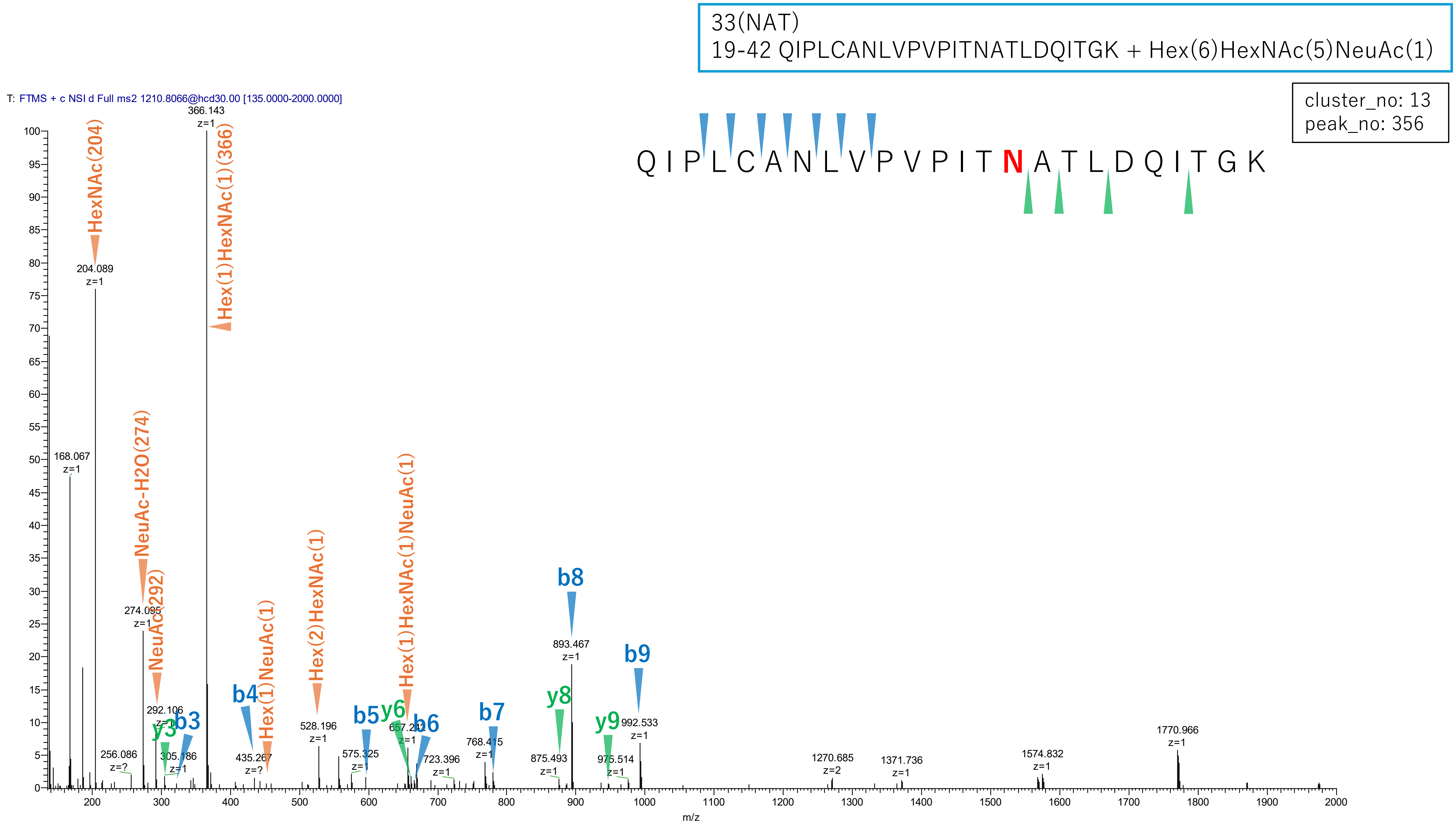


**Figure S2-78. MS2 spectra of glycopeptides assigned for hAGP.**

103(NGT)
102-113 ENGTVSRYEGGR + Hex(6)HexNAc(5)NeuAc(3)

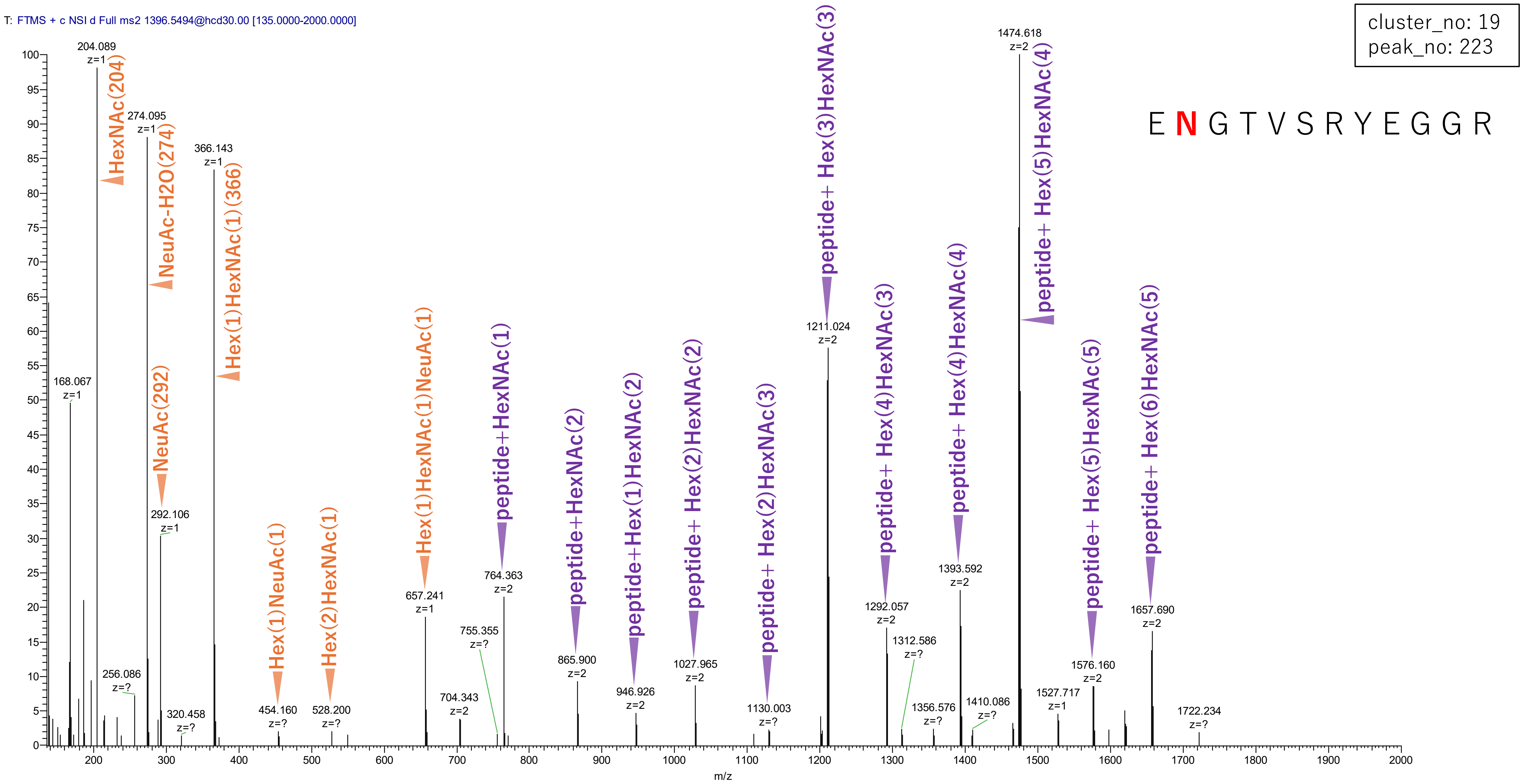


Figure S2-79. MS2 spectra of glycopeptides assigned for hAGP.

103(NGT)
102-113 ENGTVSRYEGGR + Hex(7)HexNAc(6)NeuAc(4)

cluster_no: 19
peak_no: 343

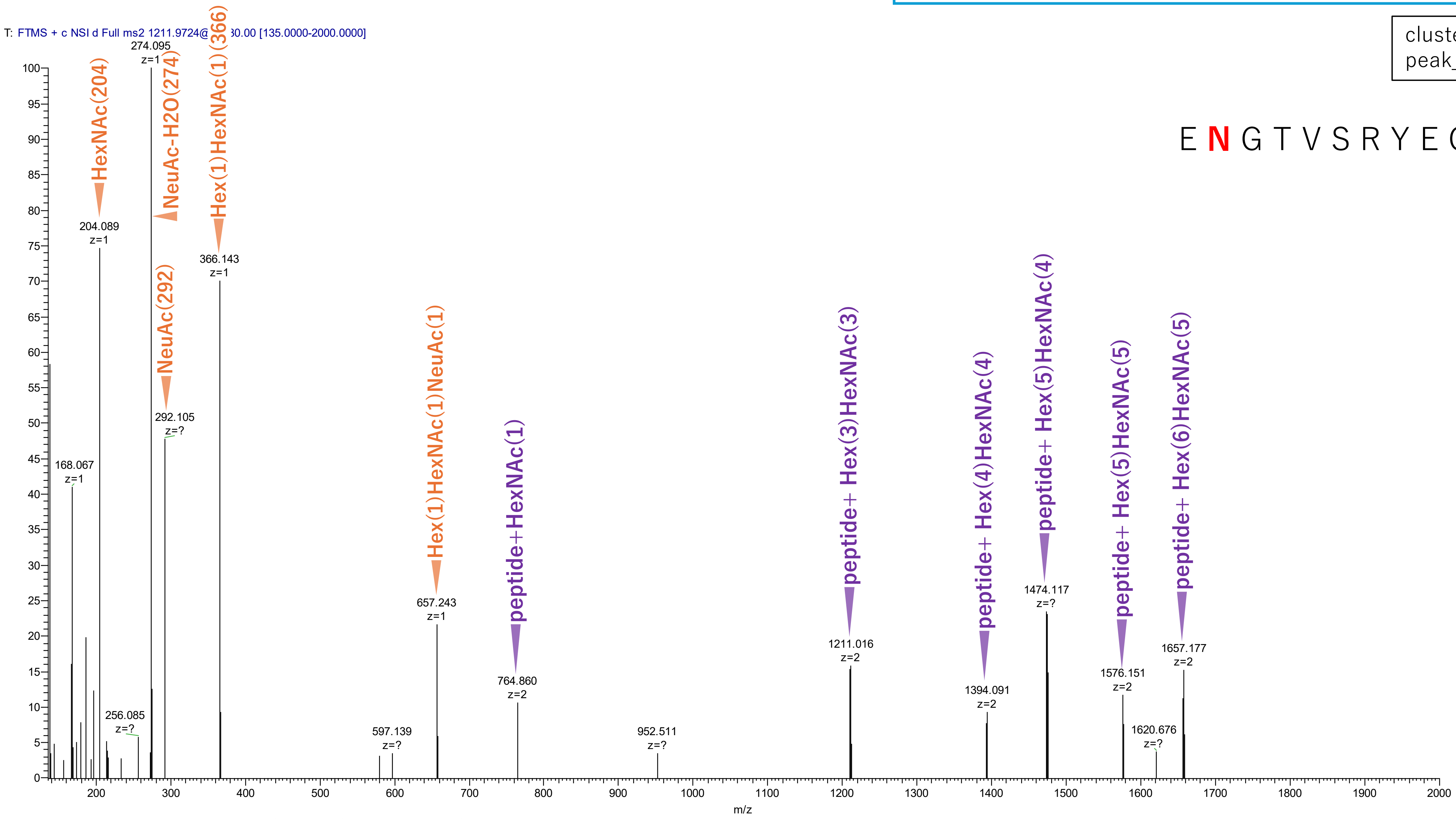


Figure S2-80. MS2 spectra of glycopeptides assigned for hAGP.

103(NGT)
102-113 ENGTVSRYEGGR + Hex(6)HexNAc(5)dHex(1)NeuAc(3)

cluster_no: 19
peak_no: 352

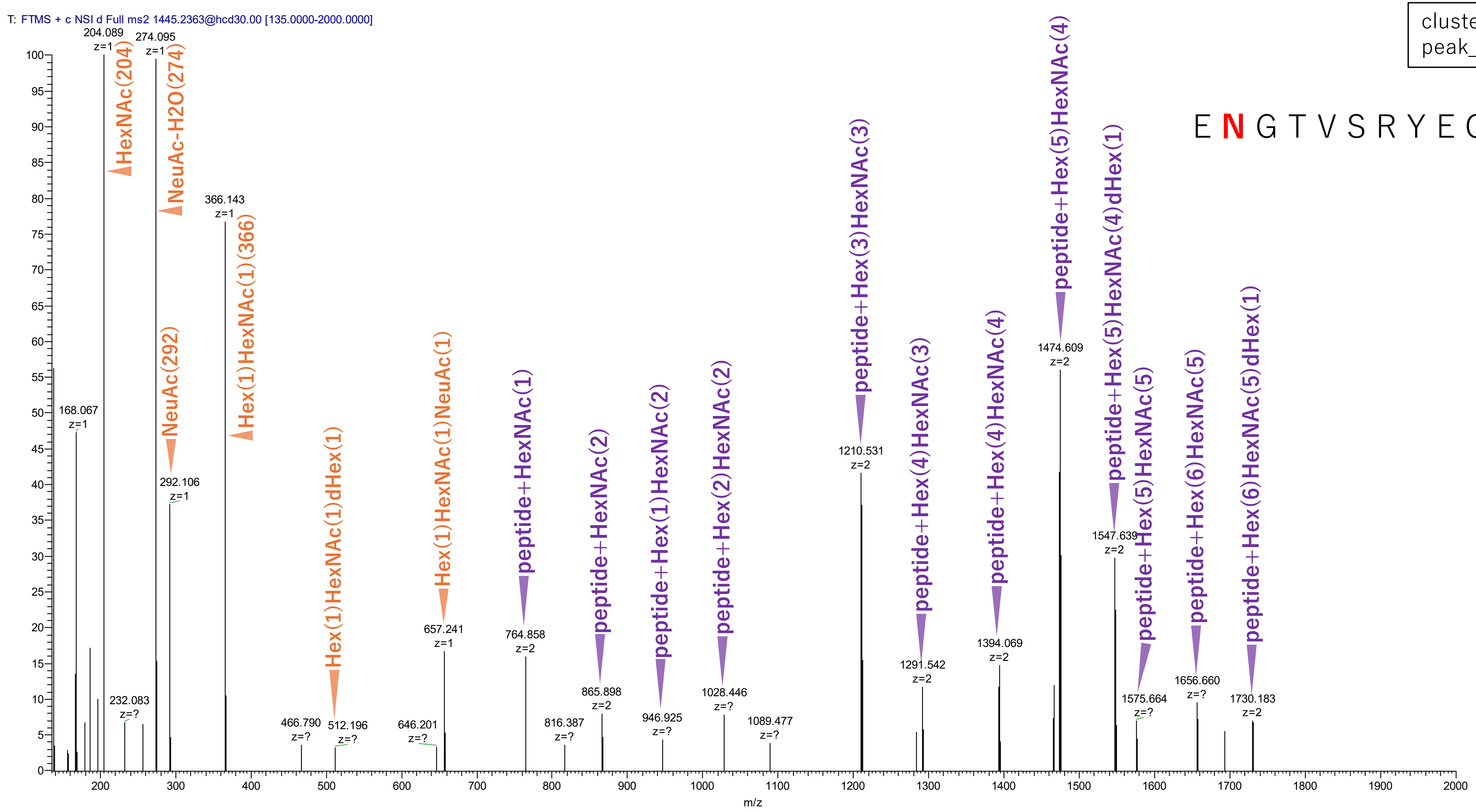


**Figure S2-81. MS2 spectra of glycopeptides assigned for hAGP.**

103(NGT)
102-113 ENGTVSRYEGGR + Hex(7)HexNAc(6)dHex(1)NeuAc(4)

cluster_no: 19
peak_no: 479

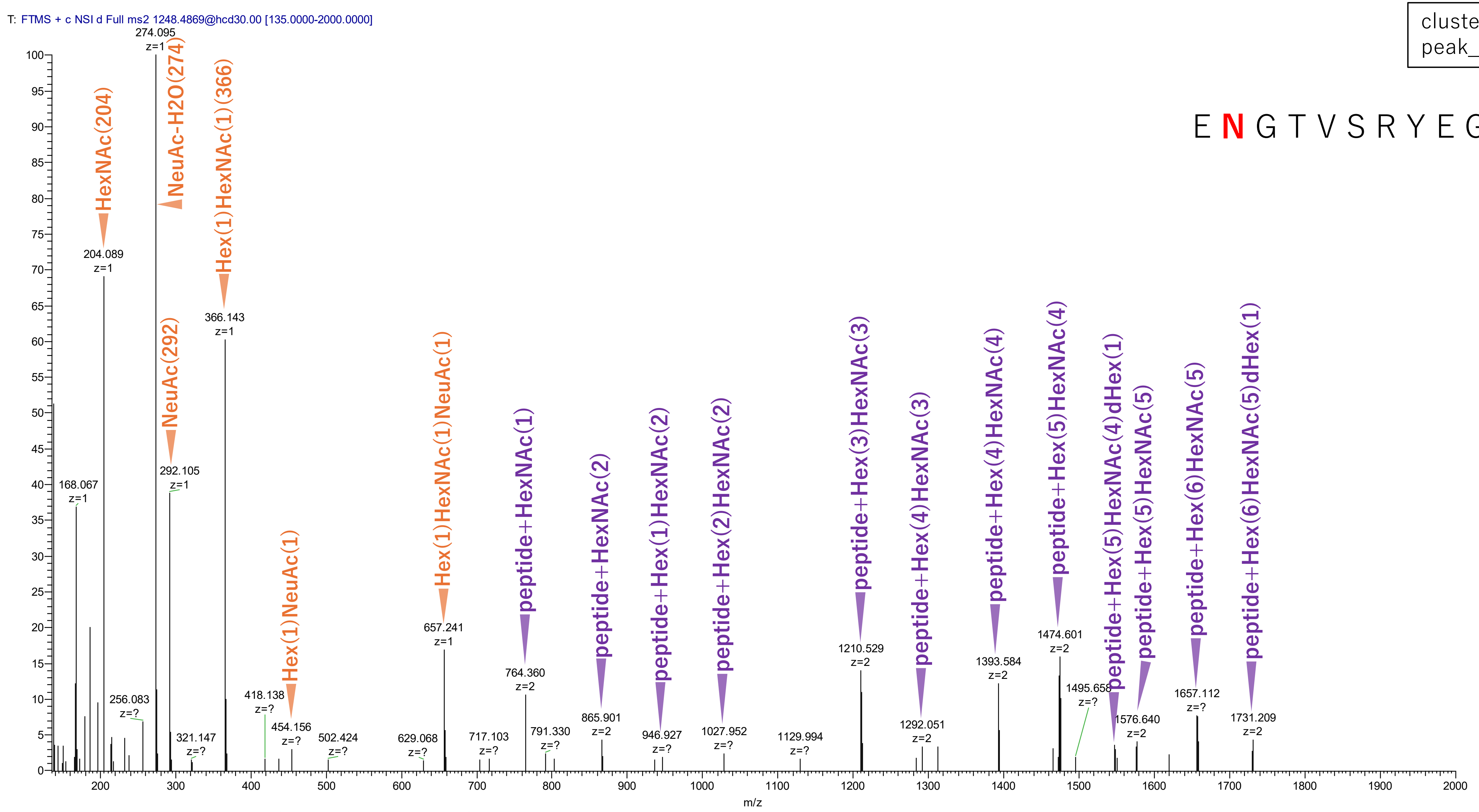


**Figure S2-82. MS2 spectra of glycopeptides assigned for hAGP.**

103(NGT)
102-113 ENGTVSRYEGGR + Hex(7)HexNAc(6)NeuAc(3)

cluster_no: 19
peak_no: 587

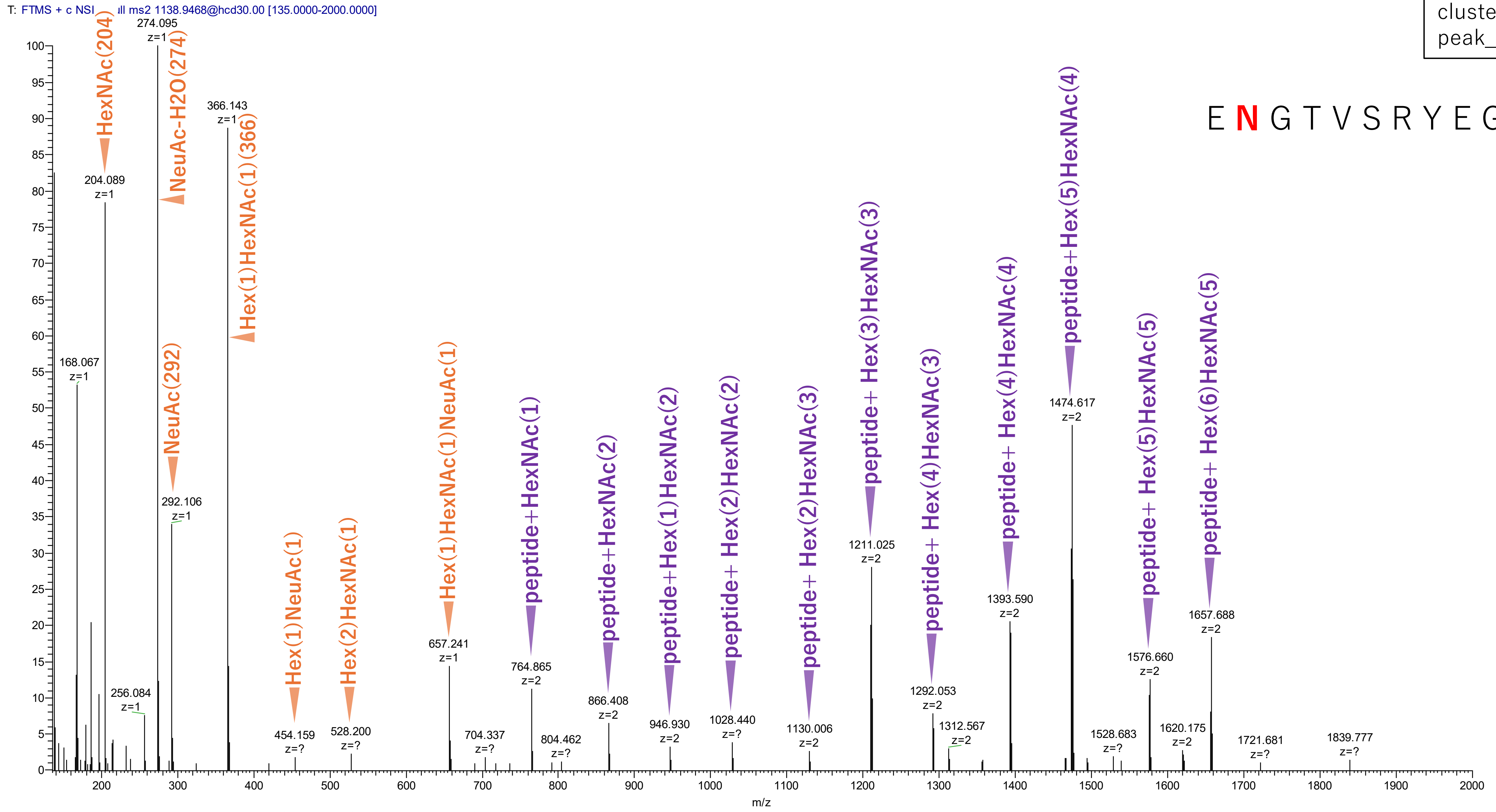


Figure S2-83. MS2 spectra of glycopeptides assigned for hAGP.

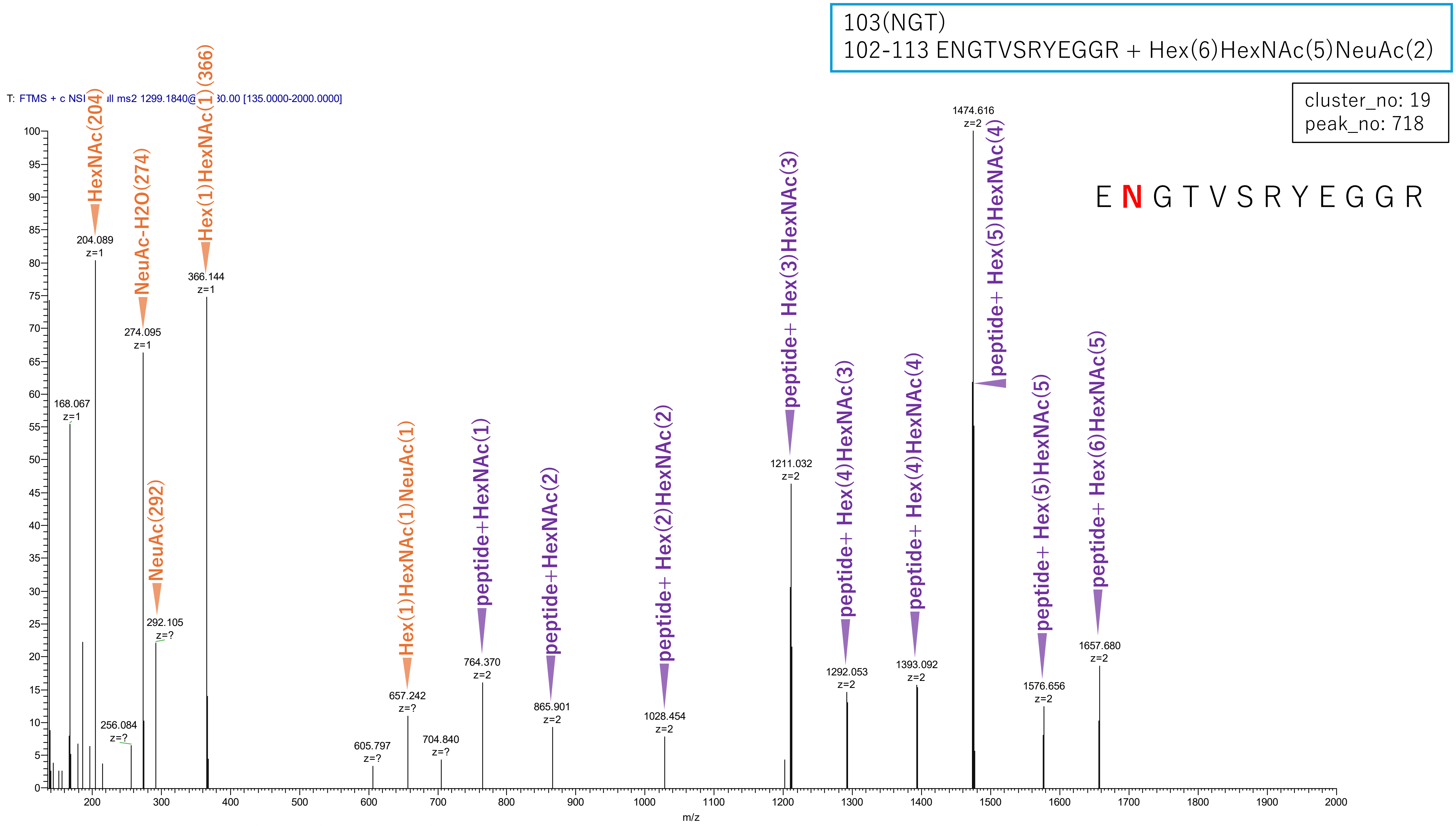


**Figure S2-84. MS2 spectra of glycopeptides assigned for hAGP.**

103(NGT)
102-113 ENGTVSRYEGGR + Hex(5)HexNAc(4)NeuAc(2)

cluster_no: 19
peak_no: 723

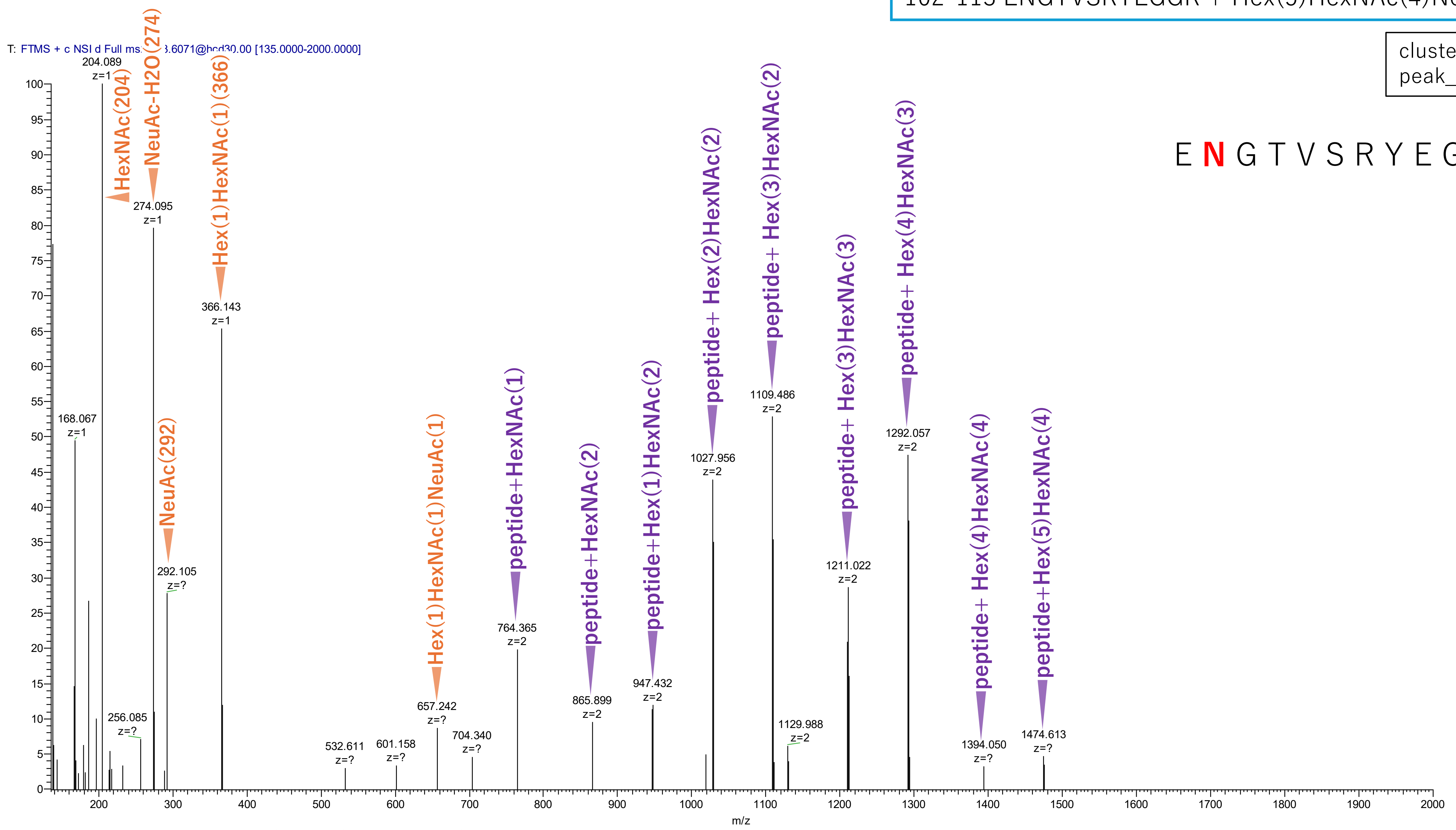


**Figure S2-85. MS2 spectra of glycopeptides assigned for hAGP.**

103(NGT)
102-113 ENGTVSRYEGGR + Hex(7)HexNAc(6)dHex(1)NeuAc(3)

cluster_no: 19
peak_no: 853

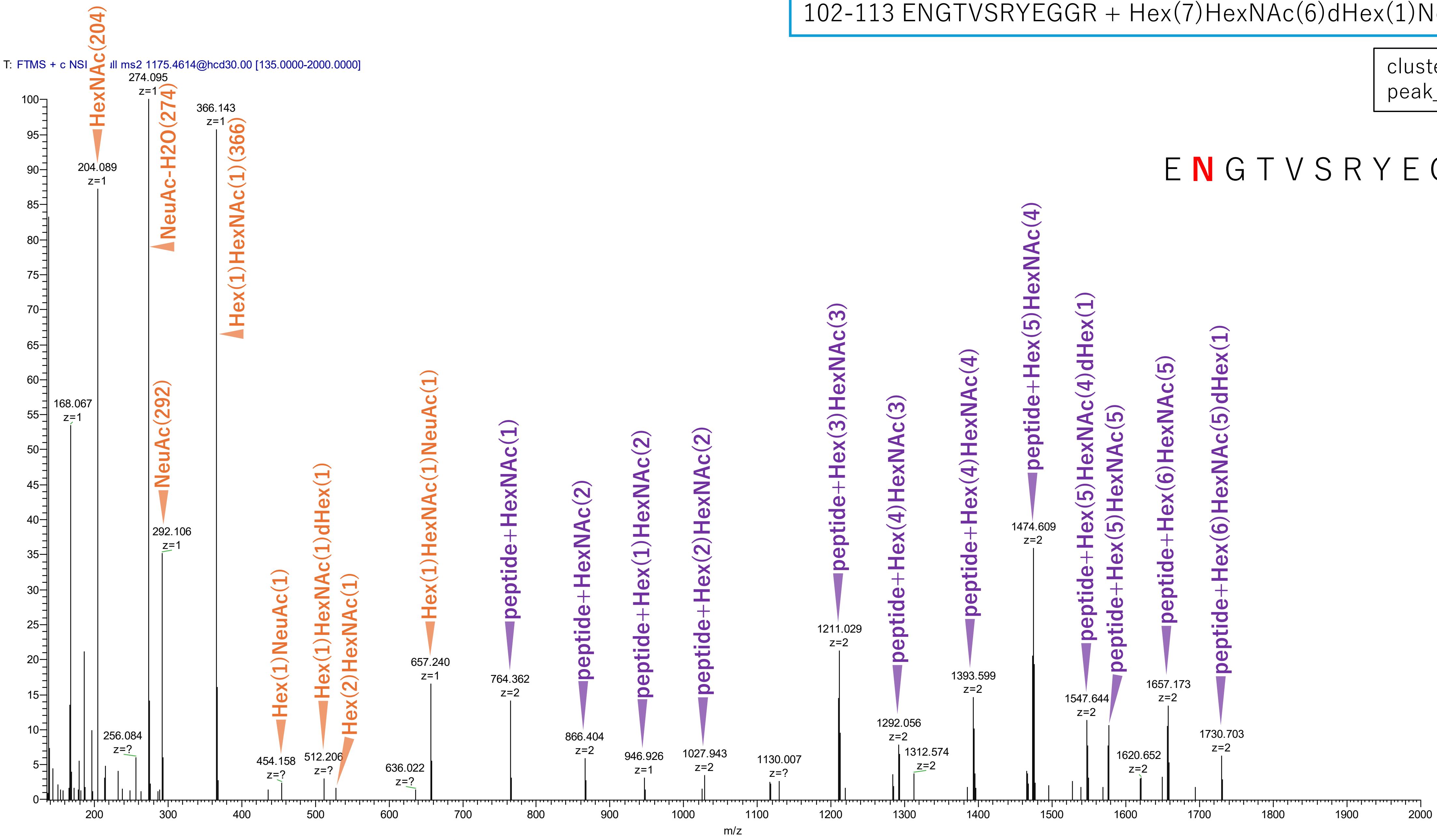


**Figure S2-86. MS2 spectra of glycopeptides assigned for hAGP.**

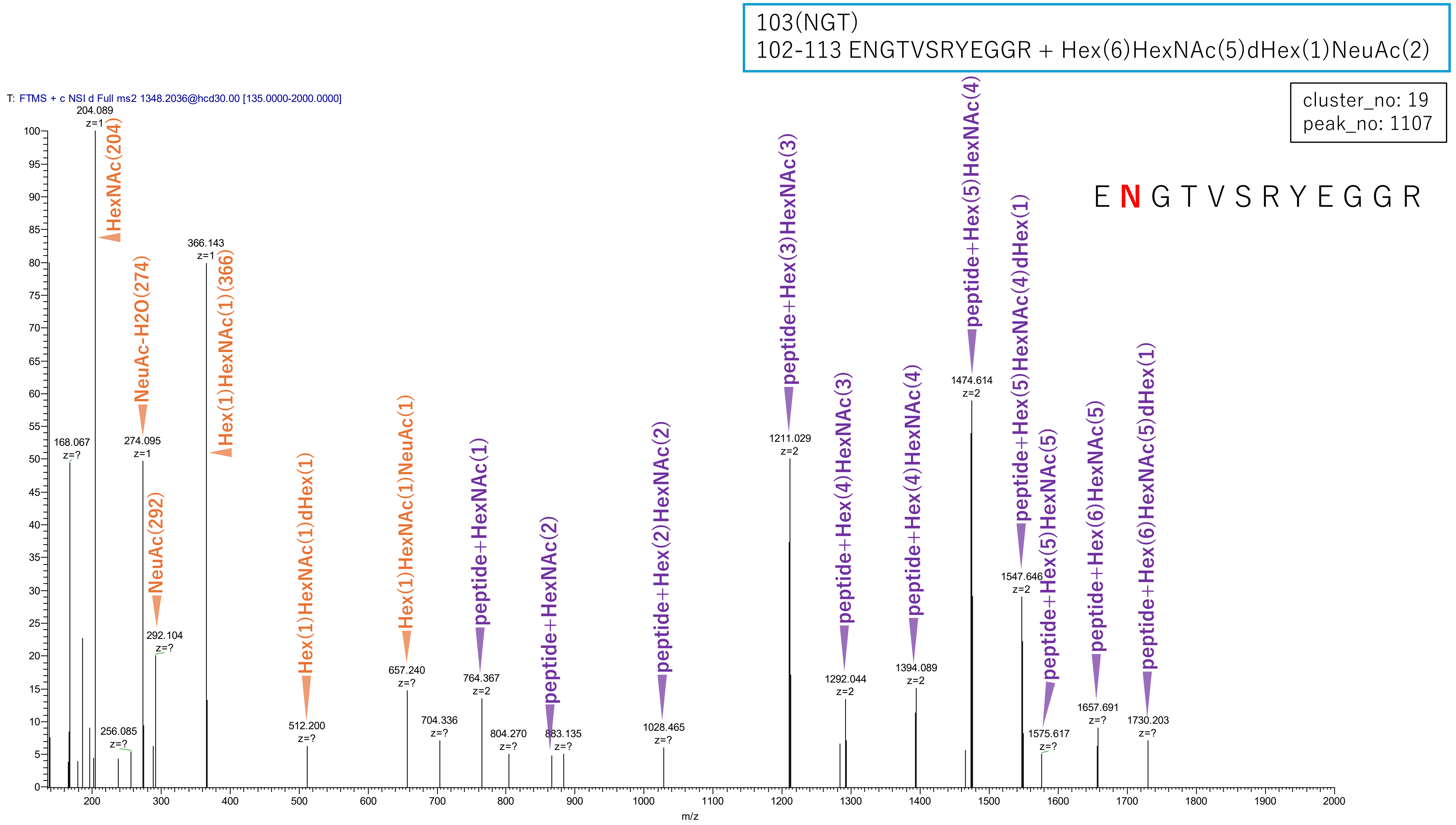


**Figure S2-87. MS2 spectra of glycopeptides assigned for hAGP.**

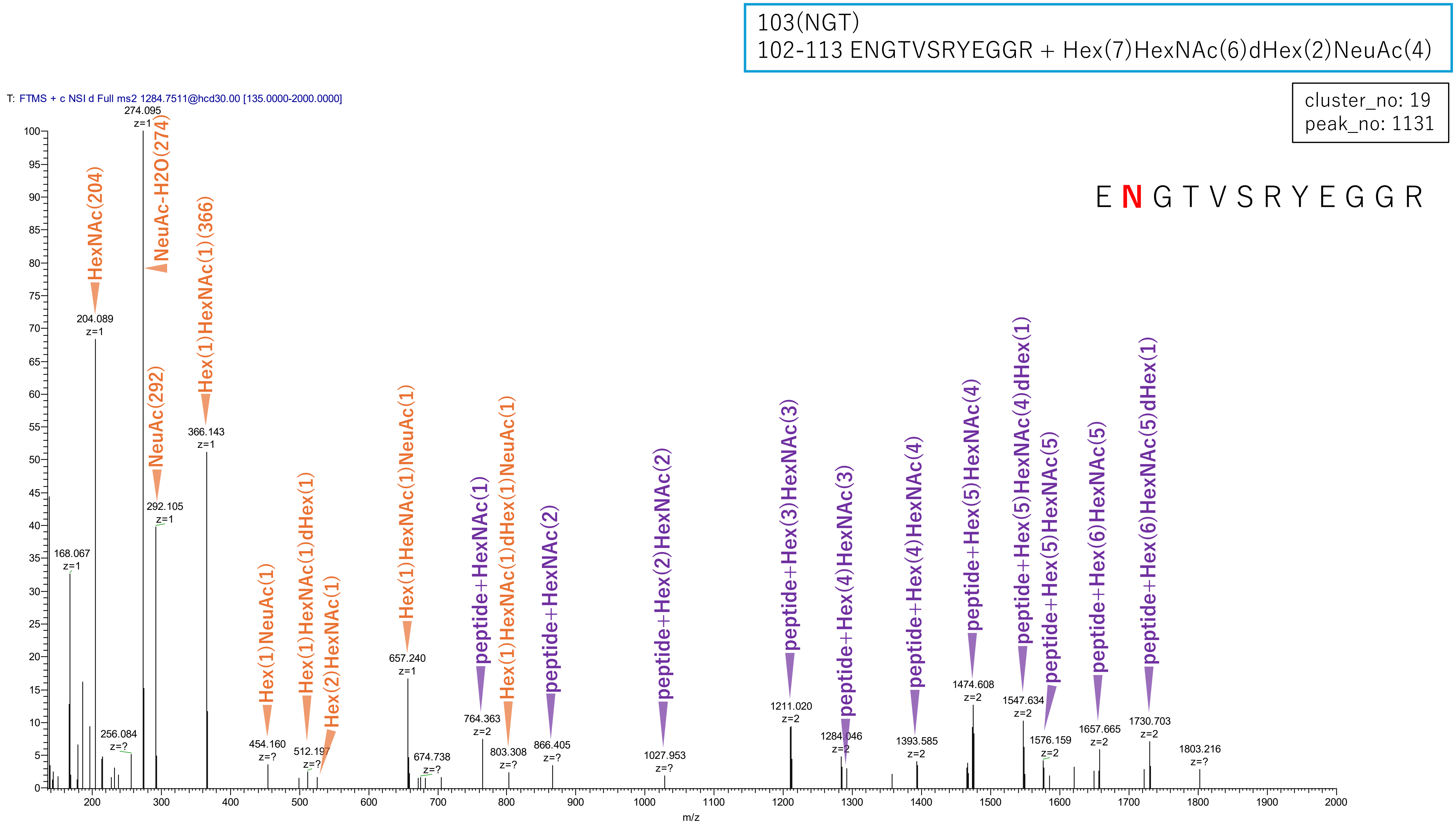


**Figure S2-88. MS2 spectra of glycopeptides assigned for hAGP.**

103(NGT)
102-113 ENGTVSRYEGGR + Hex(7)HexNAc(6)dHex(3)NeuAc(4)

cluster_no: 19
peak_no: 1956

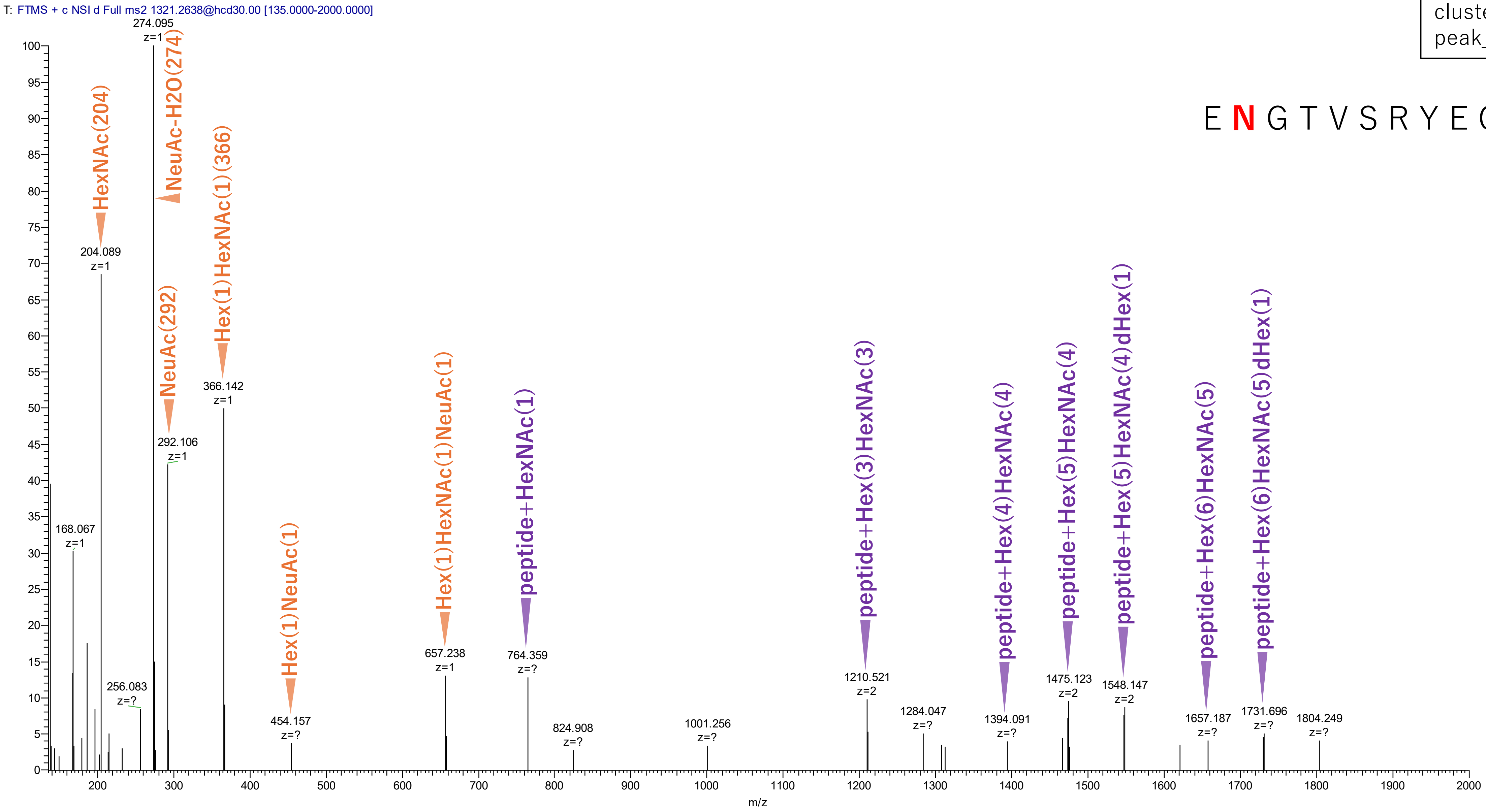


**Figure S2-89. MS2 spectra of glycopeptides assigned for hAGP.**

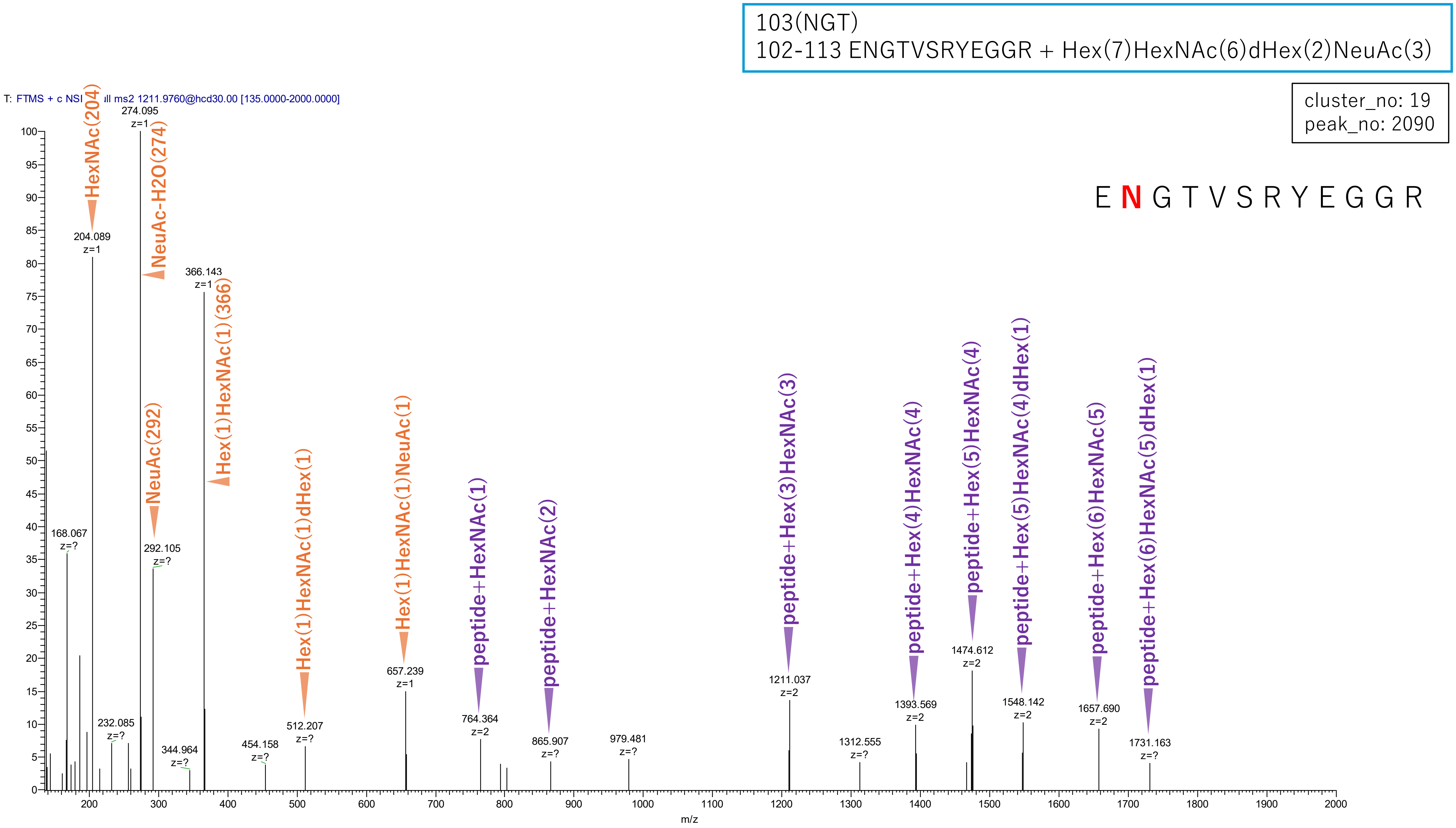


**Figure S2-90. MS2 spectra of glycopeptides assigned for hAGP.**

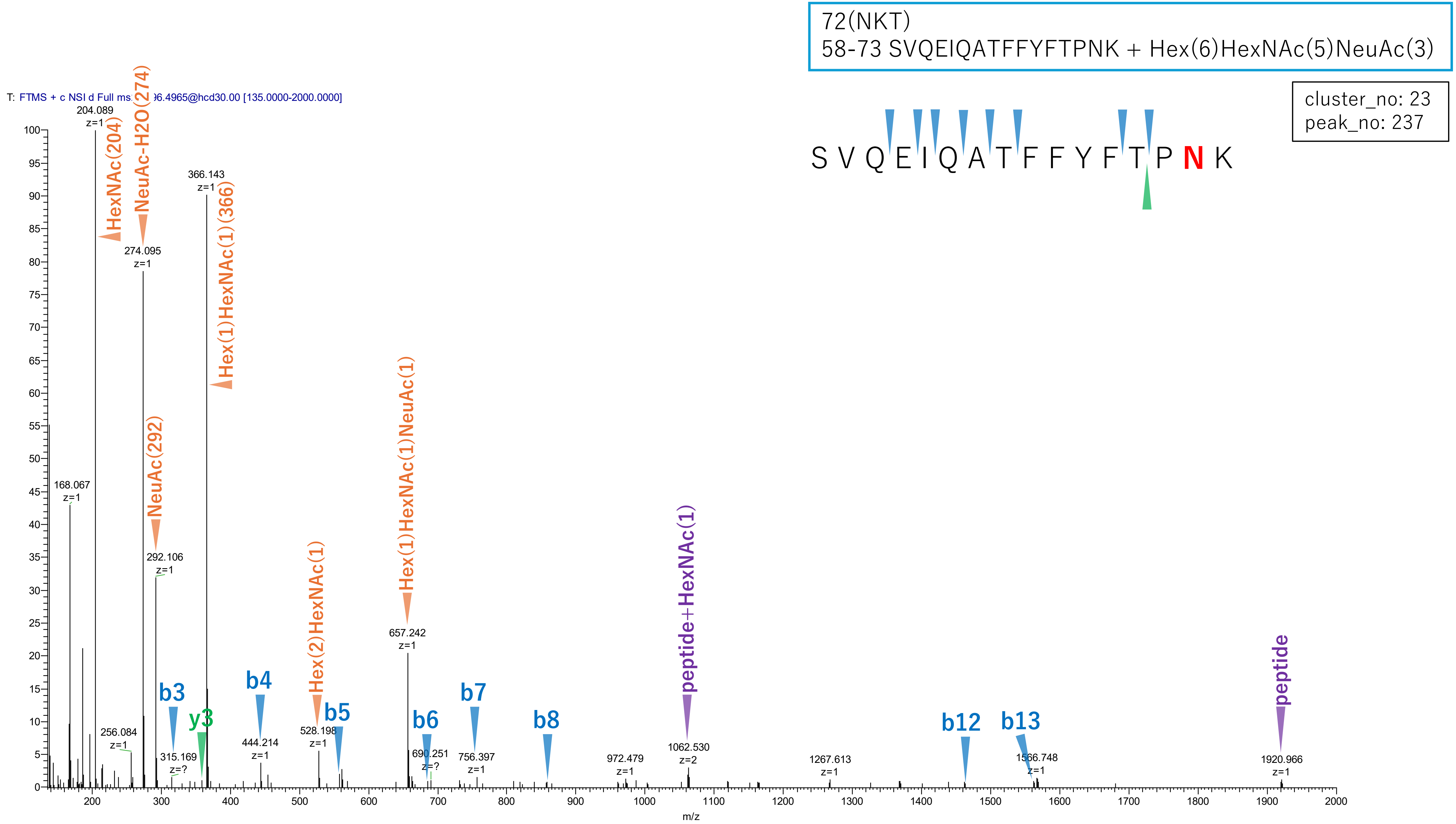


Figure S2-91. MS2 spectra of glycopeptides assigned for hAGP.

72(NKT)
58-73 SVQEIQATFFYFTPNK + Hex(5)HexNAc(4)NeuAc(2)

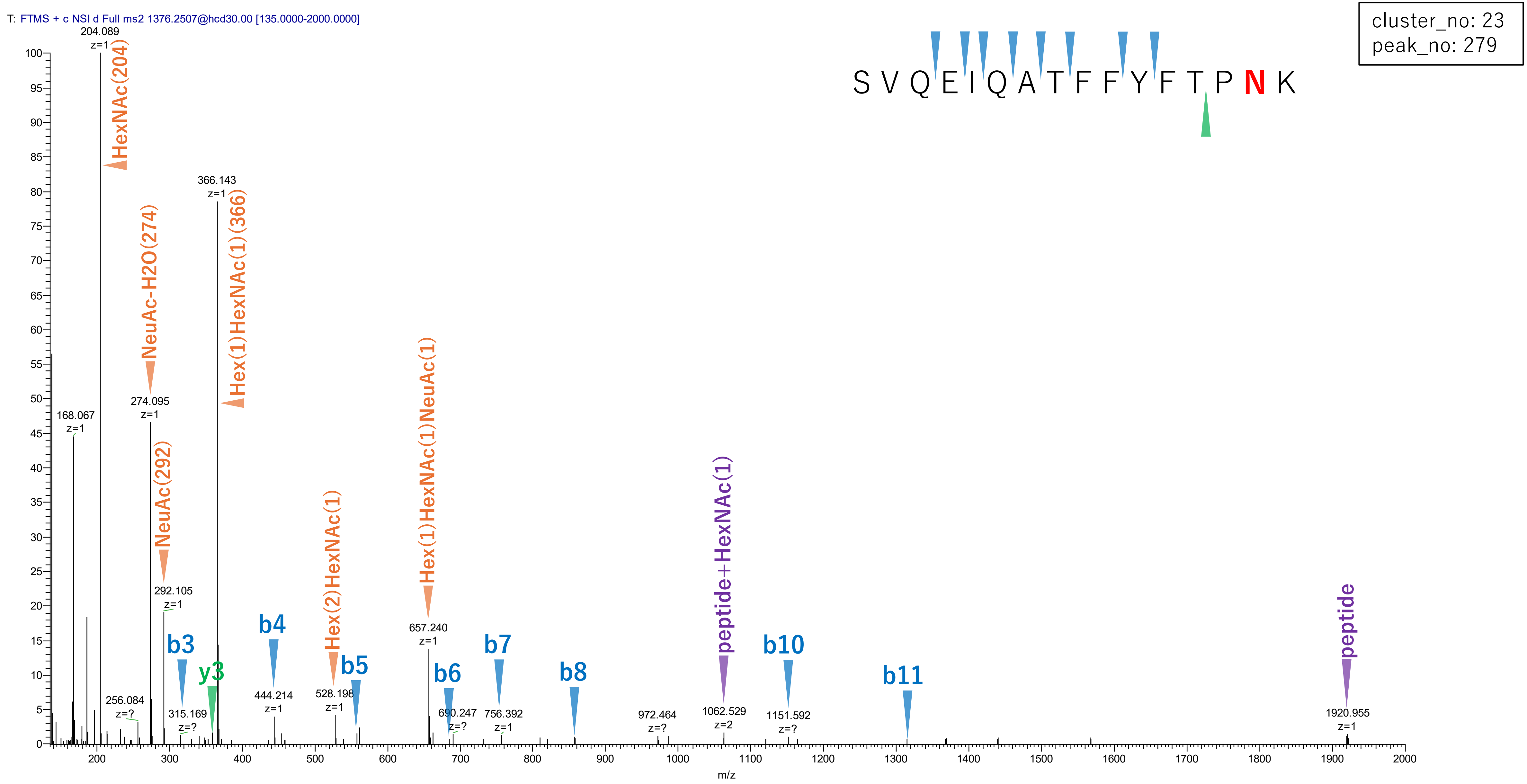


Figure S2-92. MS2 spectra of glycopeptides assigned for hAGP.

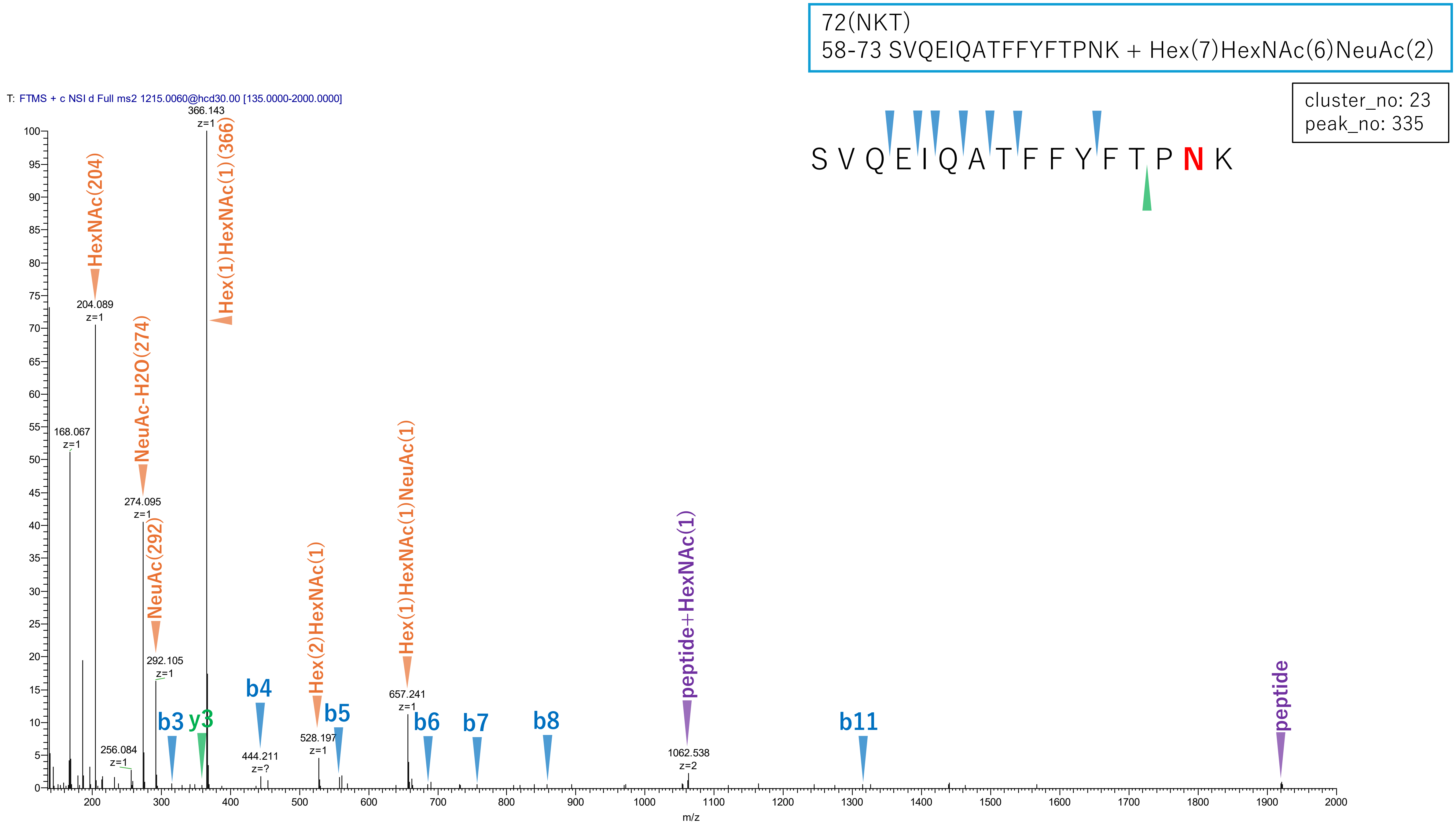


**Figure S2-93. MS2 spectra of glycopeptides assigned for hAGP.**

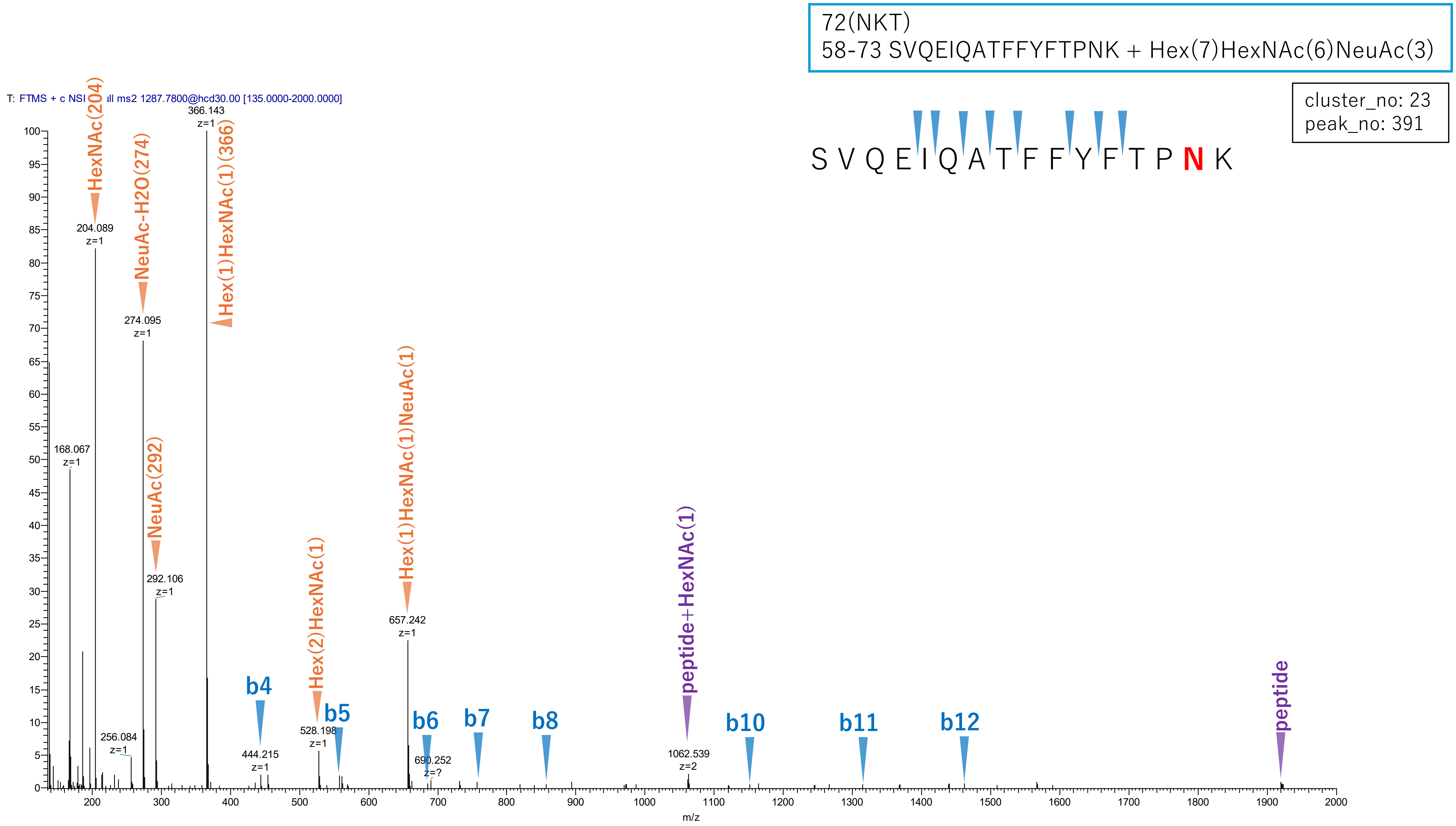


Figure S2-94. MS2 spectra of glycopeptides assigned for hAGP.

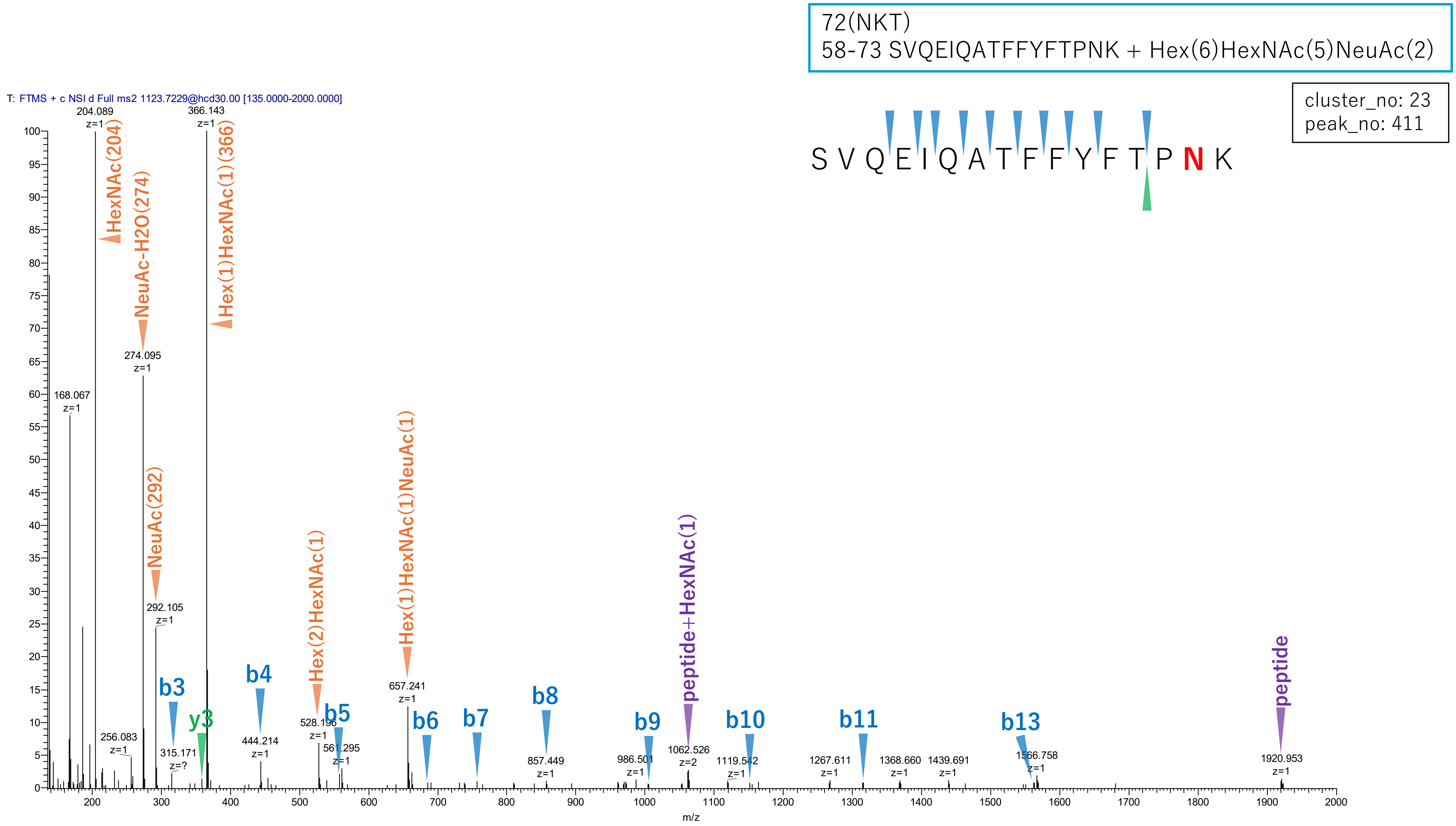


**Figure S2-95. MS2 spectra of glycopeptides assigned for hAGP.**

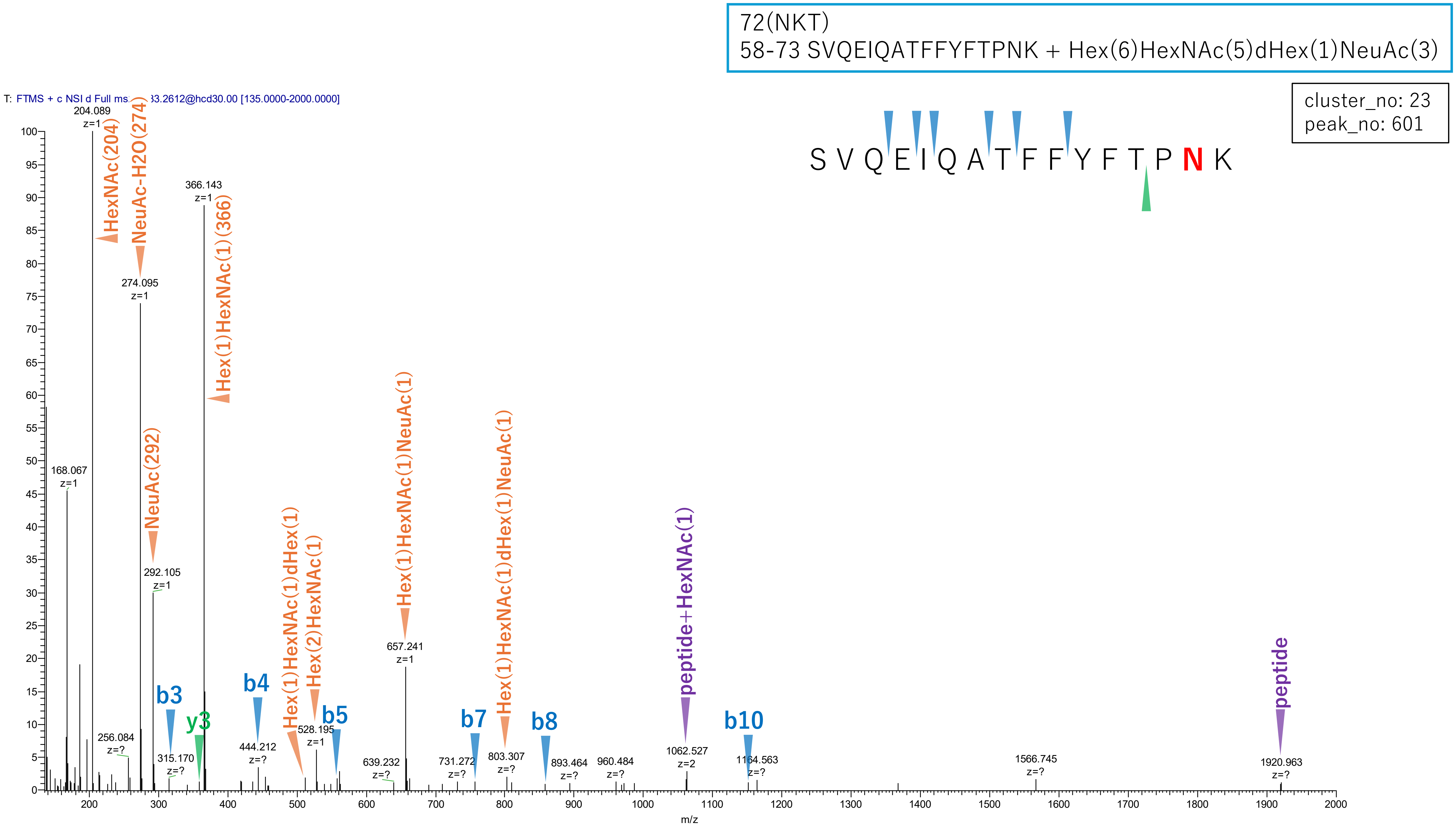


**Figure S2-96. MS2 spectra of glycopeptides assigned for hAGP.**

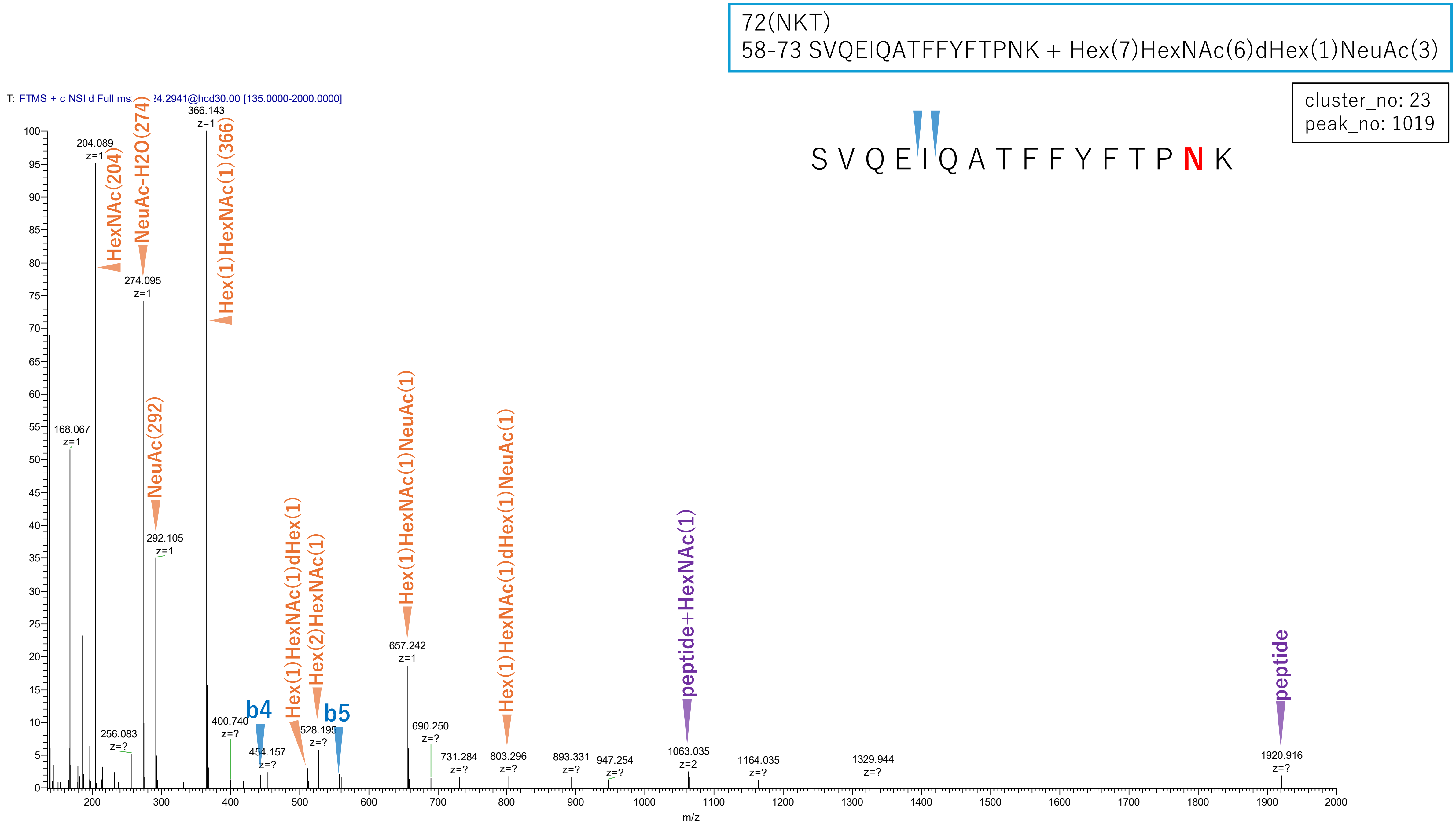


**Figure S2-97. MS2 spectra of glycopeptides assigned for hAGP.**

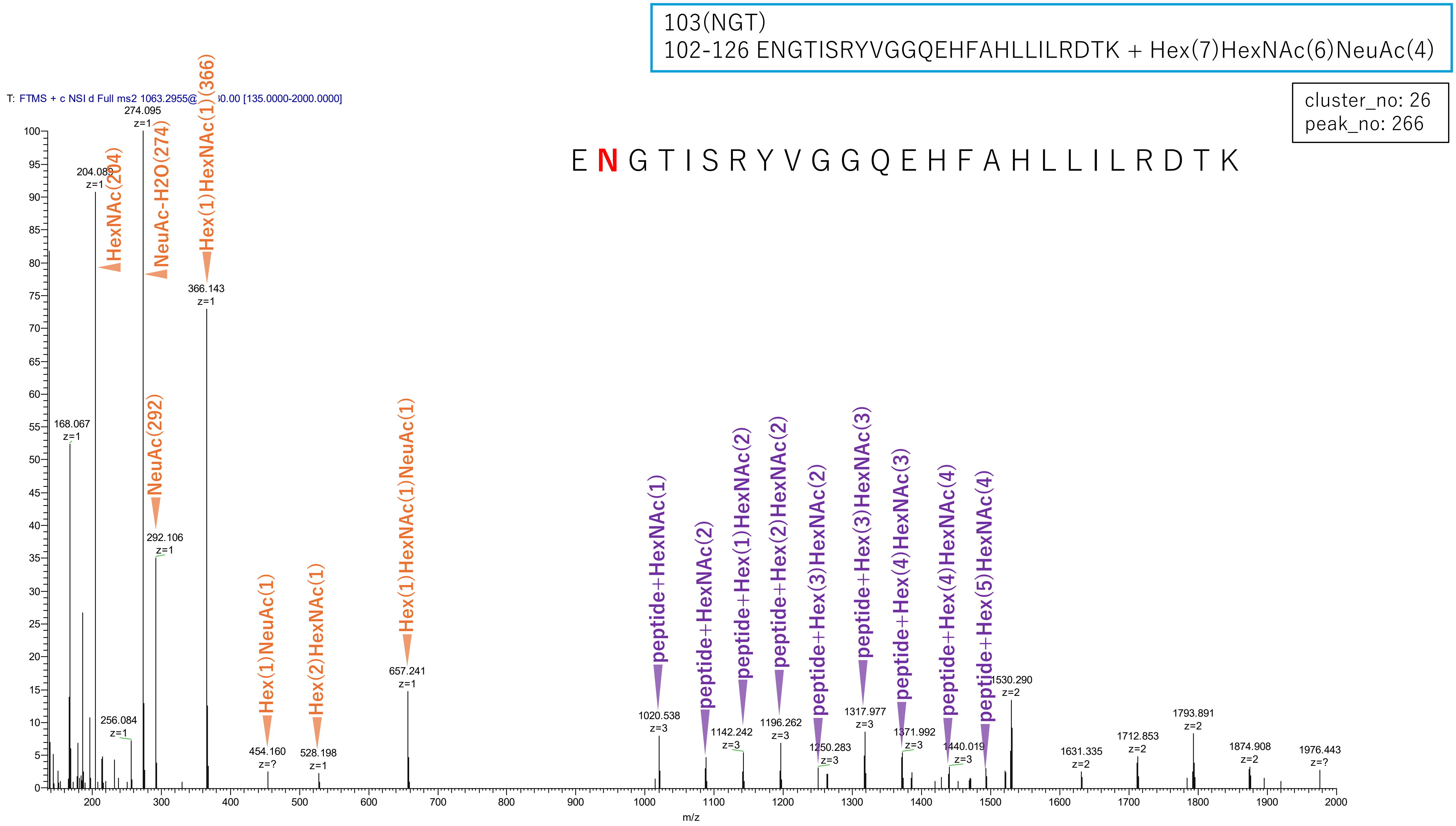


**Figure S2-98. MS2 spectra of glycopeptides assigned for hAGP.**

103(NGT)
102-126 ENGTISRYVGGQEHFAHLLILRDTK + Hex(6)HexNAc(5)NeuAc(3)

cluster_no: 26
peak_no: 465

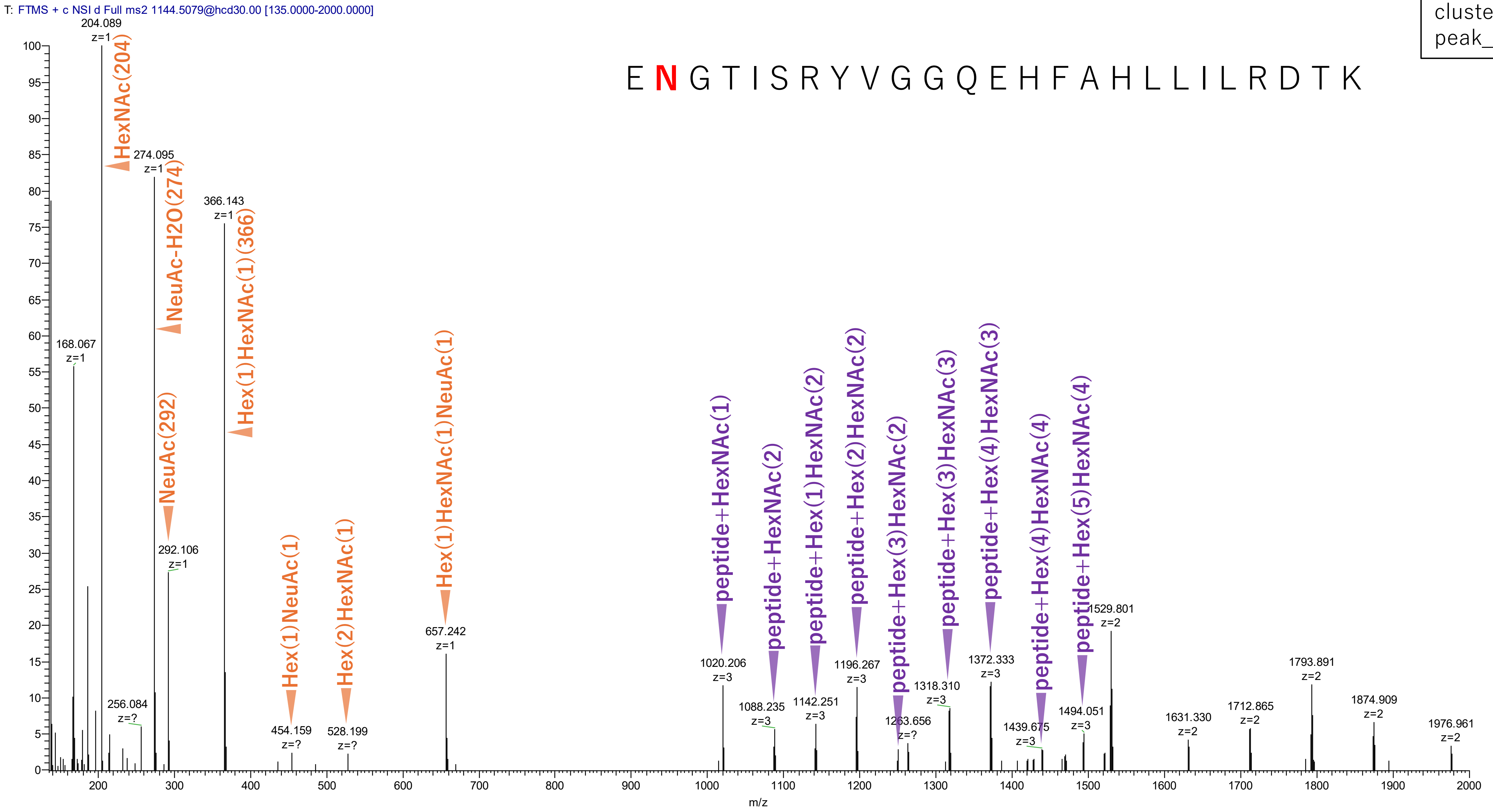


**Figure S2-99. MS2 spectra of glycopeptides assigned for hAGP.**

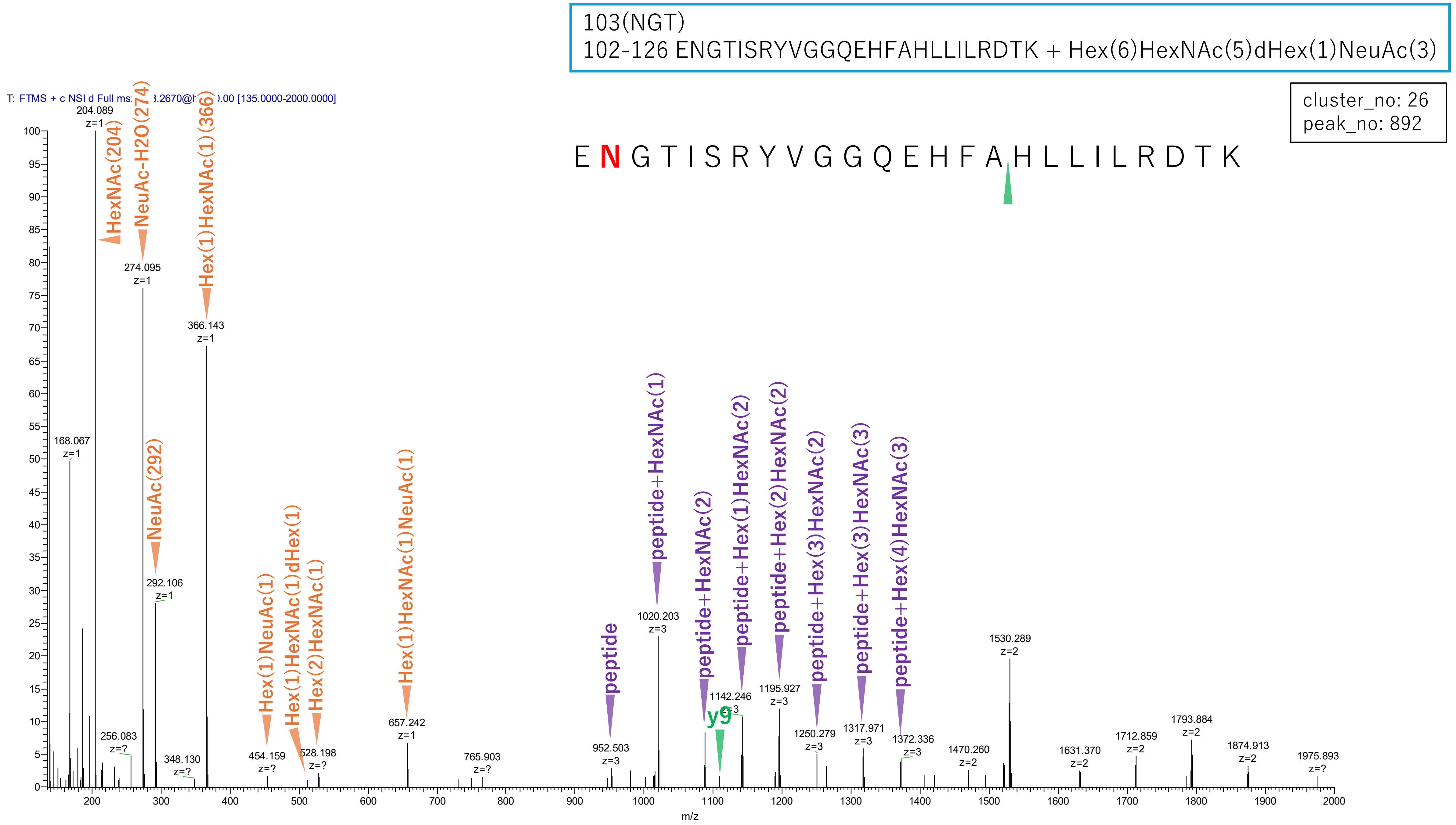


**Figure S2-100. MS2 spectra of glycopeptides assigned for hAGP.**

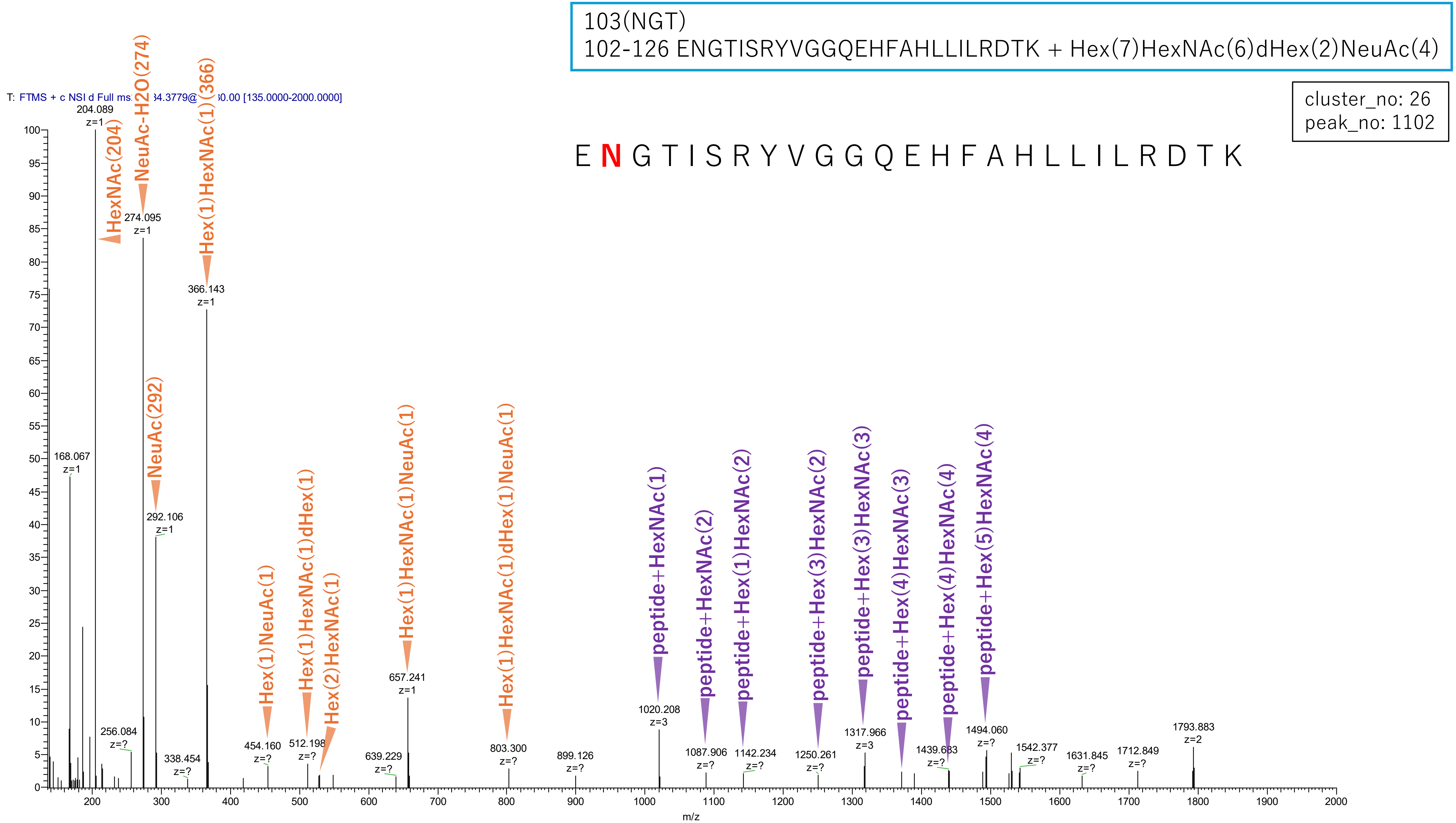


Figure S2-101. MS2 spectra of glycopeptides assigned for hAGP.

103(NGT)
102-126 ENGTISRYVGGQEHFAHLLILRDTK + Hex(7)HexNAc(6)dHex(1)NeuAc(3)

cluster_no: 26
peak_no: 1908

E N G T I S R Y V G G Q E H F A H L L I L R D T K

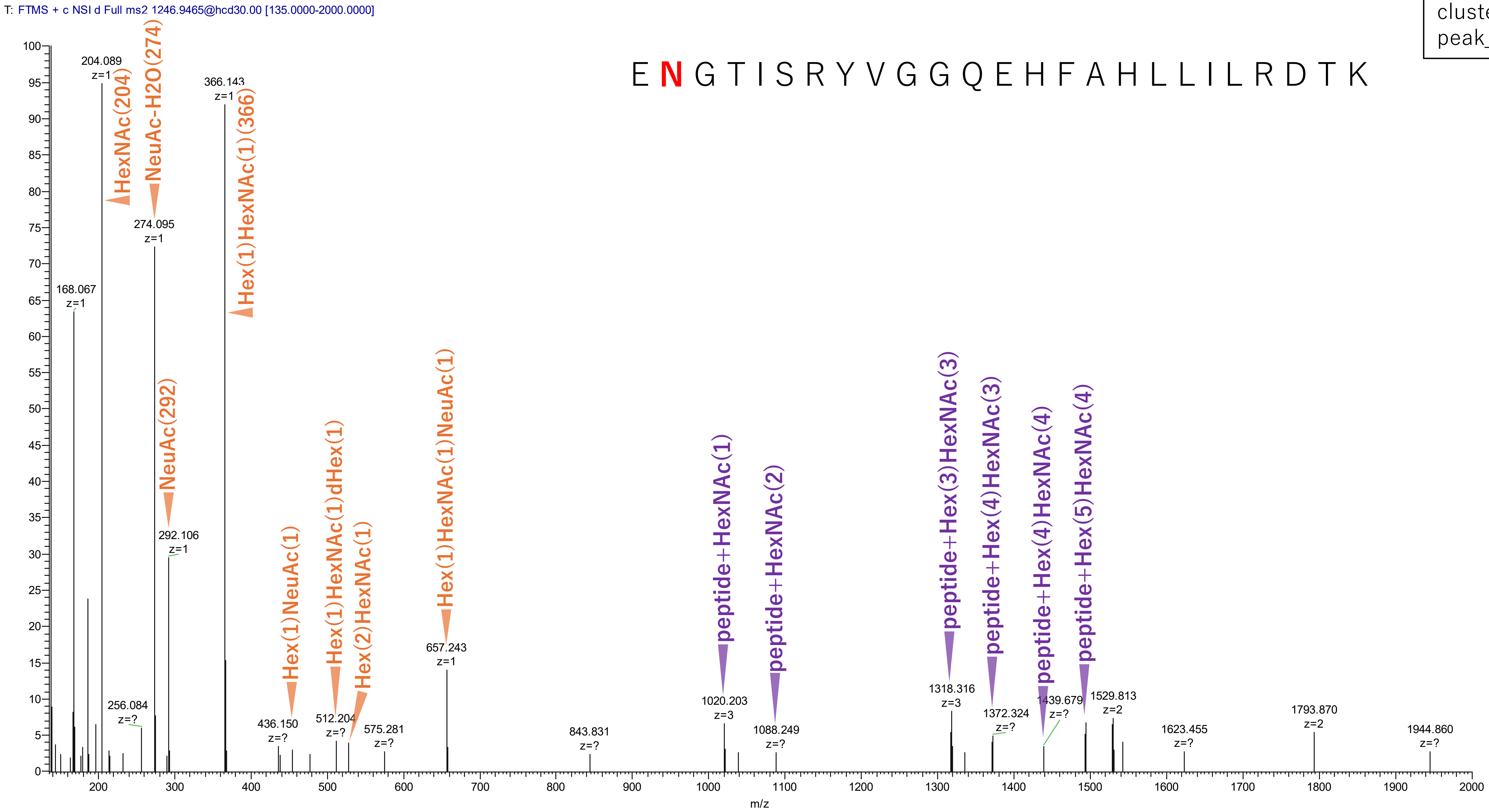


**Figure S2-102. MS2 spectra of glycopeptides assigned for hAGP.**

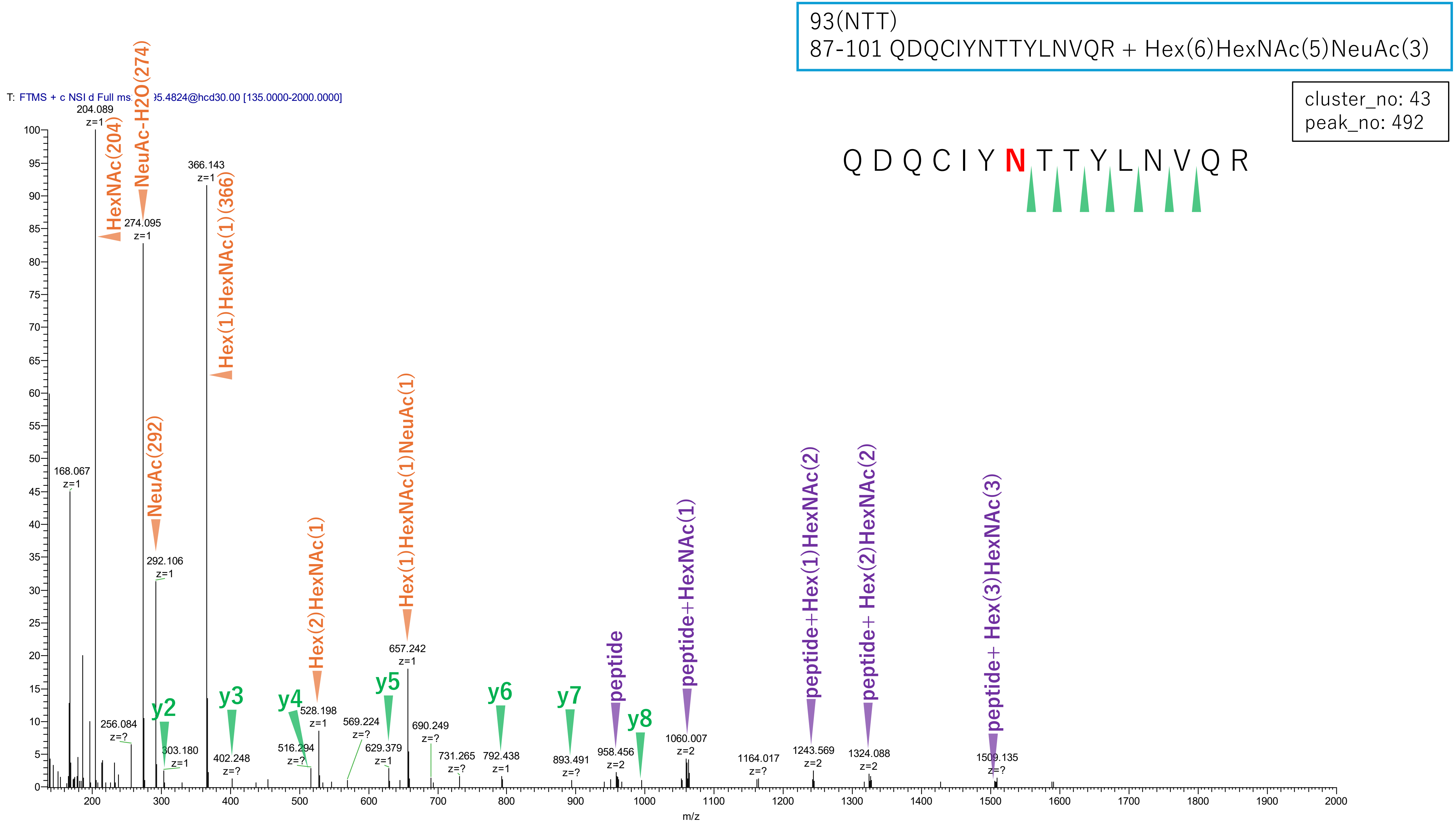


**Figure S2-103. MS2 spectra of glycopeptides assigned for hAGP.**

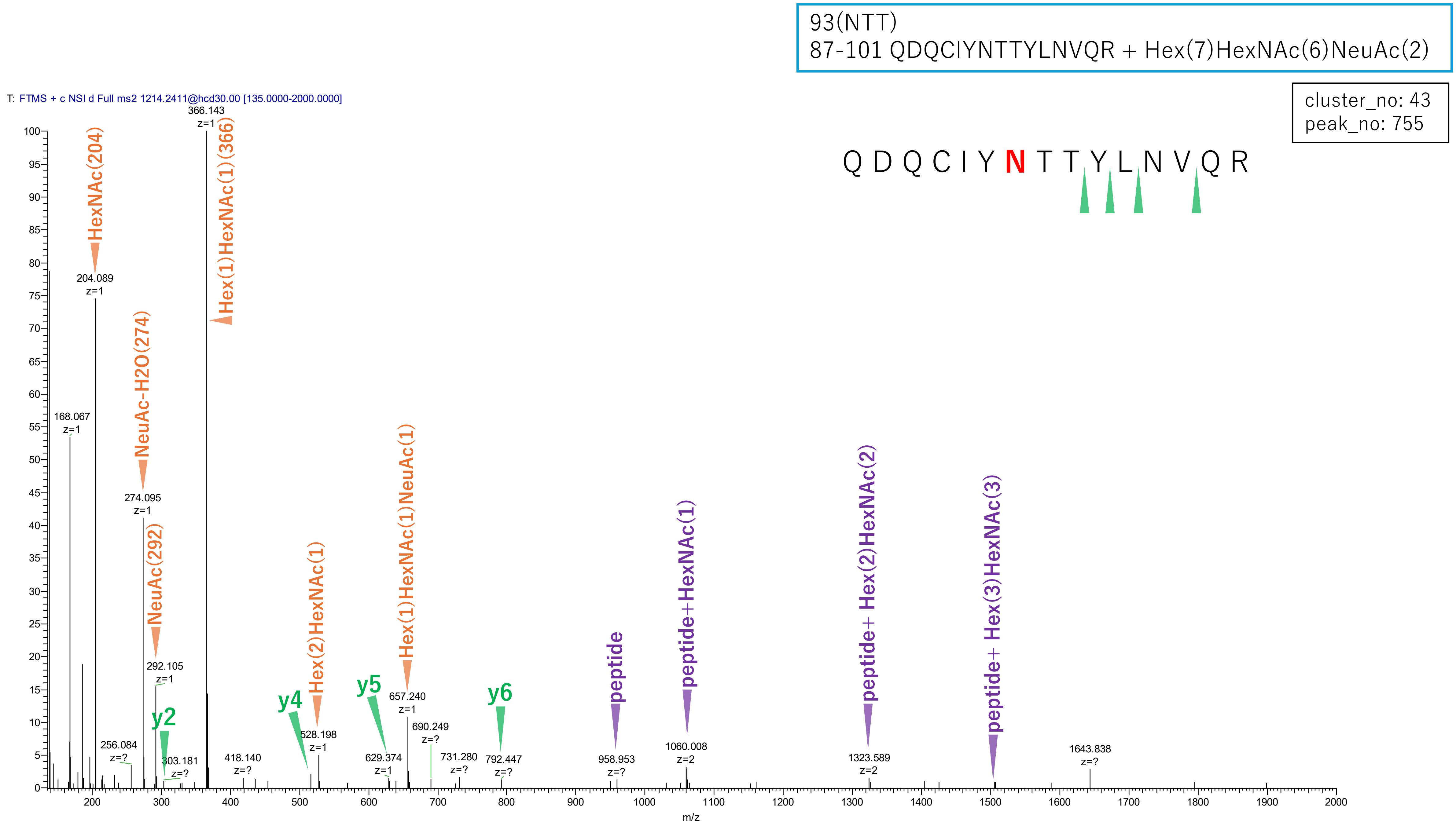


**Figure S2-104. MS2 spectra of glycopeptides assigned for hAGP.**

93(NTT)
87-101 QDQCIYNTTYLNVQR + Hex(6)HexNAc(5)dHex(1)NeuAc(3)

cluster_no: 43
peak_no: 969

Q D Q C I Y **N** T T Y L N V Q R

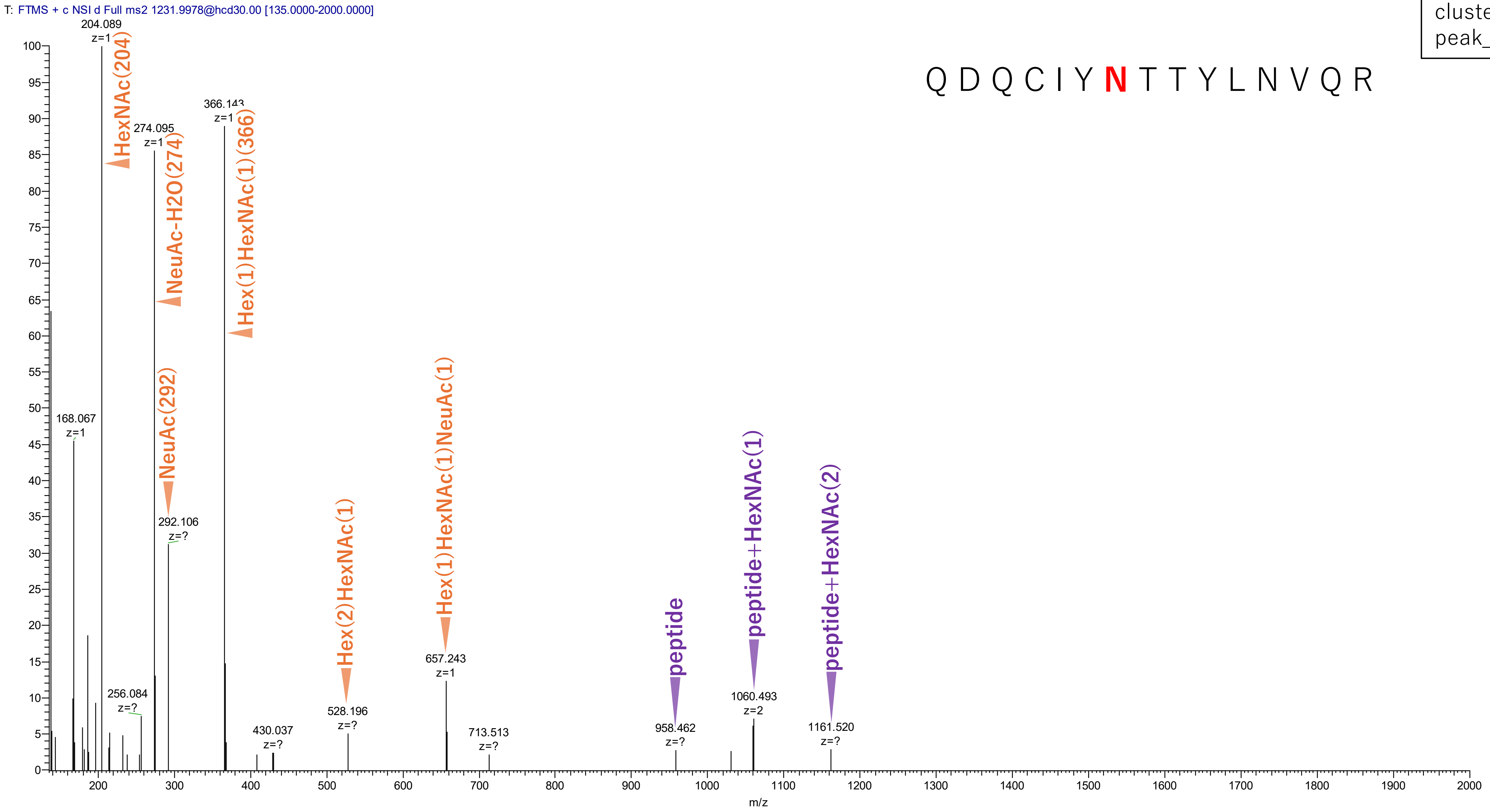


Figure S2-105. MS2 spectra of glycopeptides assigned for hAGP.

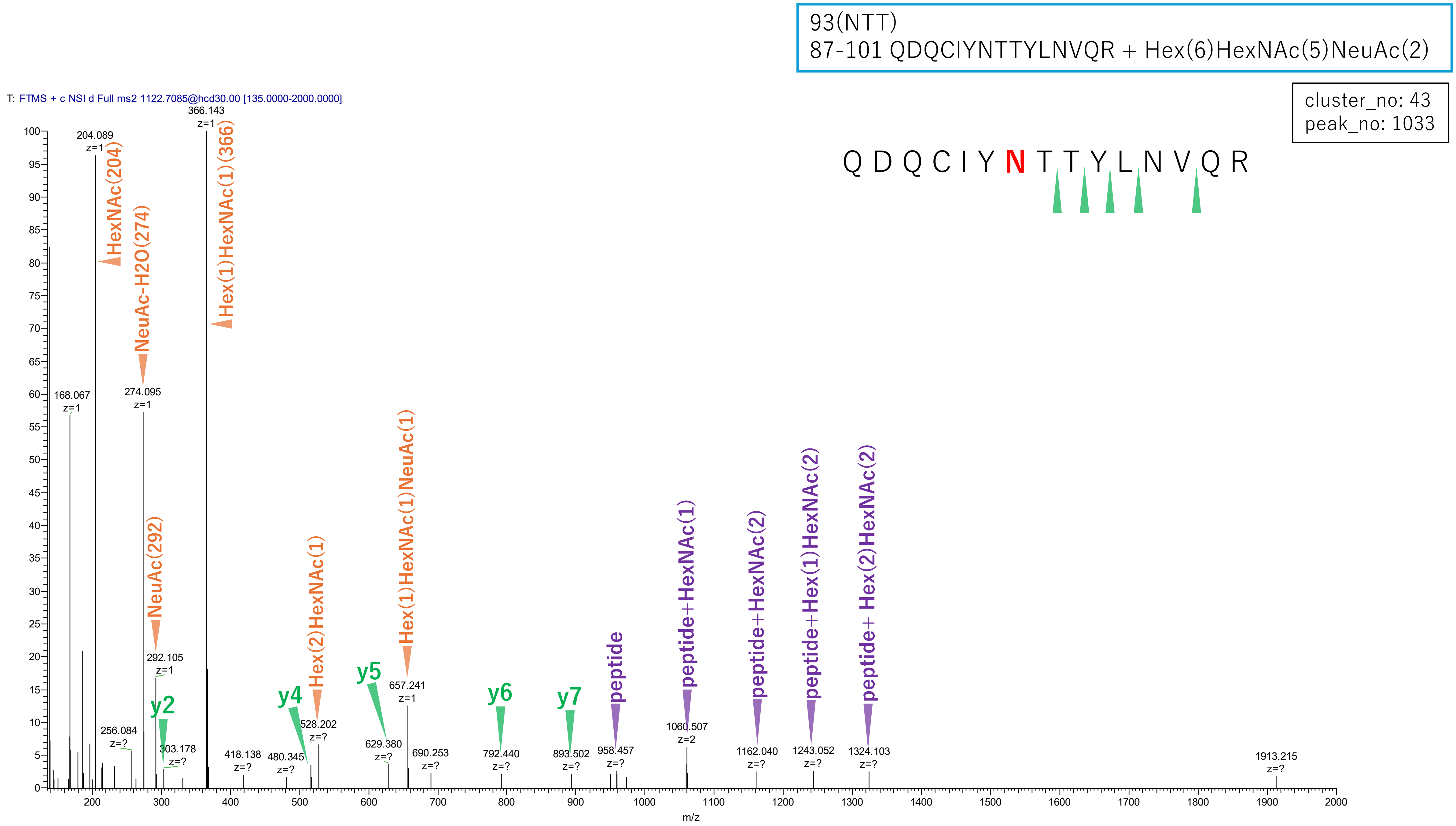


**Figure S2-106. MS2 spectra of glycopeptides assigned for hAGP.**

103(NGT)
102-123 ENGTISRYVGGQEHFAHLLILR + Hex(6)HexNAc(5)NeuAc(3)

cluster_no: 48
peak_no: 859

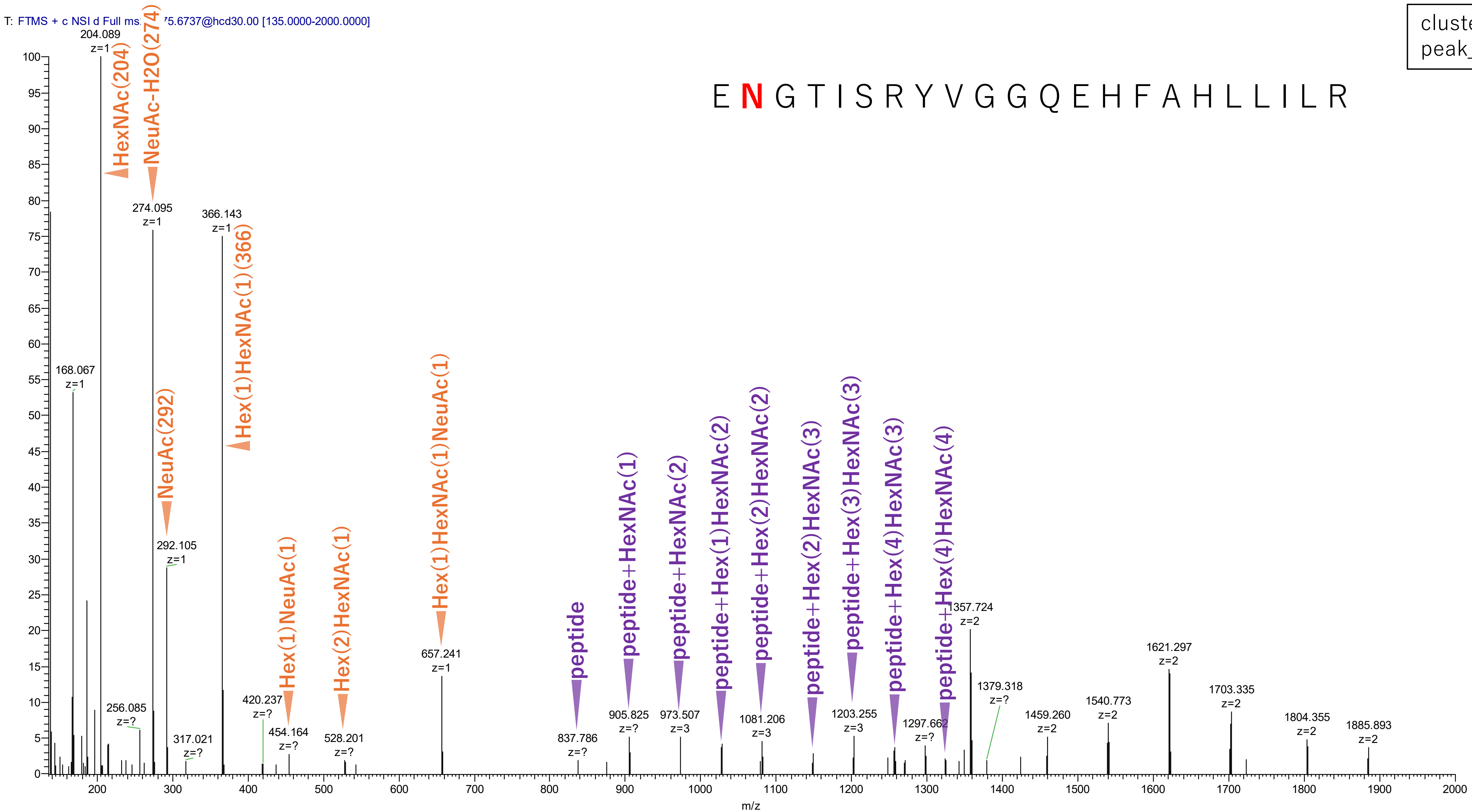


Figure S2-107. MS2 spectra of glycopeptides assigned for hAGP.

103(NGT)
102-123 ENGTISRYVGGQEHFAHLLILR + Hex(7)HexNAc(6)dHex(1)NeuAc(4)

cluster_no: 48
peak_no: 1023

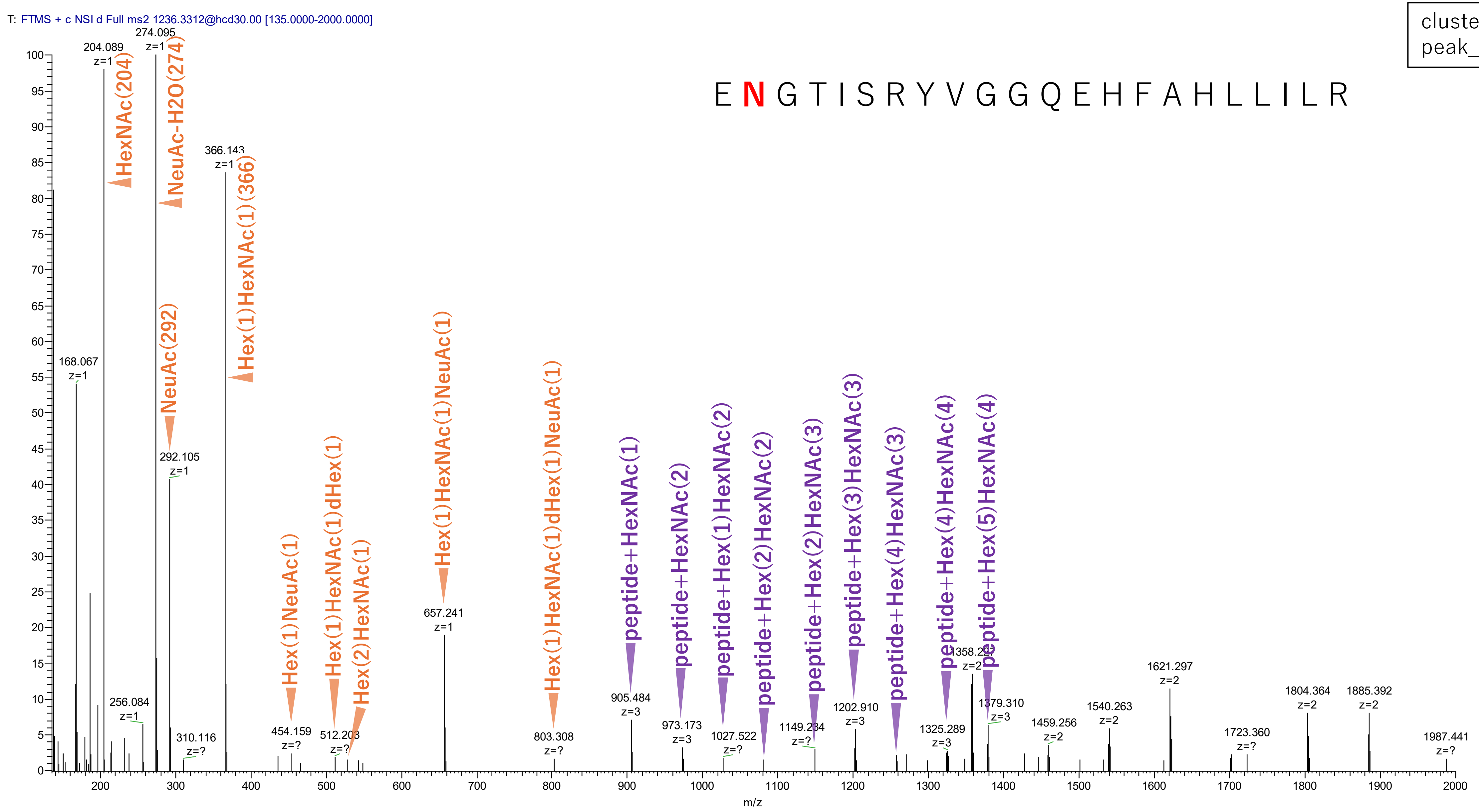


Figure S2-108. MS2 spectra of glycopeptides assigned for hAGP.

103(NGT)
102-123 ENGTISRYVGGQEHFAHLLILR + Hex(7)HexNAc(6)NeuAc(3)

cluster_no: 48
peak_no: 1564

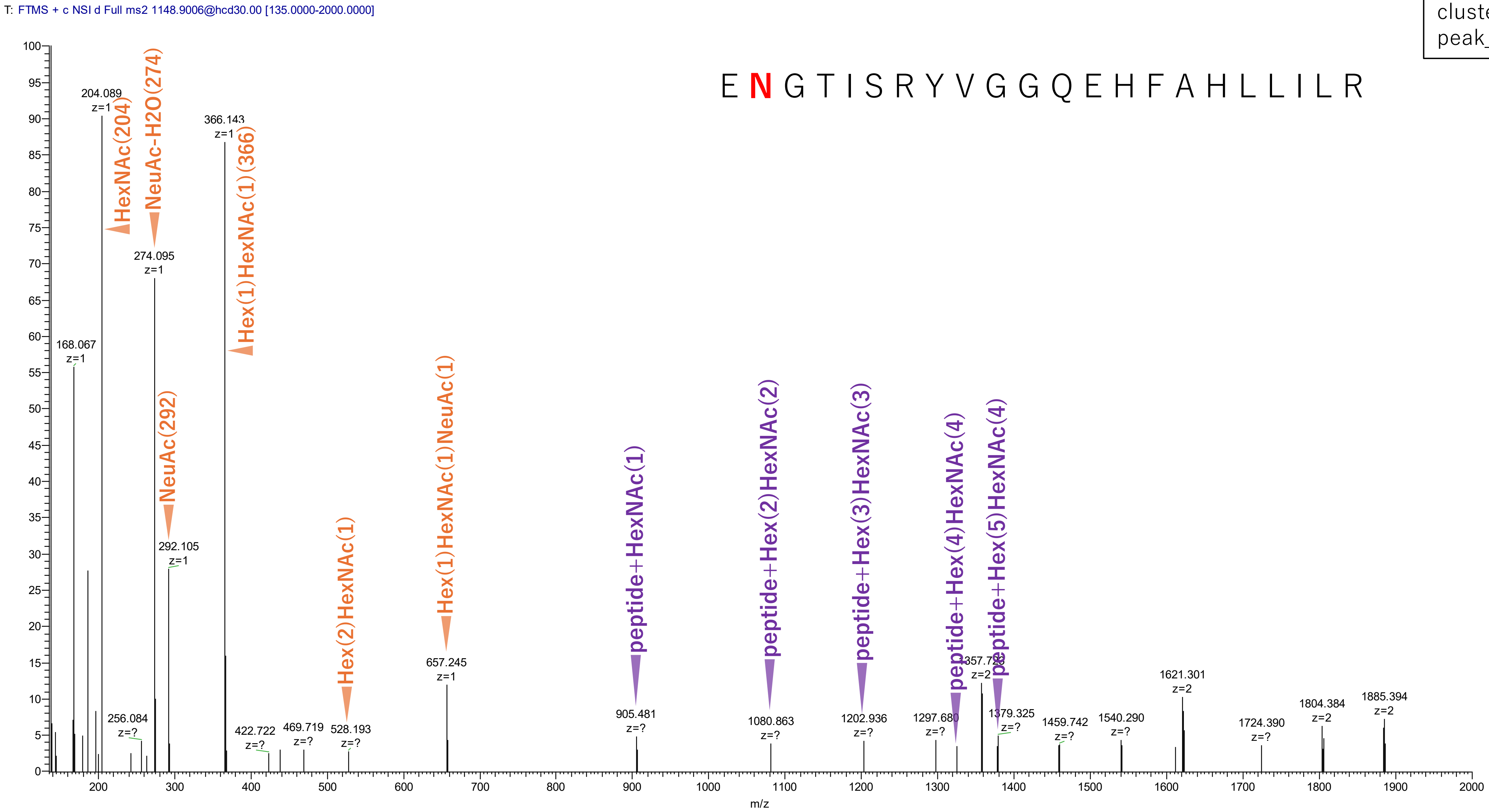


**Figure S2-109. MS2 spectra of glycopeptides assigned for hAGP.**

103(NGT)
102-123 ENGTISRYVGGQEHFAHLLILR + Hex(6)HexNAc(5)dHex(1)NeuAc(3)

cluster_no: 48
peak_no: 1613

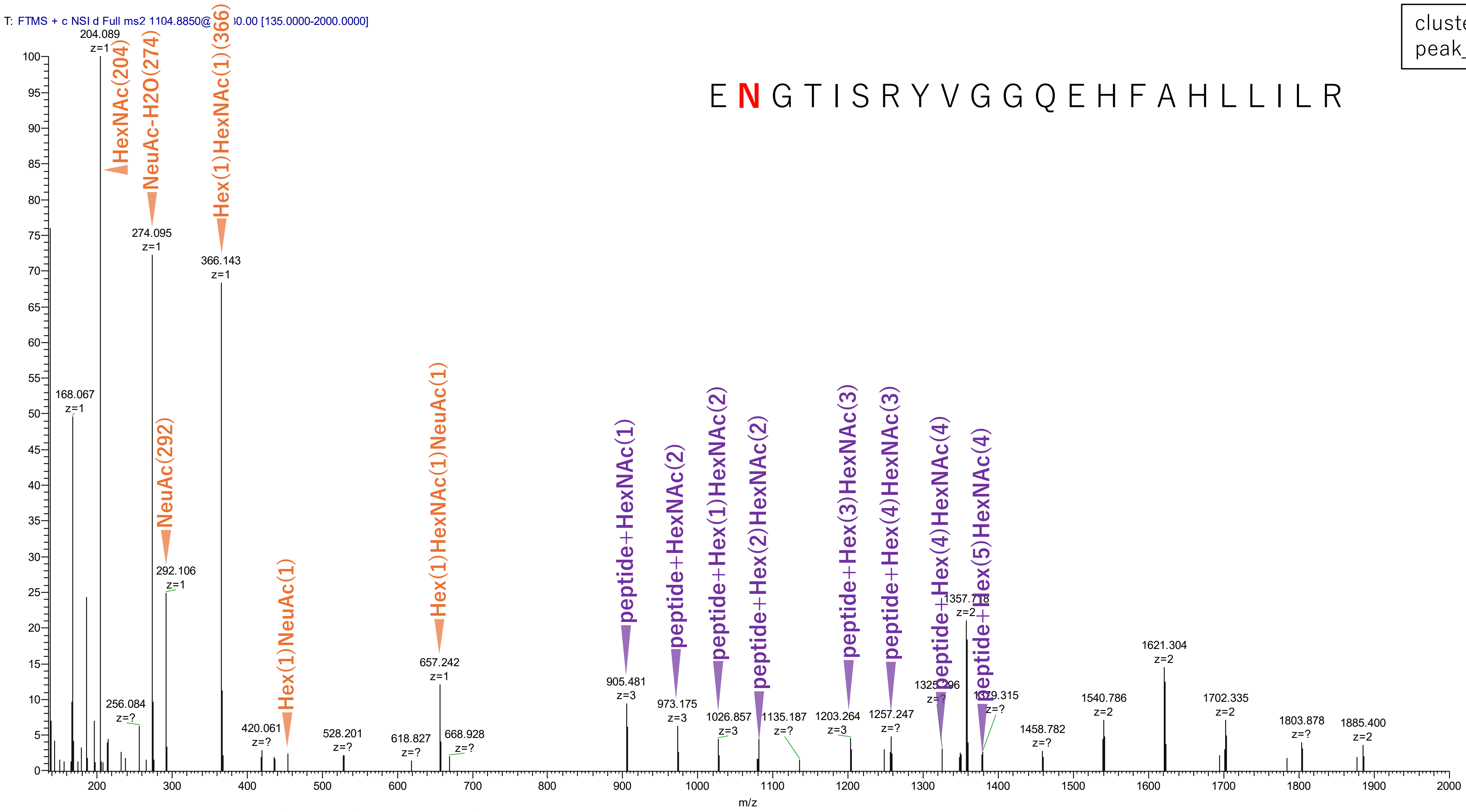


Figure S2-110. MS2 spectra of glycopeptides assigned for hAGP.
