## Supplementary material for "GRable version 1.0: A software tool for site-specific glycoform analysis with improved MS1-based glycopeptide detection with parallel clustering and confidence evaluation with MS2 information": Document S1

#### User Manual

Last update: April 10, 2024

### 1. Outline of GRable

#### 1.1. Introduction

GRable is a freely available online tool to find site-specific glycoforms of glycopeptides. This tool is unique in that it utilizes an MS1-based glycoproteomic method named “Glyco-RIDGE” (Glycan heterogeneity-based Relational Identification of Glycopeptide signals on Elution profile).

#### 1.2. Principle of the Glyco-RIDGE method

First, this method identifies glycopeptide signals based on the chromatographic properties of glycopeptides and mass differences due to the glycan heterogeneity. That is, glycopeptides having the same core peptide but different glycans elute within a narrow range of retention time (RT), so glycopeptide signals with a similar RT and mass differences corresponding to the masses of glycan units can be assigned as a cluster, without MS2 spectrum analyses. In parallel, core peptides present in the glycopeptide sample are identified by IGOT-LC/MS/MS. Then, considering that the mass value of glycopeptide is the sum of those of core peptide and glycan, the combination of peptide and glycan for each glycopeptide is searched from these three lists: the mass and RT lists of glycopeptides, peptides, and glycans).

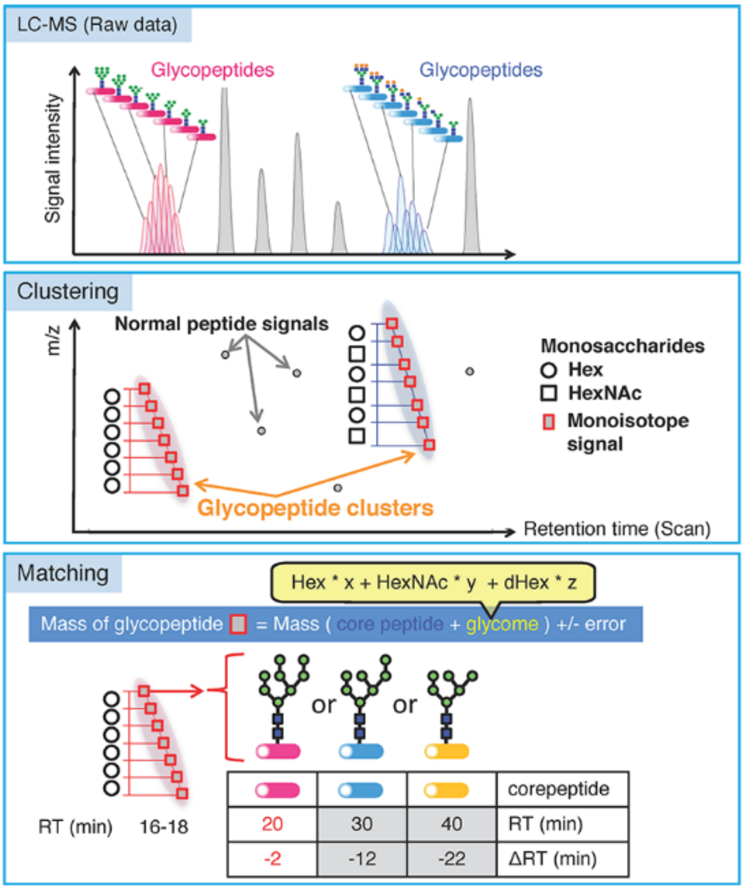

**Figure 1. Overview of the Glyco-RIDGE method**

(Narimatsu *et al. J Proteome Res,* 2018)

#### 1.3. Contact information

If you belong to an academic research institute, you can use the full version without functional limitations by concluding a joint research agreement between your organization and AIST.

Please contact us:.

### 2. System overview

#### 2.1. Overview of data processing

This software proceeds along seven steps. In the GRable Version 1.0, Step 2 (deconvolution) is currently unapplicable. Each step except for Step 2 is executed step-by-step after uploading required data and search parameter settings at a main window of the software. The results of Steps 3 to 5 can be visually confirmed at a viewer of the main interface. Detailed results of Steps 4 to 7 can be exported as an Excel file with each setting.

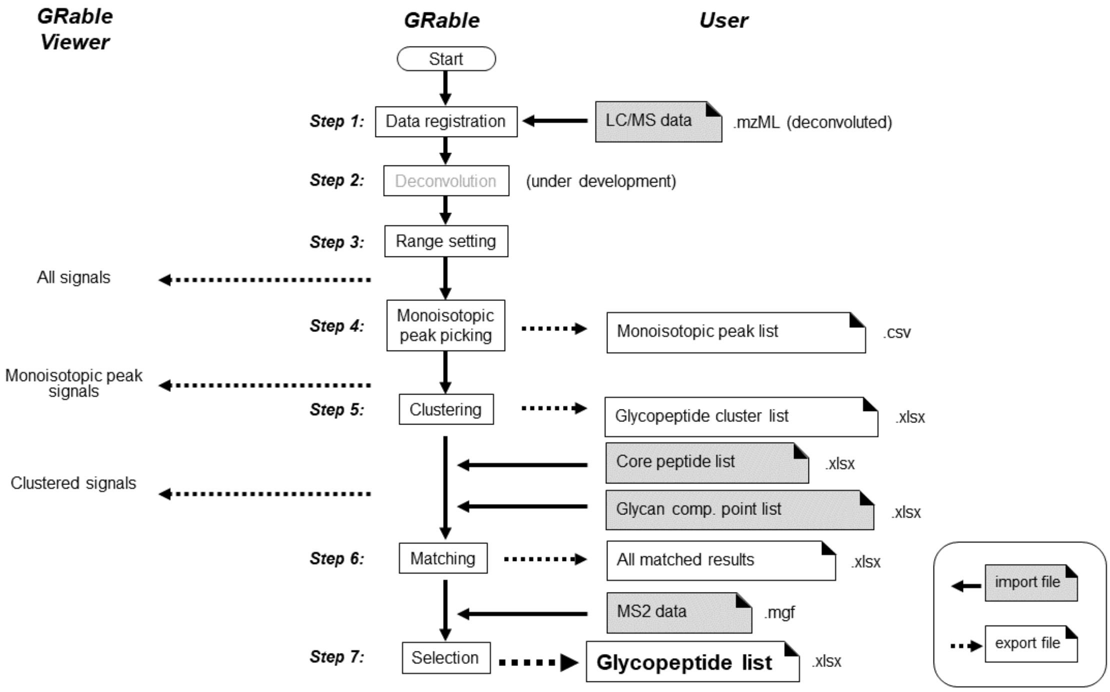

**Figure 2-1. Overview of data processing by GRable.**

#### 2.2. Requirements for input data

GRable requires the following 4 files [Note 1]. The details of input files are described below (Section 4.).

1) **LC/MS data of glycopeptides (.mzML):** The LC/MS data (.raw) should be deconvoluted and exported as an mzML format using Proteome Discoverer. ([Note 2, 3])
2) **Core peptide list (.xlsx):** This list contains information on existing core peptides identified by PNGase-mediated deglycosylation followed by LC/MS analysis. ([Note 4])
3) **Glycan point list (.xlsx):** This list is used for giving a point to individual member according to matched glycan compositions. Score of a cluster is sum of points obtained by each cluster member. The score will be used primarily to evaluate matching result for the cluster. It is possible to give high points to compositions with a high probability of existence, and to give penalties to compositions that are biosynthetically impossible.
4) **LC/MS/MS data of glycopeptides (.mgf):** Peak list of MS2 spectra extracted from the raw data of 1).

Notes

1. All data should be less than 300 MB each.
2. GRable is developed using Thermo Scientific data. Therefore, data from other manufacturers' equipment are currently not supported. Also, LC/MS using C18 column is assumed.
3. The analytical data require a mass accuracy better than 5 ppm. This is because GRable assigns glycopeptide signals using the accurate mass difference between glycopeptides with the same core

peptide and estimates glycan composition based on the mass difference between the core peptide and the glycopeptide. Low mass accuracy increases the possibility of incorrect matches.

4. This analysis should preferably be performed under the same LC conditions as the measurement of glycopeptide sample. In the case of highly purified protein, presumed peptide sequences can be included in the list.
5. By the GRable algorithm, multiple core peptides may match to single cluster. Then, it uses some surrounding information to select the most plausible match among them, e.g., sum of glycan points of cluster member, RT difference with core peptide, signal intensity of core, etc. In addition, if a cluster member has acquired MS2, the soft will use that information.
6. In the future, we plan to develop an updated version that can handle data from manufacturers other than Thermo Scientific, and that allows individual user management.

#### 2.3. GUI and functionality

Figure 2-2 shows the main window of the GRable Version 1.0 with the numbers of functions listed in Table 2-1. Note that functions that are present but not described in this manual are under construction and thus not guaranteed.

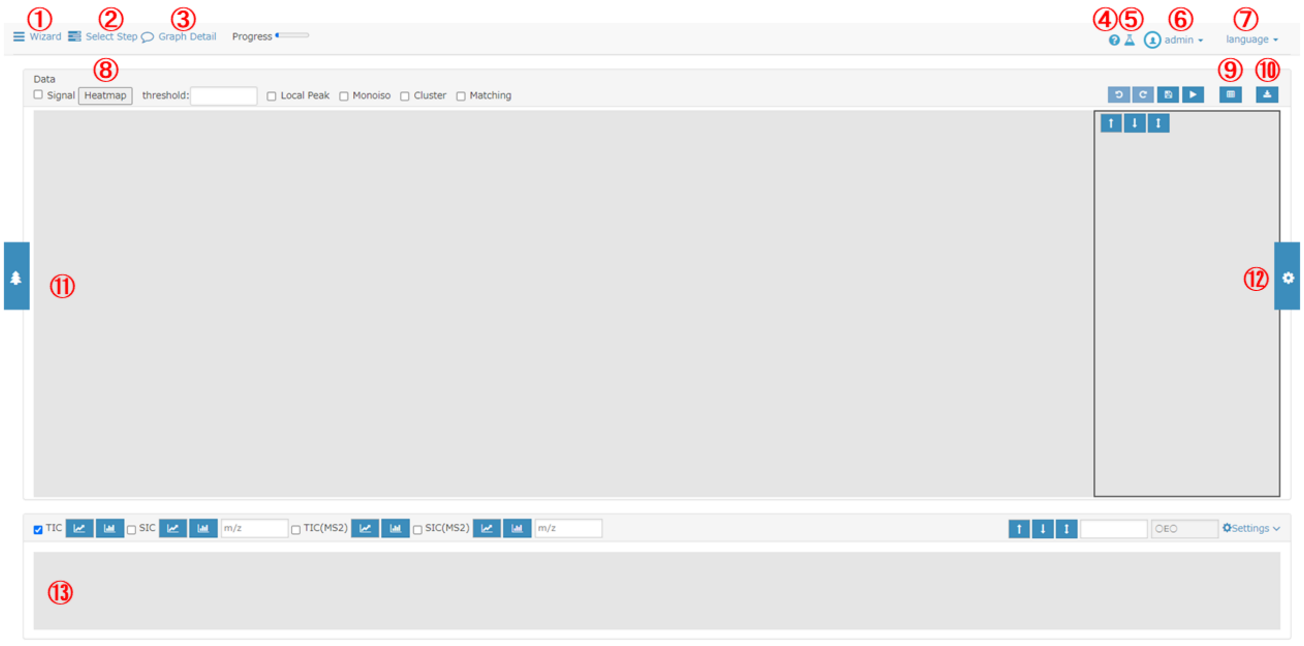

**Figure 2-2. Main window.**

**Table 2-1. List of functions.**

Function No. is the same shown in Figure 2-2.

| No. | Name of function | Description |
|---|---|---|
| 1 | Wizard | Select step you want to do in the module. |
| 2 | Select step | Select step you want to do in the dropdown list. |
| 3 | Graph Detail | Displays setting parameters of the displayed graph. |
| 4 | System information | Displays system update information. |
| 5 | Monoisotopic Mass Table | Displays a Monoisotopic Mass Table. |

| No. | Name of function | Description |
|---|---|---|
| 6 | User/project management | Manages users and projects and log out. |
| 7 | Language | Selects display language (Japanese or English). |
| 8 | Heatmap | Displays signal intensity as a heatmap. |
| 9 | Option | Displays 3D display or monoisotopic peak list. |
| 10 | Download | Downloads graph image data, clustering analysis results, and matching results. |
| 11 | Tree | Displays data and analysis results in a tree format and performs display, analysis, and reanalysis. |
| 12 | Settings | Displays the setting panel for heatmap operations (enlargement/reduction, etc.). |
| 13 | Chromatograms | Displays the Chromatograms graph. |

### 3. Operation

#### 3.1. Login

To log in to the system, you need to enter a login ID and password on the login window (Figure 3-1). When both are appropriate, you can low in the software.

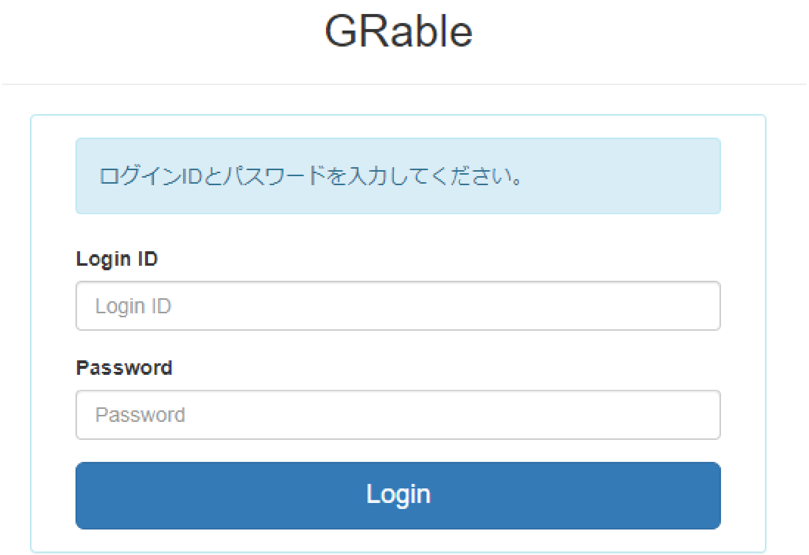

**Figure 3-1.　Login window.**

#### 3.2. Language selection

The display can be switched between in Japanese and English using the language selection menu (Figure 2-2⑦) on the upper right of the main window.

#### 3.3. Data registration (Step 1)

##### 3.3.1. Functional overview

GRable allows the registration of LC/MS data of glycopeptides in mzML format, which ensures great versatility and ease of data processing. This software is also designed to deal with charge-deconvoluted and centroid spectra of each scan. This software is tested and optimized using the mzML data prepared with

Proteome Discoverer (version 2.4 or later; Thermo Fisher Scientific), in which deconvolution is performed with a workflow consisting of Xtract. To do the Glyco-RIDGE analysis, high accuracy of mass measurement is strongly required, e.g., mass resolution of MS1 by orbitrap analyzer is 120,000 or 240,000 (at m/z 200) and a lock mass at 445.12003 is necessary.

###### 3.3.2. Operation procedure

To register data for analysis, press the wizard button (Figure ①) on the main window to display the data registration module (Figure 3-2).

① Select the type of data to be registered. Select “LC/MS” for uploading mzML format files after deconvolution.
② Enter the analysis data set name.
③ Select data. After data uploading, the check will be added.
④ Press the Register button.
⑤ When "Registration completed" appears, the process is finished. Proceed the next step.
⑥ If you want to cancel this step in progress, press the "Cancel" button.

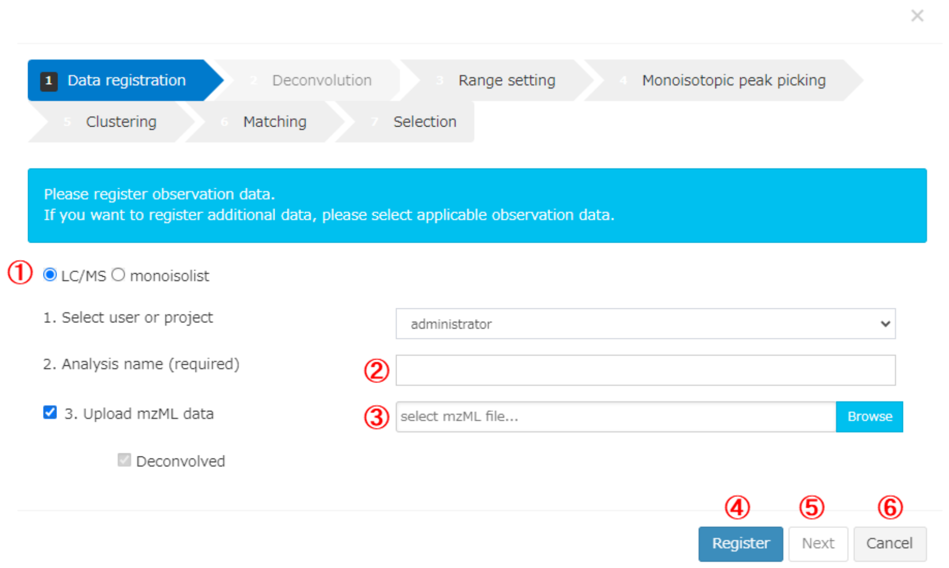

**Figure 3-2. Data registration module.**

##### 3.4. Range setting (Step 3)

###### 3.4.1. Functional overview

This step (Figure 3-3) is designed to set the RT range, mass range, and minimum signal intensity (threshold) over which the analysis will be performed. Analysis for a data of long gradient elution and broad mass ranges are time consuming. The signals selected in this step (all signals) can be seen as a heatmap in the viewer main window. The resulting peak list is exported as a CSV (comma-separated values) file because of the huge size of the data. (In the subsequent steps, the setting parameters and the results were exported as an Excel (.xlsx) file.)

##### 3.4.2. Operation procedure

To perform range setting, press the wizard button (Figure 2-2①) from the main module to display the range setting module (Figure 3-3).

① Enter an analysis name.
② Set parameters (Table 3-1). Note that "Load settings" is unapplicable in this version.
③ Select whether the setting is saved.
④ Click the "Run" button to start analysis.
⑤ When "Range setting completed" is displayed, the process is finished. Please proceed the next step.
⑥ If you want to cancel this step in progress, press the "Cancel" button.

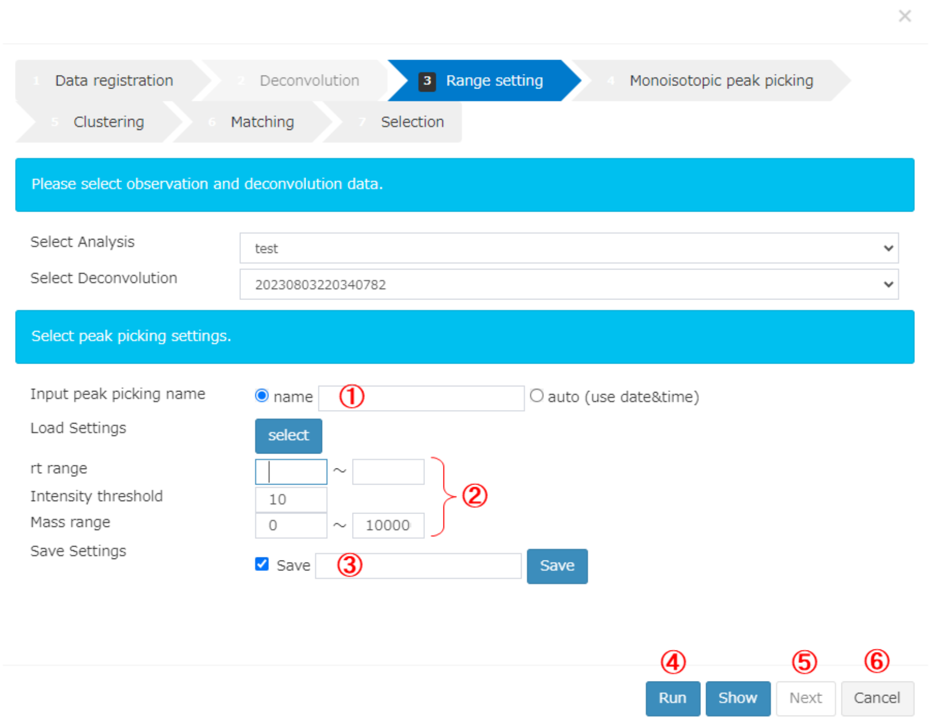

**Figure 3-3. Range setting module.**

**Table 3-1. Details of range settings.**

| No. | Parameter | Description |
|---|---|---|
| 1 | rt range | Set RT range (min) of data used for analysis |
| 2 | Intensity threshold | Set signal intensity threshold of data used for analysis |
| 3 | Mass Range | Set mass range (upper and lower limits) of data used for analysis |

#### 3.5. Monoisotopic peak picking (Step 4)

##### 3.5.1. Functional overview

This step (Figure 3-4) is intended to find and group signals of the identical ion based on three parameters of time (scan), mass (MH+), and intensity, and to obtain the monoisotopic mass of each signal group at the peak time, using our unique algorithms. At the previous step, signals without isotope signals were removed as noise, so all signals have one isotope signal at least. First, all signals within the analysis range are surveyed to find local peak signal. If a peak signal is found, the same signals within the error tolerance (set at 5 ppm typically) are repeatedly searched from the neighboring scans (toward both sides of RT) and grouped together. The same searches are performed for other isotope signals of the peak signal and obtain a single signal group. The minimum size of the signal group is set to 3 scans × 4 isotope signals as default. Three scan spectra centered on each group peak scan are then accumulated and the resulting spectra are analyzed by the shape analysis module. In this module, monoisotopic signals are searched from the highest signal toward lower mass. The monoisotopic signal threshold (lowest intensity) to the highest signal is calculated from the oligomer of carbamidomethylcysteine with the lowest number of carbons per 100 masses (=3.1, mean = 4.3, maximum = 6.1 (Phe)). Default values are shown in a setting module (Figure 3-5). If the relative intensity of a candidate monoisotopic signal is less than this value, the signal is not considered as monoisotopic and the search ends. After this search, a monoisotopic peak list of all ions is created (which can be exported as a table consisting of peak RTs, monoisotopic masses, and peak intensities). The picked monoisotopic peak signal can be seen in the viewer overlaid on all signals.

##### 3.5.2. Operation procedure

To perform monoisotopic peak picking, press the wizard button (Figure 2-2①) from the main window to display the monoisotopic peak picking module (Figure 3-4). Recommended parameters are set as a default.

① Select the “Monoisotopic peak picking” tab.
② Enter an analysis name.
③ Set parameters (Table 3-2). Note that “Load settings” is unapplicable in this version.
④ Select whether the setting is saved.
⑤ Click the “Start” button to start analysis. When “Analysis complete” is displayed, the process is finished.
⑥ If you want to cancel this step in progress, press the "Cancel" button.

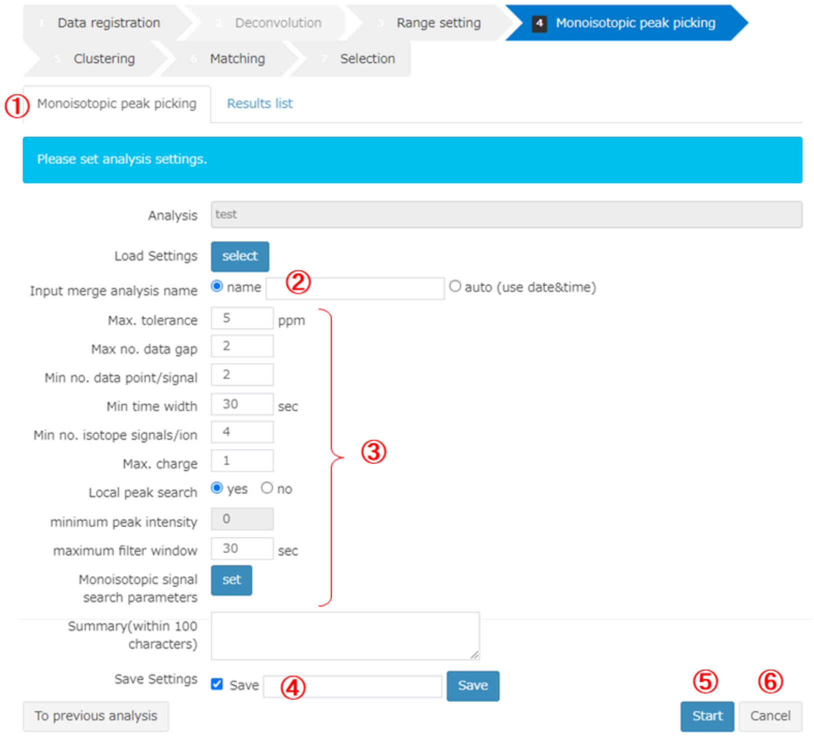

**Figure 3-4. Monoisotopic peak picking module.**

**Table 3-2. Details of monoisotopic peak picking settings.**

| No. | Parameter | Description |
|---|---|---|
| 1 | Max. tolerance | Sets the maximum mass tolerance among a series of scans for treating signals as ones derived from the identical ions. |
| 2 | Max no. data gap | Sets the number of allowable lack of scans for treating signals within a single ion group. |
| 3 | Min no. data point/signal | Sets the minimum number of scans of a single group. |
| 4 | Min time width | It is unnecessary to set this parameter because it is not utilized for data processing in this version. |
| 5 | Min no. isotope signals/ion | Sets the minimum number of isotopes. |
| 6 | Max. charge | This parameter should be set as 1, because only deconvoluted data is used for analysis. |
| 7 | Local peak search | If “yes” is selected, local peak detection is performed using the Maximum filter and uses it as the starting point for the monoisotopic peak search. If “NO” is selected, all signals are used as the starting point for monoisotopic peak search in descending order of intensity. Note that it is recommended to select “yes”, because this process takes much time without the Maximum filter. |
| 8 | minimum peak intensity | The minimum intensity of signal as a starting point for monoisotopic peak search. |
| 9 | maximum filter window | Specifies the filter size (time width) of the Maximum filter for local peak search. |
| 10 | Monoisotopic signal search parameters | Define the parameters for determination of the monoisotopic peak position based on the relative intensities of isotopes in a dialog box (Figure 3-5). |

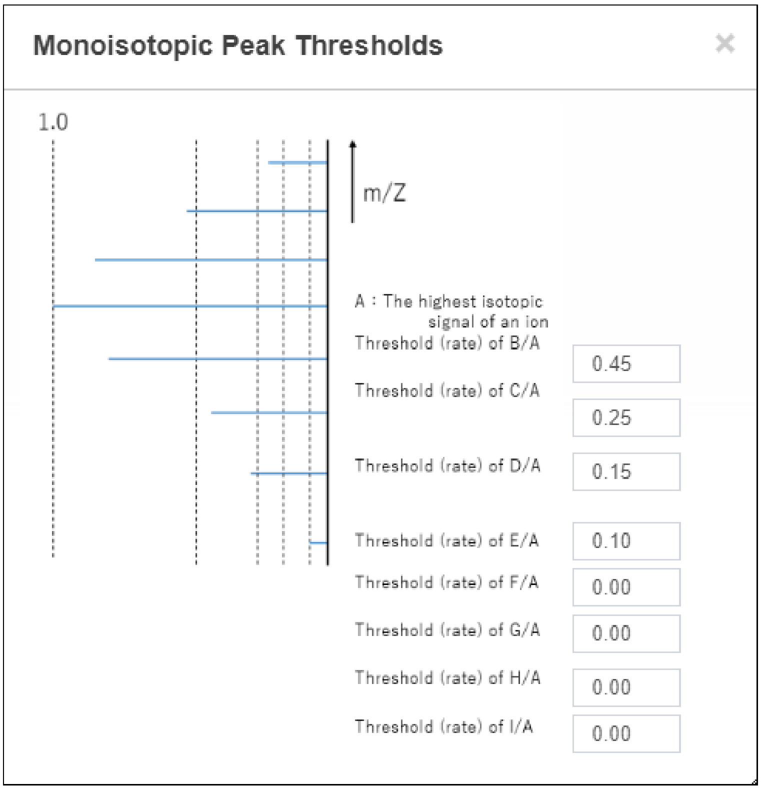

**Figure 3-5. Setting window for monoisotopic signal search parameters.**

#### 3.6. Clustering (Step 5)

##### 3.6.1. Functional overview

In this step (Figure 3-6), a series of signals of glycopeptide group are found as a cluster based on the elution behavior and mass difference of its members. By setting maximum RT difference and maximum error of mass difference for glycan units (such as Hex, HexNAc, and dHex), a pair of putative glycopeptide group signals is searched for all monoisotopic peaks. If a single signal (node) has multiple relations (edges), the pairs are combined to form a cluster. By repeating similar search, a group with at least a user-defined number of members is considered as a single glycopeptide cluster. The minimum number of members in a cluster is set to 4 as a default.

##### 3.6.2. Operation procedure

To perform clustering, press the wizard button (Figure 2-2①) from the main window to display the clustering module (Figure 3-6).

① Select the “Clustering” tab.
② Enter an analysis name.
③ Set parameters (Table 3-3). Note that “Load settings” is unapplicable in this version.
④ Select whether the setting is saved.
⑤ Click the “Start” button to start analysis. When “Analysis complete” is displayed, the process is finished.
⑥ If you want to cancel this step in progress, press the "Cancel" button.

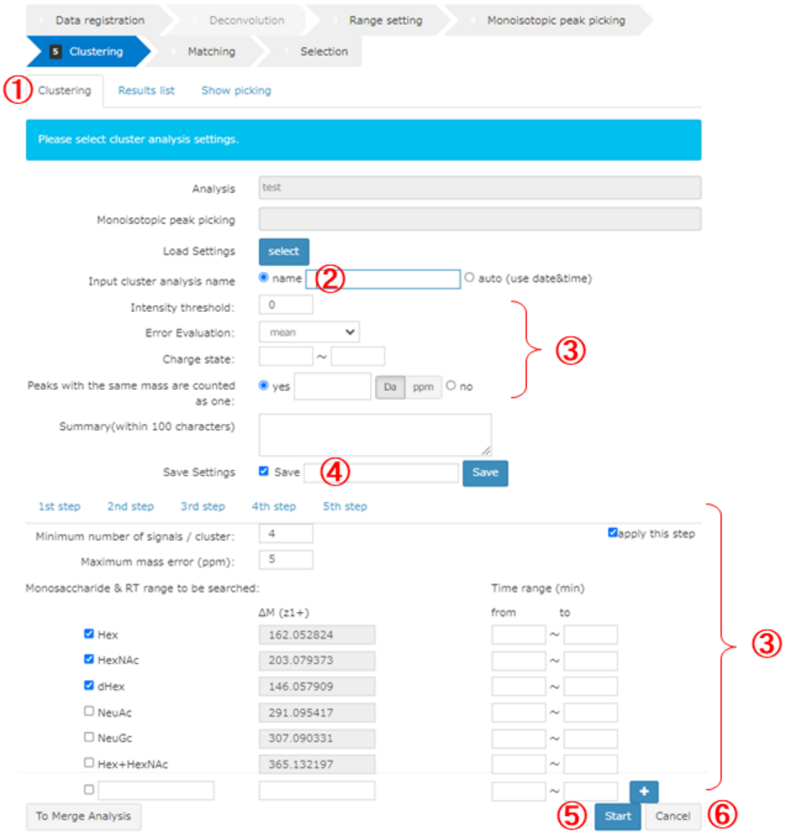

**Figure 3-6. Clustering module.**

**Table 3-3. Details of clustering settings.**

| No. | Parameter | Description |
|---|---|---|
| 1 | Intensity threshold | Sets the minimum intensity threshold. |
| 2 | Error Evaluation | Sets the error evaluation method ("mean" or "sum SQ of SQ"). The default setting is "mean", which is a stricter setting. GRable calculates the difference between M1 and M2 (masses of two monoisotopic signals). To calculate the difference in ppm, the difference (ΔΔM obs-calc) between the observed mass difference (ΔM obs=M2-M1) and calculated mass value of any glycan unit (e.g., M(Hex); set by user as shown below) is divided by either value of:<br>- mean: (M1+M2)/2<br>- sum SQ of SQ: $\sqrt{(M1)^2 + (M2)^2}$<br>If the difference (ppm) is smaller than the maximum mass tolerance (set by user as shown below), the two monoisotopic signals are considered in the relationship with a difference of the glycan unit. |
| 3 | Charge state | Note that this parameter should remain blank, because only deconvoluted data is used for analysis. |
| 4 | Peaks with the same mass are counted as one | Selects "yes" or "no". If "yes" is selected (by default), peaks with the same mass, but divided to multiple peaks, are considered as single peak (composition) when the number of members of a cluster is evaluated. Note that peaks with the same mass are highlighted in yellow in an exported sheet. |
| 5 | 1st step (2nd step…) | Sets the parameters for each step of parallel clustering.<br>By clicking, you can switch tabs and enter settings for each step. |
| 6 | apply this step | Checks the checkbox for each step whether the step is included in parallel clustering. Users can apply up to five steps in one analysis. |
| 7 | Minimum number of signals / cluster | Sets the minimum number of signals for one cluster. |
| 8 | Maximum mass error(ppm) | Sets the maximum mass tolerance. |
| 9 | Monosaccharide & RT range to be searched | Specifies types of glycan units (mono- and oligo-saccharides) along with their masses and RT ranges to be considered. Hex, HexNAc, dHex, NeuAc, NeuGc, Hex+HexNAc can be entered by default, and the number of free entry fields can be increased up to 10. |

#### 3.7. Matching (Step 6)

##### 3.7.1. Functional overview

In this step (Figure 3-7), GRable searches combination of core peptide and glycan composition that match with the mass of putative glycopeptide within the allowed mass error (user setting), according to the following equation:

Observed M(glycopeptide) = calculated M(core peptide identified) + M(Hex)*i + M(HexNAc)*j + M(dHex)*k + M(NeuAc)*l (where M is a mass value, and i, j, k, and l are integers)

To do this, a list of core peptide candidates is required. The list can be prepared by the PNGase-mediated deglycosylation followed by LC/MS analysis of the same glycopeptide sample. Identification of deglycosylated peptides by LC/MS is one to two orders of magnitude more sensitive than glycopeptide identification, allowing core peptide listing using only 5-10 % of the sample used for glycopeptide analysis. As described in Selection step below, RT differences between glycopeptides and deglycopeptides are one of clue for selecting correct match, so LC/MS data is recommended to be obtained on the same day and sequential runs of analysis. In addition, a glycan point list is also needed. Glycan compositions considered for matching are given as the number range of each glycan units (user setting). As default, the ranges of Hex, HexNAc, dHex, and NeuAc are set to 0-10, 1-10, 0-4, and 0-4, respectively. Then, using the set of 3 masses of glycopeptides, core peptides, and glycans, matched combination are searched according to the equation 1. For each glycopeptide, point is given according to the glycan composition matched. The points are set in the glycan point list (given by user). Glycan points are classified into three kinds; 1 = compositions produced by common/familiar biological glycan processing, 0 = possible but not common, and -1 = unusual from common biological pathway. Users can set this allocation of points, e.g., higher points are assigned to the compositions which were observed in actual glycome analysis. All matched results can be obtained as an Excel file.

##### 3.7.2. Operation procedure

To perform matching, press the wizard button (Figure 2-2①) from the main window to display the matching module (Figure 3-7).

① Select the “Matching” tab.
② Enter an analysis name.
③ Set parameters (Table 3-4). Note that “Load settings” is unapplicable in this version.
④ Select whether the setting is saved.
⑤ Click the “Start” button to start analysis. When “Analysis complete” is displayed, the process is finished.
⑥ If you want to cancel this step in progress, press the "Cancel" button.

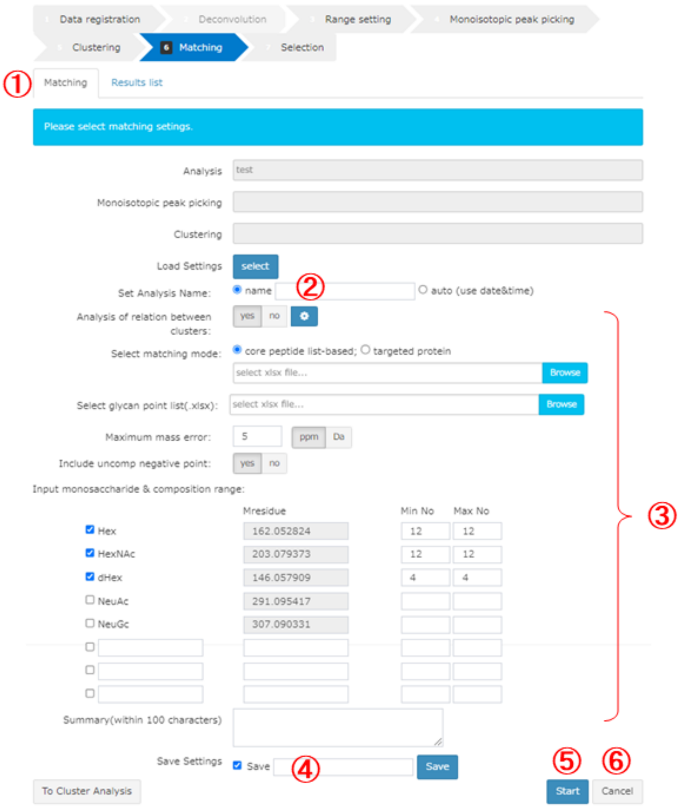

**Figure 3-7.　Matching module.**

**Table 3-4.　Details of matching settings.**

| No. | Parameter | Description |
|---|---|---|
| 1 | Analysis of relation between clusters | Selects whether inter-cluster analysis is executed. If “yes” is selected, enter the settings by clicking on the button on the right side to display the setting dialog (Figure 3-8). In the search difference setting, specify types of relations to be considered along with mass tolerance (Da). “Delta RT (min)” is a RT range for evaluating other cluster members. “Minimum rate of related members” is a threshold of the parentage of members detected in this search; when the members over the threshold are detected, reference and target clusters are treated as “related” and the target cluster is indicated in a export file. |
| 2 | Select matching mode | Select “core peptide list-based” mode and upload a core peptide list (in a fixed format). Note that “target protein” mode is not available in this version. |
| 3 | Select glycan point list | Uploads a glycan point list. |
| 4 | Maximum mass error | Sets the maximum mass tolerance (ppm or Da). |
| 5 | Include uncomp negative point | Selects whether negative points are given for unusual glycan compositions defined in a glycan point list (in a fixed format). |
| 6 | Input monosaccharide & composition range | Specifies types of glycan units (mono- or oligo-saccharide) along with their masses and minimum/maximum numbers to be considered. Hex, HexNAc, dHex, NeuAc, and NeuGc can be entered by default, and there are also three free entry fields. Be sure that the glycan units are totally identical to the ones written in a glycan point list; unless an alert will appear. |

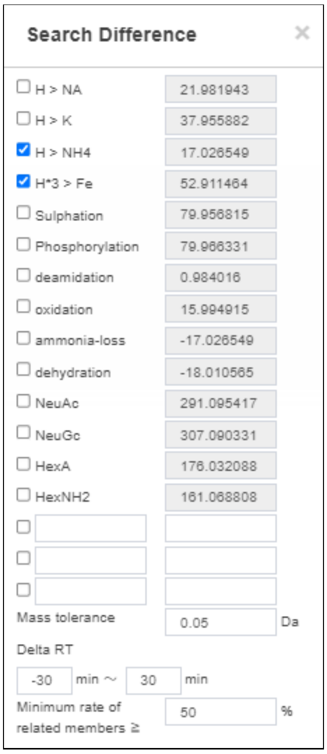

**Figure 3-8. Setting dialog of inter-cluster analysis.**

#### 3.8. Selection (Step 7)

##### 3.8.1. Functional overview

In the matching step, multiple combinations are often suggested for one glycopeptide cluster. To select the most plausible combination among the candidates, GRable collects the following information from the results and additional MS2 information to evaluate their reliability at the cluster level as well as the single glycopeptide level.

Information

1) **Core peptides abundance:** The number of core peptides identified from a small aliquot of glycopeptide sample by the IGOT-LC/MS/MS method greatly exceeds the number of clusters detected by the Glyco-RIDGE method. This suggests that detected glycopeptide clusters are likely to have core peptides with high abundance or high ionization efficiency among the peptides contained in the sample. Accordingly, if multiple core peptides were matched to single glycopeptide group (cluster), the most abundant core peptide is most plausible, and thus the results of the most abundant core are displayed at the top of the list.
2) **Glycan abundance:** In the process of IGOT-LC/MS/MS for preparing a core peptide list, the major glycan compositions (glycomes) of the sample can be determined by collecting the released glycans and analyzing them, for example, by MALDI-MS. As is the case with core peptide abundance, it is expected that the major glycan compositions are likely to attach to major glycopeptides. Thus, GRable utilize a glycan point list, in which glycan compositions that are detected actually in the glycome of sample are given a point. In the matching module, the sum of the points within the cluster is calculated as a total score, which is used for selection of the plausible match. That is, a match with higher score is considered better among candidate matches for each cluster. When an actual glycan composition list is unavailable, a glycan point list can be made considering the N-glycan processing pathways; in the list, common and unusual compositions have positive and negative points, respectively. In this selection module, users can select whether negative points are considered or not.
3) **RT difference between glycopeptides and corresponding core peptides:** Since the RT of a core peptide is close to those of neutral glycopeptides and often delayed behind them, core peptides with the RT outside the setting are unlikely to be true. However, when a sialic acid is added to a glycopeptide, the RT becomes longer, which is contrast with the case when a neutral glycan is added. Thus, if the matched glycans are sialylated, it should be paid attention to evaluate using RT difference.
4) **Mass accuracy:** In the clustering and matching steps, mass tolerance is usually set to 5 ppm, which is slightly tighter than that used for database searches in proteome analysis (i.e., 7 ppm). To guarantee highly reliable results, in the selection step, delta mass is usually set to 2 ppm. For this reason, MS1 should be acquired under the higher resolution, as described above (Section 2.2).
5) **Utilization with MS2 information:** Although this Glyco-RIDGE method is performed only with MS1 information in the clustering and matching steps to listing the glycopeptide candidates, MS2 information is also utilized for selecting the most plausible matching. In the selection step, it is checked whether MS2 information is obtained for each cluster member. In this module, users can select whether MS2 information is used for analysis, and if it is acquired, the following information is retrieved and used for selection.

i. Presence of diagnostic ions: The presence or absence of diagnostic ions derived from glycans (e.g., HexNAc(204) and Hex+HexNAc(366)) and their signal intensity are obtained automatically for all the MS2 scans. This is supportive information for each matching result, confirming that the ion is derived from a glycopeptide.

ii. Presence of glycopeptide ions: GRable checks the presence of ions of a peptide and a peptide remaining the innermost GlcNAc of N-glycan, called Y0 and Y1, respectively, and ions that occur frequently around them (e.g., Y1+Fuc and Y2). If two of these ions are found and the mass of observed or calculated Y0 is identical to that of a matched peptide within a given mass tolerance, the result is a strong endorsement of the matching result.

iii. DB search results: GRable does not identify a glycopeptide directly based on MS2 spectrum analysis, but it collects the MS2-based search results for the evaluation of the matching results. In a selection result sheet, a peptide sequence corresponding the Y0 mass in a core peptide list is provided, which facilitates easy evaluation of the result.

##### 3.8.2. Operation procedure

To perform selection, press the wizard button (Figure 2-2①) from the main window to display the selection module (Figure 3-9).

① Select the “Selection” tab.
② Enter an analysis name.
③ Set parameters (Table 3-5). Note that “input Analysis Setting” is unapplicable in this version.
④ Select whether the setting is saved.
⑤ Click the “Start” button to start analysis. When “Analysis complete” is displayed, the process is finished.
⑥ If you want to cancel this step in progress, press the "Cancel" button.

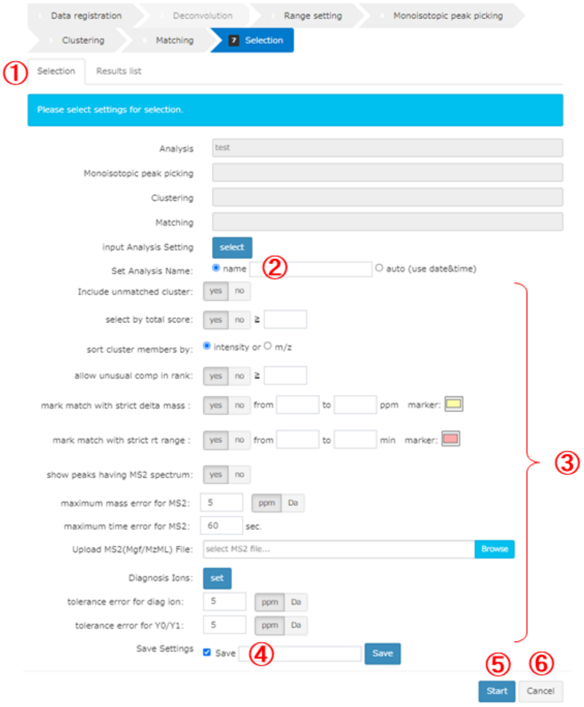

**Figure 3-9. Selection module.**

**Table 3-5. Details of Selection setting.**

| No. | Parameter | Description |
|---|---|---|
| 1 | Include unmatched cluster | If “yes” is selected, only the cluster information will be included in matching results for clusters that did not match at all. |
| 2 | select by total score | Selects whether a threshold of total score is used for selection. If “yes” is selected, matching results contain only matches whose total score is greater than or equal to the threshold value set by the user. For clusters that have no matches above the prescribed score, only the cluster information will be included in matching results. |
| 3 | sort cluster members by | Selects the sorting method for cluster members.<br>- “intensity”: sort in descending order of intensity<br>- “m/z”: sort in descending order of mass |
| 4 | allow unusual comp in rank | If “no” is selected, signals with rank equal to or greater than the user's specified number and with unusual comp will be not contained in selection results. |
| 5 | mark match with strict delta mass | If “yes” is selected, matches that have difference of masses (delta(ppm)) between observed and theoretical glycopeptides within the mass range (specified by user in ppm) will be considered “plausible” and marked with color (specified by user) in an export file. Note that this parameter is used only for marking to allow manual inspection, and thus all the selected glycopeptide signals (even with out of range) are included in the selection results. |
| 6 | mark match with strict rt range | If “yes” is selected, matches that have difference of RTs (delta(RT)) between a glycopeptide and core peptide within the RT range (specified by user in min) will be considered “plausible” and marked with color (specified by user) in an export file. Note that this parameter is used only for marking to allow manual inspection, and thus all the selected glycopeptide signals (even with out of range) are included in the selection results. |
| 7 | show peaks having MS2 spectrum | To utilize MS2 information, selects “yes” and specifies the subsequent parameters. If “yes” is selected, glycopeptide signals with MS2 information will be marked with “1” in a “MS2?” column of a selection result sheet. |
| 8 | maximum mass error for MS2 | Sets the maximum mass tolerance (ppm or Da) for searching corresponding MS2 spectra for each clustered glycopeptide. If the difference of masses (deltaMH+(Da)) between a clustered glycopeptide (m/z) and calculated one (MH+calc) based on MS2 information (i.e., charge and precursor mass) are smaller than the threshold, the corresponding MS2 information will be indicated in a “MS2 info for cluster” sheet. |
| 9 | maximum time error for MS2 | Sets the maximum RT tolerance (sec) for searching corresponding MS2 spectra for each clustered glycopeptide. If the difference of RTs (delta rt) between a clustered glycopeptide (rt) and MS2 signal |

| No. | Parameter | Description |
|---|---|---|
| | | (rt(min)) are smaller than the threshold, the corresponding MS2 information will be indicated in a “MS2 info for cluster” sheet. |
| 10 | Upload MS2(Mgf/MzML) File | Uploads MS2 data (in mgf or mzML format). When a mzML format file is used, be sure to use MS2 data before deconvolution. |
| 11 | Diagnosis Ions | Specifies glycan fragment ions used as diagnostic ions to confirm that the MS2 spectra is surely derived from a glycoprotein. The panel that appears with the “set” button allows setting type of diagnostic ions with their mass (Da) (Figure 3-10). |
| 12 | tolerance error for diag ion | Sets the mass tolerance (ppm or Da) between theoretical and observed fragment ions for evaluating the presence of diagnostic ions specified by user as above. |
| 13 | tolerance error for Y0/Y1 | Sets the mass tolerance (ppm or Da) between theoretical and observed fragment ions for evaluating the presence of Y0 and its related ions. |

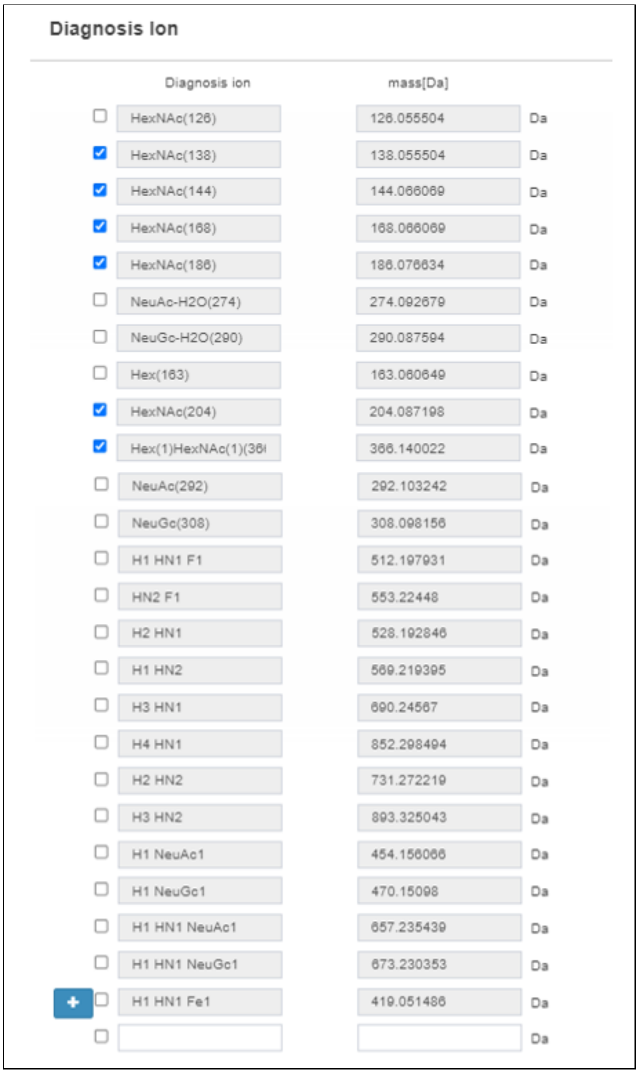

**Figure 3-10.  Setting dialog of diagnosis ion.**

#### 3.9. Data tree

Data tree displays results of each step for one Analysis project in a tree-view format. To display the tree, move the cursor to the tree button on the left of the main window (Figure 2-2⑪) to display the data tree (Figure 3-11). By right clicking data of interest in the tree, a menu will appear for executing functions (Table 3-6).

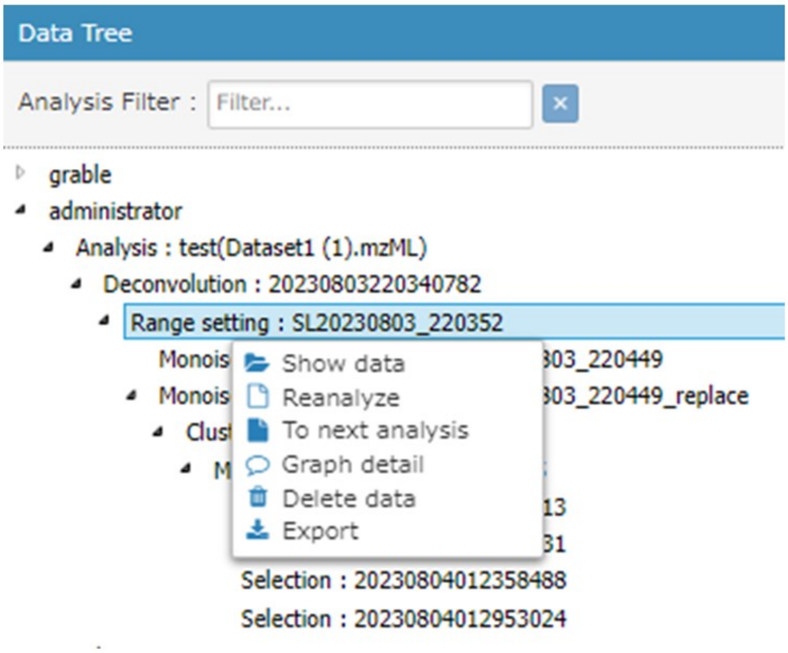

**Figure 3-11. Representative image of data tree.**

**Table 3-6. Details of data tree functions.**

| No. | Function | Description |
|---|---|---|
| 1 | Show data | This function is to display the selected result data on a viewer in the main window. Details of the viewer are described in Section 3.10. This operation is valid only for results of "range setting", "monoisotopic peak picking", "clustering", and "matching" steps. |
| 2 | Reanalyze | This function is to re-analyze the selected data with another different setting in a selected step. The parameters shown in the setting dialog are the same ones with the selected data. |
| 3 | To next analysis | This function is to proceed to next step using the selected result data. |
| 4 | Graph detail | This function is to display the settings of selected analysis in a separate dialog. |
| 5 | Delete data | This function is to delete selected data. |
| 6 | Export | This function is to export input data and analysis results along with search parameter settings. To export them, check the items to be exported in the download dialog (Figure 3-12) and click on the "download" button. Details of exported files are described in Section 5. Note that the file size of peak list is large and thus it is recommended to uncheck for the peak list unless necessary. The peak list is separately exported as a .csv file. The other selected items are exported in one Excel file. |

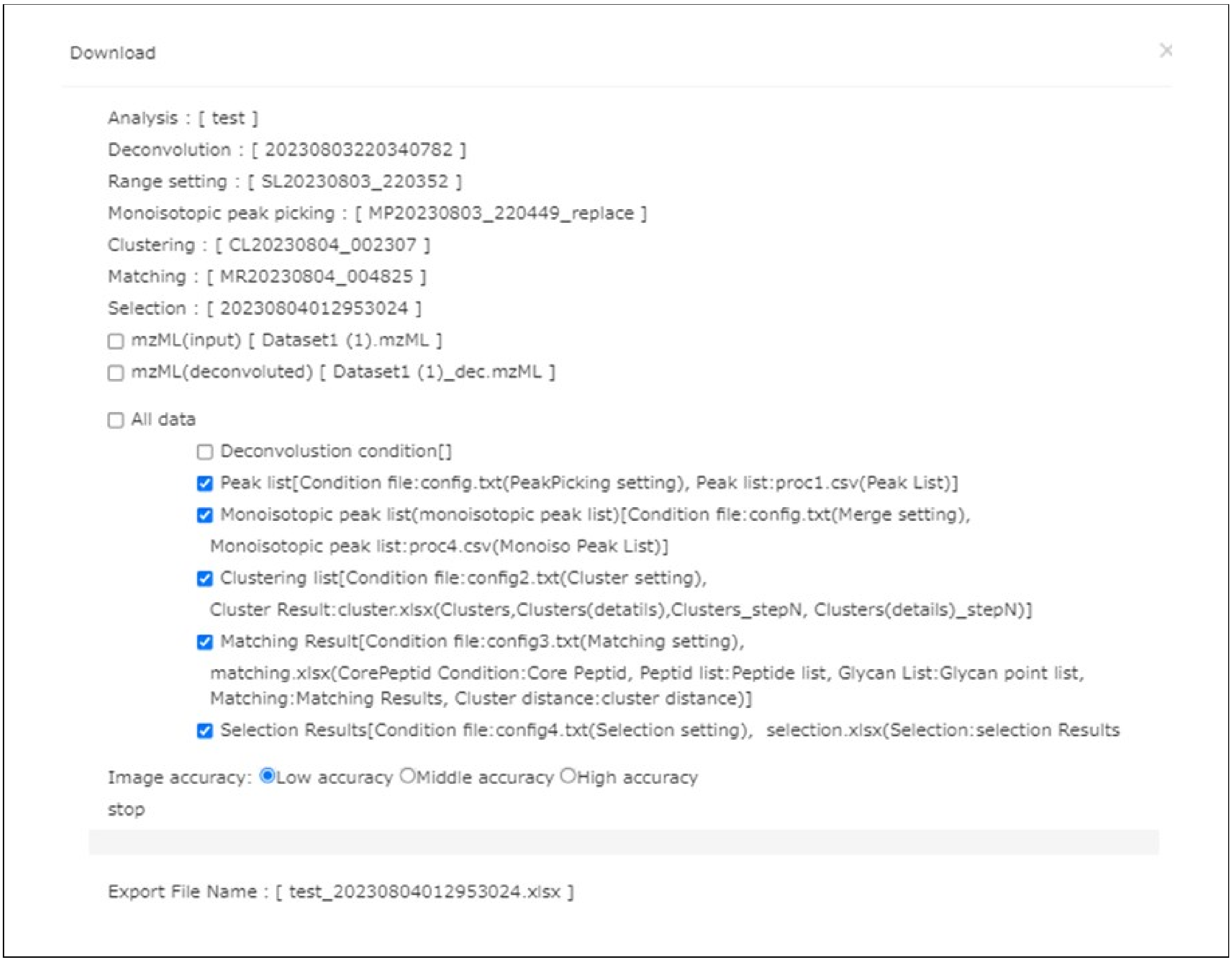

**Figure 3-12. Download dialog.**

#### 3.10. Viewer

The viewer in the main window (Figure 3-13) is equipped for a graphical visualization of analysis results. The data to be shown can be selected in a tree view as documented above (Section 3.9) and its visualization can be optimized using functional buttons (Table 3-13).

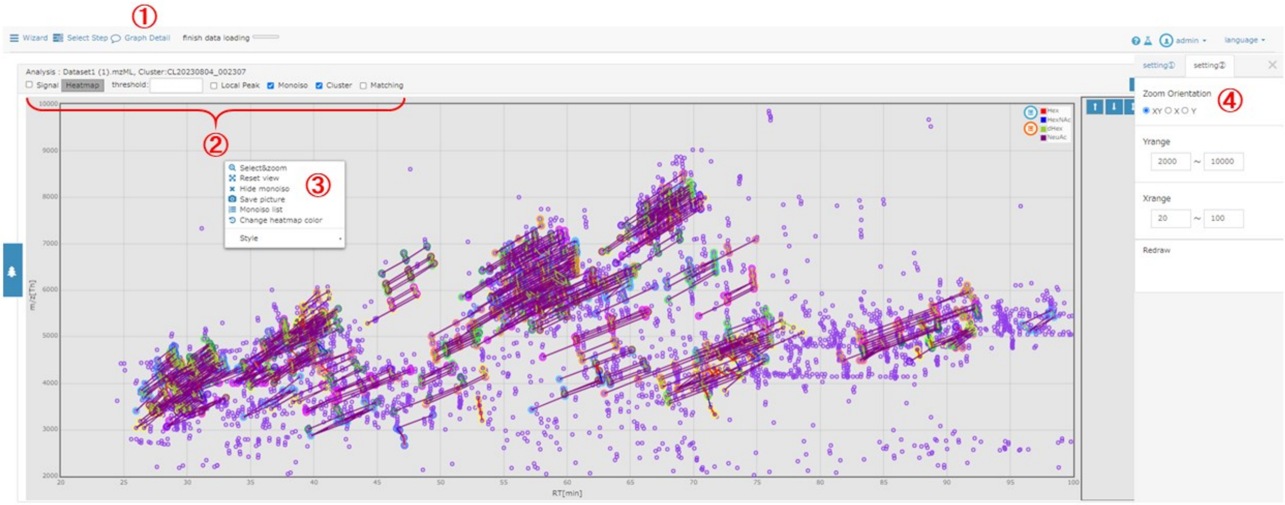

**Figure 3-13. Viewer in the main window.**

**Table 3-13. Details of viewer functions.**

| No. | Function | Description |
|---|---|---|
| 1 | Graph Detail | This function is to display the settings of selected analysis in a separate dialog. |
| 2 | Panels | "Analysis" indicates analysis name along with the name of selected results. A "Heatmap" button and threshold entry form are valid only when the "Signal" is checked. User can the display mode of data among "Local Peak", "Monoiso", "Cluster", and "Matching".<br><br>- **Local Peak**: Displays all the detected local peaks.<br>- **Monoiso**: Displays all the detected monoisotopic peaks in black circle.<br>- **Cluster**: Displays all the detected glycopeptide clusters, in which line colors indicate the relationship of glycan units between two glycopeptide signals.<br>- **Matching**: Displays only the matched glycopeptide clusters when the "Cluster" is unchecked.<br><br>Note that display of "Signal" and "Local Peak" is slow and thus it is recommended to uncheck these buttons. |
| 3 | Select&zoom | This function is to display the enlarged graph within range specified by using a mouse. |
| 3 | Reset view | The function is to return to default setting. |
| 3 | Hide monoiso | This function is to display only monoisotopic peaks by hiding clusters. |
| 3 | Save picture | This function is to save the image of viewer as a PNG file. |
| 3 | Monoiso list | This function is to display a dialog of monoisotopic peak list. Details of the operation of this list are described in Section 3.11. |
| 3 | Change heatmap color | This function is unapplicable in this version. |
| 3 | Style | Using this function, the style of monoisotopic peaks can be changed. |
| 4 | Zoom Orientation | Specifies the orientation when the viewer is enlarged and reduced using a mouse wheel. |
| 4 | Y range / X range | Specifies the range of data (X: RT and Y: m/z) to be visualized. When the "Redraw" button is clicked, the setting values will be applied. |

#### 3.11. Monoisotopic peak list

##### 3.11.1. Functional overview

The monoisotopic peak list (Figure 3-14) displays monoisotopic peaks obtained by the monoisotopic peak picking (Step 4) in a dialog format. To display this list, show the analysis results after Step 4 according to the aforementioned instruction (Section 3.10) and then press the "option" button in the main window (Figure 2-2⑨). This dialog is moveable, expandable and contractable. The items of the list are summarized in Table 3-14.

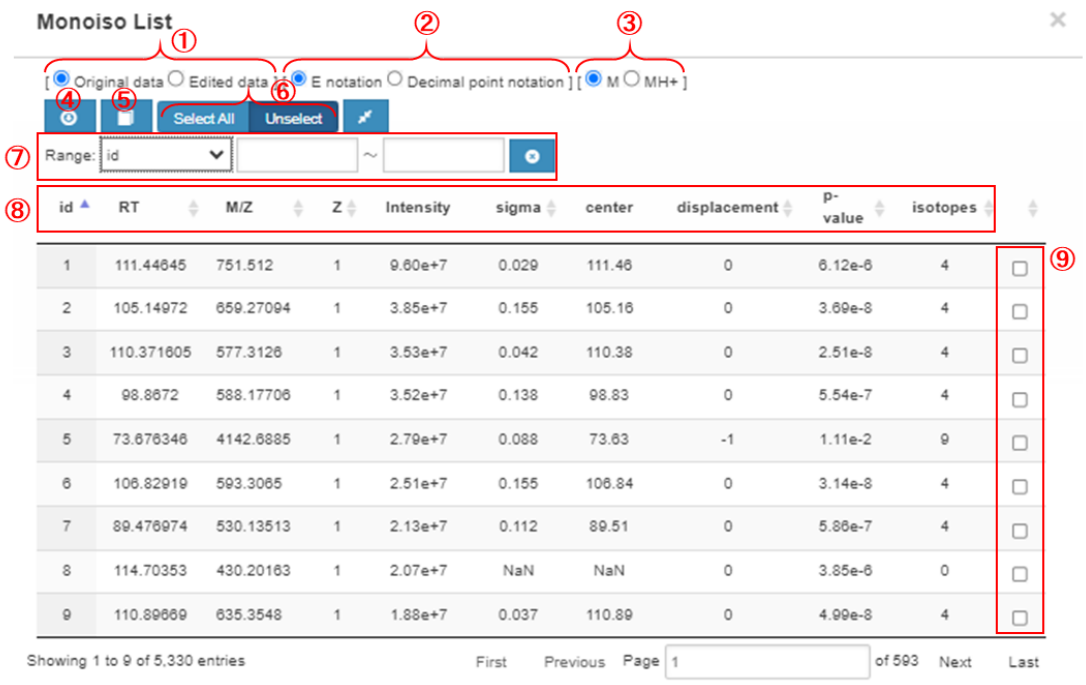

**Figure 3-14. Monoisotopic peak list.**

**Table 3-14. Details of monoisotopic peak list items.**

| No. | Function / Item | Overview |
|---|---|---|
| 1 | Data switching | This function is to switch between original data and edited data. In an initial state, original data are displayed. |
| 2 | Decimal display switching | The display of “intensity” can be switched between E format and decimal format. An initial state is the display in the E format. |
| 3 | M / MH$^+$ switching | The display of “M/Z” can be switched between M and MH$^+$. An initial state is the display in the M format. |
| 4 | Data download | The monoisotopic peak list can be downloaded in .csv format. |
| 5 | Save edits | The modified list can be saved. This button is used for data correction function described below. |
| 6 | Select All / Unselect | All signals in the list can be selected or unselected together. This button is used for data correction function described below. |
| 7 | Filter | Signals in the list can be filtered for each item with ranges specified by user. To use this function, select an item in the dropdown list and then enter the range you want to display in the textbox. The filtering can be reset by press the “x” button. Note that only half-width numbers can be entered. This button is used for data correction function described below. |
| 8 | id | ID number of the isotope group |
| 8 | RT | RT of the isotope group |
| 8 | M/Z | m/z of the monoisotopic peak |
| 8 | Z | Charge state of the monoisotopic peak. |
| 8 | Intensity | Signal intensity of the monoisotopic peak |
| 8 | Sigma | Width of chromatographic distribution (average approximately 0.1) |
| 8 | Center | RT of the monoisotopic peak. |

| No. | Function / Item | Overview |
|---|---|---|
| 8 | displacement | The degree of monoisotopic shift expressed as a difference from a multinomial distribution fitting curve in the spectrum. The theoretical spectrum is created based on abundance ratios of four isotopes including C, N, O, and S. |
| 8 | p-value | The *P*-value for the fitting of an observed spectrum to its theoretical spectrum. |
| 8 | isotopes | The number of isotopes |
| 9 | Check box | Each ID can be selected/unselected individually by checking/unchecking this check box. This function is used for data correction function described below. |

##### 3.11.2. Details of the correction function

The data correction function can be used in the monoisotopic peak list (Figure 3-15) according to the following procedure.

① Select “replace” in the dropdown list.

② Enter the range of “displacement” values to be corrected. At once the range is entered, the list is updated.

③ Click the “Select All” to check all the selected IDs. Note that this step will take time.

④ When the update is finished, “SelectAll Complete” will be appeared and all the check box in the list will be checked.

⑤ Click the “Save edits” button, enter an analysis name, and then click the “create” button to save the edited data as a new one. Note that this step will take time.

⑥ The updated data will appear in the data tree of the analysis (Section 3.9).

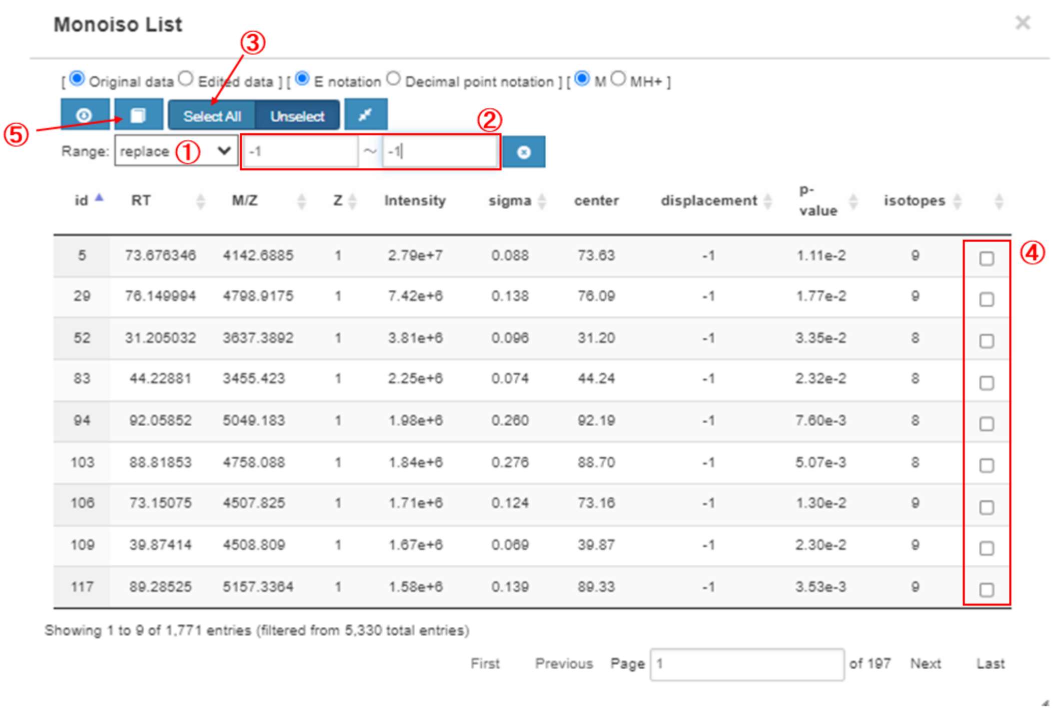

**Figure 3-15. Procedure of correction function.**

### 4. Preparation details of import files

#### 4.1. Core peptide list (.xlsx)

To prepare the core peptide list used for matching (Step 6), we recommend performing isotope-coded glycosylation site-specific tagging (IGOT)-LC-MS/MS as follows. An aliquot of glycopeptides is treated with peptide-*N*-glycosidase F (PNGase F) in $^{18}O$-labeled water to remove *N*-glycan and to label deglycosylated Asn as $^{18}O$-labeled Asp (i.e., IGOT). The IGOT-treated deglycopeptides is analyzed by LC/MS/MS and the MS2 data were used for database search using Mascot. Using Mascot export file, the core peptide list is created in a fixed format (Figure 4-1). In the list, some items should be manually added to Mascot export file, some of which are mandatory for analysis (Table 4-1). If IGOT-LC-MS/MS is unavailable, the core peptide list can be creased based on proteome results. For analysis of a highly purified glycoprotein, it is also possible to list up peptides predicted based on protease digestion patterns, albeit there is a caution for RT (See Table 4-1). Note that unknown parameters should be indicated as “0” (not to remain blank) to ensure correct performance.

**Note that items minimally required for preparing a core peptide list are the calculated mass, retention time of each peptide and at least one peptide identifier such as CP (core peptide) number.** Accordingly, other information on core peptides including protein accession number, protein description, peptide position, modification, signal intensity, and protein sequence, can be liked to assigned glycopeptides after analysis. To simplify this operation, we provide a format of core peptide list for including these information in advance.

<table>
<tr><th></th><th>A</th><th>B</th><th>C</th><th>D</th><th>E</th><th>F</th><th>G</th><th>H</th><th>I</th><th>J</th><th>K</th><th>L</th><th>M</th><th>N</th></tr>
<tr><td>1</td><td>CP No.</td><td>prot_acc</td><td>prot_desc</td><td>pep_star</td><td>pep_end</td><td>pep_seq</td><td>pep_var_mod_pos</td><td>Siteseqp</td><td>Nigot</td><td>pep_calc</td><td>Mpep</td><td>prot_seq</td><td>intensity</td><td>RT</td></tr>
<tr><td>2</td><td>1</td><td>P19652</td><td>>sp|P19652|A1AG</td><td>19</td><td>38</td><td>QIPLCANLVPVPITNATLDR</td><td>3.00000000000000200000.0</td><td>33(NAT)/</td><td>1</td><td>2190.16</td><td>2187.17</td><td>MALSWV</td><td>1.1E+08</td><td>89.9836</td></tr>
<tr><td>3</td><td>2</td><td>P02763</td><td>>sp|P02763|A1AG</td><td>19</td><td>42</td><td>QIPLCANLVPVPITNATLDQITGK</td><td>3.00000000000000002000000000.</td><td>33(NAT)/</td><td>1</td><td>2561.37</td><td>2558.38</td><td>MALSWV</td><td>3.5E+07</td><td>95.6326</td></tr>
<tr><td>4</td><td>3</td><td>P19652</td><td>>sp|P19652|A1AG</td><td>19</td><td>38</td><td>QIPLCANLVPVPITNATLDR</td><td>0.00000000000000200000.0</td><td>33(NAT)/</td><td>1</td><td>2207.19</td><td>2204.2</td><td>MALSWV</td><td>3E+07</td><td>89.9673</td></tr>
<tr><td>5</td><td>4</td><td>P19652</td><td>>sp|P19652|A1AG</td><td>19</td><td>42</td><td>QIPLCANLVPVPITNATLDRITGK</td><td>3.00000000000000002000000000.</td><td>33(NAT)/</td><td>1</td><td>2589.41</td><td>2586.42</td><td>MALSWV</td><td>7390537</td><td>90.2897</td></tr>
<tr><td>6</td><td>5</td><td>P19652</td><td>>sp|P19652|A1AG</td><td>102</td><td>113</td><td>ENGTVSRYEGGR</td><td>0.020000000000.0</td><td>103(NGT</td><td>1</td><td>1326.61</td><td>1323.62</td><td>MALSWV</td><td>6000682</td><td>34.6546</td></tr>
<tr><td>7</td><td>6</td><td>P00738</td><td>>sp|P00738|HPT_</td><td>236</td><td>249</td><td>VVLHPNYSQVDIGL</td><td>0.00000200000000.0</td><td>241(NYS)</td><td>1</td><td>1555.81</td><td>1552.82</td><td>MSALGA</td><td>4919720</td><td>69.4705</td></tr>
<tr><td>8</td><td>7</td><td>P02765</td><td>>sp|P02765|FETUA</td><td>145</td><td>165</td><td>VCQDCPLLAPLNDTRVVHAAK</td><td>0.0000000000002000000000.0</td><td>156(NDT</td><td>1</td><td>2379.19</td><td>2376.2</td><td>MKSLVLLI</td><td>3312263</td><td>63.1591</td></tr>
<tr><td>9</td><td>8</td><td>P19652</td><td>>sp|P19652|A1AG</td><td>87</td><td>101</td><td>QNQCFYNSSYLNVQR</td><td>0.000000200000000.0</td><td>93(NSS)/</td><td>1</td><td>1922.85</td><td>1919.86</td><td>MALSWV</td><td>2250460</td><td>60.5409</td></tr>
<tr><td>10</td><td>9</td><td>P19652</td><td>>sp|P19652|A1AG</td><td>19</td><td>42</td><td>QIPLCANLVPVPITNATLDRITGK</td><td>0.00000000000000002000000000.</td><td>33(NAT)/</td><td>1</td><td>2606.44</td><td>2603.45</td><td>MALSWV</td><td>1832450</td><td>90.0245</td></tr>
<tr><td>11</td><td>10</td><td>P19652</td><td>>sp|P19652|A1AG</td><td>26</td><td>38</td><td>LVPVPITNATLDR</td><td>0.0000000200000.0</td><td>33(NAT)/</td><td>1</td><td>1410.8</td><td>1407.81</td><td>MALSWV</td><td>1713041</td><td>64.1101</td></tr>
</table>

**Figure 4-1. Representative image of core peptide list.**

**Table 4-1. Details of core peptide list.**

| No. | Manually addition | Mandatory | Item | Description |
|---|---|---|---|---|
| 1 | Required | No, but reccomended* | CP No. | Core peptide number. *Note that this information is not required for matching but recommended to be included as a peptide identifier. It is also recommended to be manually numbered in descending order of signal intensity, as an indicator of the relative abundance of signal intensity. |
| 2 | | | prot_acc | Protein accession No. |
| 3 | | | prot_desc | Protein description |
| 4 | | | pep_start | Start position of peptide within a protein |
| 5 | | | pep_end | End position of peptide within a protein |

| No. | Manually addition | Mandatory | Item | Description |
|---|---|---|---|---|
| 6 | | No, but reccomended* | pep_seq | Peptide sequence. *Note that this information is not required for matching but used in the Selection step; when a peptide having the identical mass to predicted Y0 mass is found in a core peptide list, the peptide sequence will be shown in the Selection results sheet. |
| 7 | | | pep_var_ mod_pos | Position of the following variable modifications:<br>1: ammonia-loss (peptide N-term, carbamidomethyl C)<br>2: Delta:H(-1)N(-1)18O(1)(N)<br>3: Gln->pyro-Glu (peptide N-term, Q)<br>4: oxidation (M) |
| 8 | Required | | Siteseqpos | Position of IGOT-labeled Asn residue (+2.98822096000004) within a protein along with its consensus sequence |
| 9 | Required | | Nigot | Number of IGOT-labeled Asn residues. Note that this parameter is not mandatory for analysis but used for calculation of glycan compositions after subtraction of Hex(3)HexNAc(2) (trimannosyl core) in the Matching results sheet. |
| 10 | | | pep_calc_ mr | Calculated mass of peptide |
| 11 | | yes | Mpep | Calculated mass of peptide without IGOT labeling. Note that this information is the most important to obtain results. |
| 12 | | | prot_seq | Protein sequence |
| 13 | | No, but reccomended* | intensity | Signal intensity. *Note that this information is not required for matching but recommended to be included as an indicator of the abundance of peptide. When candidate core peptides are identified using Mascot. MS2 total ion current can be used because of the ease of availability. |
| 14 | Required | yes* | RT | RT (min) of MS2 fragment ion. Note that RT (sec) in mascot export files should be converted into RT (min). *Note that for analysis of purified glycoproteins, analysis may work well without RT, although it causes a decrease in the reliability of results. |

#### 4.2. Glycan point list (.xlsx)

The glycan point list used for the matching (Step 6) can be prepared based on glycans observed in glycome analysis of the same glycoprotein sample and/or the biosynthetic pathway of glycans. In the glycan point list (Figure 4-2), glycan compositions (highlighted in pink) are listed along with a given point in a “point” raw. Unusual glycan compositions can be also specified by listing along with a negative point in a “unusual” raw. These unusual glycan compositions are used for analysis only when the “uncomp negative point” parameter is “yes” in the matching setting (Table 3-4). Note that the name of glycan unit (e.g., Hex, HexNAc, dHex, and NeuAc) should be the same as those in the matching setting (Table 3-4).

| | A | B | C | D | E | F | G |
|---|---|---|---|---|---|---|---|
| 1 | | point | Hex | HexNAc | dHex | NeuAc | |
| 2 | point | 1 | 0 | 1 | 0 | 0 | |
| 3 | point | 1 | 0 | 1 | 1 | 0 | |
| 4 | point | 1 | 0 | 2 | 0 | 0 | |
| 5 | point | 1 | 0 | 2 | 1 | 0 | |
| 6 | point | 1 | 1 | 2 | 0 | 0 | |
| 7 | point | 1 | 1 | 2 | 1 | 0 | |
| 8 | point | 1 | 2 | 2 | 0 | 0 | |
| 9 | point | 1 | 2 | 2 | 1 | 0 | |
| 10 | point | 1 | 3 | 2 | 0 | 0 | |
| 11 | point | 1 | 3 | 2 | 1 | 0 | |
| 12 | point | 1 | 4 | 2 | 0 | 0 | |
| 13 | point | 1 | 4 | 2 | 1 | 0 | |
| 14 | point | 1 | 5 | 2 | 0 | 0 | |
| 15 | point | 1 | 5 | 2 | 1 | 0 | |
| 16 | point | 1 | 6 | 2 | 0 | 0 | |
| 17 | point | 1 | 6 | 2 | 1 | 0 | |
| 18 | | | | | | | |
| 19 | unusual | -1 | 9 | 2 | 1 | | |
| 20 | unusual | -1 | 9 | 2 | 2 | | |
| 21 | unusual | -1 | 9 | 2 | 3 | | |
| 22 | unusual | -1 | 9 | 2 | 4 | | |
| 23 | unusual | -1 | 8 | 2 | 1 | | |
| 24 | unusual | -1 | 8 | 2 | 2 | | |
| 25 | unusual | -1 | 8 | 2 | 3 | | |
| 26 | unusual | -1 | 8 | 2 | 4 | | |

**Figure 4-2. Representative image of glycan point list.**

### 5. Details of export files

Analysis results can be exported using the “Export” function in the data tree (Section 3.9). Peak list (Table 5-1) is separately exported as a .csv file. The other selected items (Table 5-2) are exported in one Excel file.

**Table 5-1. Details of peak list.**

| Item | Description |
|---|---|
| SCAN | Scan number of MS1 spectra |
| RT | RT (min) of a local peak |
| MASS | Observed mass (m/z) of a local peak |
| INTENSITY | Signal intensity of a local peak |
| GID | Glycopeptide ID that is the same with the “monoiso no” in the “Monoiso Peak List” sheet |
| FLAG | This parameter is not worked. |

**Table 5-2. List of data exported in an Excel file.**

| Sheet name | Related step | Description | Details of items |
|---|---|---|---|
| Range setting | Range setting | Parameters specified for this step | |

| Sheet name | Related step | Description | Details of items |
|---|---|---|---|
| Monoiso Peak List | Monoisotopic peak picking | List of monoisotopic peaks detected in this step | Table 5-3 |
| Monoiso Peak Picking log | Monoisotopic peak picking | Log information for this step | |
| Monoiso Peak Picking setting | Monoisotopic peak picking | Parameters specified for this step | |
| Clustering setting | Clustering | Parameters specified for this step | |
| Clusters | Clustering | List of clusters detected in this step | Table 5-4 |
| Clusters(details) | Clustering | Detailed information of clusters detected in this step | Table 5-5 |
| Clusters_stepX | Clustering | List of clusters detected only in the applied setting (stepX) | Table 5-4 |
| Clusters(details)_stepX | Clustering | Detailed information of clusters detected only in the applied setting (stepX) | Table 5-5 |
| Matching setting | Matching | Parameters specified for this step | |
| Glycan point list | Matching | Glycan point list identical to an input file | See Section 4.2 |
| Peptide list | Matching | Core peptide list identical to an input file | See Section 4.1 |
| Matching Results | Matching | List of all matches of glycopeptide cluster, core peptide, and glycan compositions | Table 5-6 |
| relation between clusters | Matching | Inter-cluster analysis results (*Included only when inter-cluster analysis is applied) | Table 5-7 |
| Selection setting | Selection | Parameters specified for this step | |
| Selection Results | Selection | List of matches selected based on the specified parameters | Table 5-6 |
| MS2 info for clusters | Selection | MS2 information for each selected cluster | Table 5-8 |
| All MS2 info | Selection | MS2 information for all scans in input MS2 data | Table 5-8 |
| logging | Selection | Log information for this step | |
| peptide chart | Selection | 2D-map (RT – Mass of predicted Y0 ion) for core peptides observed in the "All MS2 info" sheet | Table 5-8 |

**Table 5-3. Details of monoisotopic peak list.**

| Item | Description |
|---|---|
| monoiso no | Monoisotopic peak No. that are numbered in a descending order of peak intensity |
| monoiso retention time | RT (min) of monoisotopic peak |
| monoiso mass | Mass (M) of monoisotopic peak |
| peak intensity | Signal intensity of a local peak |

| Item | Description |
|---|---|
| monoiso rtid | This parameter is not worked. |
| monoiso intensity | Signal intensity of a monoisotopic peak |
| charge | Charge state |
| center of distribution | RT of the center of chromatographic distribution |
| sigma of distribution | Width of chromatographic distribution (average approximately 0.1) |
| valid groups | Number of isotopes |
| multinomial pvalue | The P-value for the fitting of an observed spectrum to its theoretical spectrum. |
| composition | This parameter is a part of algorithm for the fitting of an observed spectrum to its theoretical spectrum. |
| suggest monoiso displacement | The degree of monoisotopic shift expressed as a difference from a multinomial distribution fitting curve in the spectrum. |
| suggested monoiso mass | Mass of a monoisotopic peak selected after check using the correction function |
| pvalue suggested monoiso | The P-value for a suggested monoisotopic peak |
| suggested monoiso signal found | This parameter is not worked. (Always “0”) |
| original Mz | Mass of a monoisotopic peak selected before the correction function |
| update flag | If there is an update using the correction function, “1” is indicated. |

**Table 5-4. Details of Clustering result sheets.**

| Item | Description |
|---|---|
| cluster_no | Cluster No. that is numbered in a descending order of the maximum of intensity within the cluster. |
| peak_no | Monoisotopic peak No. that is the same as “monoiso no” in the monoisotopic peak list. |
| charge | Charge state of the monoisotopic peak |
| m/z | m/z of the monoisotopic peak<br>(*When “Peaks with the same mass are counted as one” is “yes” in the setting, the masses having the same mass withing the cluster are highlighted in yellow) |
| rt | RT (min) of the monoisotopic peak |
| intensity | Signal intensity of the monoisotopic peak |
| no_member | No. of members for the cluster<br>(*When “Peaks with the same mass are counted as one” is “yes” in the setting, the masses having the same mass withing the cluster are counted as one) |
| step | Step No. that found the monoisotopic peak |

**Table 5-5. Details of Clustering result (details) sheet.**

| Item* | Description |
|---|---|
| cluster_no | Cluster No. |
| charge | Charge state of the monoisotopic peak |
| peak_no | Monoisotopic peak No. that is the same as “monoiso no” in the monoisotopic peak list. |
| rt | RT (min) |
| scan_no | Scan No. |
| m/z | Mass ($MH^+$) |
| mass | Mass (M) |
| intensity | Signal intensity |
| target_peak_no | Monoisotopic peak No. that is the same as “monoiso no” in the monoisotopic peak list. |
| rt | RT (min) |
| scan_no | Scan No. |
| m/z | Mass ($MH^+$) |
| mass | Mass (M) |
| intensity | Signal intensity |
| residue | A glycan unit equivalent to the difference between reference and target peaks |
| diff_mass | Difference in observed masses between reference and target peaks |
| diff_theor | Theoretical mass difference between reference and target peaks, which corresponds to the theoretical mass of the glycan unit |
| delta | Delta value (Da) between “diff_mass” and “diff_theor”. |
| diff(ppm) | Delta value (ppm) converted from “delta” by the method selected for “Error Evaluation” of the setting |
| diff_rt | Difference in RTs between reference and target peaks |
| step | Step No. that found the monoisotopic peak |

*Red: information on reference peaks. Blue: information on target peaks examined for relationship with reference peaks.

**Table 5-6. Details of matching and selection result sheets.**

| Item *1 | Description |
|---|---|
| cluster_no | Cluster No. |
| member | No. of the members within the cluster |
| peak_no | Monoisotopic peak no. that is the same as “monoiso no” in the monoisotopic peak list. |
| charge | Charge state |
| scan_no | Scan No. of the monoisotopic peak |
| m/z | Mass ($MH^+$) |
| rt | RT (min) |
| intensity | Signal intensity |

| Item *1 | Description |
|---|---|
| M(gpep, obs) | Observed mass (M) |
| step | Step No. that found the monoisotopic peak |
| CP No. | Core peptide No. |
| prot_acc | Protein accession No. |
| prot_desc | Protein description |
| pep_start | Start position of peptide within a protein |
| pep_end | End position of peptide within a protein |
| pep_seq | Peptide sequence |
| pep_var_mod_pos | Position of the following variable modifications:<br>1: ammonia-loss (peptide N-term, carbamidomethyl C)<br>2: Delta:H(-1)N(-1)18O(1)(N)<br>3: Gln->pyro-Glu (peptide N-term, Q)<br>4: oxidation (M) |
| Siteseqpos | Position of IGOT-labeled Asn residue (+2.98822096000004) within a protein along with its consensus sequence |
| Nigot | Number of IGOT-labeled Asn residues |
| pep_calc_mr | Calculated mass of peptide |
| Mpep | Calculated mass of peptide without IGOT labeling |
| prot_seq | Protein sequence |
| intensity | Signal intensity of MS2 fragment ion |
| RT | RT (min) of MS2 fragment ion. Note that RT (sec) in mascot export files should be converted into RT (min) |
| rank | Ion score rank in the mascot search results |
| Hex | No. of Hex residues in the matched glycan composition |
| HexNAc | No. of HexNAc residues in the matched glycan composition |
| dHex | No. of dHex residues in the matched glycan composition |
| NeuAc | No. of NeuAc residues in the matched glycan composition |
| Hex[-core] | No. of Hex residues after subtracting by the trimannosyl core of N-glycan (i.e. -3) in the matched glycan composition |
| HexNAc[-core] | No. of HexNAc residues after subtracting by the trimannosyl core of N-glycan (i.e. -2) in the matched glycan composition |
| dHex | No. of dHex residues in the matched glycan composition |
| NeuAc | No. of NeuAc residues in the matched glycan composition |
| unusual comp | If the glycan composition is defined as “unusual” in the glycan point list, “1” will be indicated. |
| point | A point given for the glycan composition |
| factor | A coefficient value for weighting a glycan composition of interest. The factor is “1” unless other values are specified in the glycan point list. Note that this function is not guaranteed in this version. |
| score | A score for the glycan composition calculated by multiplying “point” and |

| Item *1 | Description |
|---|---|
| | "factor" |
| total score | The sum of scores for all the members of the cluster |
| mass | Calculated mass (M) of theoretical glycopeptide in a combination of the core peptide and glycan composition |
| delta(Da) | Delta value (Da) between observed and calculated masses for the glycopeptide |
| delta(ppm) | Delta value (ppm) converted from delta(Da) |
| delta(RT) | Delta value (min) between RTs of the glycopeptide and core peptide |
| MS2? *2 | If the MS1 spectra has any MS2 information, "1" will be indicated with a link to the corresponding data in the "MS2 info for clusters" sheet. |
| predict peptide *2 | This is identical to the "peptide[seq]" information in the "MS2 info for clusters" sheet. |
| Y0 (z=1) *2 | Y0 mass ($MH^+$; z=1) calculated using the mass of core peptide |
| Y0 (z=2) *2 | Y0 mass ($MH^+$; z=2) calculated using the mass of core peptide |
| Y0 (z=3) *2 | Y0 mass ($MH^+$; z=3) calculated using the mass of core peptide |
| Y1 (z=1) *2 | Y1 mass ($MH^+$; z=1) calculated using the mass of core peptide |
| Y1 (z=2) *2 | Y1 mass ($MH^+$; z=2) calculated using the mass of core peptide |
| Y1 (z=3) *2 | Y1 mass ($MH^+$; z=3) calculated using the mass of core peptide |

*1 Red: information on glycopeptide clusters. These items are the identical to those in the "Cluster(details)" sheet (Table 5-5). Blue: information on core peptides and glycan compositions. These items are the identical to those in the "Peptide list" sheet (Table 4-1).

*2 These items will be indicated only when the "show peaks having MS2 spectrum" is "yes" in the setting (Table 3-5).

**Table 5-7. Details of inter-cluster analysis result sheet.**

| Item | Description |
|---|---|
| cluster_no* | Cluster No. |
| #member* | No. of members in the cluster |
| group_no* | Group No. for annotating each glycopeptide (C, cluster; G, group)<br>(*The peaks with the same mass are counted as one.) |
| peak_no* | Monoisotopic peak No. that is the same as "monoiso no" in the monoisotopic peak list. |
| charge* | Charge state |
| m/z* | m/z of the monoisotopic peak |
| rt* | RT (min) of the monoisotopic peak |
| intensity* | Signal intensity of the monoisotopic peak |
| #members | Non-redundant No. of the combination of reference and target peaks with a relationship specified by user. When the ratio of members is over the threshold specified in the setting (Figure 3-8), these peaks are considered as "related". |
| Delta m/z | Difference in masses (m/z) between reference and target peaks |

| Item | Description |
|---|---|
| Relation | The relationship between reference and target peaks, which is identified by comparing the "Delta m/z" with the value entered in the setting. |
| Direction | Direction of relationship that indicates which peak is a naked or adduct) |
| Delta Delta m/z | Delta value (Da) between the "Delta m/z" and the value entered in the setting.<br>When this value is lower the threshold specified in the setting, these peaks are considered as "related". |
| Delta rt | Delta value (min) of RTs between reference and target peaks. When this value is within the range specified in the setting, these peaks are considered as "related". |

*Column A-H indicate items for Reference cluster, whereas column I-P are for Target clusters examined whether it has a relationship specified by user.

**Table 5-8. Details of MS2 information sheets.**

| Item *1 | Description |
|---|---|
| cluster_no | Cluster No. |
| peak_no | Monoisotopic peak No. that is the same as "monoiso no" in the monoisotopic peak list. |
| charge | Charge state |
| m/z | Observed mass ($MH^+$) of the monoisotopic peak |
| rt | RT (min) of the monoisotopic peak |
| intensity | Signal intensity of the monoisotopic peak |
| no_member | Ascending numbering for monoisotopic peaks in the cluster |
| precursors | Ascending numbering for precursor ions for one monoisotopic peak |
| MH+calc | Calculated mass ($MH^+$) of the glycopeptide based on the mass of precursor ion and its charge state |
| deltaMH+(Da) | Delta value (Da) between observed and calculated masses ($MH^+$) of the monoisotopic peak |
| delta rt | Delta value (min) between RTs of a monoisotopic peak (MS1) and its corresponding MS2 spectra |
| scan | Scan No. of MS2 spectra annotated in the uploaded mgf file. Note that the link is inactive in this version. |
| rt(min) | RT (min) of MS2 spectra |
| charge | Charge state of the precursor ion |
| precursor mass | Mass ($MH^+$) of the precursor ion in the indicated charge state |
| Intensity | Signal intensity of the precursor ion |
| z[Y0/Y1] | Charge state of Y0 and Y1 ions |
| matches[Y0/Y1] | No. of matches for glycopeptide-derived ions |
| Y0 mass predict | Predicted mass (M) of Y0 ion |
| peptide[seq] | CP No. and sequence of the core peptide that has the mass (M) corresponding to the "Y0 mass predict". |

| Item *1 | Description |
|---|---|
| | (*Present only if there is a corresponding core peptide in the list) |
| peptide[mass] | Observed mass (M) of the core peptide |
| peptide-Y0 mass | Delta value (Da) between the “peptide[mass]” and “Y0 mass predict” |
| Y0 mass obs *2 | Observed mass of the Y0 ion |
| Y0 int *2 | Signal intensity of the Y0 ion |
| Y1-Y0 dif *2 | Delta value (Da) between masses of Y1 and Y0 ions |
| isotope count | Ascending numbering of isotopes |
| HexNAc(138) obs *3 | Observed mass of a diagnostic ion (specified by user) |
| HexNAc(138) dif *3 | Delta value (Da) between observed and calculated masses of a diagnostic ion (specified by user). When the delta value is lower than the value of “tolerance error for diag ion” value entered in the setting (Table 3-5), the observed ion will be considered as a diagnostic ion. |
| HexNAc(138) int *3 | Signal intensity of a diagnostic ion (specified by user) |
| used in cluster *4 | Cluster No. that includes a monoisotopic peak corresponding to the MS2 scan |
| precursor intensity *4 | Signal intensity of the precursor ion |

*1 Red: information on assigned clusters that are shown in the “Selection Results” sheet.

*2 Similar items are also indicated for other glycopeptide-related ions (Y0x, Y1, Y1F, and Y2).

*3 Similar items are also indicated for other diagnostic ions specified by user.

*4 Indicated only in the “MS2 info for clusters” sheet.
